# The first functional characterization of ancient interleukin-15-like (IL-15L) reveals shared and distinct functions of the IL-2, -15 and -15L family cytokines

**DOI:** 10.1101/644955

**Authors:** Takuya Yamaguchi, Axel Karger, Markus Keller, Eakapol Wangkahart, Tiehui Wang, Christopher J. Secombes, Azusa Kimoto, Mitsuru Furihata, Keiichiro Hashimoto, Uwe Fischer, Johannes M. Dijkstra

## Abstract

The ancient cytokine interleukin 15-like (IL-15L) was lost in humans and mice but not throughout mammals. This is the first study to describe IL-15L functions, namely in the fish rainbow trout. Fish have only one α-chain receptor gene *IL-15Rα*, whereas in mammalian evolution this gene duplicated and evolved into *IL-15Rα* plus *IL-2Rα*. Trout IL-2, IL-15 and IL-15L all could bind IL-15Rα and were able to induce phosphorylation of transcription factor STAT5. Reminiscent of the mammalian situation, trout IL-15 was more dependent on “in *trans*” presentation by IL-15Rα than IL-2. However, whereas trout IL-15 could also function as a free cytokine as known for mammalian IL-15, trout IL-15L function showed a total dependency on in *trans* presentation by IL-15Rα. Trout lymphocytes from the mucosal tissues gill and intestine were sensitive to IL-15, but refractory to IL-2 and IL-15L, which is reminiscent of sensitivities to IL-15 in mammals. Distinguishing engagement of the IL-2Rα/IL-15Rα receptor chain may explain why IL-2 and IL-15 were selected in evolution as major growth factors for regulatory T cells and lymphocytes in mucosal tissues, respectively. Trout IL-15L efficiently induced expression of *IL-4* and *IL-13* homologues in CD4^-^CD8^-^IgM^-^ splenocytes, and we speculate that the responsive cells within that population were type 2 innate lymphoid cells (ILC2). In contrast, trout IL-15 efficiently induced expression of *interferon γ* and *perforin* in CD4^-^CD8^-^IgM^-^ splenocytes, and we speculate that in this case the responsive cells were natural killer (NK) cells. In fish, in apparent absence of IL-25, IL-33 and TSLP, primitive IL-15L may have an important role early in the type 2 immunity cytokine cascade. Among trout thymocytes, only CD4^-^CD8^-^ thymocytes were sensitive to IL-15L, and different than in mammals the CD4^+^CD8^+^ thymocytes were quite sensitive to IL-2. In addition, the present study provides (i) the first molecular evidence for inter-species cytokine with receptor chain interaction across fish-mammal borders, and (ii) suggestive evidence for a tendency of IL-2/15/15L cytokines to form homodimers as an ancient family trait. This is the first comprehensive study on IL-2/15/15L functions in fish and it provides important insights into the evolution of this cytokine family.

## Introduction

Interleukin 2 (IL-2) was one of the first cytokines to be detected. This was due to the remarkable power of IL-2 to induce and sustain T lymphocyte proliferation *in vitro*, and IL-2 was originally named “T cell growth factor” (TCGF) (1–3). Many years later, IL-15, a molecule closely related to IL-2, was discovered (4), and it took even longer to realize that IL-15 was especially potent/stable in combination with its “heterodimer partner” IL-15Rα (5–7). Nowadays, recombinant IL-2 and IL-15 (with or without IL-15Rα), or antibodies blocking their action, provide important tools for *in vitro* culturing of lymphocytes and for treating disease in the clinic or in preclinical models [reviewed in (8)]. The mammalian IL-2 versus IL-15 situation is not fully understood yet, and analysis of this cytokine family in non-mammalian species may provide additional insights.

*IL-15-like* (*IL-15L*) gene was originally discovered in teleost fish (9–11), but later an intact gene was also discovered in cartilaginous fish (12), reptiles, non-eutherian mammals, and some eutherian mammals including cattle, horse, pig, cat, mouse lemur, rabbit and hedgehog (13). In rodents and higher primates only remnants of *IL-15L* were found, and IL-15L function is not expected in those species (13).

The cytokines IL-2, IL-15, and IL-15L are close relatives within a larger subfamily of cytokines that also includes IL-4, IL-7, IL-9, IL-13, IL-21 and thymic stromal lymphopoietin (TSLP), most of which bind receptors that contain an IL-2Rγ chain (aka “common cytokine-receptor γ-chain” or “γ_c_”) (13–15).

The following describes the IL-2 and IL-15 functions as discovered for mammals. IL-2 and IL-15 signal through the heterodimer type I receptor IL-2Rβ·IL-2Rγ and can induce very similar transcription profiles (16). Both IL-2 and IL-15 importantly activate transcription factor STAT5 (15, 17). Whereas free IL-2 and IL-15 molecules can bind with low efficiency to IL-2Rβ·IL-2Rγ heterodimers, the cytokine-specific and efficient receptor complexes are formed by the heterotrimers IL-2Rα·IL-2Rβ·IL-2Rγ and IL-15Rα·IL-2Rβ·IL-2Rγ, respectively (16, 18–21). The IL-2Rα and IL-15Rα chains do not belong to the type I receptor chain family, but important parts of their ectodomains belong to the complement control protein (CCP) domain family (aka “sushi” or “SRC” domains). IL-2 is secreted predominantly by activated T cells, while IL-2Rα is abundant constitutively on the surface of regulatory T cells (T_regs_) and enhanced on several leukocyte populations after their activation, most notably on effector T cells (22–24). IL-2 interacts primarily in free secreted form with membrane-bound IL-2Rα·IL-2Rβ·IL-2Rγ complexes, and in this situation the IL-2Rα chain is said to be provided “in *cis*”. IL-2 secretion by activated T cells forms part of a self-stimulatory loop for these cells, but also provides a negative feedback loop through the stimulation of T_regs_ (25). In contrast to IL-2, the IL-15 protein is predominantly expressed together with IL-15Rα by antigen presenting cells such as monocytes and dendritic cells (23). Membrane-bound or shed/secreted IL-15·IL-15Rα complexes can stimulate other cells that express IL-2Rβ·IL-2Rγ, and in this situation the IL-15Rα chain is said to be provided “in *trans*” (26–28). The IL-15 to IL-15Rα binding mode is characterized by unusually high affinity in the picomolar range (5, 20), and, although in experiments IL-15 was shown to be able to function as an independent secreted cytokine, it was calculated that in human serum all IL-15 may be bound to soluble forms of IL-15Rα (28). Both IL-2 and IL-15 can stimulate a variety of lymphocytes, but whereas a dominant effect of IL-2 concerns the above mentioned T_reg_ stimulation (29, 30), IL-15 is particularly important for stimulation of natural killer (NK) cells, intra-epithelial lymphocytes (IELs), and CD8^+^ T cells (7, 31, 32).

Mammalian *IL-2Rα* and *IL-15Rα* genes were derived from a gene duplication event (21), probably early in tetrapod species evolution from an *IL-15Rα* type gene, after which the IL-2Rα to IL-2 binding mode substantially diverged (13). In contrast, sequence comparisons suggest that the binding mode of IL-15 and IL-15L to IL-15Rα, as elucidated for mammalian IL-15 (33, 34), did not change during evolution of jawed vertebrates (13). In teleost (modern bony) fish, consistent with sequence motif conservation (13), and in the absence of an IL-2Rα molecule (13, 35), both IL-2 and IL-15 were found to bind with IL-15Rα, although IL-15 with a higher affinity (35).

In teleost fish, the *IL-2* and *IL-15* loci are well conserved (13), and some studies have been done on the recombinant cytokines [reviewed in (36)]. Importantly, reminiscent of the proliferation functions in mammals, rainbow trout IL-2 and IL-15 in the supernatants of transfected cells were both able to sustain long term culturing of lymphocytes from trout head kidney (a fish lymphoid organ) that expressed markers of CD4^+^ T cells (37, 38).

Hitherto, the only functional property determined for IL-15L was its interaction with IL-15Rα, which we showed using recombinant bovine proteins (13). In contrast to the situation in mammals, bona fide *IL-15L* genes are well conserved throughout fishes (13), so we speculated that fish IL-15L might have a more robust and easier to identify function. In the present study, we started with analyses of both rainbow trout and bovine IL-2, IL-15 and IL-15L, after which we concentrated on the rainbow trout model because only for that species we detected IL-15L function. Functions of the recombinant trout cytokines were investigated using both supernatants of transfected mammalian cells and isolated proteins after expression in insect cells. Comparisons between rainbow trout IL-2, IL-15 and IL-15L functions, and their different dependencies on IL-15Rα, revealed ancient similarities of this cytokine system with the mammalian situation. Unexpected were the very different, and even opposing, immune effects that rainbow trout IL-15 and IL-15L could have on some lymphocyte populations.

## Results

### Identification, expression analysis, and sequence comparisons of rainbow trout IL-15La and -b

Two rainbow trout *IL-15L* genes, *Il-15La* and *IL-15Lb*, could be identified in genomic sequence databases (Fig. 1) and were amplified from cDNA (Figs. S1A and -B). They map to the rainbow trout reference genome chromosomes 27 and 24, which have been recognized as a pair of chromosomes sharing ohnologous regions derived from a whole genome duplication early in the evolution of salmonid fishes (39). By 5’-RACE analysis and database comparisons a number of AUG triplets in 5’ untranslated regions (5’UTRs) of both trout *IL-15La* and *IL-15Lb* were found (Figs. S1A and -B, and Table 1), as reported for *IL-15L* of other fish species (9, 11) (Table 1), for mammalian *IL-15L* (13) (Table 1) and for fish and mammalian *IL-15* (9, 10, 40–42). These additional *AUG* triplets suggest that efficient translation may need some special conditions and that the transcript amounts may not be directly representative of the protein amounts (40, 41). *IL-15La* was found constitutively expressed in many tissues of healthy trout, whereas *IL-15Lb* showed a more restricted expression pattern (Figs. 2 and S1C). Fig. 2 and Fig. S1C-1 show our experimental RT-qPCR and semi-quantitative RT-PCR data, respectively, while Fig. S1C-2 shows the relative numbers of matches in tissue-specific single read archive (SRA) datasets of the NCBI database. Despite variation between trout individuals, rather consistent findings were that trout *IL-15Lb* expression was relatively high in gill, and both trout *IL-15La* and *IL-15Lb* expression were relatively low in head kidney (Figs. 2 and S1C). In genomic sequence databases of a related salmonid fish, Atlantic salmon (*Salmo salar*), *IL-15La* and *IL-15Lb* could also be found (Fig. 1), and comparison of these sequences with tissue-specific RNA-based SRA datasets indicated that *IL-15La* and *IL-15Lb* expression in Atlantic salmon agree with the above summary for trout (Fig. S1C-2). Fig. 2 shows that *IL-15La* transcripts were also found in trout macrophages, and epithelial and fibroblast cell lines.

**Fig. 1.**
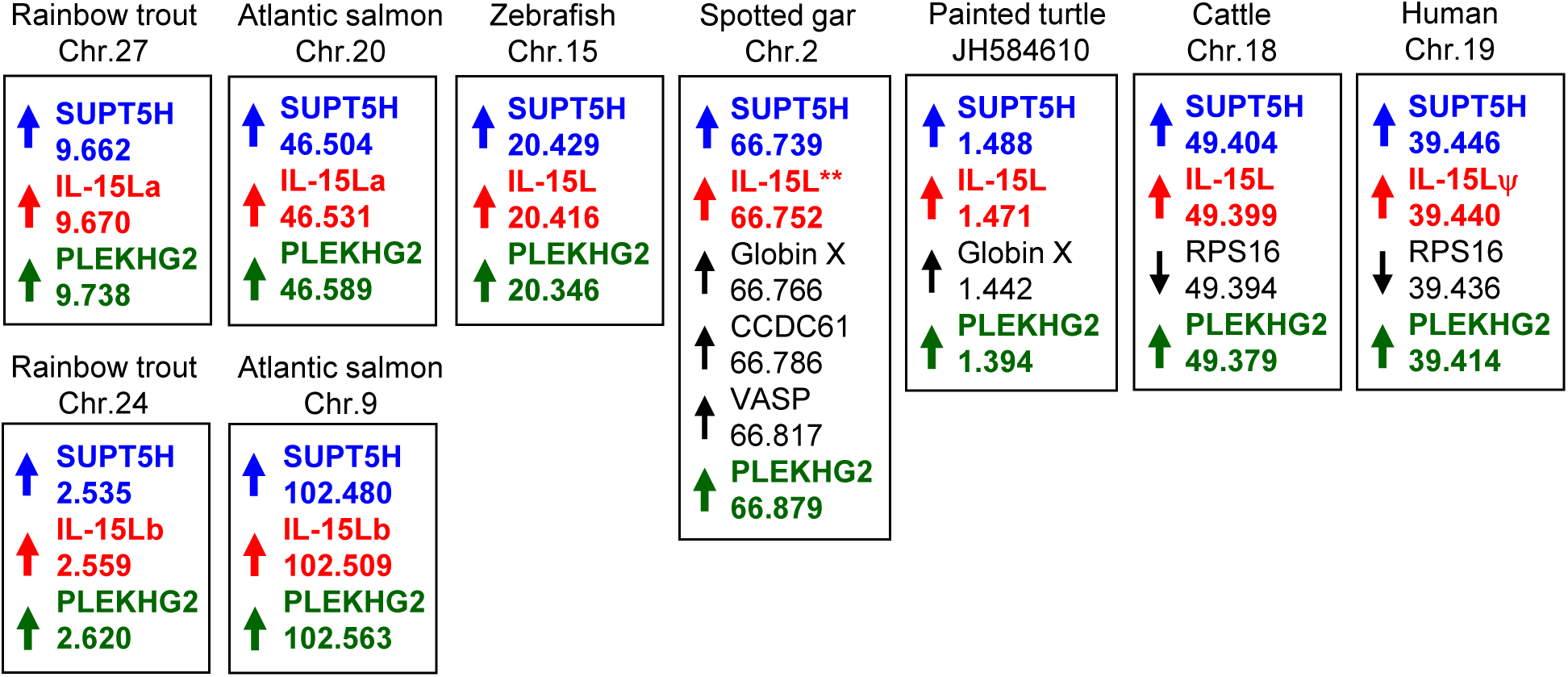
Genomic location of *IL-15L* gene. In rainbow trout, Atlantic salmon (*Salmo salar*), zebrafish (*Danio rerio*), spotted gar (*Lepisosteus oculatus*), painted turtle (*Chrysemys picta*), cattle, and human, *IL-15L* loci are found close to suppressor of *Ty 5 homolog* (*SUPT5H*) and *pleckstrin homology domain containing family G member 2* (*PLEKHG2*) genes. Human *IL-15L* is a pseudogene (IL-15Lψ) (13). Rainbow trout *IL-15La* and *IL-15Lb* were found in the rainbow trout reference genome sequence (NCBI database Omyk_1.0) chromosomes 27 and 24, respectively. Atlantic salmon *IL-15La* and *IL-15Lb* were found in the Atlantic salmon reference genome sequence (NCBI database ICSASG_v2) chromosomes 20 and 9, respectively. *For spotted gar, full-length *IL-15L* consensus gene sequence could not be found in the determined genomic sequence (13), but full-length cDNA information is available in the NCBI TSA database (see Fig. 3). Depicted zebrafish, spotted gar, turtle, cattle and human data are based on (Pre-) Ensembl datasets GRC.z10, LepOcu1, ChrPicBel3.0.1, UMB 3.1 and GRCh38.p12, respectively. Arrows indicate gene orientations, and numbers indicate the nucleotide positions in megabase of probable ORF start codons (for all genes in trout, salmon, turtle and human, the *IL-15L* genes in the other species, and *PLEKHG2* in spotted gar) or the gene 5’ ends according to annotation in the Ensembl database. Orientation between scaffolds was adapted to match gene orientations.

**Fig. 2.**
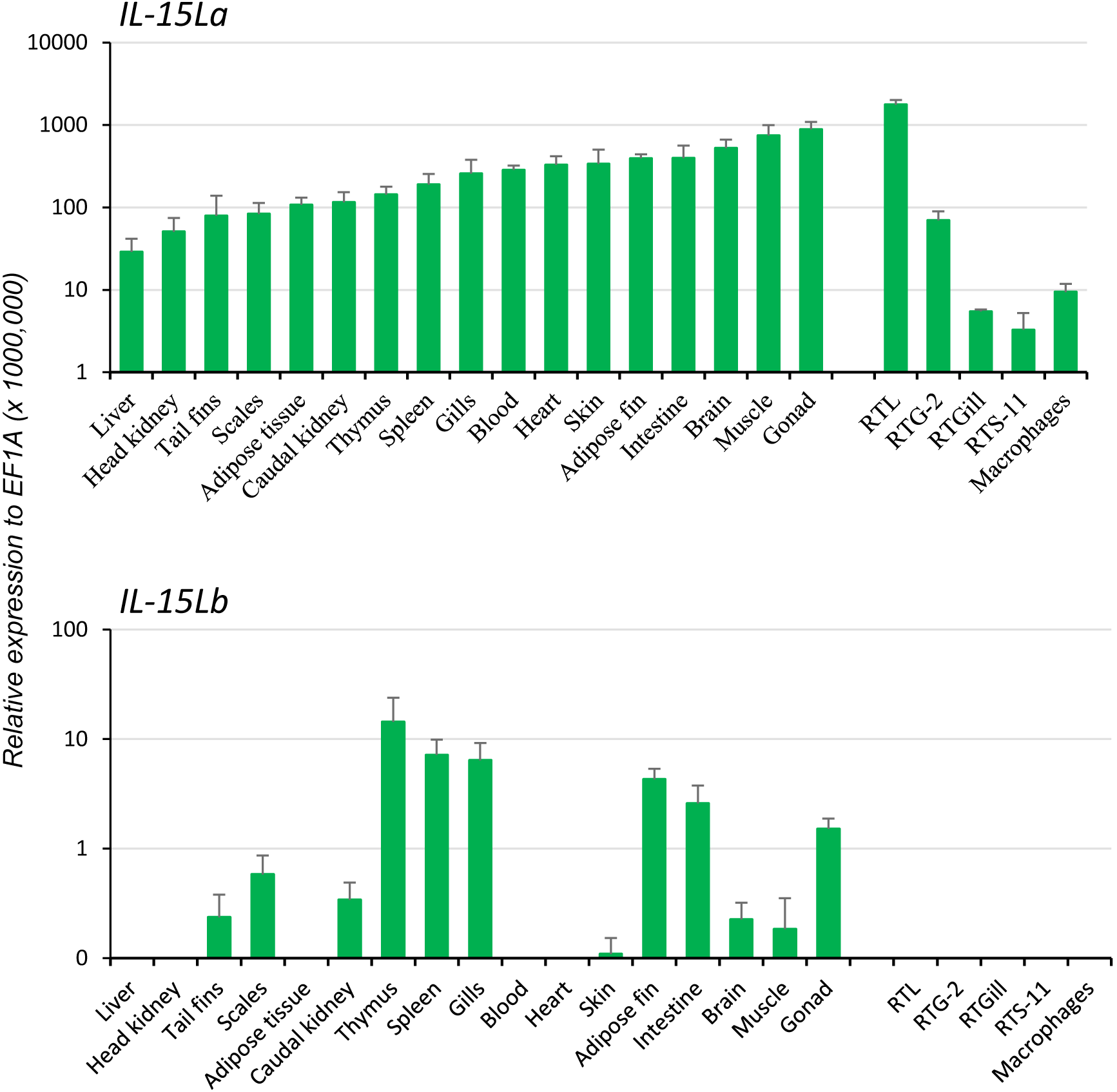
Expression of *IL-15La* and *IL-15Lb* in rainbow trout tissues and cells. Seventeen tissues from six rainbow trout individuals were sampled: blood, tail fins, scales, skin, muscle, adipose fin, thymus, gills, brain, adipose tissues, spleen, liver, heart, gonad, head kidney (HK), caudal kidney and intestine. The relative expression of each *IL-15L* gene was normalized against the expression level of the housekeeping gene *EF1A*. The expression levels in four trout cell lines — a monocyte/macrophage-like cell line RTS-11 from spleen, an epithelial cell line RTL from liver, and fibroid cell lines RTG-2 from gonad and RTGill from gills — and in primary HK macrophages were determined in a similar way. The figure shows the means+SEM values, with n=6 for the trout tissues and n=4 for the cell lines and primary macrophage cultures.

**Table. 1.**
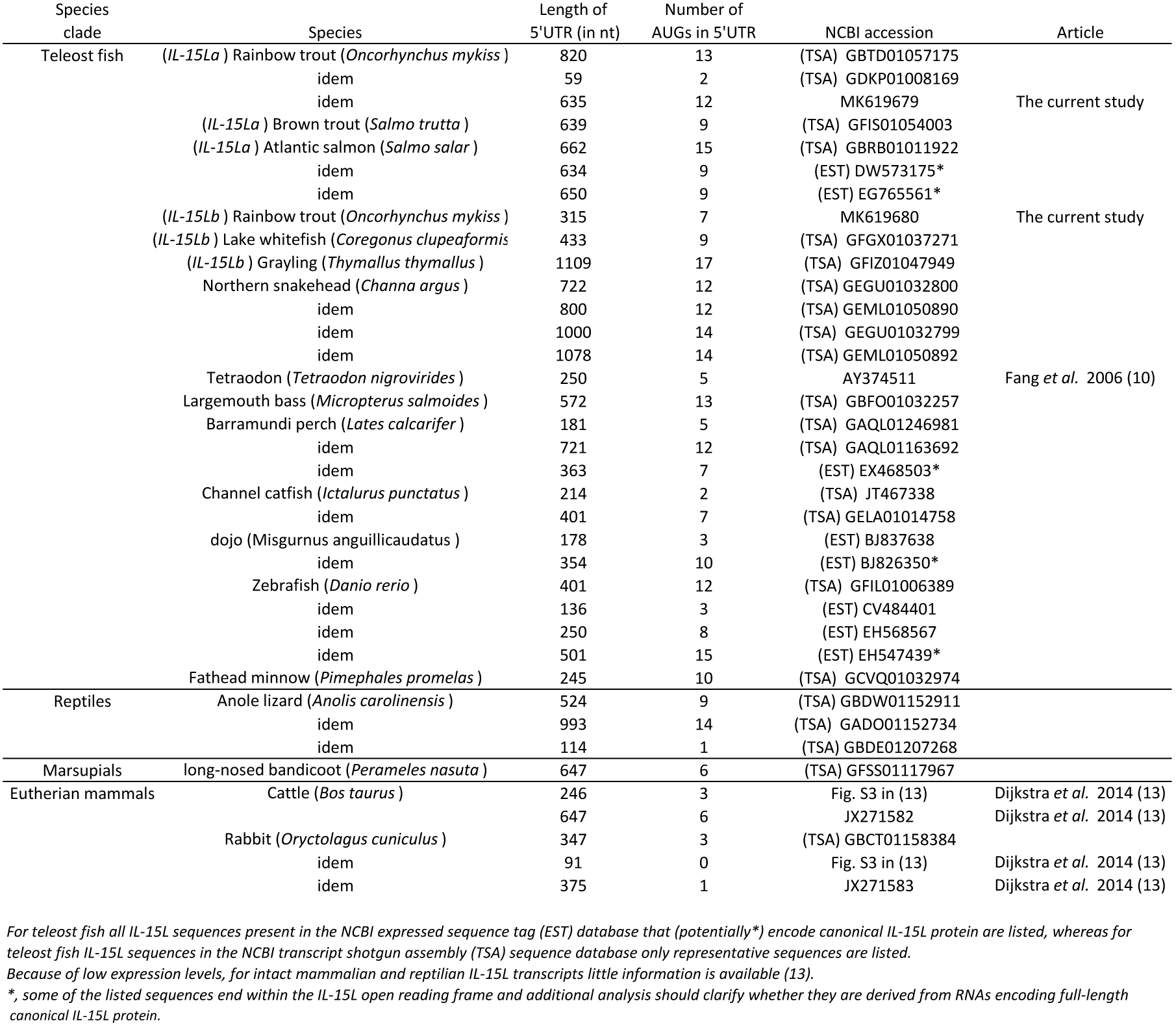
Number of AUG codons in the 5'UTR of reported *IL-15L* transcript sequences.

The deduced amino acid sequences of trout IL-15La and IL-15Lb are aligned in Fig. 3 together with related cytokines. Residues that are rather typical for IL-15L (13) are shaded green. Phylogenetic tree analysis comparing these highly diverged cytokines does not provide conclusive information on their evolution (13), but when such analysis is performed on only the IL-15L sequences the result (Fig. S1D) is consistent with the location-based assumption (see above) that rainbow trout *IL-15La* and *IL-15Lb* are paralogues which were generated by the whole genome duplication in an ancestor of salmonids (39). As we discussed previously (13), although the conservation of the overall sequences is poor, residues of mammalian IL-15 that are known to bind IL-15Rα are well conserved throughout IL-15, IL-15L, and fish IL-2. In Fig. 3, blue and pink shading mark residues of binding patches 1 and 2 determined for mammalian IL-15 to IL-15Rα binding (33, 34), with the most important residues (34) indicated with a circle above. Residues of mammalian IL-2, IL-15, and IL-4 which are known to be of major importance for interaction with their respective type I receptors (16, 43–47) are shaded red in Fig. 3, and so are residues of the other cytokines for which a similar importance may be expected (13). Although some of the alignments of the highly diverged α-helix A and C regions in Fig. 3 are quite speculative [for a better discussion of the alignment see (13)], among IL-15L sequences an acidic residue (D/E) in α-helix A, an arginine in α-helix C, and a glutamine or glutamic acid (E/Q) in α-helix D that may participate in type I receptor binding are rather well conserved.

**Fig. 3.**
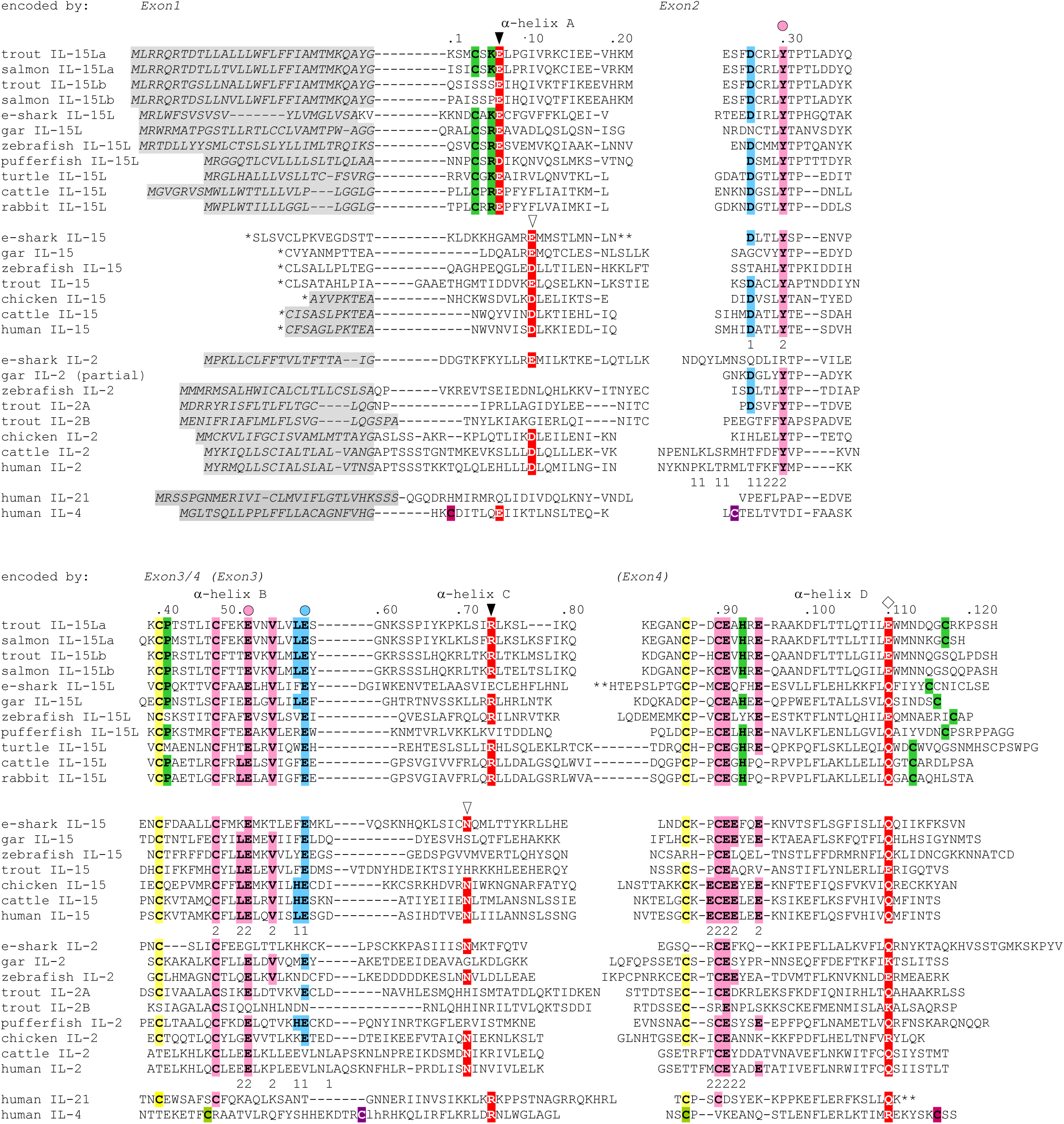
Alignment of deduced rainbow trout IL-15La and IL-15Lb with other IL-15L, IL-15 and IL-2 amino acid sequences. Identical colored shading of cysteines refers to known or expected disulfide bridges. Residues rather characteristic for IL-15L (13), including a cysteine pair, are shaded green. In the interaction of mammalian IL-15 and IL-2 with their respective receptor chains IL-15Rα or IL-2Rα, two binding patches “1” and “2” were distinguished, with patch 2 quite similar and patch 1 quite different between IL-15·IL-15Rα and IL-2·IL-2Rα (33, 34); the participating residues in human IL-15 and IL-2 are indicated with the respective number 1 or 2 below them. To highlight the conservation of the IL-15 to IL-15Rα binding mode, residues in the alignment that are identical to the human or murine IL-15 patch 1 residues are shaded blue, and residues that are identical to the human or murine IL-15 patch 2 are shaded pink; the most important residues for IL-15 to IL-15Rα binding (34) are indicated by a colored circle above the alignment. Residues which are known or are expected to be of importance for interaction with type I receptors are shaded red (13, 16, 43–47); the open triangles indicate residues important for interaction of mammalian IL-15 and IL-2 with the IL-2Rβ chain, the closed triangles indicate positions which in IL-4 are important for interaction with IL-4Rα chain, and the diamonds are indicated above a glutamine or arginine which in mammalian IL-2, IL-15 or IL-4 is important for binding IL-2Rγ chain (16, 45–47). *, for IL-15 sequences the leader peptide amino acid sequences encoded by exons upstream from family consensus exon 1 are not shown. **, for elephant shark IL-15L, elephant shark IL-15 and human IL-21 a stretch is not shown for lay-out reasons. Residue numbering follows trout IL-15La. Italic font and gray shading, (predicted) leader peptides. Gaps, open spaces relate to exon borders whereas hyphens connect residues encoded by the same exon. For comparisons with additional cytokines see reference (13). Names of species are: trout, rainbow trout (*Oncorhynchus mykiss*); salmon, Atlantic salmon (*Salmo salar*); e-shark, elephant shark (*Callorhinchus milii*); gar, spotted gar (*Lepisosteus oculatus*); zebrafish (*Danio rerio*); pufferfish, green spotted pufferfish (*Tetraodon nigroviridis*); turtle, painted turtle (*Chrysemys picta*); cattle (*Bos taurus*); human (*Homo sapiens*). Database accessions for the sequences are [for the sequences also see references (1, 10)]: e-shark IL-15L, GenBank KA353649; gar IL-15L, GenBank GFIM01029449; zebrafish IL-15L, GenBank NP_001009558; rainbow trout IL-15La, GenBank MK619679; rainbow trout IL-15Lb, GenBank MK619680; Atlantic salmon IL-15La, GenBank GBRB01011922; Atlantic salmon IL-15Lb, predicted from GenBank AGKD04000049; pufferfish IL-15L, predicted from Ensembl “TETRAODON8” and described by Fang et al. 2006; turtle IL-15L, GenBank XP_008171403; cattle IL-15L, NP_001288142; rabbit IL-15L, NP_001288189; e-shark IL-15, JW878023; gar IL-15, Ensembl “LepOcu1”; zebrafish IL-15, GenBank AAZ43090; trout IL-15, GenBank AJ555868; chicken IL-15, GenBank AAD38392; cattle IL-15, AAA85130; human IL-15, AAA21551; e-shark IL-2 (alias IL-2-like), predicted from the elephant genome project sequence which has GenBank accession AAVX02000000; gar IL-2, predicted from Ensembl “LepOcu1”; zebrafish IL-2, predicted from Ensembl “Zv9”; trout IL-2A, GenBank NM_001164065; trout IL-2B, GenBank HE805273; chicken IL-2, GenBank AAC96064; cattle IL-2, GenBank AAA30586; human IL-2, GenBank 0904306A; human IL-21, Genbank AAG29348; human IL-4, GenBank AAA59149. For comparisons with additional cytokines see (13).

Very recently, it was described that rainbow trout has two quite different IL-2 molecules, IL-2A and IL-2B, which have overlapping but distinct functions (48). Fig. 3 shows that compared to fish IL-2 consensus the rainbow trout IL-2B molecule lost cysteines and some residues for IL-15Rα binding, a topic for future studies. In the present study, we only analyze rainbow trout IL-2A, which for simplicity we call “IL-2”. Since trout IL-15Lb lost an IL-15L consensus cysteine pair (the green shaded cysteines in Fig. 3), we speculated that trout IL-15La function would be more representative of canonical IL-15L function, and therefore most research in the present study was dedicated to this protein version.

### Trout IL-15La can be N-glycosylated

Trout IL-15La has a single N-glycosylation motif [NxS/T (49)] at position 61 (Fig. 3). Human HEK293T cells were transfected with DNA plasmid expression vectors encoding FLAG-tagged versions of bovine IL-2, IL-15 and IL-15L, and trout IL-2, IL-15, IL-15La and IL-15Lb (for sequences of expression vectors see Fig. S2). After 24 h, cell lysates were prepared and treated with PNGase-F or without (mock treatment), and then the samples were subjected to anti-FLAG Western blot analysis. This revealed shifts in apparent molecular weight indicative of N-glycosylation for most investigated cytokines but not for bovine IL-15L and trout IL-15Lb (Figs. 4 and S5A). The results are consistent with these latter two cytokines not having an N-glycosylation motif (Fig. 3).

**Fig. 4.**
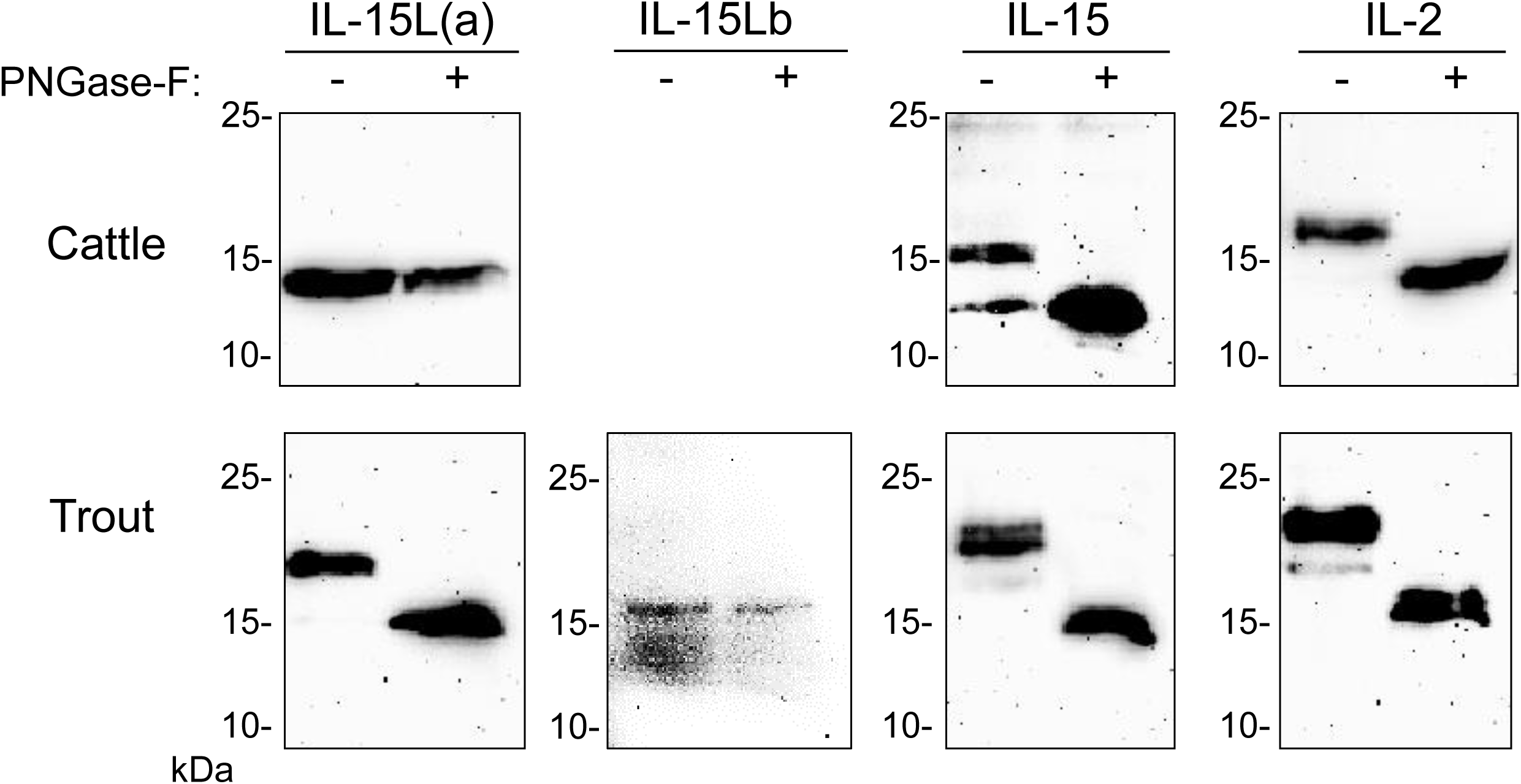
Bovine IL-2 and IL-15, and rainbow trout IL-2, IL-15 and IL-15La in transfected HEK293T cells are N-glycosylated. Lysates of HEK293T cells transfected for the indicated bovine or trout cytokine were digested with PNGase-F (+) or mock treated (-) and analyzed by Western blotting using anti-FLAG mAb. All analyzed molecules except bovine IL-15L and trout IL-15Lb showed a shift in apparent molecular weight consistent with removal of N-linked oligosaccharides. For full-size blots and additional controls see Fig. S5A.

### Cross-reactivities between trout and bovine cytokines and IL-15Rα

Previously, we showed by a combination of DNA plasmid transfection and anti-FLAG flow cytometry experiments that FLAG-tagged bovine IL-15L could be found on the surface of HEK293T cells if they were co-transfected for bovine IL-15Rα but not if co-transfected for bovine IL-2Rα (13). In the present study we repeated this analysis, but in addition included recombinant expression of trout IL-15Rα, and of FLAG-tagged bovine IL-2 and IL-15, and trout IL-2, IL-15, IL-15La and IL-15Lb. The results of representative experiments are shown in Fig. 5, while Table S1 summarizes the results of all experiment repeats that were done. The interaction between bovine IL-2Rα chain and bovine IL-2 was mutually specific (Fig. 5A). Bovine IL-15 and IL-15L, and trout IL-2, IL-15, IL-15La and IL-15Lb, could only be detected, or were detected at higher amounts, at the cell surface if the cells were co-transfected for either bovine or trout IL-15Rα (Figs. 5B and 5C). The cross-species interactions appeared to be especially efficient for bovine IL-15Rα co-expressed with trout IL-2, IL-15 and IL-15L (Fig. 5B), but were also observed for trout IL-15Rα co-expressed with bovine IL-15 and IL-15L (Fig. 5C). For unknown reasons, recombinant bovine IL-15 (in which the leader sequence had been replaced for that of IL-2; see Fig. S2) was also detectable at the cell surface in the absence of co-transfected receptor chains (Figs. 5-Ab, -Bb and -Cb).

**Fig. 5.**
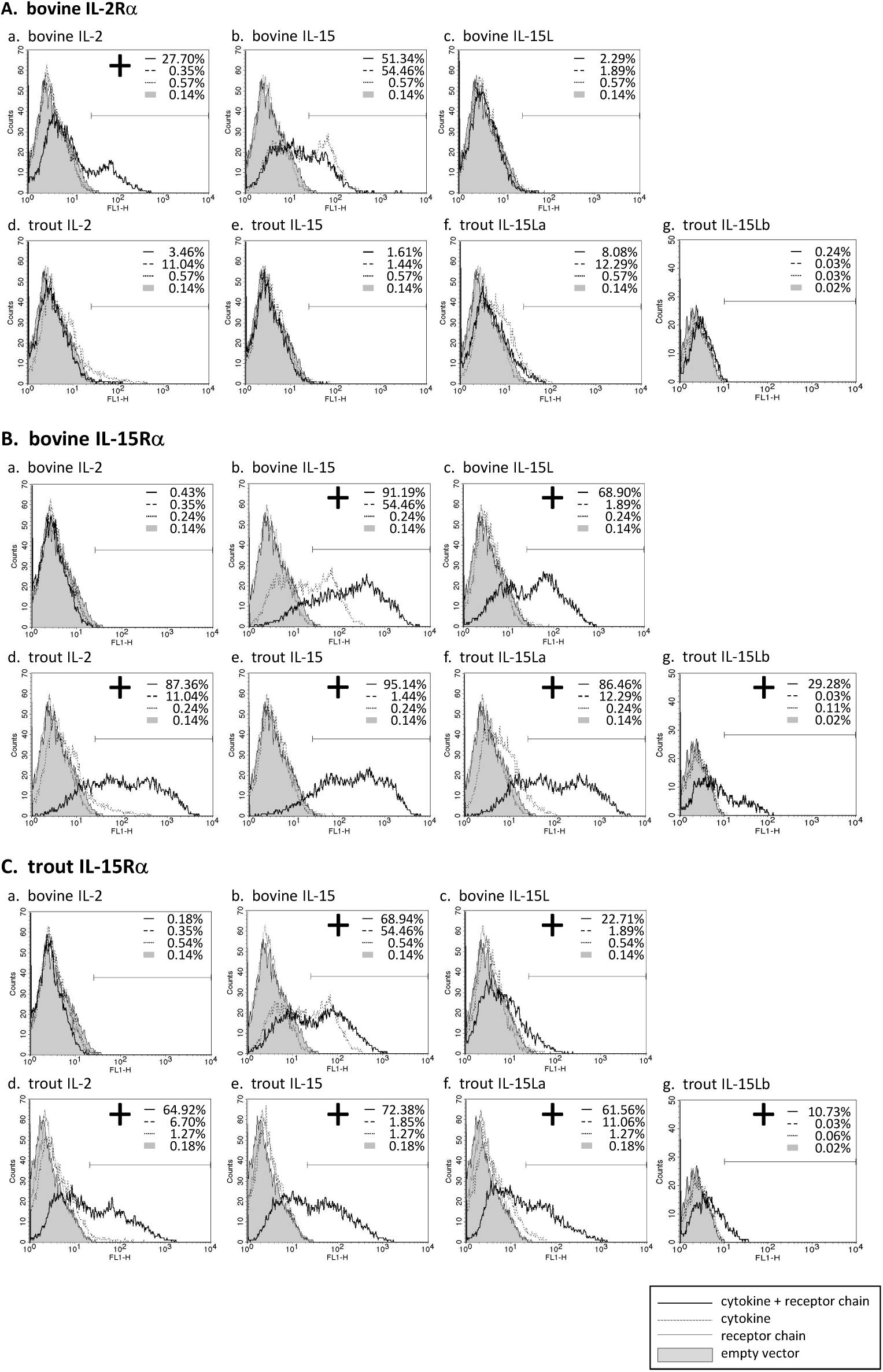
Cell surface presentation of bovine and rainbow trout IL-2, IL-15 and IL-15L by IL-2Rα or IL-15Rα of those species. Live transfected human HEK293T cells were analyzed by flow cytometry using anti-FLAG mAb to detect the presence of the FLAG-tagged cytokines on the cell surface. The percentages of anti-FLAG-labeled cells were compared between cells transfected for cytokine alone (dashed line), cells co-transfected for cytokine plus receptor chain (bold line), cells transfected for the receptor chain alone (dotted line) and cells transfected with empty vector (gray). Bovine IL-2Rα (A), bovine IL-15Rα (B) and trout IL-15Rα (C) were tested in combination with each cytokine (a-g). If the percentage of anti-FLAG-labeled cells was higher in the cells co-transfected for the receptor chain than in the cells transfected for the cytokine alone, the cytokine was considered to be bound to the receptor chain (shown as “+”). For a summary of experiment repeat results see Table S1.

### Dependency on soluble IL-15Rα for efficient stable secretion of bovine and trout IL-15 and IL-15L by transfected HEK293T cells

Previously, we found that recombinant bovine IL-15L could only be found in the supernatant of transfected cells if co-transfected for soluble IL-15Rα (sIL-15Rα) (13). In the present study that research was extended by also investigating the effect of co-transfection for species-specific sIL-15Rα on the stable secretion of bovine IL-15 and of trout IL-2, IL-15 and IL-15L, and that of co-transfection for bovine sIL-2Rα on bovine IL-2. Figs. 6A+6B, and 6C+6D, show the Western blot results of representative experiments in which the bovine and trout molecules were expressed, respectively, comparing the cytokines present in the supernatant (Figs. 6A and 6C) to those present in the matching cell lysates (Figs. 6B and 6D). Fig. S5B shows experiment repeats and the uncropped blot results, and in addition includes the Western blot analyses for detection of the receptor chains. The data in Figs. 6 and S5B consistently indicate that bovine and trout IL-15 and IL-15L are dependent for their abundance in the supernatant on the co-expression with, or fusion to (Fig. S5B), sIL-15Rα. As reported before (13), no IL-15L was detectable in supernatants of cells transfected for bovine IL-15L alone (Fig. 6A). When transfected for only trout IL-15La, small amounts of the cytokines could be detected in the supernatant, but these increased markedly upon co-transfection for, or genetic fusion to, sIL-15Rα (Figs. 6C and S5B). Trout IL-15Lb was consistently found in lower amounts than the other cytokines, even in the transfected cell lysates, especially in the absence of sIL-15Rα (Fig. 6D) which seems to have a stabilizing role and to be necessary for finding any trout IL-15Lb in the cell supernatant (Fig. 6C). Similar to IL-15L, the presence of bovine and trout IL-15 in the supernatant was considerably boosted by the co-transfection for sIL-15Rα (Figs. 6A and 6C). Stable secretion of IL-2 of cattle and trout did not depend on receptor chain co-expression (Figs. 6A and 6C), and especially bovine IL-2 was efficiently released from the cells (compare Fig. 6A with Fig. 6B). In Fig. 6, although somewhat arbitrarily, very efficient secretion, intermediate efficient secretion, and poor secretion are highlighted with arrows, estimated from comparison of Fig. 6A with 6B, and of Fig. 6C with 6D. Whether the increased amounts of IL-15 and IL-15L in the supernatants in the presence of sIL-15Rα were caused by enhanced secretion, improved stability, or by both, needs further investigation. We interpret the band of ∼37 kDa observed for the cell lysate samples containing trout IL-15La as a possible IL-15La homodimer (Fig. 6D); similar sized trout IL-15La protein complexes can also be seen in additional Western blot figures in Figs. S5A and S5B, and were also observed for purified IL-2 (see below).

**Fig. 6.**
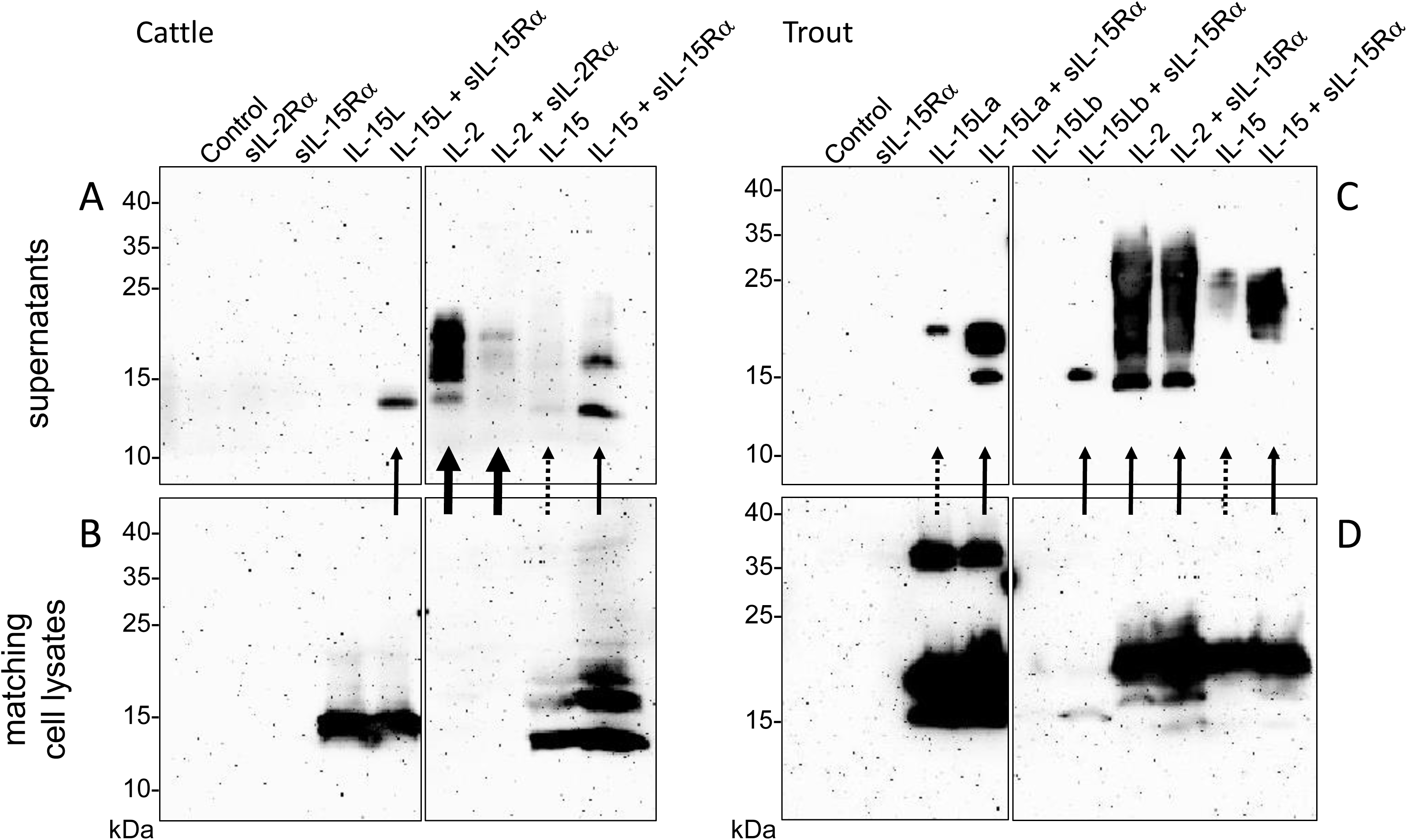
Co-expression with soluble IL-15Rα enhances presence of bovine and trout IL-15 and IL-15L in the supernatant of transfected cells. HEK293T cells were transfected for FLAG-tagged cytokines and/or species-specific Myc-tagged soluble IL-2Rα or IL-15Rα, and the matching supernatants (A, cattle; C, trout) and cell lysates (B, cattle; D, trout) were compared by Western blot analysis using anti-FLAG mAb. Thick, thin and dashed arrows highlight very efficient secretion, intermediate efficient secretion, and poor secretion, respectively, as they are deduced from comparison between the supernatant and cell lysate results. For experiment repeats and additional information see Fig. S5B.

### Trout IL-2, IL-15 and IL-15L in supernatants of transfected HEK293T cells induce STAT5 phosphorylation in distinct lymphocyte populations; trout IL-15La and IL-15Lb stimulate CD4^-^ CD8^-^ (double negative, DN) thymocytes

Preliminary experiments in which total leukocytes of different rainbow trout tissues were stimulated with cytokine-containing supernatants of transfected cells did not reveal induction of phosphorylated STAT5 (pSTAT5) by IL-15L. Therefore, we tried to increase the sensitivity by first sorting the CD8α-positive and -negative (CD8^+^ and CD8^-^) morphological lymphocyte fractions (FSC^low^/SSC^low^ in flow cytometry; mostly called “lymphocytes” from here) using an established monoclonal antibody (50) (Fig. S3). These cells were incubated with supernatants of HEK293T cells transfected for trout cytokines and/or for trout sIL-15Rα, or with control supernatant, and then pSTAT5 amounts were compared by Western blot analysis. Results are shown in Figs. 7 and S5C. In several experiments, but not consistently in all experiments, non-tagged versions of the cytokines were included [named IL-2(N), IL-15(N), IL-15La(N) and IL-15Lb(N)], to exclude the possibility that a FLAG-tag effect was responsible for the experimental outcome. Trout IL-2 efficiently stimulated both CD8^+^ and CD8^-^ lymphocytes of the systemic lymphoid tissues spleen and head kidney, and also of the thymus (Fig. 7). However, IL-2 was not found to stimulate lymphocytes from gill, and only had a weak stimulatory effect on CD8^+^ and CD8^-^ populations isolated from intestine (highlighted by blue bars in Fig. 7). That IL-15 was more efficient than IL-2 in the stimulation of lymphocytes from intestine and gill was evident because the induced pSTAT5 amounts were higher while the amounts of recombinant cytokine used for stimulation were smaller (Fig. S5C). Trout IL-15La and IL-15Lb containing supernatants did not detectably induce pSTAT5 in any of the investigated cell populations, except for CD8^-^ thymocytes (Figs. 7 and S5C; highlighted by red bars in Fig. 7; IL-15Lb data are only shown in Fig. S5C). The stimulation by IL-15La and IL-15Lb appeared to be fully dependent on the co-presence of sIL-15Rα (Figs. 7 and S5C), although it should be realized that in absence of sIL-15Rα the concentrations of IL-15La and IL-15Lb in the supernatant were very low or absent (see Fig. 6).

**Fig. 7.**
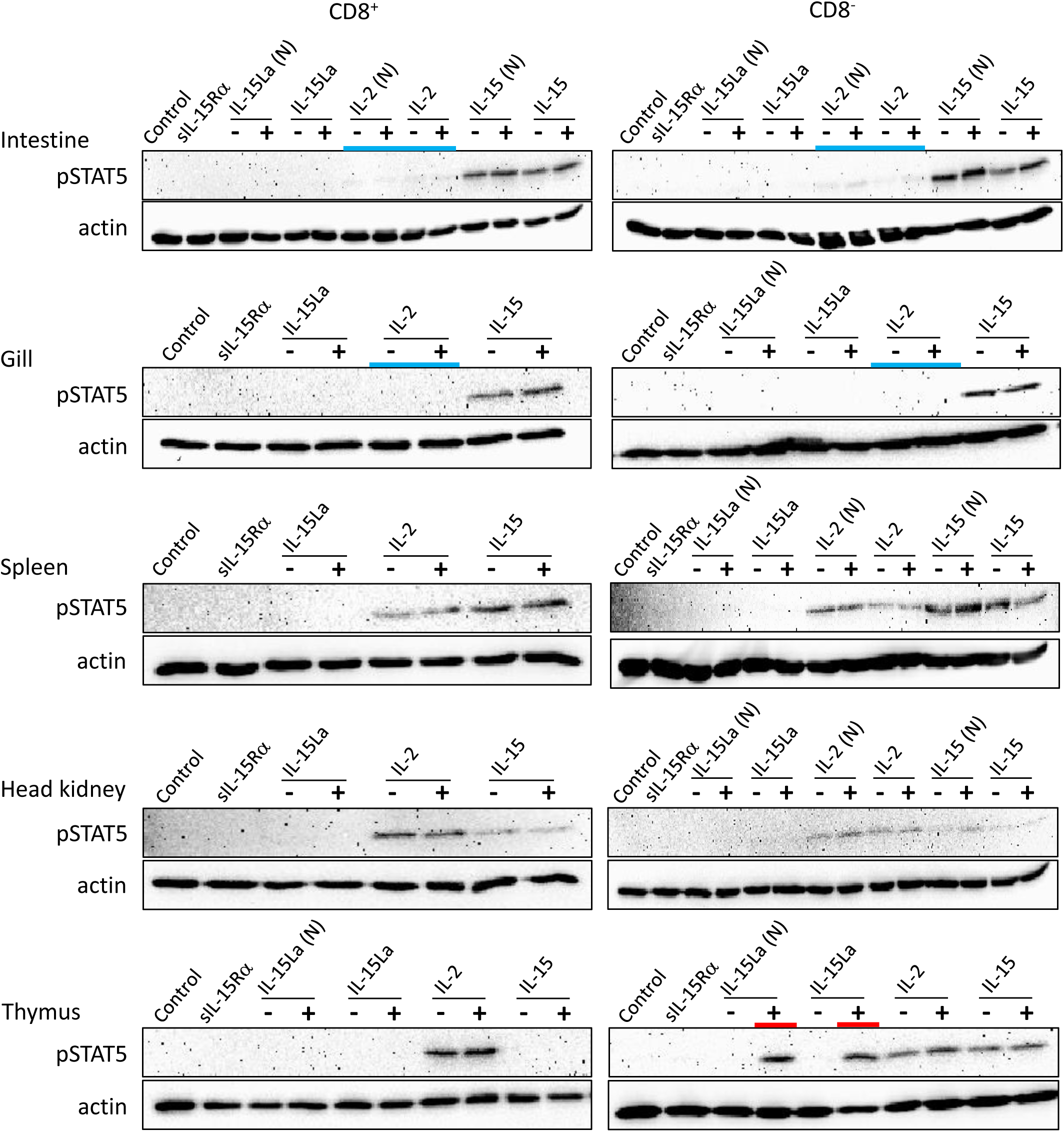
Phosphorylation of STAT5 in CD8^+^ and CD8^-^ fractions of trout lymphocytes isolated from several tissues, induced after incubation for 15 min with recombinant trout cytokine containing HEK293T cell supernatants. Western blot analysis using anti-pSTAT5 mAb. Lysates of mock-treated cells were loaded as “Control”. Symbols “-” and “+” indicate whether cytokines were co-expressed with trout sIL-15Rα. Highlighted with colors are the ability of IL-15La plus sIL-15Rα to stimulate DN thymocytes (red) and the (relative) inefficiency of IL-2 to stimulate lymphocytes from the mucosal tissues intestine and gill (blue). An addition “(N)” indicates that no tag was added to the recombinant cytokine (see Fig. S2). See Fig. S5C for experiment repeats and additional information.

During our studies, monoclonal antibodies against rainbow trout CD4-1 and CD4-2 became available (51); whether CD4-1 and CD4-2 have similar or different functions is not known, but in CD4-positive lymphocytes they commonly are co-expressed (51). To further investigate which thymocytes of trout were stimulated by IL-15La and IL-15Lb, thymocytes were labeled with an anti-CD8α monoclonal antibody with a different isotype (see Fig. S3A) than the above-mentioned (50) and additionally labeled for CD4 (using a mixture of anti-CD4-1 and anti-CD4-2; Fig. S3). Upon stimulation with supernatants of cells transfected for the various trout cytokines, it was found that IL-15La and IL-15Lb induced STAT5 phosphorylation in only unstained thymocytes (i.e. double negative or DN thymocytes; Figs. 8 and S5D; highlighted with red bars in Fig. 8). As observed for the CD8^-^ thymocytes (Figs. 7 and S5C), this stimulation was dependent on co-presence of, or fusion to, sIL-15Rα (Figs. 8 and S5D; the “RLI” protein is a fusion version). A notable observation is that cells stained for both CD4 and CD8 molecules (double positive or DP thymocytes) were only sensitive to IL-2 and not to IL-15 or to IL-15L (Figs. 8 and S5D; highlighted with a magenta bar in Fig. 8).

**Fig. 8.**
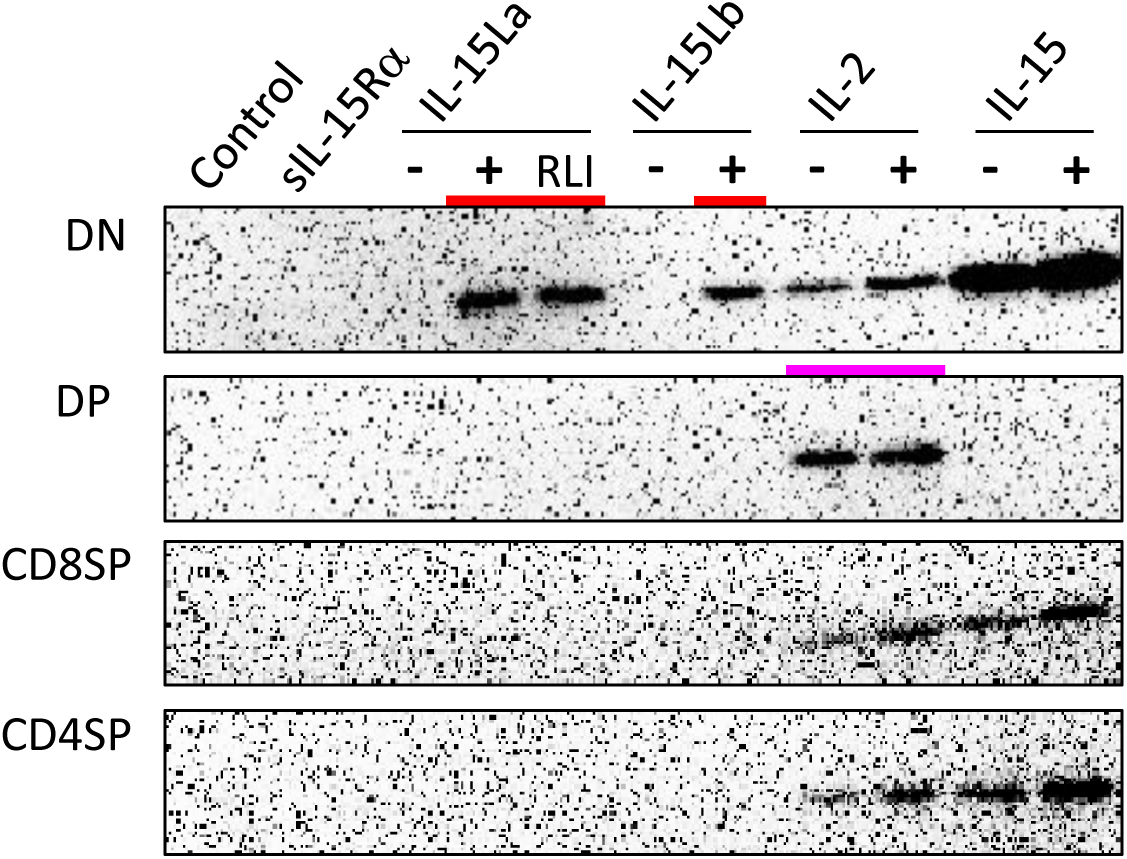
Phosphorylation of STAT5 in CD8^-^CD4^-^ (DN), CD8^+^CD4^+^ (DP), CD8^+^CD4^-^ (CD8SP), and CD8^-^CD4^+^ (CD4SP) fractions of trout thymocytes, induced after incubation for 15 min with recombinant trout cytokine containing HEK293T cell supernatants. Western blot analysis using anti-pSTAT5 mAb. Lysates of mock-treated cells were loaded as “Control”. Symbols “-” and “+” indicate whether cytokines were co-expressed with trout sIL-15Rα. Highlighted with colors are the ability of IL-15La plus sIL-15Rα, IL-15Lb plus sIL-15Rα, and a fusion of trout IL-15La to, in this case, human sIL-15Rα (RLI), to stimulate DN thymocytes (red) and the ability of IL-2 to stimulate DP thymocytes (magenta). See Fig. S5D for experiment repeats and additional information.

### Expression of trout IL-2, IL-15 and IL-15L in insect cells

To enable experiments under quantitatively controlled conditions, trout FLAG-tagged IL-2, IL-15 and IL-15La, and trout Myc-tagged sIL-15Rα were expressed in insect cells using a baculovirus system. Expression of the cytokines was undertaken with or without co-expression of trout sIL-15Rα, and in the case of IL-15 and IL-15La also as genetic fusions with trout sIL-15Rα; the fusion products were named IL-15-RLI and IL-15La-RLI. Recombinant proteins were isolated from the supernatant using anti-FLAG agarose and the resulting preparations were analyzed by size exclusion chromatography and by Coomassie staining and Western blotting after SDS-PAGE. Western blot analyses revealed that the sIL-15Rα proteins could only be isolated by anti-FLAG agarose when co-expressed with FLAG-tagged IL-2, IL-15 or IL-15La (Figs. S4A-to-C), confirming the interaction of all three cytokines with IL-15Rα as already shown with different experiments in Fig. 5. Size exclusion chromatography results indicated that IL-15, IL-15La, sIL-15Rα and IL-2+sIL-15Rα preparations may be unstable and prone to aggregation, and these preparations were not used for functional assays. For functional studies of IL-2 a preparation was used which mainly behaved as an apparent homodimer during size exclusion chromatography (Fig. S4F) as described for mammalian IL-2 preparations (52); Western blot analysis of the purified trout IL-2 also suggested the ability to form homodimers (Fig. S4F). Since initial analyses indicated functional similarity between the noncovalent associations and genetically linked forms of IL-15 or IL-15La with sIL-15Rα (Figs. S5F-1 and -2; see also Fig. 8), and because of the convenience and apparent stability, the preparations of the genetic fusion products IL-15-RLI and IL-15La-RLI (Figs. S4D and S4E) were selected over the noncovalent associations for further functional studies. When using sensitive cells, the trout IL-2, IL-15-RLI and IL-15La-RLI proteins were found to induce pSTAT5 from concentrations of 40 pM or less (Fig. S5F), which is reminiscent of the working concentrations found for recombinant IL-2 and complexes of IL-15 with sIL-15Rα in human systems (16).

### High concentrations of trout IL-15La-RLI induce STAT5 phosphorylation in trout splenocytes

Three different concentrations (5, 25 and 125 nM) of recombinant IL-2, IL-15-RLI and IL-15La-RLI proteins isolated from insect cells were used to stimulate CD4^+^CD8^-^ (CD4SP [single positive]), CD4^-^CD8^+^ (CD8SP), and CD4^-^CD8^-^ (DN) lymphocyte fractions of thymus, intestine and spleen, while for the thymus this analysis also included the CD4^+^CD8^+^ (DP) fraction [which is only abundant in that tissue; (51)]. Even when using high concentrations of purified cytokines, important findings obtained by using supernatants of HEK293T cells (Figs. 7, 8, S5C and S5D) were confirmed; for example, intestinal lymphocytes were hardly responsive to IL-2, and DP thymocytes were stimulated only by IL-2 (Figs. 9 and S5E; highlighted by blue and magenta bars, respectively, in Fig. 9). Also, the sensitivity of DN thymocytes to IL-15La+sIL-15Rα (in this case as RLI fusion form) was confirmed (highlighted by a red bar in Fig. 9). However, now, at the highest tested concentration of purified IL-15La-RLI, also preparations of DN and CD8SP splenocytes, and CD4SP and CD8SP thymocytes, were detectably stimulated (highlighted by orange bars in Fig. 9; more visible for thymocytes in Fig. S5E-2). An additional observation was that pSTAT5 levels in DN splenocytes were not very responsive to IL-2 treatment (highlighted with a green bar in Fig. 9; see also Fig. S5C-6).

**Fig. 9.**
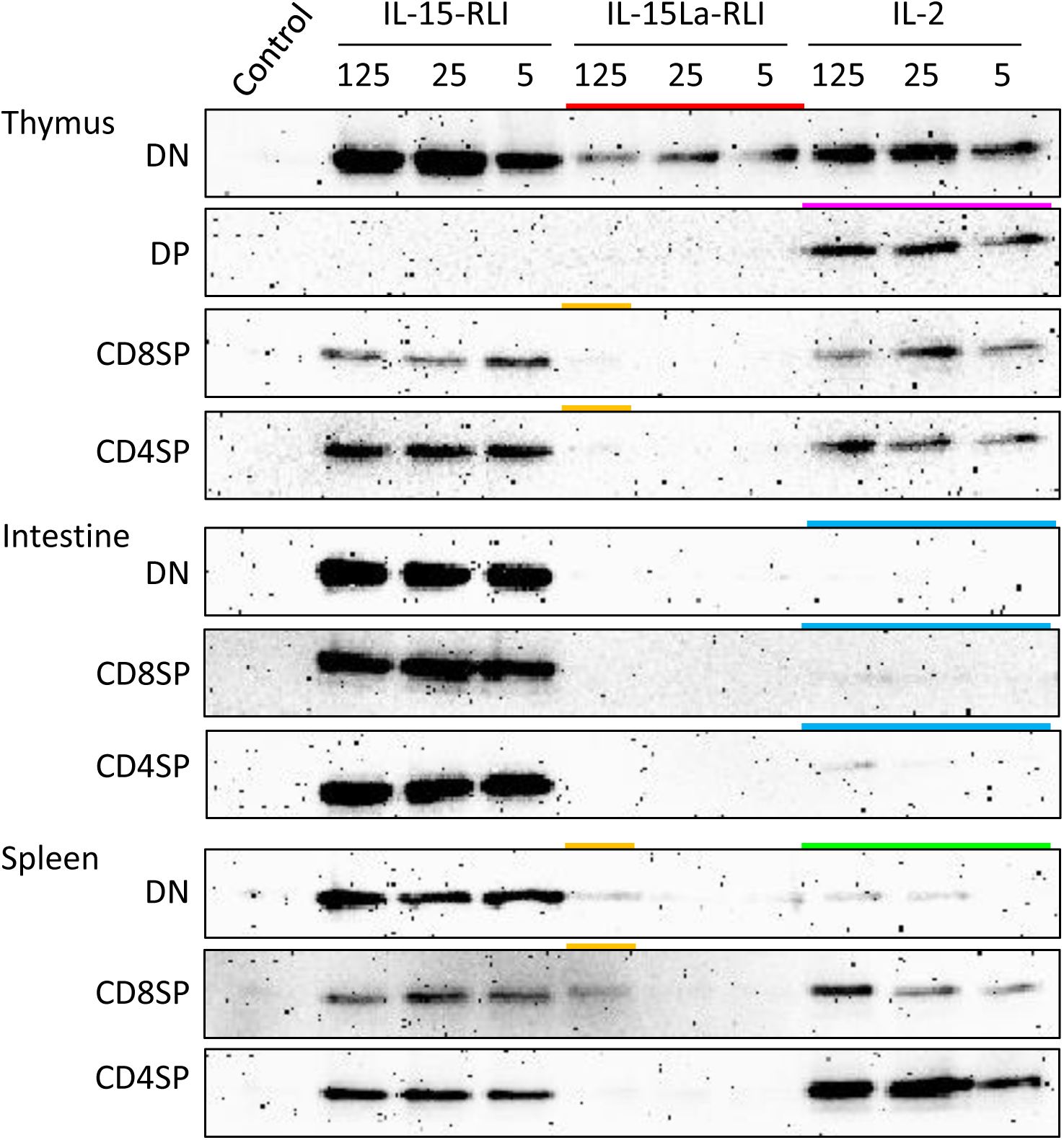
Phosphorylation of STAT5 in lymphocyte fractions of trout thymus, intestine and spleen, induced after incubation for 15 min with purified recombinant trout cytokines (produced in insect cells) IL-2, IL-15-RLI and IL-15La-RLI at 5, 25 and 125 nM. Western blot analysis using anti-pSTAT5 mAb. DN, CD8^-^CD4^-^; DP, CD8^+^CD4^+^; CD8SP, CD8^+^CD4^-^; CD4SP, CD8^-^CD4^+^. Lysates of mock-treated cells were loaded as “Control”. Highlighted with colors are the ability of IL-15La-RLI to stimulate DN thymocytes (red), the ability of IL-2 to stimulate DP thymocytes (magenta), the (relative) inefficiencies of IL-2 to stimulate intestinal lymphocytes (blue) and DN splenocytes (green), and the weak ability of IL-15La-RLI to stimulate DN and CD8SP splenocytes and CD8SP and CD4SP thymocytes (orange). See Fig. S5E for additional information.

### Trout IL-15 (+sIL-15Rα) induces expression of type 1 immunity marker genes in trout total splenocytes but trout IL-15L+sIL-15Rα induces expression of type 2 immunity marker genes

After preliminary experiments, judging the technical feasibility and reproducibility of experiments and results, and the fact that splenocytes were sensitive to IL-15L+sIL-15Rα as shown by the pSTAT5 analysis (Fig. 9), we decided to concentrate on trout splenocytes (and not cells from other tissues) for further RT-qPCR analysis after cytokine stimulation. Purified trout IL-2, IL-15-RLI and IL-15La-RLI were incubated at 0.2, 1 and 5 nM concentrations with total splenocytes, and after 4 h and 12 h incubation the RNA of the cells was isolated and subjected to RT-qPCR analysis to assess the expression levels of type 1 immunity marker genes *interferon γ* (*IFNγ*) and *perforin*, and type 2 immunity marker genes *IL-4/13A*, *IL-4/13B1* and *IL-4/13B2*. IL-2 significantly enhanced *IFNγ*, *perforin*, *IL-4/13B1* and *IL-4/13B2*; IL-15-RLI significantly enhanced *IFNγ* and *perforin*; and IL-15La-RLI significantly enhanced *IL-4/13A*, *IL-4/13B1* and *IL-4/13B2* (Fig. 10). To ensure that the observations were not caused by preparation artifacts, similar experiments were performed with supernatants of transfected HEK293T cells. The results (Fig. S6A) are comparable to those in Fig. 10, and provide the important additional observations that non-covalent complexes between IL-15La and sIL-15Rα, and between IL-15Lb and sIL-15Rα, also specifically enhanced expression of *IL-4/13A*, *IL-4/13B1* and *IL-4/13B2*. A further finding, consistent with the pSTAT5 assay results (Figs. 7, 8, S5C and S5D), was that IL-15 with and without sIL-15Rα seemed to have similar potencies in enhancing *IFNγ* and *perforin* expression, but that IL-15La and IL-15Lb fully depended on co-expression with sIL-15Rα for function (Fig. S6A).

**Fig. 10.**
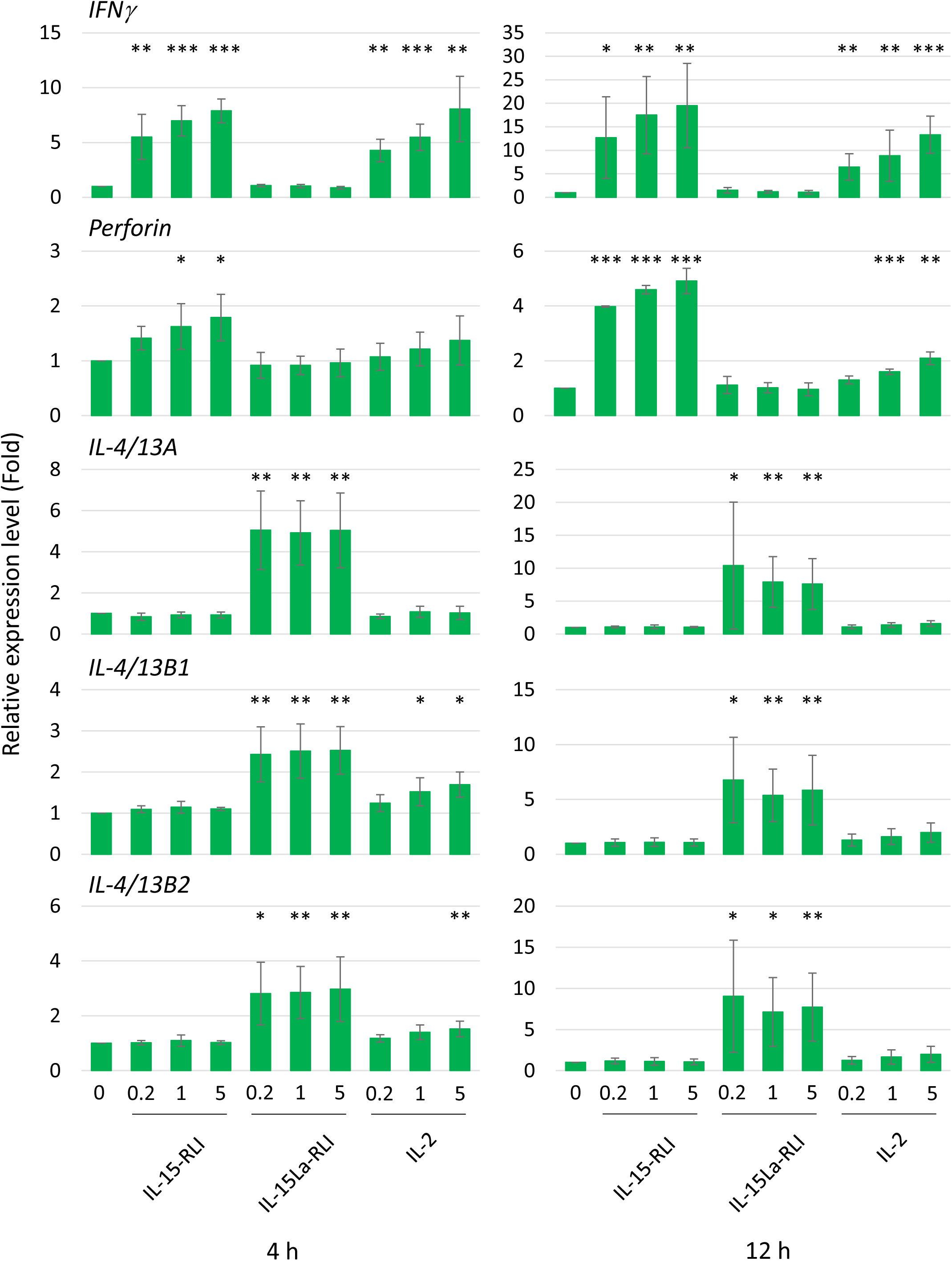
Relative expression levels of *IFNγ*, *perforin*, *IL-4/13A*, *IL-4/13B1* and *IL-4/13B2* in trout total splenocytes induced after incubation for 4 h and 12 h with purified recombinant trout cytokines IL-2, IL-15-RLI and IL-15La-RLI (produced in insect cells) at 0.2, 1 and 5 nM. Expression levels were normalized to *EF1A* expression and the values for the mock-treated control were set to 1 in each experimental panel. The average values of four biological experiments are shown together with error bars representing SD. In cases in which the average value was more than 1.5-fold higher than in the matching controls, one, two, or three asterisks indicate p-values smaller than 0.05, 0.01, or 0.001, respectively, based on paired samples T-test for the log-adjusted values of relative expression levels in samples and matching controls (97).

### Trout IL-15La-RLI efficiently induces type 2 immunity marker gene expression in CD4^-^CD8^-^IgM^-^ splenocytes

An additional stimulation experiment was performed using 0.2 and 5 nM concentrations of purified trout IL-2, IL-15-RLI and IL-15La-RLI for stimulation of sorted CD4^+^, CD8^+^, IgM^+^ and CD4^-^ CD8^-^IgM^-^ (triple negative or TN) fractions of spleen morphological lymphocytes. On average, the relative abundancies of each of the four fractions were: 24% CD4^+^ cells, 6% CD8^+^ cells, 39% IgM^+^ cells, and 31% TN cells (Fig. S3E). From previous studies it follows that, as in mammals, and although probably none of the populations was fully homogeneous, the trout CD4^+^ cells included helper and regulatory TCRαβ^+^ T cells (35, 51, 53), the CD8^+^ cells included cytotoxic TCRαβ^+^ T cells (50, 54), the IgM^+^ cells probably predominantly represented IgM^+^ B cells [e.g. (55)], and the TN cells probably were a mixture of several cell populations [e.g. (55–58)]. RT-qPCR analysis revealed that among the four populations the TN cells expressed the highest constitutive and cytokine-induced expression levels of cytokine genes *IFNγ*, *IL-4/13A*, *IL-4/13B1* and *IL-4/13B2* (Fig. 11). The highest constitutive levels of *perforin* were found in CD8^+^ cells (Fig. 11), but, for interpretation at the single cell level, it should be realized that this may be a more homogenous population than the TN cells. Only in the TN cells the *perforin* levels were found significantly enhanced after cytokine stimulation (Figs. 11 and S6B). Expression patterns induced by the individual cytokines were similar as observed for total splenocytes (Fig. 10), with IL-15-RLI efficiently enhancing the type 1 immunity marker genes *IFNγ* and *perforin*, with IL-15La-RLI efficiently enhancing the type 2 immunity marker genes *IL-4/13A*, *IL-4/13B1* and *IL-4/13B2*, and with IL-2 efficiently enhancing the type 1 immunity marker genes *IFNγ* and *perforin* but also the type 2 immunity marker gene *IL-4/13B1* (Fig. 11). Different from the observations for trout total splenocytes (Fig. 10), however, was that IL-15-RLI was found to have (p<0.05) a stimulatory effect on *IL-4/13A* expression by TN cells, although the levels of *IL-4/13A* induced by IL-15-RLI were much lower than induced by IL-15La-RLI (Fig. 11). Such IL-15 activity would not be in conflict with reports for mammals, since although the overall dominant effect of mammalian IL-15 is the stimulation of type 1 immunity (7, 31, 32), in isolated experiments mammalian IL-15 was found able, for example, to induce the expression of the type 2 immunity cytokine IL-4 in mast cells (59).

**Fig. 11.**
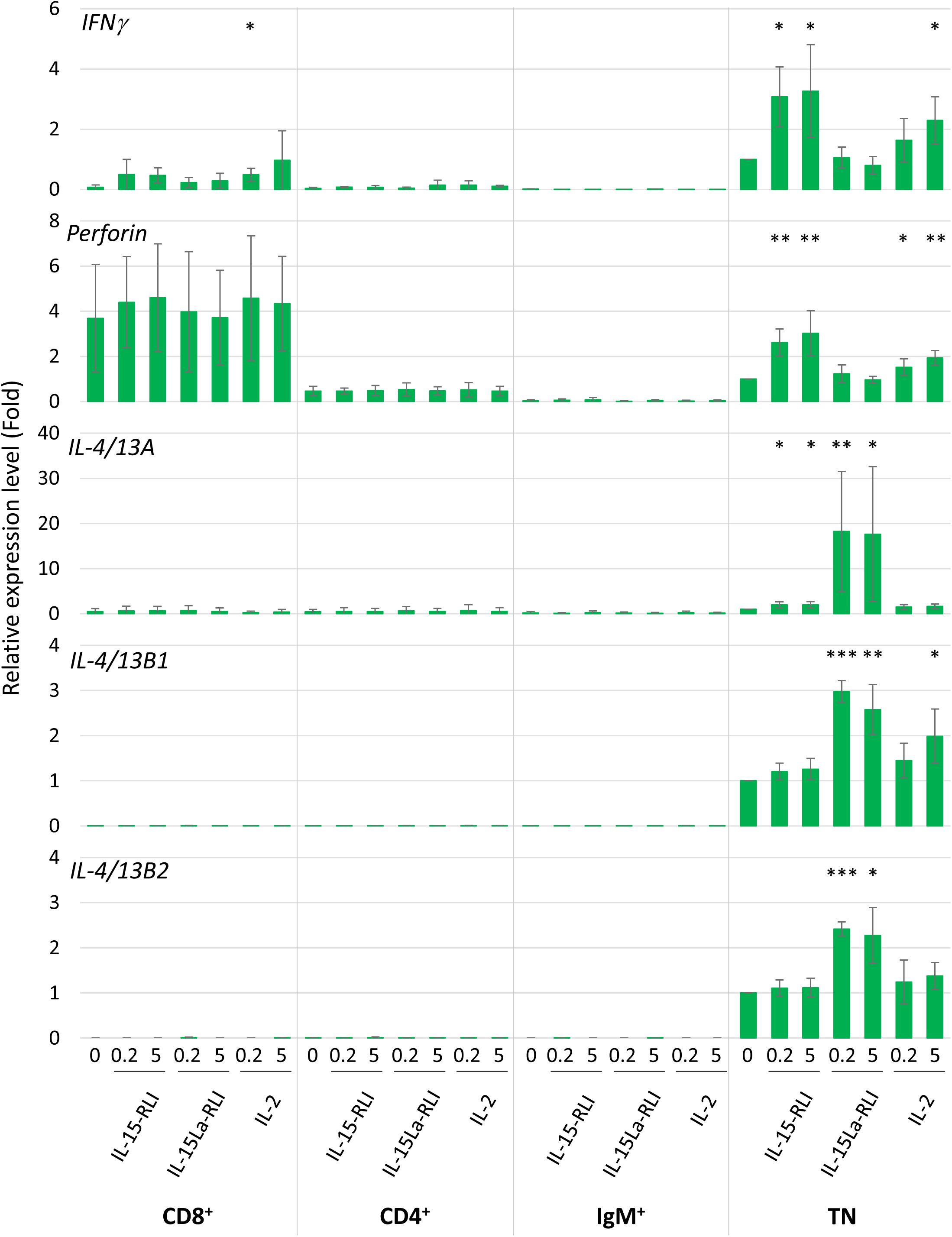
Relative expression levels of *IFNγ*, *perforin*, *IL-4/13A*, *IL-4/13B1* and *IL-4/13B2* in CD4^+^, CD8^+^, IgM^+^ and CD4^-^CD8^-^IgM^-^ (TN) trout spleen morphological lymphocytes after incubation for 12 h with purified recombinant trout cytokines IL-2, IL-15-RLI and IL-15La-RLI (produced in insect cells) at 0.2 and 5 nM. Expression levels were normalized to *EF1A* expression and the values for the mock-treated TN control were set to 1 in each experimental panel. The average values of four biological experiments are shown together with error bars representing SD. Asterisks indicate cases with estimated significance as described for Fig. 10. A table with the underlying Ct values is shown in Fig. S6B(a). For depiction at a larger scale and an alternative analysis of the results for the CD4^+^, CD8^+^ and IgM^+^ cells, see Figs. S6B-(b) and -(c).

IL-2 and IL-15 are known as important growth and survival factors for distinct populations of lymphocytes (3, 25, 30, 31, 60), and the observation in the present study that there is no stringent correlation between cytokine-mediated induction of pSTAT5 and marker gene expression (compare Figs. 9 and 11) may relate to the fact that cell growth/survival and cell functional activity are not identical processes. It should also be realized that mammalian IL-2 and IL-15 can activate more transcription factors than only their dominantly activated transcription factor STAT5 (15, 17, 59), and future studies should establish antibodies for allowing a more extensive analysis of activated transcription factors in fish. Future research in fish should also try to establish antibodies against potential receptors of the IL-2/15/15L family and other cell surface markers so that sensitive cell populations can be further characterized.

## Discussion

The current study shows within species, and cross-species, interactions between the cytokines trout IL-2, IL-15, IL-15La and IL-15Lb, and bovine IL-15 and IL-15L, and the receptor chain IL-15Rα of both cattle and trout (Fig. 5). We are not aware of any other reports on fish-mammalian cross-species interactions involving cytokines and receptor chains, or cytokines and their heterodimer complex partners. Trout and cattle shared their last common ancestor around 416 million years ago (61), emphasizing how ancient the IL-2/15/15L-to-IL-15Rα interaction system is. The result was not unexpected, because residues in IL-15 and IL-15Rα for ligand-receptor binding are very well conserved from cartilaginous fish to mammals, and the respective IL-15 residues are also well conserved in IL-15L and in fish IL-2 (Fig. 3) (13). Whilst mammalian IL-15 binds IL-15Rα with an unusually high affinity (5, 20), mammalian IL-2 binds IL-2Rα with much lower affinity (20), agreeing with the relatively poor conservation of the relevant binding residues among tetrapod IL-2 and IL-2Rα (Fig. 3) (13), and the differences in stability of free mammalian IL-2 and IL-15 (62–64). It was estimated that *IL-2Rα* originated from an *IL-15Rα* duplication early in tetrapod evolution (13, 35), but when in tetrapod evolution IL-2 and IL-2Rα acquired their mutual specificity (Fig. 5) (19, 20, 65) is unclear. For example, chicken IL-2 (66) is still very similar to IL-15 (Fig. 3) and in the past was even mistaken for it (67), and it would be interesting to investigate its alpha receptor chain binding specificity.

The stable secretion of human IL-15 is significantly enhanced by co-expression with soluble IL-15Rα (64, 68). Likewise, stable secretion of bovine and trout IL-15 and IL-15L was largely enhanced by co-expression with soluble IL-15Rα (Fig. 6). Furthermore, as found for human IL-2 (40), bovine and trout IL-2 were stably secreted in the absence of co-expression with the respective soluble receptor alpha chain, IL-2Rα or IL-15Rα (Fig. 6). Therefore, it can be concluded that during evolution the propensities of IL-2 to act as a free cytokine and of IL-15 and IL-15L to behave as a “heterodimer” with IL-15Rα were already established at the level of fish. Compared to IL-15, IL-15L appears to be even more dependent on in *trans* presentation with IL-15Rα than found for IL-15, both in regard to apparent stability (Fig. 6) and function (Figs. 7, 8 and S6A). In the literature, the established abilities of mammalian IL-2 to be presented in *trans* (69), and of mammalian IL-15 to function as a free cytokine [e.g. (21, 34)], are sometimes forgotten. However, that trout IL-2 was also readily found at the surface of IL-15Rα co-expressing cells (Fig. 5) and trout IL-15 was also able to function as a free cytokine (Figs. 7, S5C and S6A), suggest that both the in *cis* and in *trans* pathways are functionally relevant ancient traits of both cytokines. Future studies should focus on the identification of the signaling receptors for the trout cytokines, and further investigate potential functional differences between the free and IL-15Rα-bound cytokine forms.

One important reason for the selection of IL-2 over IL-15 for acquiring a dominant role in T_reg_ stimulation during evolution was probably that its free diffusion can aid in the recruitment of T_regs_ to sites of inflammation (25). In a pufferfish, CD4^+^IL-15Rα^+^ naïve lymphocytes were found to express *FOXP3* and to have immunosuppressive functions, while CD4^+^IL-15Rα^-^ lymphocytes from this fish did not express *FOXP3* (35). Furthermore, the ability of zebrafish FOXP3 to induce T_reg_-like functions has been shown or suggested (70, 71). Therefore, despite the fact that fish do not have a separate IL-2Rα chain (13, 35), a preferred usage by fish IL-2 of IL-15Rα in *cis* may allow the cytokine to have a similarly important role in T_reg_ stimulation as in mammals. Different uses of the receptor alpha chain may also have caused, during evolution, IL-15 to be selected over IL-2 for important roles in the stimulation of lymphocytes of mucosal tissues (Fig. 8) (31, 72–76), because IL-15 presentation at the cell membrane allows the power of cytokine signaling to be retained within confined niches. In short, our data reveal that important characteristics relating to the mechanistic and functional “dichotomy” (16) observed for mammalian IL-2 and IL-15 were already established in a common ancestor of mammals and teleost fish.

Size exclusion chromatography (and also Western blot data) suggest that trout IL-2 molecules produced in insect cells form homodimers (Fig. S4F), and homodimer structures have also been described in some studies for recombinant mammalian IL-2 (52). Homodimer structures may also explain a large band observed upon Western blot analysis of trout IL-15La expressed in transfected mammalian cells (Figs. 6, S5A and S5B) or purified from insect cells (Fig. S4B), and there is evidence that, at least under some conditions, human IL-15 can form noncovalent homodimers (77, 78). A related short-chain four α-helix bundle cytokine for which homodimer formation is known is IL-5 (79), but IL-2 and IL-15 are generally considered to be monomers. However, given the indications for dimer formation in both fish and mammals, the possibility that IL-2/15/15L family cytokines may potentially form homodimers as a functionally relevant ancient trait should be critically evaluated in future studies.

*IL-15L* intact gene appears to have been lost in amphibians, birds, and many mammals (13). We have not found a function for bovine IL-15L as yet, and the present study is the first to report on IL-15L functions, including the ability of rainbow trout IL-15L to stimulate DN thymocytes and CD4^-^CD8^-^IgM^-^ splenocytes. We are not aware of any other ancient cytokine shared between fish and mammals for which the function hitherto was not known.

The developmental path of mammalian T lymphocytes within the thymus is from an early DN stage towards an intermediate DP stage, after which the cells mature to become CD4SP or CD8SP T cells that ultimately can leave the thymus (80). Fish thymocyte progressive development has not been studied in detail, but available knowledge of fish thymus organization, gene expression, and functions of mature T cells [e.g. (50, 51, 81, 82), reviewed in (53)] suggest a similar development to mammals. Probably, as in mammals (80), DN thymocytes in trout importantly consist of several stages of early T cells. In addition, as in mammals, the trout DN thymocytes likely include some B cells, although they are scarce in trout thymus [e.g. (50)], and, based on findings in mammals, may include several developmental stages of natural killer (NK) cells and innate lymphoid cells (ILCs), including multipotent precursors that may also develop into T cells (83, 84). Future research should try to identify more precisely the (sub-) population of fish DN thymocytes which is sensitive to IL-15L.

While most of the results obtained in the present study for trout IL-2 and IL-15 agree well with reports for mammals, an exception is the detected sensitivity of trout DP thymocytes to IL-2 (Figs. 8 and 9). In addition to being refractory to IL-2 and IL-15, mammalian DP thymocytes have low sensitivity to the STAT5 activating cytokine IL-7 (85, 86). Of relevance to these findings is the observation that in mice in which IL-7 sensitivity was induced at the DP stage (by genetic engineering), IL-7 stimulation could induce thymocyte development into mature CD8^+^ T cells in the absence of the normal requirement for positive selection mediated by TCR-pMHC interaction, thus bypassing a critical step in T cell education (87). Hence, it is puzzling that trout DP thymocytes are so sensitive to IL-2. Future work should try to determine whether this *ex vivo* finding has relevance within the fish thymus, try to discover where in the fish thymus IL-2 is expressed, and investigate whether the fish DP population can be divided into IL-2 responding and non-responding populations. Possibly, the IL-2-sensitive DP thymocytes are T_reg_ cells expressing relatively high levels of IL-15Rα [see mammalian study (88)], but antibodies against trout IL-15Rα which could help investigate this matter are not yet available.

Trout splenocytes were found sensitive to IL-15La-RLI as indicated by STAT5 phosphorylation (Fig. 9), and these cells were chosen for a detailed analysis by RT-qPCR analysis. In total splenocytes, trout IL-2 enhanced expression of the type 1 immunity marker genes *IFNγ* and *perforin*, and also of the type 2 immunity marker genes *IL-4/13B1* and *IL-4/13B2* (Figs. 10 and S6A), which is reminiscent of previous findings for IL-2 in trout (48, 89) and mammals (60, 90–92). In contrast, if using these target cells, trout IL-15, free or complexed with IL-15Rα, only induced the type 1 cytokine marker genes *IFNγ* and *perforin* (Figs. 10 and S6A), activities agreeing with previous findings for mammalian IL-15 (32, 93) and free trout IL-15 (42). The important novel finding from this study is that trout IL-15Rα-complexed IL-15L only enhanced expression of the type 2 immunity marker genes *IL-4/13A*, *IL-4/13B1* and *IL-4/13B2* (Figs. 10, 11 and S6A), and so can have an opposite immune function relative to IL-15. When separating trout spleen lymphocyte subpopulations using antibodies against CD4, CD8 and IgM, the highest levels of *IL-4/13A*, *IL-4/13B1* and *IL-4/13B2* expression were found for CD4^-^CD8^-^IgM^-^ cells, especially after stimulation with IL-15La-RLI (Fig. 11), suggesting that this cell population contains a subpopulation which is very important for type 2 immunity. Based on comparison with mammalian studies, and recent indications for the existence of such cells in fish (58), we suspect that these cells are similar to mammalian type 2 innate lymphoid cells (ILC2) which are specifically dedicated to type 2 immunity [reviewed in (94)]. Meanwhile, after stimulation with trout IL-15-RLI, the trout CD4^-^CD8α^-^IgM^-^ splenocytes upregulated *IFNγ* and *perforin* (Fig. 11), perhaps involving a cell subpopulation similar to mammalian NK cells because these cells are particularly sensitive to IL-15 (7, 31, 32, 93). Neither ILC2 nor NK cells have been properly identified in fish, and the present study provides additional support for their existence. The *IL-4/13* genes are homologues of mammalian *IL-4* and *IL-13* (95, 96), and IL-15L is the first cytokine found to specifically induce their expression in fish. In mammals, the cytokines TSLP, IL-25 and IL-33 are important for stimulating ILC2 cells, and we speculate that absence of one or more of these molecules in fish, as their genes have not been detected so far (53), may explain the stricter evolutionary conservation of IL-15L in fishes compared to tetrapod species.

In conclusion, the present study reveals that the mechanistic and functional dichotomies between IL-2 and IL-15 are an ancient phenomenon, as evidenced by their conservation in both fish and mammals. Furthermore, we identified an unexpected cytokine candidate for playing a role in the early stages of the type 2 immunity cytokine cascade in fish, namely IL-15L, which is closely related to the type 1 immunity cytokine IL-15. These findings are an important step in characterizing the IL-2/15/15L cytokine family and for understanding the original blueprint of the cytokine network in jawed vertebrates.

## Materials and methods

Detailed descriptions of materials and methods are provided in the supplementary Text S1. Animal experiments were in agreement with relevant guidelines for animal welfare.

## Acknowledgements

TY and UF were supported by the EU FP7 grant 311993 (TARGETFISH) and the German Research Council grant No. FI 604/7-1. JMD was supported by the Ministry of Education, Culture, Sports, Science and Technology, Japan, Grants-in-Aid for Scientific Research No. 25450319. TW received funding from the MASTS pooling initiative (The Marine Alliance for Science and Technology for Scotland), that is funded by the Scottish Funding Council (grant reference HR09011). EW was supported by the Ministry of Science and Technology of Thailand and Mahasarakham University. We thank Mrs. Susann Schares for excellent technical assistance.

