## Supplement for "The first functional characterization of ancient interleukin-15-like (IL-15L) reveals shared and distinct functions of the IL-2, -15 and -15L family cytokines"

**This PDF file includes:**

- Table of contents
- Supplementary text
- Figs. S1 to S6
- Table S1
- References for SI reference citations

**Table of contents** **Page**


Supplementary text (*Materials and methods*) 3

Fig. S1: Additional information on *IL-15L* nucleotide and deduced amino acid sequences 20

Fig. S2: Expression vectors used in this study 28

Fig. S3: Isolation of rainbow trout lymphocyte subpopulations by flow cytometry 48

Fig. S4: Preparations of recombinant trout cytokines, soluble IL-15Rα (sIL-15Rα) and RLI fusion proteins from insect cells 59

Fig. S5: Additional Western blot results 70

Fig. S6: Additional information on RT-qPCR experiments 103

Table S1: Percentages of $FLAG^+$ cells among live HEK293T cells after their transfection for FLAG-tagged trout or bovine IL-2/15/15L-family cytokines with or without co-transfection for trout or bovine IL-15Rα or bovine IL-2Rα 111

References for SI reference citations 112

Supplementary Text

#### Materials and methods

**Table of Contents** **Page**


Rainbow trout 5

Rainbow trout permanent cell lines and primary head kidney (HK) macrophage cultures 5

Permanent human and insect cell lines 6

Database searches and analysis of nucleotide and deduced amino acid sequences 6

Isolation of RNA, synthesis of cDNA, PCR amplifications, sequencing and cloning into expression vectors 7

Reverse transcription quantitative real-time PCR (RT-qPCR) analysis of trout *IL-15La* and *IL-15Lb* tissue distribution 8

Expression of recombinant proteins in human HEK293T cells 8

Transfection

Analysis of transfected HEK293T cells by flow cytometry

Expression of soluble cytokine (-complexes) in HEK293T cells

SDS-PAGE and Western blotting 10

Expression of recombinant proteins in insect cells 10

Transfection of recombinant bacmid DNA into High Five cells

Isolation of recombinant baculoviruses by plaque assay

Titration of recombinant baculoviruses by endpoint dilution assay

Infection of Sf9 cells with recombinant baculoviruses

Purification of recombinant cytokines from insect cell supernatants 12

Gel filtration chromatography 12

Deglycosylation assay 13

Establishment of a monoclonal antibody (mAb) against rainbow trout CD8α 13

Isolation of rainbow trout lymphocyte subpopulations 14

Stimulation of trout leukocytes with recombinant cytokines 15

Stimulation of trout leukocyte subpopulations and Western blot analysis

Stimulation of trout leukocytes and RT-qPCR analysis

Statistics 16

Tables of primer sequences 17

***Rainbow trout.***

Rainbow trout (*Oncorhynchus mykiss*) weighing between 80 and 300 gram were used in this study at three different facilities:

1. Inland Station, National Research Institute of Aquaculture (NRIA; Mie, Japan). Fish were fed commercial dry pellets and kept in 15 °C flow-through water. For semi-quantitative RT-PCR analysis, a trout individual (Trout-1) of strain Tokyo, Tokyo Metropolitan Fisheries Experimental Station (Tokyo, Japan), was investigated. For determining *IL-15La* and *IL-15Lb* sequences, 5'-RACE analysis and semi-quantitative RT-PCR, a trout individual (Trout-2) of the homozygous clonal rainbow trout strain C25 was used. These homozygous isogeneic trout had been produced from outbred strain Nagano at the Nagano Prefectural Fisheries Experimental Station (Nagano, Japan), by gynogenesis over two generations by suppression of mitosis and meiosis in the first and second generations, respectively (1). Clonality had been confirmed by DNA fingerprinting. For convenient propagation of the strains, some of the gynogenetic animals had been subjected to a treatment with methyltestosterone and developed as homozygous neomales.

2. Scottish Fish Immunology Research Centre (SFIRC), the University of Aberdeen, UK. Fish were fed commercial dry pellets and kept in 15±1 °C recirculating water. The six trout individuals used for RT-qPCR analysis had been purchased from the Mill of Elrich Trout Fishery (Aberdeenshire, Scotland, UK).

3. Friedrich-Loeffler-Institut (FLI), Federal Research Institute for Animal Health (Insel Riems-Greifswald, Germany). Fish were fed commercial dry pellets and kept at 15 °C in a partially recirculating water system. The investigated trout individuals belonged to the homozygous clonal strain C25 (see above). Most of the experiments described in the present study were done at the FLI.

Fish handing and experimental protocols complied with the guidelines for animal welfare in the respective countries and institutes.

***Rainbow trout permanent cell lines and primary head kidney (HK) macrophage cultures.***

Four rainbow trout cell lines were used for gene expression analysis: a monocyte/macrophage-like cell line RTS-11 from spleen (2), an epitheloid cell line RTL from liver (3), the fibroblastoid cell lines RTG-2 from gonad (4), and RTGill from gills (5). Cells were maintained in Leibovitz (L-15) medium (Invitrogen) containing 30 % fetal bovine serum (FBS; Labtech International, for RTS-11 cells) or 10 % FBS (for the other three cell lines and for primary HK macrophages) and antibiotics (100 U penicillin/ml and 100 μg streptomycin/ml; Invitrogen) at 20 °C. Primary HK macrophage cultures from four individual trout at the SFIRC were prepared as outlined by Costa *et al.* (6).

***Permanent human and insect cell lines.***

*HEK (human embryo kidney) 293T cells.* HEK293T cells were used for transient expression of recombinant proteins. Cells were maintained in minimal essential medium (MEM) supplied with 10 % FBS at 37 °C in a 2.5 % $CO_2$ atmosphere.

*High Five and Sf9 cells.* Two insect cell lines, High Five and Sf9, were used for producing recombinant proteins. These cells were maintained in Grace's Insect medium supplied with lactalbumin hydrolysate, yeast extract and 5 % FBS, at 26 °C.

These cell lines and media were obtained from the Collection of Cell Lines in Veterinary Medicine (CCLV) at FLI.

***Database searches and analysis of nucleotide and deduced amino acid sequences.***

BLAST similarity searches were performed on sequence datasets of the National Center for Biotechnology Information (NCBI; http://blast.ncbi.nlm.nih.gov/Blast.cgi) (7) and the Ensembl database of the European Bioinformatics Institute (EBI; https://www.ensembl.org/) (8). Retrieved sequences were analyzed using genetic analysis software GENETYX (Version 12.0.3) and FGENESH gene prediction software (www.softberry.com) (9). For deduced amino acid sequences, the leader peptides and $\alpha$-helix structures of mature proteins were predicted using SignalP (http://www.cbs.dtu.dk/services/SignalP/) (10) and Jpred 4 softwares (http://www.compbio.dundee.ac.uk/jpred4/index_up.html) (11), respectively. Alignments of deduced amino acid sequences were performed manually, based on comparisons of more sequences, and considerations of gene and protein structures and of phylogeny (12), and also considering the clarity of the figure. For construction of a phylogenetic tree, see Fig. S1D.

Read numbers per $10^8$ reads of *IL-15La* and *IL-15Lb* were determined by similarity searches against tissue-specific single read archive (SRA) datasets using the BLAST search function at NCBI. For rainbow trout, the SRA datasets of Bioproject PRJEB4450 (NCBI datasets ERX297509-to-297524) (13), Bioproject PRJNA389609 (NCBI datasets SRX2894150-to-2894164) (14) and Bioproject PRJNA380337 (NCBI datasets SRX2668643-to-2668653 and SRX2668655-to-2668657; Norwegian University of Life Sciences) were investigated. For Atlantic salmon, the SRA datasets of Bioproject PRJNA260929 (NCBI datasets SRX1046658, SRX1052181, SRX1052182, SRX1052184, SRX1052187-to-1052192; Norwegian University of Life Sciences) and Bioproject PRJNA72713 (NCBI datasets SRX608567, SRX608569, SRX608571, SRX608574, SRX608575, SRX608579, SRX608583, SRX608588, SRX608594, SRX608599, SRX608607, SRX608616, SRX608620, SRX608621; University of Victoria) were investigated. The species-specific *IL-15La* or *IL-15Lb* ORF sequences were subjected to "Megablast" analysis (blastn) using default settings except that the "max target sequences" number was changed to 20,000 and the "word size" was changed to 64. To ensure specificity of the Megablast analysis, only matches with score values ≥187 for PRJEB4450, ≥185 for PRJNA389609 and PRJNA72713, ≥233 for PRJNA380337, ≥192 for Bioproject PRJNA260929 were counted.

***Isolation of RNA, synthesis of cDNA, PCR amplification, sequencing and cloning into expression vectors.***

Total RNA samples of trout were isolated from tissues by two-fold purification with TRIzol (Gibco) and stored at the NRIA. Equal amounts of RNA were transcribed into cDNA using Superscript transcriptase (Invitrogen). A cDNA sample from spleen of Trout-2 was used for the amplification of the full-length *IL-15La* open reading frame (ORF) using primer set Trout_IL-15La_CDS and ExTaq polymerase kit (Takara) while a cDNA sample from gill of Trout-2 was used for the amplification of the full-length *IL-15Lb* ORF using primer set Trout_IL-15Lb_CDS. These primer sequences are shown in "Table of primer sequences A" below. For 5'-RACE analysis cDNA samples were synthesized from total RNA of spleen and gill of Trout-2 using the SMARTER RACE cDNA amplification system (Clontech). The first PCR was performed using spleen cDNA (for IL-15La) or gill cDNA (for IL-15Lb) with NUP primer (provided with the kit) and a specific primer for corresponding gene. Subsequently, nested PCR was performed using each first PCR product with UPM primer (provided with the kit) and a specific inner primer for corresponding gene. The sequences of primers used for 5'-RACE analysis are shown in Table of primer sequences A. The first PCR schedule was 94 °C for 5 min, 5 x (94 °C for 30 sec, 72 °C for 1:30 min), 10 x (94 °C for 30 sec, 70 °C for 30 sec, 72 °C for 1 min), 25 x (94 °C for 30 sec, 68 °C for 30 sec, 72 °C for 1 min), 72 °C for 7 min. After diluting the product of the first reaction (1/200), the amplification schedule for the nested PCR was 94 °C for 5 min, 32 x (94 °C for 30 sec, 60 °C for 1 min, 72 °C for 30 sec), 72 °C for 7 min. The amplified *IL-15La* and *IL-15Lb* full-length ORF and 5'-RACE fragments were prepared for sequencing by standard TA-cloning with the pGEM T-Vector System (Promega). The sequences of multiple clones were determined by dideoxy chain termination method and using an automated sequencer to exclude PCR errors. Assembled sequences of the overlapping full-length ORF and 5'-RACE amplifications of rainbow trout *IL-15La* and *IL-15Lb* were deposited to GenBank and are available as accessions MK619679 and MK619680, respectively.

For semi-quantitative analysis of tissue distribution of transcripts, PCR was performed with the ExTaq polymerase kit, using equal amounts of cDNA solution as templates, and the primer sets Trout_IL-15La, Trout_IL-15Lb and Trout_EF1A (Table of primer sequences B), for amplification of fragments of *IL-15La*, *IL-15Lb* and *elongation* *factor 1 alpha* (*EF1A*), respectively. For the semi-quantitative PCR analysis of *IL-15La* and *IL-15Lb* expression, the amplification schedule was: 94 °C for 5 min, 32 x (94 °C for 30 sec, 60 °C for 30 sec, 72 °C for 40 sec), 72 °C for 7 min; and for *EF1A* amplification, the schedule was: 94 °C for 5 min, 25 x (94 °C for 30 sec, 60 °C for 30 sec, 72 °C for 30 sec), 72 °C for 7 min.

For construction of DNA expression vectors, gene sequences were amplified from cDNA or commercially ordered, and, often after PCR-mediated gene modifications, cloned into commercial DNA plasmid vectors by using appropriate restriction enzymes behind the CMV-IE promoter, or into the baculovirus transfer vector pFBD-P10Uhis-ieGFP behind the p10-promoter (for cloning details see Fig. S2). The vector pFBD-P10Uhis-ieGFP is based on the vector pFBDΔXhoI_Histag (15) which is a derivate of pFastBac-Dual (Invitrogen) in which the PolH promoter region was replaced by a CMV-IE promoter driven GFP expression cassette (Dr. Günther M. Keil, personal communication). The expression vectors were multiplied in *E. coli* and isolated by standard techniques. To check whether the sequences were correctly inserted, all DNA expression vectors were sequenced by dideoxy chain termination method and using an automated sequencer.

***Reverse transcription quantitative real-time PCR (RT-qPCR) analysis of trout IL-15La and IL-15Lb tissue distribution.***

Six healthy rainbow trout were used at the SFIRC for RT-qPCR analysis of *IL-15La* and *IL-15Lb* tissue distribution. The RNA preparations from trout tissues, cell lines, and primary HK macrophage cultures, and the following RT-qPCR analysis, were performed as described previously (16, 17). The relative expression levels of each IL-15L gene were normalized against the expression level of *EF1A*. A common reference containing equal molar amounts of purified PCR products of trout *IL-15La*, *IL-15Lb* and *EF1A* was used for the quantification. The primer sets used for amplification were Trout_IL-15La_qPCR, Trout_IL-15Lb_qPCR and Trout_EF1A_qPCR (Table of primer sequences B).

***Expression of recombinant proteins in human HEK293T cells.***

*Transfection.* HEK293T cells were transfected using X-tremeGENE HP DNA Transfection Reagent (Roche) as described in our previous study (18), with slight changes. For co-expression of cytokines with IL-15Rα or IL-2Rα, HEK293T cells at a 80 - 90 % confluency were co-transfected with 2 μg of the cytokine-encoding plasmid together with 2 μg of IL-15Rα-encoding plasmid or IL-2Rα-encoding plasmid (in total 4 μg) per well. To express only cytokines or receptor α chains, HEK293T cells were co-transfected with 2 μg of the respective plasmid together with 2 μg of “empty” pcDNA3.1 or pRc/CMV2 commercial vector (Invitrogen). Negative control cells were transfected with 4 μg of empty vector. The recombinant molecules were expressed by using the following expression vectors (for sequences see Fig. S2): bovine IL-2, pRcCMV2-*Bos-IL-2-FLAG*; bovine IL-15; pRcCMV2-*Bos-IL-15-FLAG*; bovine IL-15L, pRcCMV2-*Bos-IL-15L-FLAG*; bov.IL-15Lhyb-h-RLI, pcDNA3.1-*IL-2-Lead-RLI-bov.IL-15Lhyb*; bovine (full-length) IL-15Rα, pcDNA3.1-*Bos-IL-15Rα-Myc-His*; bovine soluble IL-15Rα (aka sIL-15Rα), pcDNA3.1-*Bos-solIL-15Rα-Myc-His*; bovine (full-length) IL-2Rα, pcDNA3.1-*Bos-IL-2Rα-Myc-His*; bovine soluble IL-2Rα (aka sIL-2Rα), pcDNA3.1-*Bos-solIL-2Rα-Myc-His*; trout IL-2, pcDNA3.1-*trout-IL-2-FLAG*; trout IL-2(N), pcDNA3.1-*trout-IL-2-(non-tagged)*; trout IL-15, pcDNA3.1-*trout-IL-15-FLAG*; trout IL-15(N), pcDNA3.1-*trout-IL-15-(non-tagged)*; trout IL-15La, pcDNA3.1-*trout-IL-15La-FLAG*; trout IL-15La(N), pcDNA3.1-*trout-IL-15La-(non-tagged)*; trout IL-15Lb, pcDNA3.1-*trout-IL-15Lb-FLAG*; trout IL-15Lb(N), pcDNA3.1-*trout-IL-15Lb-(non-tagged)*; trout IL-15La-h-RLI, pcDNA3.1-*IL-2-Lead-RLI-trout-IL-15La*; trout (full-length) IL-15Rα, pcDNA3.1-*trout-IL-15Rα-Myc*; trout soluble IL-15Rα (aka sIL-15Rα), pcDNA3.1-*trout-solIL-15Rα-Myc*.

*Analysis of transfected HEK293T cells by flow cytometry*. In order to check the binding ability of receptor α-chains for each cytokine, HEK293T cells were co-transfected with plasmids encoding full-length (transmembrane) forms of IL-15Rα or IL-2Rα and plasmids encoding the cytokines, or, as negative controls, transfected with only one of these plasmids or with empty vector alone (see above). Two days after transfection, HEK293T cells were collected, washed and stained with mouse ANTI-FLAG M2 Monoclonal Antibody (Sigma) and anti-mouse IgG, IgM (H+L) secondary antibody conjugated with Alexa Fluor 488 (Thermo Fisher Scientific) diluted according to the manufacturer's instructions. The stained HEK293T cells were analyzed with a FACSCalibur flow cytometer (BD Biosciences). Conditions were adjusted by setting the thresholds for conjugate controls. Dead cells were excluded from analysis by propidium iodide (PI) staining. The data were analyzed using BD CellQuest Pro Software (BD Biosciences).

*Expression of soluble cytokine (-complexes) in HEK293T cells.* HEK293T cells were co-transfected with plasmids encoding soluble forms of IL-15Rα or IL-2Rα and plasmids encoding the cytokines, or transfected with only one of these plasmids, or (as negative control) with empty vector alone. Medium of HEK293T cells was replaced to EX-CELL Serum-Free Medium (Sigma) before transfection. Two days after transfection, 2 ml of supernatant was collected from each well, and filtered through a 0.22 μm pore PVDF membrane (Syringe Driven Filter Unit, Millex-GV). For analysis by Western blotting as shown in main text Fig. 6 and supplementary Fig. S5B, the 2 ml supernatants were concentrated to 40 - 50 μl by ultrafiltration with a 3 kDa nominal molecular weight cutoff membrane (Amicon Ultracel - 3K, Millipore), yielding the "concentrated supernatant" samples. For leukocyte stimulation experiments, supernatants were used without concentration ("unconcentrated supernatant" samples). Remaining HEK293T cells in each well were collected, pelleted and lysed in 100 µl of NP40 Cell Lysis Buffer (Thermo Fisher Scientific) supplied with Protease Inhibitor Cocktail (Sigma), and used for further analysis as "cell lysate" samples.

***SDS-PAGE and Western blotting.***

Fifteen µl of the samples were mixed with 5 µl of 4 x reducing Laemmli Sample Buffer (Bio-Rad), heated for 3 min at 95 °C and electrophoresed using (unless mentioned otherwise) 12 % poly-acrylamide gels and standard procedures (19) and with PageRuler Prestained Protein Ladder (Thermo Fisher Scientific) as molecular weight marker. After electrophoretic separation, proteins were either visualized by treatment with Coomassie Brilliant Blue (CBB) R-250 staining solution (BioRad) or prepared for Western blotting by transfer to Amersham Hybond P 0.45 PVDF membranes (GE Healthcare) using a Trans-Blot Turbo Transfer System (Bio-Rad). Membranes were blocked by incubation in StartingBlock (TBS) Blocking Buffer (Thermo Fisher Scientific) and subsequently incubated overnight at 4 °C with mouse ANTI-FLAG M2 Monoclonal Antibody, Myc-Tag (9B11) Mouse mAb or Phospho-Stat5 XP Rabbit mAb (Cell Signaling Technology) specific for phosphorylated Tyr694 (Tyr694 and surrounding residues are conserved between trout and mouse STAT5), made up in blocking buffer. The membranes were washed with TBS/0.1 % Tween-20, followed by incubations with HRP-Conjugated Goat Anti-mouse IgG (Pierce) or HRP-linked Anti-rabbit IgG (Cell Signaling Technology) in blocking buffer for 1 - 2 h. All antibodies were used at the concentrations recommended by the manufacturer. Bands were visualized by chemiluminescence reaction (SuperSignal West Pico Chemiluminescent Substrate, Thermo Fisher Scientific) and documented on a VersaDoc 4000 MP workstation (BioRad) using Quantity One software (BioRad). As a loading control, membranes which had been subjected to pSTAT5-detection were stripped by incubation in 0.1 M glycine-HCI buffer (pH 2.8) for 2 h with gentle shaking at room temperature and subsequently reprobed for actin using mAb C4 (Millipore).

***Expression of recombinant proteins in insect cells.***

*Construction of recombinant bacmid DNA.* Recombinant plasmids were isolated and transformed to DH10Bac competent cells (Invitrogen) with standard procedure, after which bacmid DNA was isolated. The recombinant plasmids are explained in Fig. S2, with the names of the encoded recombinant proteins and plasmids as follows: trout IL-2, pFBD-P10Uhis-ieGFP-*trout-IL-2-FLAG*; trout IL-15, pFBD-P10Uhis-ieGFP-*trout-IL-15-FLAG*; trout IL-15La, pFBD-P10Uhis-ieGFP-*trout-IL-15La-FLAG*; trout soluble IL-15Rα (aka sIL-15Rα), pFBD-P10Uhis-ieGFP-*trout-solIL-15Rα-Myc*; trout IL-15-RLI, pFBD-

P10Uhis-ieGFP-*trout-IL-15-RLI*; trout IL-15La-RLI, pFBD-P10Uhis-ieGFP-*trout-IL-15La-RLI*.

*Transfection of recombinant bacmid DNA into High Five insect cells.* High Five cells were seeded into a 6-well plate and incubated at 26 °C for 1 h. A transfection mix with 5 µg bacmid DNA and 6 µl X-tremeGENE reagent in 100 µl α-MEM (Sigma) was prepared for each cytokine and incubated at room temperature for 40 min. These transfection mixes were diluted with 900 µl Insect-XPRESS medium, and then dropped onto the High Five cells. After 5 h incubation, the supernatant was replaced by 2 ml of fresh Insect-XPRESS medium per well and continued to be cultured. After 3 days cultivation, cells and supernatants were collected and stored at -80 °C.

*Isolation of recombinant baculoviruses by plaque assay.* Sf9 cells were seeded into 6-well plates and incubated for 30 min at room temperature. Aliquots of the transfected High Five cells and their supernatants were thawed and diluted from $10^0$ to $10^{-2}$ in Grace's insect medium, and 100 µl of each dilution was added to the well. After 1 h cultivation at 26 °C, supernatants were removed and cultures were overlaid with 1% low-melting agarose containing Grace's insect medium. After 3 days cultivation, GFP-positive plaques were picked and resuspended individually in 1 ml of Grace's insect medium. Each resuspended plaque was transferred into flasks with $10^5$ Sf9 cells to be infected. Infection progress was monitored by GFP fluorescence. After 5 - 7 days cultivation, Sf9 cells and their supernatants were collected, aliquoted and kept at -80 °C as recombinant baculovirus stocks.

*Titration of recombinant baculoviruses by endpoint dilution assay.* Aliquots of recombinant baculovirus stocks were thawed and diluted from $10^{-1}$ to $10^{-8}$ in Grace's insect medium, and 100 µl of each virus dilution was pipetted into 96-well plates in quadruplicate. Subsequently, 6 x $10^4$ freshly harvested Sf9 cells/well were added. After 5 - 7 days incubation at 26 °C, the numbers of GFP-positive wells were counted and virus titers were calculated as endpoint dilution assay TCID50 [$TCID_{50} = D^{(n/p+0.5)}$ x 1/sample volume (ml); D = dilution factor; n = number of positive wells; p = number of parallel values].

*Infection of Sf9 cells with recombinant baculoviruses.* To obtain recombinant cytokines, Sf9 cells were infected in suspension (1 x $10^6$ /ml) or in T175 flasks with Insect-XPRESS medium. For suspension culture, 0.1 % Pluronic-F68 (Gibco) was added. For infection with recombinant baculoviruses encoding IL-15-RLI or IL-15La-RLI an MOI of 2 - 3 was used, while infection with recombinant baculovirus encoding IL-2 was carried out with an MOI of 0.5 in order to reduce aggregation events. After 4 - 6 days incubation at 26 °C, supernatants were collected, filtered through a 0.22 µm membrane and kept at 4 °C until further analysis or purification.

***Purification of recombinant cytokines from insect cell supernatants.***

Filtered supernatants from infected Sf9 cells were mixed with ANTI-FLAG M2 Affinity Gel (Sigma) at a ratio of 600:1 (e.g., 300 ml of supernatant was mixed with 0.5 ml of affinity gel). After overnight incubation at 4 °C, the mixtures were poured into 10 ml columns with 35 µm filter pore size (MoBiTec), washed with approximately 150 ml of TBS and eluted six times with 1 ml aliquots of 0.1 M glycine HCl, pH 3.5 into vials containing 20 µl of 1 M Tris, pH 8.0. Subsequently the buffer was exchanged and concentrated to 300 µl PBS (-) using Amicon Ultracel - 3K centrifugal filters. Protein concentrations were determined with Pierce BCA Protein Assay Kit (Thermo Fischer Scientific) according to the manufacturer's instructions. Amount of substance (mol) for each recombinant protein was calculated based on the protein amount in the preparations and molecular weight. The molecular weight of recombinant proteins was estimated based on their amino acid sequences using Compute pI/Mw tool (https://web.expasy.org/compute_pi/). Purified recombinant proteins were analyzed immediately by Western blotting or gel filtration chromatography, or they were supplemented with 0.1 % Bovine serum albumin (BSA) and 50 % glycerol (recombinant protein storage buffer) for storage at -20 °C.

***Gel filtration chromatography.***

To analyze purified recombinant proteins, gel filtration chromatography was carried out on a Superose 12 column (30 cm length, 1 cm diameter, Pharmacia) using a HPLC system (BT 9200 Titan pump, BT 9520 IN UV monitor, Eppendorf Biotronik) for solvent delivery and UV monitoring at 280 nm. For the equilibration with PBS (-) and for the separation of proteins, a constant flow rate of 0.5 ml/min was maintained throughout the experiment. Samples of recombinant proteins containing between 10 - 15 µg were diluted in 0.5 ml PBS (-), injected using a 0.5 ml sample loop (Rheodyne), and fractionated (1 fraction/0.5 ml). The fractionated samples corresponding to peaks in the chromatogram were concentrated to 40 - 50 µl by ultrafiltration with an Amicon Ultracel - 3K filter and were analyzed by SDS-PAGE and Western blotting. The calibration of the column was carried out using standard procedures and the same chromatographic conditions as described for the recombinant proteins. The void volume was estimated with Blue dextrane (Sigma). For the calibration, elution volumes of calibrant proteins (Sigma) were plotted against the decadic logarithms of their molecular weights and linear regression models were calculated which were then used for the determination of molecular weights of the recombinant proteins.

***Deglycosylation assay.***

PNGase-F digestion of the lysates of transfected HEK293T cells and purified recombinant cytokine preparations was performed using the PNGase-F kit (New England Biolabs) with similar procedures as described in our previous study (18). Prior to assay, the concentration of purified recombinant protein preparations in PBS (-) was adjusted to approximately 500 ng/30 μl. Nine μl of the cell lysates or the recombinant cytokines were subjected to assay as suggested by the manufacturer, and incubated in the presence or absence (mock control) of PNGase-F. Digested samples were subjected to buffer exchange to PBS (-) and concentrated to 40 - 50 μl by ultrafiltration with an Amicon Ultracel - 3K filter. SDS-PAGE and Western blotting analysis were performed as described above except for using 16 % poly-acrylamide gels for analysis of the purified cytokines [Figs. S4D(e), S4E(e) and S4F(e)].

***Establishment of a monoclonal antibody (mAb) against rainbow trout CD8α.***

Although an anti-trout CD8α mAb had already been established by our group (20), a new mAb clone named 7α8c with a different immunoglobulin isotype was established enabling multi-color immunostaining with mAbs against other molecules. MAb 7α8c was established as previously described (21, 22) with slight modifications as follows: Rats were immunized twice into the tail base with Normal Rat Kidney cells expressing trout CD8α (20) emulsified in Complete Freund's Adjuvant (Sigma). Hybridomas were cloned twice by limiting dilution, and one of the resulting clones, designated as 7α8c, was selected for further experiments. Supernatants were stored as 50 % glycerol stocks at -20 °C. For validation of mAb specificity, HEK293T cells expressing trout CD8α-HA (established by Takizawa *et al.*, unpublished) were tested for the reactivity with mAb 7α8c by flow cytometry using FACSCanto II (BD Biosciences). As a positive expression control, an anti-HA mAb was applied. Anti-rat IgG Alexa Fluor 488 (Thermo Fisher scientific) and anti-Mouse IgG, IgM (H+L) Alexa Fluor 488 Secondary Antibody were used as secondary conjugates, respectively. To further prove the specificity of mAb 7α8c, $mAb^+$ and $mAb^-$ lymphocyte subpopulations were flow sorted from trout intestine as described in the next paragraph. Total RNA was extracted from the subpopulations using NucleoSpin RNA kit (Macherey-Nagel), and subjected to semi-quantitative one step RT-PCR analysis with primers for *β-actin* and *CD8α* (Table of primer sequences B) as described previously (20). The cycle numbers used for *β-actin* and *CD8α* were 25 cycles and 37 cycles, respectively.

***Isolation of rainbow trout lymphocyte subpopulations.***

Leukocytes from trout thymus, gill, head kidney (HK), spleen and intestine were isolated as described previously (20). Briefly, the cell suspensions were layered onto an isotonic Percoll (GE Healthcare) gradient (r = 1.075 g/ml) and centrifuged at 650 x g for 40 min. After centrifugation, cells lying at the interface were collected and washed twice with cold mixed medium (MM): Iscove's DMEM/Ham's F12 (Gibco) at a ratio of 1:1, supplemented with 10 % fetal bovine serum (FBS) and 100 U penicillin/ml and 100 μg streptomycin/ml. If leukocytes isolation was followed by flow sorting, leukocytes from 4 - 8 clonal individuals were pooled to get sufficient amounts of lymphocytes (Fig. S3). The cell suspensions were kept on ice until further preparation.

To isolate $CD8\alpha^+$ and $CD8\alpha^-$ lymphocytes, leukocytes from thymus, gill, HK, spleen and intestine were stained with anti-CD8α (clone 13.2D; rat IgG2a isotype) (18) and anti-rat IgG Alexa Fluor 488. To isolate CD8 single positive [SP], CD4SP, double positive [DP] and double negative [DN] lymphocytes, leukocytes from thymus, spleen and intestine were stained with anti-CD4-1 (rat IgG2a isotype) (22), anti-CD4-2 (rat IgG2b isotype) (22) and anti-CD8α (clone 7α8c; rat IgG1 isotype [Fig. S3A]) mAbs. Stained cells were detected with anti-rat IgG2a-PE (eBioscience), anti-rat IgG2b-PE (eBioscience) and anti-rat IgG1-FITC conjugates (BD Bioscience), respectively. To isolate CD8SP, CD4SP, IgMSP and triple negative [TN] lymphocytes, splenocytes were stained with anti-CD4-1, anti-CD4-2 and anti-CD8α mAbs as described above in addition to anti-IgM (mouse IgG1 isotype) mAb (23). Stained cells were detected with anti-rat IgG2a-eFluor 660 (eBioscience), anti-rat IgG2b-eFluor 660 (eBioscience), anti-rat IgG1-FITC, and anti-mouse IgG1-Brilliant Violet 421™ conjugates (Biolegend), respectively.

Cell suspensions were incubated on ice with mAbs and corresponding secondary conjugates for 30 min and then washed twice with MM after each respective staining steps. For negative controls, conjugate controls were prepared as described above. Doublets were excluded by FSC-A/FSC-H gating, and dead cells were excluded by DAPI (4', 6-diamidino-2-phenylindole)- or PI- staining. Secondary antibodies were tested before the experiments to exclude possible cross-reactions between isotype/species specific conjugates. Stained cells were sorted using a BD FACSAria™ Fusion flow cytometer (BD Biosciences). For flow sorting, only “lymphocyte gate” cells ($FSC^{low}/SSC^{low}$; aka “morphological lymphocytes”) where considered. For sorting of IgMSP and TN lymphocytes, only the $CD4^-CD8^-$ DN population was considered. After all conditions were adjusted by setting the thresholds for conjugate controls and the compensation parameters, test sortings were performed for each of the populations to confirm their purity. The purity of each sorted lymphocyte subpopulation was at least 99.2 %. Data on flow cytometry were analyzed using FlowJo V10 software (Tree Star). Sorted-lymphocyte subpopulations were cultivated overnight with MM (containing 20 % FBS) at 15 °C under a 2.5 % $CO_2$ atmosphere, prior to the next experimental steps.

***Stimulation of trout leukocytes with recombinant cytokines.***

*Stimulation of trout leukocyte subpopulations and Western blot analysis.* $CD8^+$ and $CD8^-$ lymphocytes were isolated from trout thymus, gill, HK, spleen and intestine while CD8SP, CD4SP, DP and DN lymphocytes were isolated from trout thymus, spleen and intestine, as described above. In each experiment, equal numbers of lymphocyte subpopulations (1 - 3 x $10^5$/incubation, depending on the yields achieved after flow sorting) were resuspended in 400 µl of supernatants containing recombinant cytokines (1:1 dilution in MM) from transfected HEK293T or in 400 µl of purified recombinant proteins diluted in MM (5 nM, 25 nM and 125 nM) and incubated for 15 min at 15 °C. As negative controls, lymphocyte subpopulations were incubated with supernatants from mock-transfected HEK293T cells (1:1 dilution in MM) or protein storage buffer (see paragraph *Purification of recombinant cytokines with anti-FLAG agarose*) diluted with MM. After incubation, lymphocytes were pelleted at 1,500 x g for 5 min at 4 °C, and lysed with 17 µl of NP40 Cell Lysis Buffer supplied with Protease Inhibitor Cocktail and Halt Phosphatase Inhibitor Cocktail (Thermo Fisher Scientific). Then the samples were subjected to Western blot analysis for detection of phosphorylated STAT5 similar to as described above except for using 8 % poly-acrylamide gels.

*Stimulation of trout leukocytes and RT-qPCR analysis.* Trout total splenocytes were freshly isolated as described above. Samples were used per individual fish in "total splenocytes" stimulation experiments, but pooled from multiple clonal fish prior to flow sorting into CD8SP, CD4SP, IgMSP and TN lymphocytes. Total splenocytes were incubated for 4 h or 12 h with 400 µl of supernatants containing recombinant cytokines (1:1 dilution in MM) from transfected HEK293T cells or with 400 µl of purified recombinant proteins diluted in MM (5 nM, 25 nM and 125 nM). The subpopulations were incubated for 12 h with 400 µl of purified recombinant proteins diluted in MM (0.2 nM and 5 nM). For negative controls, total splenocytes and their subpopulations were mock-treated as described above. In each experiment, equal numbers of total splenocytes (2 x $10^5$) or lymphocyte subpopulations (1 - 3 x $10^5$) were incubated. After incubation at 15 °C, cells were pelleted as described above, followed by treatment with 350 ul of Lysis Buffer RA1 (supplied by NucleoSpin RNA kit) containing 1/100 2-mercaptoethanol.

Total RNA was extracted from the incubated cells using NucleoSpin RNA kit and aliquots corresponding to 0.5 - 1.6 x $10^5$ cells were reverse transcribed into cDNA using SensiFAST™ cDNA Synthesis Kit (Bioline) according to the manufacturer's instruction. The resulting cDNA was diluted 1:6 with distilled water. Five µl of the resultant cDNA was used for qPCR detection of expression of *IFNγ, Perforin, IL-4/13A, IL-4/13B1, IL-4/13B2* and *EF1A* using primer sets Trout_IFNγ (1/2)_qPCR, Trout_PFN1_qPCR, Trout_IL-4/13A_qPCR, Trout_IL-4/13B1_qPCR, Trout_IL-4/13B2_qPCR and Trout_EF1A_qPCR

(Table of primer sequences B). qPCRs were performed using SensiFAST™ SYBR Lo-ROX Kit (Bioline) with Stratagene Mx3000P and MxPro software version 4.10 (Agilent Technologies). The comparative quantitation mode was chosen with default settings except for the amplification conditions (120 s at 95 °C, followed by 40 cycles of 5 s at 95 °C, 11 s at 60 °C and 15 s at 72 °C). After amplification, melting curve analysis of PCR products was performed in order to exclude the presence of unspecific products or primer dimer synthesis. All reactions were done in duplicate for technical replication. Based on the two replicates, the Ct values were calculated automatically by MxPro software, and used for analysis. The relative expression levels in the lymphocytes were normalized to the expression levels of *EF1A* using the equation $2^{-\Delta\Delta Ct}$ (24). For the graphs in Figs. 10 and S6A, the *EF1A*-normalized gene expression levels were normalized to those of the relevant control samples (4 h or 12 h) in the same experiment which were set as 1, while such normalization was applied for the graphs in Figs. 11 and S6B by setting each TN control sample as the standard. This method was chosen because it focuses on the fold-differences induced within cell samples, and in the case of Fig. 11 also allows an instantaneous impression of the differences in expression levels between cell populations.

***Statistics.***

For the statistical evaluation of the qPCR data, the paired samples t-test was applied to the log-adjusted *EF1A*-normalized gene expression levels calculated for biological quadruplicates using IBM SPSS Statistics 25 Software (16). Calculated p-values $< 0.05$ between cytokine-treated and mock-treated samples of the same cell populations were considered to be significant.

**Table for primers sequences**

A. Primer sequences used for molecular cloning of trout *IL-15La* and *IL-15Lb*

| Primer name | Primer Sequence (5'-3') | Use | |
|---|---|---|---|
| Trout_IL-15La_CDS_F | GCTACTTCTCTACTGATCTAAAGCTC | amplification of full-length trout *IL-15La* ORF | |
| Trout_IL-15La_CDS_R | TGTCCCTGTCTATGGATGTGTCC | | |
| Trout_IL-15Lb_CDS_F | CTGCAGATGACATGGACAAGAG | amplification of full-length trout *IL-15Lb* ORF | |
| Trout_IL-15Lb_CDS_R | TCTGCTCTTATGTGAGTCCTTG | | |
| Trout_IL-15La_5'RACE_R | CAGTTGGCACCTTCTTTCTGTTTGATCAAGGACTTCAGC | 5'-RACE amplification of trout *IL-15La* | First PCR |
| Trout_IL-15La_5'RACE.nes_R | GATAATCAGCCAAAGTTGGGGTGTACAG | | Nested PCR |
| Trout_IL-15Lb_5'RACE_R | TCCAGAATCCCTAGTAATGTTGTTAAGAAATCATTTGCTGCCTG | 5'-RACE amplification of trout *IL-15Lb* | First PCR |
| Trout_IL-15Lb_5'RACE.nes_R | GTCGTAAAGCATGTGAGTGTGGACC | | Nested PCR |

Table for primers sequences *continued*

B. Primer sequences used for gene expression analysis

| Primer name | Primer Sequence (5'-3') | Use | Accession number |
|---|---|---|---|
| Trout_IL-15La_F<br>Trout_IL-15La_R | CATGTGCAGTAAAGAACTTCCCGG<br>TTTGGCTTGTATATGGGTGAGGACTTA | Semi-quantitative PCR for trout *IL-15La* | MK619679 |
| Trout_IL-15Lb_F<br>Trout_IL-15Lb_R | AAGGAGGAAGTTCACAGGATGGAATC<br>AGTAATGTTGTTAAGAAATCATTTGCTGCCTG | Semi-quantitative PCR for trout *IL-15Lb* | MK619680 |
| Trout_CD8α_F<br>Trout_CD8α-R | ATGAAAATGGTCCAAAAGTGGATGC<br>GGTTAGAAAAGTCTGTTGTTGGCTATAGG | Semi-quantitative PCR for trout *CD8α* | NM_001124263 |
| Trout_β-actin_F<br>Trout_β-actin_R | GCTGTCTTCCCCTCCATCGTC<br>GGCAGGGGTGTTGAAGGTCTC | Semi-quantitative PCR for trout *β-actin* | AF157514 |
| Trout_EF1A_F<br>Trout_EF1A_R | CCACAGGCCATCTGATCTACA<br>TGAGCTGTTTCACTCCCAGAGTGTAG | Semi-quantitative PCR for trout *EF1A* | NM_001124339 |
| Trout_IL-15La_qPCR_F<br>Trout_IL-15La_qPCR_R | TTGAGGAAGTTCACAAGATGGAATCAT<br>GTTTTGGCTTGTATATGGGTGAGGACTT | Real-time qPCR for trout *IL-15La* | MK619679 |
| Trout_IL-15Lb_qPCR_F<br>Trout_IL-15Lb_qPCR_R | GGAGGAAGTTCACAGGATGGAATCAT<br>GTCTTTTCTGGTGTAAGGATGAAGAACG | Real-time qPCR for trout *IL-15Lb* | MK619680 |
| Trout_IFNγ (1/2)_qPCR_F<br>Trout_IFNγ (1/2)_qPCR_R | AAACATAGACAAACTGAAAGTCCA<br>CGTCCAGAACCACACTCATCA | Real-time qPCR for trout *IFNγ (1/2)* | FJ184374<br>FJ184375 |
| Trout_PFN1_qPCR_F<br>Trout_PFN1_qPCR_R | CTTTGGCACGCATTACATTACCA<br>AGCTGGCCTTGCTCTCTGTCTTA | Real-time qPCR for trout *Perforin-1* | NM_001134847 |
| Trout_IL-4/13A_qPCR_F<br>Trout_IL-4/13A_qPCR_R | CACCACAAAGTGCAAGGAGTT<br>TGGTCTTGGCTCTTCACAACG | Real-time qPCR for trout *IL-4/13A* | AB574337 |
| Trout_IL-4/13B1_qPCR_F<br>Trout_IL-4/13B1_qPCR_R | GAGATTCATCTACTGCAGAGGATCATGA<br>GCAGTTGGAAGGGTGAAGCTTATTGTA | Real-time qPCR for trout *IL-4/13B1* | HG794522 |
| Trout_IL-4/13B2_qPCR_F<br>Trout_IL-4/13B2_qPCR_R | GAGACTCATCTATTGCGTATGATCATCG<br>TGCAGTTGGTTGGATGAAACTTATTGTA | Real-time qPCR for trout *IL-4/13B2* | HG794523 |
| Trout_EF1A_qPCR_F<br>Trout_EF1A_qPCR_R | CAAGGATATCCGTCGTGGCA<br>ACAGCGAAACGACCAAGAGG | Real-time qPCR for trout *EF1A* | NM_001124339 |

#### Supplementary figure 1 (Fig. S1)

Additional information on *IL-15L* nucleotide and deduced amino acid sequences

**Table of Contents** **Page**


Fig. S1A: Trout *IL-15La* cDNA sequence 20

Fig. S 1B: Trout *IL-15Lb* cDNA sequence 22

Fig. S1C: Expression levels of rainbow trout and Atlantic salmon *IL-15La* and *IL-15Lb* transcripts in various tissues 24

Fig. S1D: Rainbow trout *IL-15La* and *IL-15Lb* probably derived from a gene duplication in the salmonid lineage 27

#### **Figure S1A.** Trout *IL-15La* cDNA sequence.

Figure S1A-2 shows an assembly of two sequences obtained from spleen cDNA. One sequence contained the entire coding sequence and was amplified by primers Trout_IL-15La_CDS_F and Trout_IL-15La_CDS_R. The other sequence was obtained by nested 5'-RACE, using SMARTer RACE cDNA amplification kit (Clontech) and two PCR reactions; for the first PCR reaction the gene-specific primer Trout_IL-15La_5'RACE_R was used in conjunction with the Clontech primer UPM, which was followed by diluting the mixture and a second PCR reaction using the gene-specific primer Trout-IL-15La_5'RACE.nes_R in conjunction with the Clontech primer NUP. The 5'-RACE result is shown in Fig. S1A-1, with the arrow pointing at the analyzed band. Other than database comparisons (see main text Table 1), no efforts were dedicated to potentially finding alternative transcripts from the same gene. Primer sequences, or their complementary sequences, are underlined in Fig. S1A-2, with the primer direction indicated besides the primer name. The intron positions were determined by comparison with the genomic sequence of GenBank accession MSJN01002393, and are indicated by downward triangles. ATG motifs in the 5'UTR region are shaded yellow. Amino acids are indicated below the first nucleotides of codons.

*Fig. S1A-1.*
*Left lane is size marker in bp units. Right lane is nested 5'-RACE result for trout IL-15La.*

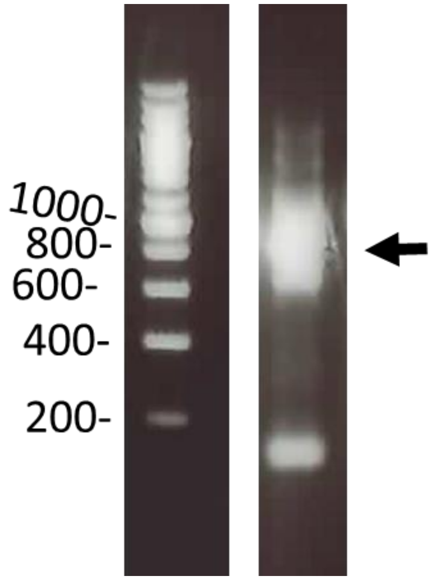

*Fig. S1A-2.*
*Assembled sequence of trout IL-15La ORF and 5’-RACE product.*

```
        10        20        30        40        50        60        70        80        90
GTGCACTAATAAAGAACTGCGGTTGTGTTCGGGTTGGACTCTCTAGGCTACACTCTTGAGTGTAAATGAGCATTTGAAATGTCAGGATTT

       100       110       120       130       140       150       160       170       180
TGTTGCAGTACAAATATTATAGATTTTAGCTACACAATTTGAACTTCAACTGTGGATTTTAAAGTATCCTATGTCAAGTCATCTGCTTCT

       190       200       210       220       230       240       250       260       270
TGACGGAATTACTTTTGTTTTCACTTTGTACCTACATACGGTGCAAACTTCAGTAGTCTTTCAAGCTGATTTACCATGTCCGGTGAGAAG

       280       290       300       310       320       330       340       350       360
CCTAGACAAGAAAAAGGAGGCAGATGCTGACGGCTTGCCTCATCTCTTCTGCTGCGCTGCTTTCCAGGACACACAAGAGAGATGCTGCAT

       370       380       390       400       410       420       430       440       450
AACAAGTCACCAGGAAGATAGAAGACAGAGTGTTTTACCCCTCATCCAAAAGTGGGGAGCCTACTGGCTAAGAAGGAGGCACATAGCTGG

       460       470       480       490       500       510       520       530       540
AGTCTTGCCTCATCTCTACTGCTTCTTCCCAGGACACACAGGAGATGTAGTGTCACTAAGAAGGCTGCAATTGACATGGACAATATGCTG

       550       560       570       580       590       600       610       620       630
AATCCCAAAACAATGGAGGATTTTAAAATAGTTATCTTAATTTTAAGCTACTTCTCTACTGATCTAAAGCTCATGTGAAAGAGTGGGCCC
                                                   Trout_IL-15La_CDS_F >
       640       650       660       670       680       690       700       710       720
ATGCCATGCTGAGGAGACAGAGGACTGACACTCTTCTAGCCCTTTTGCTGTGGTTTCTCTTCTTCATCGCCATGACAATGAAACAGGCAT
     M  L  R  R  Q  R  T  D  T  L  L  A  L  L  L  W  F  L  F  F  I  A  M  T  M  K  Q  A  Y

       730       740       750       760       770       780       790       800       810
ATGGAAAATCCATGTGCAGTAAAGAACTTCCCGGAATTGTGCGAAAATGCATTGAGGAAGTTCACAAGATGGAATCATTTGATTGCAGAC
  G  K  S  M  C  S  K  E  L  P  G  I  V  R  K  C  I  E  E  V  H  K  M  E  S  F  D  C  R  L

       820       830       840       850       860       870       880       890       900
TGTACACCCCAACTTTGGCTGATTATCAGAAGTGCCCCACGTCCACACTCATATGCTTTGAAAAAGAAGTGAATGTCCTGGTGTTAGAAT
  Y  T  P  T  L  A  D  Y  Q  K  C  P  T  S  T  L  I  C  F  E  K  E  V  N  V  L  V  L  E  S
< Trout_IL-15La_5’RACE.nes_R
       910       920       930       940       950       960       970       980       990
CTGGGAATAAGTCCTCACCCATATACAAGCCAAAACTATCCATCCGGCTGAAGTCCTTGATCAAACAGAAAGAAGGTGCCAACTGTCCAG
  G  N  K  S  S  P  I  Y  K  P  K  L  S  I  R  L  K  S  L  I  K  Q  K  E  G  A  N  C  P  D
                                              < Trout_IL-15La_5’RACE_R
      1000      1010      1020      1030      1040      1050      1060      1070      1080
ACTGTGAGGCCCACAGAGAAAGGGCAGCAAAGGATTTCTTAACAACATTGCAAACAATTCTGGAGTGGATGAACGATCAGGGGTGTCGGA
  C  E  A  H  R  E  R  A  A  K  D  F  L  T  T  L  Q  T  I  L  E  W  M  N  D  Q  G  C  R  K

      1090      1100      1110      1120
AGCCATCCAGCCATTGAGATGGACACATCCATAGACAGGGACA
  P  S  S  H  *
                            < Trout_IL-15La_CDS_R
```

#### **Figure S1B.** Trout *IL-15Lb* cDNA sequence.

Figure S1B-2 shows an assembly of two sequences obtained from spleen cDNA. One sequence contained the entire coding sequence and was amplified by primers Trout_IL-15Lb_CDS_F and Trout_IL-15Lb_CDS_R. The other sequence was obtained by nested 5'-RACE, using SMARTer RACE cDNA amplification kit (Clontech) and two PCR reactions; for the first PCR reaction the gene-specific primer Trout_IL-15Lb_5'RACE_R was used in conjunction with the Clontech primer UPM, which was followed by diluting the mixture and a second PCR reaction using the gene-specific primer Trout-IL-15Lb_5'RACE.nes_R in conjunction with the Clontech primer NUP. The 5'-RACE result is shown in Fig. S1B-1, with the arrow pointing at the analyzed band. Other than comparisons with database reports of *IL-15Lb* transcripts in related species (see main text Table 1), no efforts were dedicated to potentially finding alternative transcripts from the same gene. Primer sequences, or their complementary sequences, are underlined in Fig. S1B-2, with the primer direction indicated besides the primer name. The intron positions were determined by comparison with the genomic sequence of GenBank accession MSJN01001275, and are indicated by downward triangles. ATG motifs in the 5'UTR region are shaded yellow. Amino acids are indicated below the first nucleotides of codons.

*Fig. S1B-1.*
*Left lane is size marker in bp units. Right lane is the nested 5'-RACE result for trout IL-15Lb.*

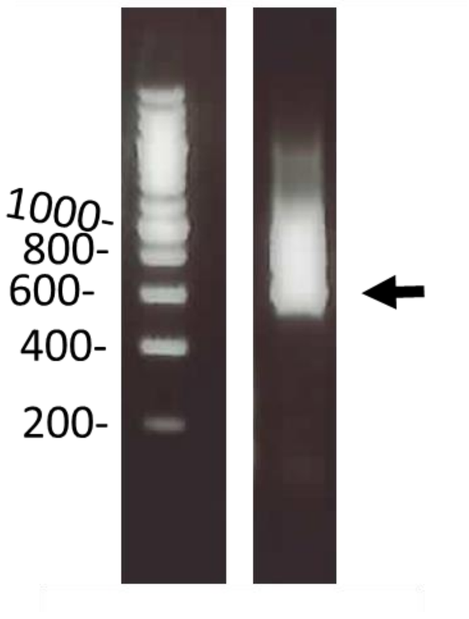

*Fig. S1B-2.*
*Assembled sequence of trout IL-15Lb ORF and 5’-RACE product.*

```
        10        20        30        40        50        60        70        80        90
GTTTTCACTTTGTACCTACCTACGATGCAAACTTCAAAAGTCGCTCAAGCTGATTTAACCATGCCAGGTGAGGAGTCTAGCCAAGAAAAA

       100       110       120       130       140       150       160       170       180
GGAGGCAGATGCTGAAGACGTGCCTAATTTCTTCTGCTGCTTTCCAGGACACACAAGAAAGAGGCTGCATAGACACACAGGGGATATACT

       190       200       210       220       230       240       250       260       270
GTGTCACAAAGAAGGCTGCAGATGACATGGACAAGAGGCTGAACCCCAAAACAATGGAGGATTTTAAACTAATTCTCCTAATTTAAGCTA
                Trout_IL-15Lb_CDS_F >
       280       290       300       310       320       330       340       350       360
TTTTCCCACTGATCTGAAGCTCCTGCGACAAAGTGGGCCCATGCCATGTTGAGGAGACAGAGAACTGGCTCTCTTCTGAACGCTTTGCTG
                                             M  L  R  R  Q  R  T  G  S  L  L  N  A  L  L

       370       380       390       400       410       420       430       440       450
TGGTTTCTCTTCTTCATTGCCATGACAATGAAACAGGCTTATGGACAATCCATTAGCAGTTCAGAAATTCACCAAATTGTGAAAACATTT
W  F  L  F  F  I  A  M  T  M  K  Q  A  Y  G  Q  S  I  S  S  S  E  I  H  Q  I  V  K  T  F

       460       470       480       490       500       510       520       530       540
ATTAAGGAGGAAGTTCACAGGATGGAATCATTTGATTGCAGACTGTACACCCCAACTTTAGCTGATTATAAGAAATGTCCCAGGTCCACA
I  K  E  E  V  H  R  M  E  S  F  D  C  R  L  Y  T  P  T  L  A  D  Y  K  K  C  P  R  S  T

       550       560       570       580       590       600       610       620       630
CTCACATGCTTTACGACAGAAGTAAAAGTCCTGATGTTAGAATATGGGAAACGTTCTTCATCCTTACACCAGAAAAGACTCACCAAACGA
L  T  C  F  T  T  E  V  K  V  L  M  L  E  Y  G  K  R  S  S  S  L  H  Q  K  R  L  T  K  R
< Trout_IL-15Lb_5’RACE.nes_R
       640       650       660       670       680       690       700       710       720
CTGACTAAATTGATGTCCTTGATAAAACAGAAGGATGGTGCCAACTGTCCACACTGTGAGGTCCACAGAGAACAGGCAGCAAATGATTTC
L  T  K  L  M  S  L  I  K  Q  K  D  G  A  N  C  P  H  C  E  V  H  R  E  Q  A  A  N  D  F

       730       740       750       760       770       780       790       800       810
TTAACAACATTACTAGGGATTCTGGAGTGGATGAACAATCAGGGGTCTCAGTTGCCAGACAGCCACTGAGATGTACACACCCATAGCCAG
L  T  T  L  L  G  I  L  E  W  M  N  N  Q  G  S  Q  L  P  D  S  H  *
 < Trout_IL-15Lb_5’RACE_R
       820       830       840
GGACATCCAAGGACTCACATAAGAGCAGA
        < Trout_IL15-Lb_CDS_R
```

**Figure S1C.** Expression levels of rainbow trout and Atlantic salmon *IL-15La* and *IL-15Lb* transcripts in various tissues.

In this figure, the results are shown for analyses of *IL-15La* and *IL-15Lb* expression levels in various tissues of rainbow trout and Atlantic salmon as determined by semi-quantitative RT-PCR (Fig. S1C-1) and by counting matches in single read archive (SRA) datasets (Fig. S1C-2). Despite variation between fish individuals, rather consistent findings were that: (i) *IL-15La* expression was more ubiquitously distributed than *IL-15Lb* expression; (ii) *IL-15Lb* expression was relatively high in gill; (iii) both *IL-15La* and *IL-15Lb* expression tended to be relatively low in head kidney. These findings agree with the RT-qPCR results shown in main text Fig. 2.

Detailed legends for Figs. S1C-1 and S1C-2 are:

Fig. S1C-1. *IL-15La and IL-15Lb expression analyzed for tissues of two trout individuals by semi-quantitative gene-specific RT-PCR.* The expected sizes of the amplified bands for *IL-15La*, *IL-15Lb* and *EF1A* were 204, 281, and 377 bp, respectively. In the photographs of the agarose gels after electrophoresis, the left lanes contain the size marker with sizes indicated in bp. The results for head kidney and gill are highlighted.

Fig. S1C-2. *IL-15La and IL-15Lb expression analyzed for tissues of three rainbow trout and two Atlantic salmon individuals by counting of gene-specific matches per $10^8$ database reads.* Read numbers per $10^8$ reads of *IL-15La* and *IL-15Lb* were determined by similarity searches against tissue-specific single read archive (SRA) datasets available at NCBI. For rainbow trout, the SRA datasets of Bioprojects PRJEB4450, PRJNA389609 and PRJNA380337 were investigated. For Atlantic salmon, the SRA datasets of Bioprojects PRJNA260929 and PRJNA72713 were investigated. Bioproject PRJEB4450 concerns tissues of a homozygous clonal 1-year-old female rainbow trout sampled 3 weeks after spawning (13). Bioproject PRJNA389609 concerns thirteen different tissues that were collected from a single immature (2-year old, 250 g) male homozygous rainbow trout of the Swanson clonal line, while the oocyte and pineal gland samples were pooled from multiple trout individuals (14). Bioproject PRJNA380337 concerns rainbow trout tissues for which detailed information has not been provided (authored by the Norwegian University of Life Sciences). Bioproject PRJNA260929 concerns tissues of a 1-year-old single homozygous female Atlantic salmon from the AquaGen aquaculture strain named “Sally” (25) (Dr. Unni Grimholt, personal communication). Bioproject PRJNA72713 concerns Atlantic salmon tissues for which detailed information has not been provided (authored by the University of Victoria). The results for the tissues head kidney and gill are highlighted.

*Fig. S1C-1.*

*IL-15La and IL-15Lb expression analyzed for tissues of two trout individuals by semi-quantitative gene-specific RT-PCR.*

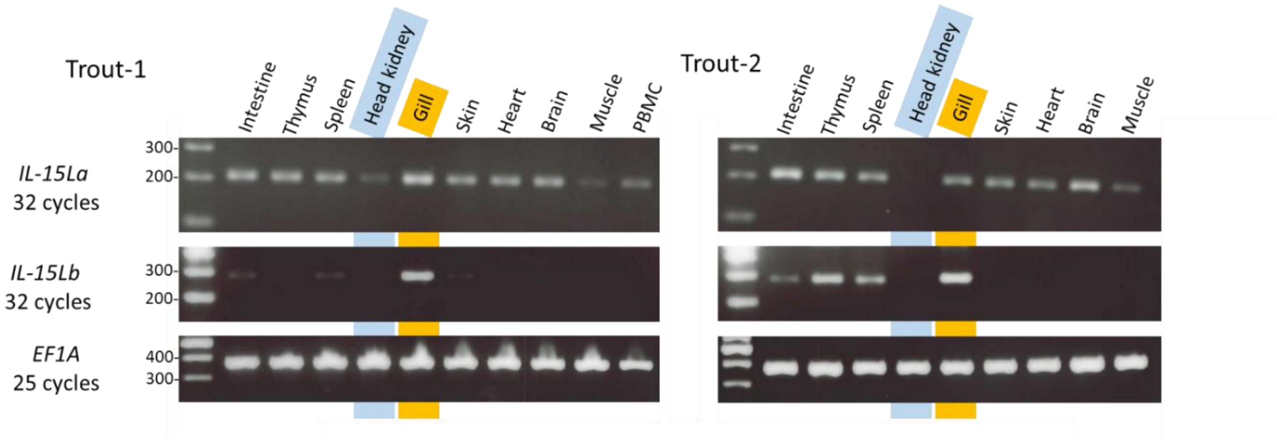

*Fig. S1C-2.*

*IL-15La and IL-15Lb expression analyzed for tissues of three rainbow trout and two Atlantic salmon individuals by counting of gene-specific matches per $10^8$ database reads.*

Matching reads / $10^8$ (100,000,000) total reads

Trout (Bioproject PRJEB4450)

| | Intestine | Spleen | Kidney | Head Kidney | Gill (branchiae) | Skin | Heart | Brain | White muscle | Red muscle | Pineal gland | Hypophysis | Liver | Stomach | Bone | Testis | Ovary | Eye | Pyloric caeca | Nose | Pancreas | Fat | Oocyte |
|---|---|---|---|---|---|---|---|---|---|---|---|---|---|---|---|---|---|---|---|---|---|---|
| IL-15La | 35 | 14 | 3 | 2 | 15 | 7 | 2 | 3 | 3 | 7 | N.A. | 0 | 23 | 8 | 5 | N.A. | 49 | N.A. | N.A. | N.A. | N.A. | N.A. | N.A. |
| IL-15Lb | 114 | 41 | 0 | 0 | 115 | 27 | 0 | 0 | 0 | 0 | N.A. | 0 | 3 | 9 | 25 | N.A. | 13 | N.A. | N.A. | N.A. | N.A. | N.A. | N.A. |

Trout (Bioproject PRJNA389609)

| | Intestine | Spleen | Kidney | Head Kidney | Gill | Skin | Heart | Brain | White muscle | Red muscle | Pineal gland | Hypophysis | Liver | Stomach | Bone | Testis | Ovary | Eye | Pyloric caeca | Nose | Pancreas | Fat | Oocyte |
|---|---|---|---|---|---|---|---|---|---|---|---|---|---|---|---|---|---|---|---|---|---|---|
| IL-15La | 38 | 17 | 22 | 17 | 55 | 9 | N.A. | 6 | 12 | 14 | 5 | N.A. | 13 | 7 | N.A. | 14 | N.A. | N.A. | N.A. | N.A. | N.A. | 13 | 3 |
| IL-15Lb | 74 | 14 | 2 | 1 | 35 | 3 | N.A. | 0 | 2 | 0 | 0 | N.A. | 0 | 3 | N.A. | 2 | N.A. | N.A. | N.A. | N.A. | N.A. | 0 | 0 |

Trout (Bioproject PRJNA380337)

| | Intestine | Spleen | Kidney | Head Kidney | Gill | Skin | Heart | Brain | Muscle | Pineal gland | Hypophysis | Liver (average of 1,2,3) | Stomach | Bone | Testis | Ovary | Eye | Pyloric caeca | Nose | Pancreas | Fat | Oocyte |
|---|---|---|---|---|---|---|---|---|---|---|---|---|---|---|---|---|---|---|---|---|---|
| IL-15La | 14 | 0 | 3 | 3 | 46 | 39 | 6 | 15 | 6 | N.A. | N.A. | 6 | N.A. | N.A. | N.A. | N.A. | 34 | 24 | N.A. | N.A. | N.A. | N.A. |
| IL-15Lb | 6 | 22 | 0 | 0 | 93 | 0 | 0 | 0 | 0 | N.A. | N.A. | 0 | N.A. | N.A. | N.A. | N.A. | 6 | 9 | N.A. | N.A. | N.A. | N.A. |

Salmon (Bioproject PRJNA260929)

| | Intestine | Spleen | Kidney | Head Kidney | Gill | Skin | Heart | Brain | Muscle | Pineal gland | Hypophysis | Liver | Stomach | Bone | Testis | Ovary | Eye | Pyloric caeca | Nose | Pancreas | Fat | Oocyte |
|---|---|---|---|---|---|---|---|---|---|---|---|---|---|---|---|---|---|---|---|---|---|
| IL-15La | 216 | 47 | N.A. | N.A. | 14 | 113 | 73 | 0 | 0 | N.A. | N.A. | 147 | N.A. | N.A. | N.A. | N.A. | N.A. | 95 | N.A. | 213 | N.A. | N.A. |
| IL-15Lb | 0 | 0 | N.A. | N.A. | 100 | 301 | 0 | 0 | 0 | N.A. | N.A. | 0 | N.A. | N.A. | N.A. | N.A. | N.A. | 354 | N.A. | 0 | N.A. | N.A. |

Salmon (Bioproject PRJNA72713)

| | Intestine | Spleen | Kidney | Head Kidney | Gill | Skin | Heart | Brain | Muscle | Pineal gland | Hypophysis | Liver | Stomach | Bone | Testis | Ovary | Eye | Pyloric caeca | Nose | Pancreas | Fat | Oocyte |
|---|---|---|---|---|---|---|---|---|---|---|---|---|---|---|---|---|---|---|---|---|---|
| IL-15La | 50 | 56 | 52 | 17 | N.A. | 12 | 65 | 10 | 49 | N.A. | N.A. | 167 | N.A. | N.A. | 23 | 7 | 0 | 192 | 67 | N.A. | N.A. | N.A. |
| IL-15Lb | 0 | 0 | 0 | 0 | N.A. | 0 | 3 | 0 | 0 | N.A. | N.A. | 0 | N.A. | N.A. | 8 | 0 | 0 | 0 | 30 | N.A. | N.A. | N.A. |

**Figure S1D.** Rainbow trout *IL-15La* and *IL-15Lb* probably derived from a gene duplication in the salmonid lineage.

The sequence divergence in the IL-2/15/15L family of cytokines is enormous and attempts to use phylogenetic tree software analysis to reliably determine the precise relationships between IL-15L and related cytokines were inconclusive [not shown and see (12)]. However, phylogenetic tree software does give reliable results when addressing more recent events in evolution, like for example the *IL-15L a-b* duplication observed in trout and salmon. Therefore, we only analyzed a selected group of molecules.

A tree was created using all the IL-15L amino acid sequences shown main text Fig. 3 with as outgroup human IL-15, all aligned as in main text Fig. 3, and only using the parts aligned with mature human IL-15. The analysis was conducted in MEGA6 (26) by UPGMA method (27). The percentage of replicate trees in which the associated taxa clustered together in the bootstrap test (10000 replicates) are shown next to the branches. The tree is drawn to scale, with branch lengths in the same units as those of the evolutionary distances used to infer the phylogenetic tree. The evolutionary distances were computed using the Poisson correction method and are in the units of the number of amino acid substitutions per site. All ambiguous positions were removed for each sequence pair. The important results are highlighted in blue and show that the salmonid *IL-15L a-b* duplication occurred after the ancestors of salmonids and zebrafish separated, and before the separation of the species lineages leading to rainbow trout and Atlantic salmon. This *IL-15L* duplication also included the neighboring genes (main text Fig. 1) and presumably was part of the whole genome duplication early in the salmonid lineage (13, 25, 28).

*Fig. S1D. Phylogenetic tree.*

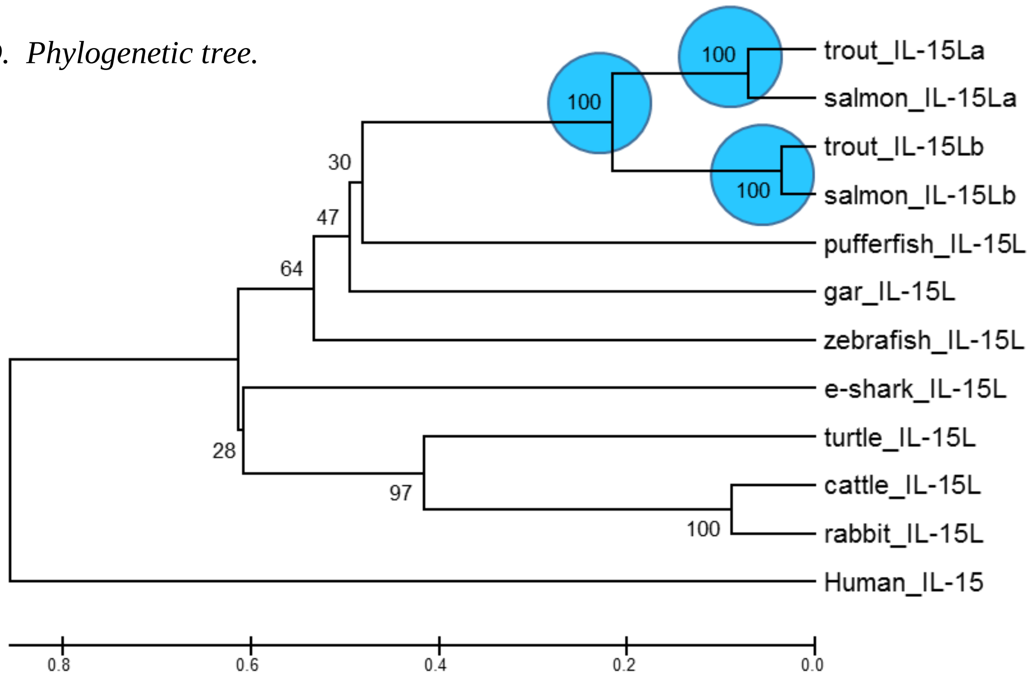

#### Supplementary figure 2 (Fig. S2)

Expression vectors used in this study

**Table of Contents** **Page**


Legend to Figs. S2A, S2B, S2C and S2D 30

Fig. S2A: Expression vectors for (modified) bovine cytokines and receptor chains 31

- pRcCMV2-*Bos-IL-2-FLAG* 31
- pRcCMV2-*Bos-IL-15-FLAG* 31
- pRcCMV2-*Bos-IL-15L-FLAG* 32
- pRcCMV2-*Bos-IL-15L-(non-tagged)* 32
- pRcCMV2-*Bos-IL-15Lhyb-FLAG* 33
- pcDNA3.1-*IL-2-Lead-RLI-bov.IL-15Lhyb* 33
- pcDNA3.1-*Bos-IL-15Rα-Myc-His* 34
- pcDNA3.1-*Bos-solIL-15Rα-Myc-His* 35
- pcDNA3.1-*Bos-IL-2Rα-Myc-His* 35
- pcDNA3.1-*Bos-solIL-2Rα-Myc-His* 36

Fig. S2B: Expression vector for a genetic fusion of human IL-15Rα and IL-15 37

- pcDNA3.1-*IL-2-Lead-RLI* 37

Fig. S2C: Expression vectors for (modified) trout cytokines and IL-15Rα 38

- pcDNA3.1-*trout-IL-2-FLAG* 38
- pcDNA3.1-*trout-IL-2-(non-tagged)* 38
- pcDNA3.1-*trout-IL-15-FLAG* 39
- pcDNA3.1-*trout-IL-15-(non-tagged)* 39
- pcDNA3.1-*trout-IL-15La-FLAG* 40
- pcDNA3.1-*trout-IL-15La-(non-tagged)* 40
- pcDNA3.1-*trout-IL-15Lb-FLAG* 41
- pcDNA3.1-*trout-IL-15Lb-(non-tagged)* 41
- pcDNA3.1-*IL-2-Lead-RLI-trout-IL-15La* 42
- pcDNA3.1-*trout-IL-15Rα-Myc* 42
- pcDNA3.1-*trout-solIL-15Rα-Myc* 43

Fig. S2D: Vectors for expressing (modified) trout cytokines and trout IL-15Rα in insect cells using a baculovirus system. 44
- pFBD-P10Uhis-ieGFP-*trout-IL-2-FLAG* 44
- pFBD-P10Uhis-ieGFP-*trout-IL-15-FLAG* 44
- pFBD-P10Uhis-ieGFP-*trout-IL-15La-FLAG* 45
- pFBD-P10Uhis-ieGFP-*trout-solIL-15Rα-Myc* 45
- pFBD-P10Uhis-ieGFP-*trout-IL-15-RLI* 46
- pFBD-P10Uhis-ieGFP-*trout-IL-15La-RLI* 46

#### Legend to Figs. S2A, S2B, S2C and S2D.

This figure shows the sequence fragments containing bovine (Fig. S2A), human (Fig. S2B), and trout (Fig. S2C) *IL-2*, *IL-15*, *IL-15L*, *IL-2Rα* and/or *IL-15Rα* sequences, or modifications thereof, which were cloned into DNA plasmid expression vectors behind a CMV immediate early promoter or into a baculovirus transfer vector behind a p10 promoter (Fig. S2D). The texts for the individual constructs explain wat the vectors encode, and how the encoded protein is named in the paper. The pink font sequence fragments indicate the restriction sites used for cloning. The DNA plasmid expression vector used for cloning was either pRc/CMV2 (Invitrogen) or pcDNA™3.1/myc-His B (Invitrogen). The baculovirus transfer vector used for cloning was pFBD-P10Uhis-ieGFP (a derivate of pFastBac-Dual [Invitrogen], see Text S1). The cloning strategies varied and were somewhat arbitrarily because experiments were done at different time points and constructs were made in different laboratories.

Several of the expression constructs with bovine *IL-2*, *IL-15L*, *IL-15Rα* and *IL-2Rα* were already described in detail in references (12) and (18), but for convenience are shown here again.

Because the information on the construction method is not so relevant, the below only briefly mentions whether the construction involved an amplification from cDNA or a purchase of a synthetic gene sequence (Invitrogen) and describes the GenBank accession number of a matching sequence. When relevant, additional information is given.

Green font nucleotide sequence: By choice of primer sequences, or synthetic gene ordering, an ACC motif was added directly upstream of the start codon to assure efficient translation.

Red font amino acid sequence: By choice of primer sequences, or synthetic gene ordering, FLAG tag sequences were added.

Orange font amino acid sequence: By choice of primer sequences, or as part of the commercial vector, Myc tag sequences were added.

Purple font amino acid sequence: As part of the commercial vector, poly-His tag sequences were added.

Other colors are explained in the individual texts.

**Figure S2A.**
**Expression vectors for (modified) bovine cytokines and receptor chains.**

#### pRcCMV2-*Bos-IL-2-FLAG*

This expression vector encodes bovine FLAG-tagged IL-2 (named IL-2).
Cloned from cDNA. The *IL-2* part of the sequence is identical with GenBank accession EU276068.
*[for cloning details see (12)]*

```
AAGCTTACCATGTACAAGATACAACTCTTGTCTTGCATTGCACTAACTCTTGCACTCGT
TGCAAACGGTGCACCTACTTCAAGCTCTACGGGGAACACAATGAAAGAAGTGAAGTCAT
TGCTGCTGGATTTACAGTTGCTTTTGGAGAAAGTTAAAAATCCTGAGAACCTCAAGCTC
TCCAGGATGCATACATTTGACTTTTACGTGCCCAAGGTTAACGCTACAGAATTGAAACA
TCTTAAGTGTTTACTAGAAGAACTCAAACTTCTAGAGGAAGTGCTAAATTTAGCTCCAA
GCAAAAACCTGAACCCCAGAGAGATCAAGGATTCAATGGACAATATCAAGAGAATCGTT
TTGGAACTACAGGGATCTGAAACAAGATTCACATGTGAATATGATGATGCAACAGTAAA
CGCTGTAGAATTTCTGAACAAATGGATTACCTTTTGTCAAAGCATCTACTCAACAATGA
CTGATTATAAGGATGACGACGATAAGTAAGGGCCC
```

```
MYKIQLLSCIALTLALVANGAPTSSSTGNTMKEVKSLLLDLQLLLEKVKNPENLKLSRM
HTFDFYVPKVNATELKHLKCLLEELKLLEEVLNLAPSKNLNPREIKDSMDNIKRIVLEL
QGSETRFTCEYDDATVNAVEFLNKWITFCQSIYSTMTDYKDDDDK
```

#### pRcCMV2-*Bos-IL-15-FLAG*

This expression vector encodes bovine FLAG-tagged IL-15 (named IL-15).
The sequence was commercially ordered (Invitrogen). The leader sequence is replaced with that of bovine IL-2 (shaded blue), and the codons were optimized for expression in human cells. The encoded IL-15 part is identical to GenBank accession ELR44697.

```
AAGCTTACCATGTACAAGATCCAGCTGCTGAGCTGTATCGCCCTGACCCTGGCCCTGGT
GGCCAACGGCAATTGGCAGTACGTGATCAACGACCTGAAAACCATCGAGCACCTGATCC
AGAGCATCCACATGGACGCCACCCTGTACACCGAGAGCGACGCCCACCCCAACTGCAAA
GTGACCGCCATGCAGTGCTTTCTGCTGGAACTGAGAGTGATCCTGCACGAGAGCAAGAA
CGCCACCATCTACGAGATCATCGAGAATCTGACCATGCTGGCCAACAGCAACCTGAGCA
GCATCGAGAACAAGACCGAGCTGGGCTGCAAAGAGTGCGAGGAACTGGAAGAGAAGTCC
ATCAAAGAGTTTCTGAAGTCCTTCGTGCACATCGTGCAGATGTTCATCAACACCAGCGA
CTACAAGGACGACGACGACAAGTGATCTAGAGGGCCC
```

```
MYKIQLLSCIALTLALVANGNWQYVINDLKTIEHLIQSIHMDATLYTESDAHPNCKVTA
MQCFLLELRVILHESKNATIYEIIENLTMLANSNLSSIENKTELGCKECEELEEKSIKE
FLKSFVHIVQMFINTSDYKDDDDK
```

#### pRcCMV2-*Bos-IL-15L-FLAG*

This expression vector encodes bovine FLAG-tagged IL-15L (named IL-15L).
Cloned from cDNA. The *IL-15L* part of the sequence is identical with GenBank accession NM_001301213.
*[for cloning details see (12)]*

```
AAGCTTACCATGTGGCTTCTCTGGACCACCCTCCTGCTGGTGCTGCCCTTGGGAGGCCT
AGGACCACTCCTCTGCCCAAGGGAGCCTTTCTACTTCCTCATTGCCATCACGAAGATGC
TGGAAAACAAAAATGATGGCAGTCTGTACACCCCAGATAATCTATTGGTGTGTCCTGCT
GAGACTCTCCGATGCTTCCGGCTGGAGTTGTCTGTGATCGGGTTTGAGGAGGGCCCATC
CGTGGGGATCGTTGTGTTCCGCCTACAGCGCCTACTGGATGCCCTGGGGTCCCAGCTGT
GGGTGATTGATCAGGGCCCTTGTCCACCCTGCGAAGGACACCCTCAGAGACCAGTCCCT
CTTTTTCTGGCCAAACTCTTGGAGTTATTACAGGGGACTTGTGCTCGGGACCTGCCCTC
AGCAGATTACAAGGATGACGACGATAAGTAATCTAGA
```

```
MWLLWTTLLLVLPLGGLGPLLCPREPFYFLIAITKMLENKNDGSLYTPDNLLVCPAETL
RCFRLELSVIGFEEGPSVGIVVFRLQRLLDALGSQLWVIDQGPCPPCEGHPQRPVPLFL
AKLLELLQGTCARDLPSADYKDDDDK
```

#### pRcCMV2-*Bos-IL-15L-(non-tagged)*

This expression vector encodes bovine non-tagged IL-15L [named IL-15L(N)].
Cloned from cDNA. The *IL-15L* sequence is identical with GenBank accession NM_001301213.
*[for cloning details see (12)]*

```
AAGCTTACCATGTGGCTTCTCTGGACCACCCTCCTGCTGGTGCTGCCCTTGGGAGGCCT
AGGACCACTCCTCTGCCCAAGGGAGCCTTTCTACTTCCTCATTGCCATCACGAAGATGC
TGGAAAACAAAAATGATGGCAGTCTGTACACCCCAGATAATCTATTGGTGTGTCCTGCT
GAGACTCTCCGATGCTTCCGGCTGGAGTTGTCTGTGATCGGGTTTGAGGAGGGCCCATC
CGTGGGGATCGTTGTGTTCCGCCTACAGCGCCTACTGGATGCCCTGGGGTCCCAGCTGT
GGGTGATTGATCAGGGCCCTTGTCCACCCTGCGAAGGACACCCTCAGAGACCAGTCCCT
CTTTTTCTGGCCAAACTCTTGGAGTTATTACAGGGGACTTGTGCTCGGGACCTGCCCTC
AGCATAATCTAGA
```

```
MWLLWTTLLLVLPLGGLGPLLCPREPFYFLIAITKMLENKNDGSLYTPDNLLVCPAETL
RCFRLELSVIGFEEGPSVGIVVFRLQRLLDALGSQLWVIDQGPCPPCEGHPQRPVPLFL
AKLLELLQGTCARDLPSA
```

#### pRcCMV2-*Bos-IL-15Lhyb-FLAG*

This expression vector encodes bovine FLAG-tagged IL-15L in which a few motifs have been replaced for that of other IL-2/15/15L family members (named IL-15Lhyb).
The part encoding the mature protein was commercially ordered (Invitrogen) with the codons optimized for expression in insect cells, and fused to sequences encoding authentic bovine IL-15L leader sequence and a C-terminal FLAG-tag by using appropriate primers and overlap PCR. The rationale behind the substitutions (shaded yellow) was that *in vitro* handling of bovine IL-15L protein revealed instability (not shown), while two of the modified motifs were different in mammals from IL-2/15/15L family consensus, and we thought that adding of the trout IL-15La glycosylation motif might aid stability. Unfortunately, an improved stability, or a function, was not found for this IL-15Lhyb molecule.

```
AAGCTTACCATGTGGCTTCTCTGGACCACCCTCCTGCTGGTGCTGCCCTTGGGAGGCCT
AGGACCTTTGTTGTGCCCCCGTGAACCCTTCTACTTCCTGCGCAAGATCACCAAGATGC
TCGAGAACAAGAACGACGGTTCCCTGTACACCCCCGACAACCTGCTCGTGTGCCCCGCT
GAAACCCTGCGCTGCTTCCGTCTGGAACTGTCCGTGATCGGTTTCGAGGAAGGCAACAA
GTCCTCCGTGGGTATCGTGGTGTTCAGGCTGCAGCGTCTGCTGGACGCTCTGGGTTCTC
AGCTGTGGGTCACCGACCAGGGACCTTGCCCTCCTTGCGAGGGTCACCCTCAACGTCCC
GTGCCTCTGTTCCTGGCTAAGCTGCTCGAGCTGCTGCAGGGAACCTGCGCTCGTGACCT
GCCCTCCGCTGATTACAAGGATGACGACGATAAGTAATCTAGA
```

```
MWLLWTTLLLVLPLGGLGPLLCPREPFYFLRKITKMLENKNDGSLYTPDNLLVCPAETL
RCFRLELSVIGFEEGNKSSVGIVVFRLQRLLDALGSQLWVTDQGPCPPCEGHPQRPVPL
FLAKLLELLQGTCARDLPSADYKDDDDK
```

#### pcDNA3.1-*IL-2-Lead-RLI-bov.IL-15Lhyb*

This expression vector is based on the RLI construct described by Mortier *et al.* 2006 (29), and encodes a genetic fusion of a human IL-2 leader, a FLAG-tag, a large part of the human IL-15Rα ectodomain, a glycine-serine linker (shaded blue), and the above described bovine IL-15Lhyb sequence (with the modified motifs shaded yellow) (named bov.IL-15Lhyb-h-RLI).
IL-15Rα does not interact with the signaling receptor IL-2Rβ·IL-2Rγ, and is thought to fixate IL-15 into the proper conformation for binding IL-2Rβ·IL-2Rγ (30); given the observed cross-species activity (main text Fig. 5), we speculated that RLI-versions of bovine and trout IL-15L would have similar stability and signaling advantages as observed for all-human RLI (29, 31, 32).
The linker plus IL-15Lhyb part were commercially ordered (Invitrogen) with the codons optimized for expression in insect cells, and fused to the N-terminal coding part by using plasmid pcDNA3.1-*IL-2-Lead-RLI*, appropriate primers and overlap PCR.

GGTACCACCATGTACAGGATGCAACTCCTGTCTTGCATTGCACTAAGTCTTGCACTTGT
CACAAACAGTGACTACAAGGATGACGATGACAAGATAGAAGGTAGGATCACATGCCCTC
CCCCCATGTCCGTGGAACACGCAGACATCTGGGTCAAGAGCTACAGCTTGTACTCCAGG
GAGCGGTACATTTGTAACTCTGGTTTCAAGCGTAAAGCCGGCACGTCCAGCCTGACGGA
GTGCGTGTTGAACAAGGCCACGAATGTCGCCCACTGGACAACCCCCAGTCTCAAATGCA
TTAGAGACCCTGCCCTGGTTCACCAAAGGCCAGCGCCACCCTCTGGAGGCTCCGGTGGA
GGTGGGAGTGGCGGTGGATCCGGTGGCGGAGGCAGCCCTTTGTTGTGCCCCCGTGAACC
CTTCTACTTCCTGCGCAAGATCACCAAGATGCTCGAGAACAAGAACGACGGTTCCCTGT
ACACCCCCGACAACCTGCTCGTGTGCCCCGCTGAAACCCTGCGCTGCTTCCGTCTGGAA
CTGTCCGTGATCGGTTTCGAGGAAGGCAACAAGTCCTCCGTGGGTATCGTGGTGTTCAG
GCTGCAGCGTCTGCTGGACGCTCTGGGTTCTCAGCTGTGGGTCACCGACCAGGGACCTT
GCCCTCCTTGCGAGGGTCACCCTCAACGTCCCGTGCCTCTGTTCCTGGCTAAGCTGCTC
GAGCTGCTGCAGGGAACCTGCGCTCGTGACCTGCCCTCCGCTTAATCTAGA

MYRMQLLSCIALSLALVTNSDYKDDDDKIEGRITCPPPMSVEHADIWVKSYSLYSRERY
ICNSGFKRKAGTSSLTECVLNKATNVAHWTTPSLKCIRDPALVHQRPAPPSGGSGGGGS
GGGSGGGGSPLLCPREPFYFLRKITKMLENKNDGSLYTPDNLLVCPAETLRCFRLELSV
IGFEEGNKSSVGIVVFRLQRLLDALGSQLWVTDQGPCPPCEGHPQRPVPLFLAKLLELL
QGTCARDLPSA

#### pcDNA3.1-*Bos-IL-15Rα-Myc-His*

This expression vector encodes full-length bovine Myc-tagged IL-15Rα (named IL-15Rα).
Cloned from cDNA. The *IL-15Rα* part of the sequence is identical with GenBank accession XM_015465884.
*[for cloning details see (12)]*

AAGCTTACCATGTCCGGGCGGCTCCGGGGCCGCGGGGCCGGCGCCCTCCCCGCGCTGGG
GCTGCTGCTGCTACTACTGCTGCTCGGATCTTCGGCCACGCCGGGCATCACCTGCCCGA
CTCCCACATCCGTGGAGCATGCAGACATCCAGGTCAAGAGTTACAGCATCAACTCCAGG
GAGCGGTATGTTTGTAATTCTGGCTTCAAGCGTAAAGCTGGGACTTCCAGCTTGACCCA
GTGTGTGTTTAACGAGACCGCGAAAGTCGCCCACTGGACCACTCCCAACCTCAAGTGCA
TCAGAGACCCCTCCCTGAGTCACCAAAGGCCACCCTCCACAGCAGCGCCTACAGGGTTG
ACCCCAGAGCCAGAGAGCCCCACCCCCTCCGGAAAAGAGCCAGATCTTACTTCCAAGTC
AGACACCAAAGTGGCCACAAGGCCAGCTACTGGACCAGGCTCCAGGCTGCCATCCACAG
CTCCTCCTGTGGGAACCACAGGGGTAGTCAGTAAGGAGACCACCTACGTCCCAGCTCAG
ACAGCAGCCAAGGCTCCGGAACACACATACCCGGCCTTGCAGGACACGCCCGGTGCATA
TCAGAACAATCCCAGAGTTGTGACCGCCGTCTCAACTGTCACTGTGCTCTTTGTAGTAT
GCCTGGTGTTTCTTCTGGGACGTTGCCTGTGGTCAAGGCGAGCCCACCAGACACCCGGT
GTTGAGATGGAGAGCATGGAGAGTGTGCCAATGACCACGGGGGCCGATGCCAGAGGGGA
GGACACAGAAATCCACCCGCATGGCCTAGGAGGCTCCGGGGACGCTGAGGCCAGCAGCG
GCCGCAGTGAAGGCCCAGCTCTTCCCCAGTCAGAGAGGACCTCGAGTCTAGAGGGCCCG
CGGTTCGAACAAAAACTCATCTCAGAAGAGGATCTGAATATGCATACCGGTCATCATCA
CCATCACCATTGA

MSGRLRGRGAGALPALGLLLLLLLLGSSATPGITCPTPTSVEHADIQVKSYSINSRERY
VCNSGFKRKAGTSSLTQCVFNETAKVAHWTTPNLKCIRDPSLSHQRPPSTAAPTGLTPE
PESPTPSGKEPDLTSKSDTKVATRPATGPGSRLPSTAPPVGTTGVVSKETTYVPAQTAA
KAPEHTYPALQDTPGAYQNNPRVVTAVSTVTVLFVVCLVFLLGRCLWSRRAHQTPGVEM
ESMESVPMTTGADARGEDTEIHPHGLGGSGDAEASSGRSEGPALPQSERTSSLEGPRFE
QKLISEEDLNMHTGHHHHHH

#### pcDNA3.1-*Bos-solIL-15Rα-Myc-His*

This expression vector encodes bovine soluble Myc-tagged IL-15Rα (named soluble IL-15Rα or sIL-15Rα).
Cloned from cDNA. The *IL-15Rα* part of the sequence is identical with GenBank accession XM_015465884.
*[for cloning details see (12)]*

AAGCTTACCATGTCCGGGCGGCTCCGGGGCCGCGGGGCCGGCGCCCTCCCCGCGCTGGG
GCTGCTGCTGCTACTACTGCTGCTCGGATCTTCGGCCACGCCGGGCATCACCTGCCCGA
CTCCCACATCCGTGGAGCATGCAGACATCCAGGTCAAGAGTTACAGCATCAACTCCAGG
GAGCGGTATGTTTGTAATTCTGGCTTCAAGCGTAAAGCTGGGACTTCCAGCTTGACCCA
GTGTGTGTTTAACGAGACCGCGAAAGTCGCCCACTGGACCACTCCCAACCTCAAGTGCA
TCAGAGACCCCTCCCTGAGTCACCAAAGGCCACCCTCCACAGCAGCGCCTACAGGGTTG
ACCCCAGAGCCAGAGAGCCCCACCCCCTCCGGAAAAGAGCCAGATCTTACTTCCAAGTC
AGACACCAAAGTGGCCACAAGGCCAGCTACTGGACCAGGCTCCAGGCTGCCATCCACAG
CTCCTCCTGTGGGAACCACAGGGGTAGTCAGTAAGGAGACCACCTACGTCCCAGCTCAG
ACAGCAGCCAAGGCTCCGGAACACACATACCCGGCCTTGCAGGACACGCCCGGTGCATA
TCAGAACAATCCCAGCTCGAGTCTAGAGGGCCCGCGGTTCGAACAAAAACTCATCTCAG
AAGAGGATCTGAATATGCATACCGGTCATCATCACCATCACCATTGA

MSGRLRGRGAGALPALGLLLLLLLLGSSATPGITCPTPTSVEHADIQVKSYSINSRERY
VCNSGFKRKAGTSSLTQCVFNETAKVAHWTTPNLKCIRDPSLSHQRPPSTAAPTGLTPE
PESPTPSGKEPDLTSKSDTKVATRPATGPGSRLPSTAPPVGTTGVVSKETTYVPAQTAA
KAPEHTYPALQDTPGAYQNNPSSSLEGPRFEQKLISEEDLNMHTGHHHHHH

#### pcDNA3.1-*Bos-IL-2Rα-Myc-His*

This expression vector encodes bovine full-length Myc-tagged IL-2Rα (named IL-2Rα).
Cloned from cDNA. Except for a single nucleotide replacement (shaded yellow), the *IL-2Rα* part of the sequence is identical with GenBank accession BC133546.
*[for cloning details see (12); through primer choice a silent mutation was introduced at the single yellow-shaded C position in order to delete a gene-internal restriction site]*

AAGCTTACCATGGAGCCCAGCTTGCTGATGTGGAGGTTCTTCGTATTCATCGTGGTACC
TGGCTGCGTGACAGAGGCTTGTCATGATGACCCTCCGAGTCTCAGAAACGCCATGTTCA
AGGTCTTCAGGTACGAGGTGGGCACCATGATAAACTGCGACTGCAAGACAGGCTTCCGC
AGAGTGTCGGCCGTCATGCGCTGCGTGGGGGACTCCAGCCACTCTGCCTGGGAAAACAG
ATGCTTCTGCAACAGCACCTCCCCTGCTAAGAACCCAGTAAAACAAGTCACTCCTGCAC
CCGAAGAACAGAGGGAGAAAAAACCCACAGATGCGCAGAACCAAACGCAGCCTCCGGAG
GAAGCTGACCTTCCAGGTCACTGTGAGGAACCGCCACCATGGGAACACGAACGTGAACC
TTTAAAGAGAGTCTACCATTTCACGCTGGGGCAGACGGTTCATTACCAGTGCGCCCAGG
GATTCAGGGCCCTACAGACCAGTCCTGCTGAAAGCACCTGCATGATGATCAACGGGGAG
CTGAGGTGGACCAGGCCCAGGCTCAAGTGCATACGTGAAGGGGAGCACGGTCAGGCTTC
AGATGACGCAGAGCCTCAGGAGAGCACGGAAGCTCCCCCTGGGAGTGGAACTTTCTTAC
CAACCAGGATGGCAGGGACCACAGATTTCCAGAAGCCCACAGATGAGATTGCAACGCTG
GATACGTTCATATTTACCACTGAGTACCAGATTGCAGTGGCCGGCTGCACCCTCCTGCT
CGCCAGCATCCTCCTCCTGAGCTGCCTCACCTGGCAGCGGAAATGGAAGAAGAACAGAA
GGACAATCTCGAGTCTAGAGGGCCCGCGGTTCGAACAAAAACTCATCTCAGAAGAGGAT
CTGAATATGCATACCGGTCATCATCACCATCACCATTGA

MEPSLLMWRFFVFIVVPGCVTEACHDDPPSLRNAMFKVFRYEVGTMINCDCKTGFRRVS
AVMRCVGDSSHSAWENRCFCNSTSPAKNPVKQVTPAPEEQREKKPTDAQNQTQPPEEAD
LPGHCEEPPPWEHEREPLKRVYHFTLGQTVHYQCAQGFRALQTSPAESTCMMINGELRW
TRPRLKCIREGEHGQASDDAEPQESTEAPPGSGTFLPTRMAGTTDFQKPTDEIATLDTF
IFTTEYQIAVAGCTLLLASILLLSCLTWQRKWKKNRRTISSLEGPRFEQKLISEEDLNM
HTGHHHHHH

#### pcDNA3.1-*Bos-solIL-2Rα-Myc-His*

This expression vector encodes bovine soluble Myc-tagged IL-2Rα (named soluble IL-2Rα or sIL-2Rα).
Cloned from cDNA. Except for a single nucleotide replacement (shaded yellow), the *IL-2Rα* part of the sequence is identical with GenBank accession BC133546.
*[for cloning details see (12); through primer choice a silent mutation was introduced at the single yellow-shaded C position in order to delete a gene-internal restriction site]*

AAGCTTACCATGGAGCCCAGCTTGCTGATGTGGAGGTTCTTCGTATTCATCGTGGTACC
TGGCTGCGTGACAGAGGCTTGTCATGATGACCCTCCGAGTCTCAGAAACGCCATGTTCA
AGGTCTTCAGGTACGAGGTGGGCACCATGATAAACTGCGACTGCAAGACAGGCTTCCGC
AGAGTGTCGGCCGTCATGCGCTGCGTGGGGGACTCCAGCCACTCTGCCTGGGAAAACAG
ATGCTTCTGCAACAGCACCTCCCCTGCTAAGAACCCAGTAAAACAAGTCACTCCTGCAC
CCGAAGAACAGAGGGAGAAAAAACCCACAGATGCGCAGAACCAAACGCAGCCTCCGGAG
GAAGCTGACCTTCCAGGTCACTGTGAGGAACCGCCACCATGGGAACACGAACGTGAACC
TTTAAAGAGAGTCTACCATTTCACGCTGGGGCAGACGGTTCATTACCAGTGCGCCCAGG
GATTCAGGGCCCTACAGACCAGTCCTGCTGAAAGCACCTGCATGATGATCAACGGGGAG
CTGAGGTGGACCAGGCCCAGGCTCAAGTGCATACGTGAAGGGGAGCACGGTCAGGCTTC
AGATGACGCAGAGCCTCAGGAGAGCACGGAAGCTCCCCCTGGGAGTGGAACTTTCTTAC
CAACCAGGATGGCAGGGACCACAGATTTCCAGAAGCCCACAGATGACTCGAGTCTAGAG

GGCCCGCGGTTCGAACAAAAACTCATCTCAGAAGAGGATCTGAATATGCATACCGGTCA
TCATCACCATCACCATTGA

MEPSLLMWRFFVFIVVPGCVTEACHDDPPSLRNAMFKVFRYEVGTMINCDCKTGFRRVS
AVMRCVGDSSHSAWENRCFCNSTSPAKNPVKQVTPAPEEQREKKPTDAQNQTQPPEEAD
LPGHCEEPPPWEHEREPLKRVYHFTLGQTVHYQCAQGFRALQTSPAESTCMMINGELRW
TRPRLKCIREGEHGQASDDAEPQESTEAPPGSGTFLPTRMAGTTDFQKPTDEIATLDTF
IFTTEYQIAVAGCTLLLASILLLSCLTWQRKWKKNRRTISSLEGPRFEQKLISEEDLNM
HTGHHHHHH

**Figure S2B.**
**Expression vector for a genetic fusion of human IL-15Rα and IL-15.**

pcDNA3.1-*IL-2-Lead-RLI*

Except for the leader sequence which is of human IL-2, the encoded sequence is identical to the RLI molecule described by Mortier *et al.* 2006 (29) which is a genetic fusion subsequently encoding a FLAG-tag, a large part of the human IL-15Rα ectodomain, a glycine-serine linker (shaded blue), and human IL-15 (named RLI).
Amplified from commercial cDNA and using appropriate primers and overlap PCR.

GGTACCACCATGTACAGGATGCAACTCCTGTCTTGCATTGCACTAAGTCTTGCACTTGT
CACAAACAGTGACTACAAGGATGACGATGACAAGATAGAAGGTAGGATCACATGCCCTC
CCCCCATGTCCGTGGAACACGCAGACATCTGGGTCAAGAGCTACAGCTTGTACTCCAGG
GAGCGGTACATTTGTAACTCTGGTTTCAAGCGTAAAGCCGGCACGTCCAGCCTGACGGA
GTGCGTGTTGAACAAGGCCACGAATGTCGCCCACTGGACAACCCCCAGTCTCAAATGCA
TTAGAGACCCTGCCCTGGTTCACCAAAGGCCAGCGCCACCCTCTGGAGGCTCCGGTGGA
GGTGGGAGTGGCGGTGGATCCGGTGGCGGAGGCAGCCTGCAGAACTGGGTGAATGTAAT
AAGTGATTTGAAAAAAATTGAAGATCTTATTCAATCTATGCATATTGATGCTACTTTAT
ATACGGAAAGTGATGTTCACCCCAGTTGCAAAGTAACAGCAATGAAGTGCTTTCTCTTG
GAGTTACAAGTTATTTCACTTGAGTCCGGAGATGCAAGTATTCATGATACAGTAGAAAA
TCTGATCATCCTAGCAAACAACAGTTTGTCTTCTAATGGGAATGTAACAGAATCTGGAT
GCAAAGAATGTGAGGAACTGGAGGAAAAAAATATTAAAGAATTTTTGCAGAGTTTTGTA
CATATTGTCCAAATGTTCATCAACACTTCTTAATCTAGA

MYRMQLLSCIALSLALVTNSDYKDDDDKIEGRITCPPPMSVEHADIWVKSYSLYSRERY
ICNSGFKRKAGTSSLTECVLNKATNVAHWTTPSLKCIRDPALVHQRPAPPSGGSGGGGS
GGGSGGGGSLQNWVNVISDLKKIEDLIQSMHIDATLYTESDVHPSCKVTAMKCFLLELQ
VISLESGDASIHDTVENLIILANNSLSSNGNVTESGCKECEELEEKNIKEFLQSFVHIV
QMFINTS

**Figure S2C.**
**Expression vectors for (modified) trout cytokines and IL-15Rα.**

pcDNA3.1-*trout-IL-2-FLAG*

This expression vector encodes rainbow trout FLAG-tagged IL-2 (named IL-2).
Cloned from cDNA. The *IL-2* part of the sequence is identical with GenBank accession FJ571512.

AAGCTTACCATGGACCGTCGTTACAGGATTTCCTTTTTGACGCTTTTTCTCACCGGTTG
TCTACAAGGAAACCCAATTCCCAGACTCCTAGCTGGAATCGATTATCTAGAAGAAAATA
TTACTTGTCCAGATTCAGTCTTCTATACACCAACTGATGTAGAGGATAGTTGCATTGTT
GCAGCATTGGCCTGTTCCATTAAGGAACTGGACACTGTGAAAGTAGAATGCCTCGATAA
CGCGGTCCATCTGGAAAGTATGCAACACCACATCAGCATGACTGCCACGGACCTACAAA
AGACGATTGATAAGGAGAACAGCACAACGGACACTTCAGAATGCATCTGTGAAGACAAG
CGGTTGGAAAAGTCTTTCAAGGACTTCATTCAGAACATAAGACATTTAACTCAAGCTCA
TGCTGCAAAGCGTCTAAGTTCAGATTACAAGGATGACGACGATAAGTAACTCGAG

MDRRYRISFLTLFLTGCLQGNPIPRLLAGIDYLEENITCPDSVFYTPTDVEDSCIVAAL
ACSIKELDTVKVECLDNAVHLESMQHHISMTATDLQKTIDKENSTTDTSECICEDKRLE
KSFKDFIQNIRHLTQAHAAKRLSSDYKDDDDK

pcDNA3.1-*trout-IL-2-(non-tagged)*

This expression vector encodes rainbow trout non-tagged IL-2 [named IL-2(N)].
Cloned from cDNA. The *IL-2* part of the sequence is identical with GenBank accession FJ571512.

AAGCTTACCATGGACCGTCGTTACAGGATTTCCTTTTTGACGCTTTTTCTCACCGGTTG
TCTACAAGGAAACCCAATTCCCAGACTCCTAGCTGGAATCGATTATCTAGAAGAAAATA
TTACTTGTCCAGATTCAGTCTTCTATACACCAACTGATGTAGAGGATAGTTGCATTGTT
GCAGCATTGGCCTGTTCCATTAAGGAACTGGACACTGTGAAAGTAGAATGCCTCGATAA
CGCGGTCCATCTGGAAAGTATGCAACACCACATCAGCATGACTGCCACGGACCTACAAA
AGACGATTGATAAGGAGAACAGCACAACGGACACTTCAGAATGCATCTGTGAAGACAAG
CGGTTGGAAAAGTCTTTCAAGGACTTCATTCAGAACATAAGACATTTAACTCAAGCTCA
TGCTGCAAAGCGTCTAAGTTCATAACTCGAG

MDRRYRISFLTLFLTGCLQGNPIPRLLAGIDYLEENITCPDSVFYTPTDVEDSCIVAAL
ACSIKELDTVKVECLDNAVHLESMQHHISMTATDLQKTIDKENSTTDTSECICEDKRLE
KSFKDFIQNIRHLTQAHAAKRLSS

#### pcDNA3.1-*trout-IL-15-FLAG*

This expression vector encodes rainbow trout FLAG-tagged IL-15 (named IL-15). Cloned from cDNA. The *IL-15* part of the sequence is identical with GenBank accession NM_001124395.

```
AAGCTTACCATGACAGGTTTTTGACAGTGCTCCTTTTTTGCATTCGTTTATTGGAGCG
CAGGACAAAGAAAAGTGTGCGATGGATCTGTCTCTTCTGGGGTTTCCATTACTATCCAC
ACCAGCGTCTGAACATTGAGCTCTGGAATTGCTTCATAATATTGAGCTGCCTGAGTGCC
ACCGCACATTTGCCCATTGCTGGTGCTGCTGAAACACACGGGATGACAATAGATGACGT
TAAAGAGCTTCAGTCGGAGTTGAAAAACTTGAAAAGTACCATAGAAAAATCAGATGCCT
GTTTGTATGCTCCTACCAACGATGACATCTACAATGACCACTGCATCTTTAAGTTCATG
CACTGTTATTTATTGGAGTTGGAGGTTGTCCTGTTTGAGGACATGTCGGTCACAGATAA
CTACCATGACGAAATAAAAACATCCATCTACCATCGGAAAAAACATTTGGAAGAACATG
AACGCCAATATAACAGTTCTAGATGCTCACCGTGTGAGGCACAGAGAGTTGCCAACTCC
ACAATATTCCTCTACAACCTGGAGCGCCTTTTGGAGAGAATAGGGCAAACTGTCAGTGA
TTACAAGGATGACGACGATAAGTAACTCGAG
```

```
MTGFLTVLLFCIRLLERRTKKSVRWICLFWGFHYYPHQRLNIELWNCFIILSCLSATAH
LPIAGAAETHGMTIDDVKELQSELKNLKSTIEKSDACLYAPTNDDIYNDHCIFKFMHCY
LLELEVVLFEDMSVTDNYHDEIKTSIYHRKKHLEEHERQYNSSRCSPCEAQRVANSTIF
LYNLERLLERIGQTVSDYKDDDDK
```

#### pcDNA3.1-*trout-IL-15-(non-tagged)*

This expression vector encodes rainbow trout non-tagged IL-15 [named IL-15(N)]. Cloned from cDNA. The *IL-15* part of the sequence is identical with GenBank accession NM_001124395.

```
AAGCTTACCATGACAGGTTTTTGACAGTGCTCCTTTTTTGCATTCGTTTATTGGAGCG
CAGGACAAAGAAAAGTGTGCGATGGATCTGTCTCTTCTGGGGTTTCCATTACTATCCAC
ACCAGCGTCTGAACATTGAGCTCTGGAATTGCTTCATAATATTGAGCTGCCTGAGTGCC
ACCGCACATTTGCCCATTGCTGGTGCTGCTGAAACACACGGGATGACAATAGATGACGT
TAAAGAGCTTCAGTCGGAGTTGAAAAACTTGAAAAGTACCATAGAAAAATCAGATGCCT
GTTTGTATGCTCCTACCAACGATGACATCTACAATGACCACTGCATCTTTAAGTTCATG
CACTGTTATTTATTGGAGTTGGAGGTTGTCCTGTTTGAGGACATGTCGGTCACAGATAA
CTACCATGACGAAATAAAAACATCCATCTACCATCGGAAAAAACATTTGGAAGAACATG
AACGCCAATATAACAGTTCTAGATGCTCACCGTGTGAGGCACAGAGAGTTGCCAACTCC
ACAATATTCCTCTACAACCTGGAGCGCCTTTTGGAGAGAATAGGGCAAACTGTCAGTTG
ACTCGAG
```

```
MTGFLTVLLFCIRLLERRTKKSVRWICLFWGFHYYPHQRLNIELWNCFIILSCLSATAH
LPIAGAAETHGMTIDDVKELQSELKNLKSTIEKSDACLYAPTNDDIYNDHCIFKFMHCY
LLELEVVLFEDMSVTDNYHDEIKTSIYHRKKHLEEHERQYNSSRCSPCEAQRVANSTIF
LYNLERLLERIGQTVS
```

#### pcDNA3.1-*trout-IL-15La-FLAG*

This expression vector encodes rainbow trout FLAG-tagged IL-15La (named IL-15La). Cloned from cDNA. The *IL-15La* part of the sequence is identical with GenBank accession MK619679.

AAGCTTACCATGCTGAGGAGACAGAGGACTGACACTCTTCTAGCCCTTTTGCTGTGGTT
TCTCTTCTTCATCGCCATGACAATGAAACAGGCATATGGAAAATCCATGTGCAGTAAAG
AACTTCCCGGAATTGTGCGAAAATGCATTGAGGAAGTTCACAAGATGGAATCATTTGAT
TGCAGACTGTACACCCCAACTTTGGCTGATTATCAGAAGTGCCCCACGTCCACACTCAT
ATGCTTTGAAAAAGAAGTGAATGTCCTGGTGTTAGAATCTGGGAATAAGTCCTCACCCA
TATACAAGCCAAAACTATCCATCCGGCTGAAGTCCTTGATCAAACAGAAAGAAGGTGCC
AACTGTCCAGACTGTGAGGCCCACAGAGAAAGGGCAGCAAAGGATTTCTTAACAACATT
GCAAACAATTCTGGAGTGGATGAACGATCAGGGGTGTCGGAAGCCATCCAGCCATGATT
ACAAGGATGACGACGATAAGTGACTCGAG

MLRRQRTDTLLALLLWFLFFIAMTMKQAYGKSMCSKELPGIVRKCIEEVHKMESFDCRL
YTPTLADYQKCPTSTLICFEKEVNVLVLESGNKSSPIYKPKLSIRLKSLIKQKEGANCP
DCEAHRERAAKDFLTTLQTILEWMNDQGCRKPSSHDYKDDDDK

#### pcDNA3.1-*trout-IL-15La-(non-tagged)*

This expression vector encodes rainbow trout un-tagged IL-15La [named IL-15La(N)]. Cloned from cDNA. The *IL-15La* part of the sequence is identical with GenBank accession MK619679.

AAGCTTACCATGCTGAGGAGACAGAGGACTGACACTCTTCTAGCCCTTTTGCTGTGGTT
TCTCTTCTTCATCGCCATGACAATGAAACAGGCATATGGAAAATCCATGTGCAGTAAAG
AACTTCCCGGAATTGTGCGAAAATGCATTGAGGAAGTTCACAAGATGGAATCATTTGAT
TGCAGACTGTACACCCCAACTTTGGCTGATTATCAGAAGTGCCCCACGTCCACACTCAT
ATGCTTTGAAAAAGAAGTGAATGTCCTGGTGTTAGAATCTGGGAATAAGTCCTCACCCA
TATACAAGCCAAAACTATCCATCCGGCTGAAGTCCTTGATCAAACAGAAAGAAGGTGCC
AACTGTCCAGACTGTGAGGCCCACAGAGAAAGGGCAGCAAAGGATTTCTTAACAACATT
GCAAACAATTCTGGAGTGGATGAACGATCAGGGGTGTCGGAAGCCATCCAGCCATTGAC
TCGAG

MLRRQRTDTLLALLLWFLFFIAMTMKQAYGKSMCSKELPGIVRKCIEEVHKMESFDCRL
YTPTLADYQKCPTSTLICFEKEVNVLVLESGNKSSPIYKPKLSIRLKSLIKQKEGANCP
DCEAHRERAAKDFLTTLQTILEWMNDQGCRKPSSH

#### pcDNA3.1-*trout-IL-15Lb-FLAG*

This expression vector encodes rainbow trout FLAG-tagged IL-15Lb (named IL-15Lb). Cloned from cDNA. The *IL-15Lb* part of the sequence is identical with GenBank accession MK619680.

AAGCTTACCATGTTGAGGAGACAGAGAACTGGCTCTCTTCTGAACGCTTTGCTGTGGTT
TCTCTTCTTCATTGCCATGACAATGAAACAGGCTTATGGACAATCCATTAGCAGTTCAG
AAATTCACCAAATTGTGAAAACATTTATTAAGGAGGAAGTTCACAGGATGGAATCATTT
GATTGCAGACTGTACACCCCAACTTTAGCTGATTATAAGAAATGTCCCAGGTCCACACT
CACATGCTTTACGACAGAAGTAAAAGTCCTGATGTTAGAATATGGGAAACGTTCTTCAT
CCTTACACCAGAAAAGACTCACCAAACGACTGACTAAATTGATGTCCTTGATAAAACAG
AAGGATGGTGCCAACTGTCCACACTGTGAGGTCCACAGAGAACAGGCAGCAAATGATTT
CTTAACAACATTACTAGGGATTCTGGAGTGGATGAACAATCAGGGGTCTCAGTTGCCAG
ACAGCCACGATTACAAGGATGACGACGATAAGTGACTCGAG

MLRRQRTGSLLNALLWFLFFIAMTMKQAYGQSISSSEIHQIVKTFIKEEVHRMESFDCR
LYTPTLADYKKCPRSTLTCFTTEVKVLMLEYGKRSSSLHQKRLTKRLTKLMSLIKQKDG
ANCPHCEVHREQAANDFLTTLLGILEWMNNQGSQLPDSHDYKDDDDK

#### pcDNA3.1-*trout-IL-15Lb-(non-tagged)*

This expression vector encodes rainbow trout un-tagged IL-15Lb [named IL-15Lb(N)]. Cloned from cDNA. The *IL-15Lb* part of the sequence is identical with GenBank accession MK619680.

AAGCTTACCATGTTGAGGAGACAGAGAACTGGCTCTCTTCTGAACGCTTTGCTGTGGTT
TCTCTTCTTCATTGCCATGACAATGAAACAGGCTTATGGACAATCCATTAGCAGTTCAG
AAATTCACCAAATTGTGAAAACATTTATTAAGGAGGAAGTTCACAGGATGGAATCATTT
GATTGCAGACTGTACACCCCAACTTTAGCTGATTATAAGAAATGtCCCAGGTCCACACT
CACATGCTTTACGACAGAAGTAAAAGTCCTGATGTTAGAATATGGGAAACGTTCTTCAT
CCTTACACCAGAAAAGACTCACCAAACGACTGACTAAATTGATGTCCTTGATAAAACAG
AAGGATGGTGCCAACTGTCCACACTGTGAGGTCCACAGAGAACAGGCAGCAAATGATTT
CTTAACAACATTACTAGGGATTCTGGAGTGGATGAACAATCAGGGGTCTCAGTTGCCAG
ACAGCCACTGACTCGAG

MLRRQRTGSLLNALLWFLFFIAMTMKQAYGQSISSSEIHQIVKTFIKEEVHRMESFDCR
LYTPTLADYKKCPRSTLTCFTTEVKVLMLEYGKRSSSLHQKRLTKRLTKLMSLIKQKDG
ANCPHCEVHREQAANDFLTTLLGILEWMNNQGSQLPDSH

#### pcDNA3.1-*IL-2-Lead-RLI-trout-IL-15La*

This expression vector is based on the RLI construct described by Mortier *et al.* 2006 (29), and encodes a genetic fusion of a human IL-2 leader, a FLAG-tag, a large part of the human IL-15Rα ectodomain, a glycine-serine linker (shaded blue), and the rainbow trout IL-15La sequence (named IL-15La-h-RLI; indicated as "RLI" in main text Fig. 8).
The linker plus *IL-15La* part were commercially ordered (Invitrogen) with the codons optimized for expression in insect cells, and fused to the N-terminal coding part by using the above described plasmid pcDNA3.1-*IL-2-Lead-RLI*, appropriate primers and overlap PCR.

GGTACCACCATGTACAGGATGCAACTCCTGTCTTGCATTGCACTAAGTCTTGCACTTGT
CACAAACAGTGACTACAAGGATGACGATGACAAGATAGAAGGTAGGATCACATGCCCTC
CCCCCATGTCCGTGGAACACGCAGACATCTGGGTCAAGAGCTACAGCTTGTACTCCAGG
GAGCGGTACATTTGTAACTCTGGTTTCAAGCGTAAAGCCGGCACGTCCAGCCTGACGGA
GTGCGTGTTGAACAAGGCCACGAATGTCGCCCACTGGACAACCCCCAGTCTCAAATGCA
TTAGAGACCCTGCCCTGGTTCACCAAAGGCCAGCGCCACCCTCTGGAGGCTCCGGTGGA
GGTGGGAGTGGCGGTGGATCCGGTGGCGGAGGCAGCAAGCAAGCTTACGGCAAGTCCAT
GTGCTCCAAGGAACTGCCCGGTATCGTGCGCAAGTGCATCGAAGAGGTGCACAAGATGG
AATCCTTCGACTGCCGTCTGTACACCCCCACCCTGGCTGACTACCAGAAGTGCCCTACC
TCCACCCTGATCTGCTTCGAGAAGGAAGTGAACGTCCTGGTCCTCGAGTCCGGCAACAA
GTCCTCCCCAATCTACAAGCCCAAGCTGTCCATCCGTCTGAAGTCCCTGATCAAGCAGA
AGGAAGGCGCTAACTGCCCCGACTGCGAGGCTCACAGAGAGCGTGCTGCTAAGGACTTC
CTGACCACCCTGCAGACCATCCTCGAGTGGATGAACGACCAGGGTTGCCGCAAGCCCTC
CTCCCACTAATCTAGA

MYRMQLLSCIALSLALVTNSDYKDDDDKIEGRITCPPPMSVEHADIWVKSYSLYSRERY
ICNSGFKRKAGTSSLTECVLNKATNVAHWTTPSLKCIRDPALVHQRPAPPSGGSGGGGS
GGGSGGGGSKQAYGKSMCSKELPGIVRKCIEEVHKMESFDCRLYTPTLADYQKCPTSTL
ICFEKEVNVLVLESGNKSSPIYKPKLSIRLKSLIKQKEGANCPDCEAHRERAAKDFLTT
LQTILEWMNDQGCRKPSSH

#### pcDNA3.1-*trout-IL-15Rα-Myc*

This expression vector encodes rainbow trout full-length Myc-tagged IL-15Rα (named IL-15Rα).
Cloned and polished from cDNA. The *IL-15Rα* part of the sequence is, except for 1 silent nucleotide exchange, identical with GenBank accession GFIN01030410.

AAGCTTACCATGCGGCTGCGTCTCCTTTCCCTCCTTCTTATCGTCAAAGTGTGTCGACT
CAATACTGTCAGTCCAGACGTGTGCCCACCCCTGTCTCATTGGAAGCTCACCAAGCCTT
TGAACACGAGGGAAATTTATCAAGAAAACGAATCCCTGCGCTACCAGTGTGTGGATGGC
TATGTGAGGAAGGCGGGAACCTCCAATCTCATCAAGTGCAAACGGATTGGTCAGGTTCT
TCAGTGGACACTCCCAACACTGATATGCATACCTGACCCATCCATTCCAACTACGATAG
AACCTTCAACAACCCACAAGACAACTCAATTTCCTACAACGCACCTCTCAGTGCCCACA

GAGAGAGTTAGCCCAAGACCAACCACTCTCAGTTCACTTTTATCTCAAACACTGCATTC
TGTCACAGAGACCCAAGAGGTCATTACTACGGTTACAGTGACAACCACCATCATCAGTG
ACATCACATCAGTGCCAGTGAATAAGTCCAGTGCGGTGACCTCTGTCTCCACGAAGACT
GTTTCTCCATCAGAGAGTTACTGTGTCCCCAACAGAGCCCCAACCTCCAGTGACTATCC
AGTTGGCGATGGAAGCACTGGGAAGAGTTCTGCTACAGCCATATATGCAGGTGTAGGTG
GGGTGGTAGTAACCATCTTGCTGGTTGCAGCGTTTGGAGTACTAATGTTTTGGAGAAGG
AAACGGATGCAGCGGCTTGATCTGTCGCCAACACCAGACGAAATTATGCAAATGAATGT
TGTGACAGAACAAAAACTCATCTCAGAAGAGGATCTGTGACTCGAG

MRLRLLSLLLIVKVCRLNTVSPDVCPPLSHWKLTKPLNTREIYQENESLRYQCVDGYVR
KAGTSNLIKCKRIGQVLQWTLPTLICIPDPSIPTTIEPSTTHKTTQFPTTHLSVPTERV
SPRPTTLSSLLSQTLHSVTETQEVITTVTVTTTIISDITSVPVNKSSAVTSVSTKTVSP
SESYCVPNRAPTSSDYPVGDGSTGKSSATAIYAGVGGVVVTILLVAAFGVLMFWRRKRM
QRLDLSPTPDEIMQMNVVTEQKLISEEDL

#### pcDNA3.1-*trout-solIL-15Rα-Myc*

This expression vector encodes rainbow trout soluble Myc-tagged IL-15Rα (named soluble IL-15Rα or sIL-15Rα).
Cloned and polished from cDNA. The *IL-15Rα* part of the sequence is, except for 1 silent nucleotide exchange, identical with GenBank accession GFIN01030410.

AAGCTTACCATGCGGCTGCGTCTCCTTTCCCTCCTTCTTATCGTCAAAGTGTGTCGACT
CAATACTGTCAGTCCAGACGTGTGCCCACCCCTGTCTCATTGGAAGCTCACCAAGCCTT
TGAACACGAGGGAAATTTATCAAGAAAACGAATCCCTGCGCTACCAGTGTGTGGATGGC
TATGTGAGGAAGGCGGGAACCTCCAATCTCATCAAGTGCAAACGGATTGGTCAGGTTCT
TCAGTGGACACTCCCAACACTGATATGCATACCTGACCCATCCATTCCAACTACGATAG
AACCTTCAACAACCCACAAGACAACTCAAGAACAAAAACTCATCTCAGAAGAGGATCTG
TGACTCGAG

MRLRLLSLLLIVKVCRLNTVSPDVCPPLSHWKLTKPLNTREIYQENESLRYQCVDGYVR
KAGTSNLIKCKRIGQVLQWTLPTLICIPDPSIPTTIEPSTTHKTTQEQKLISEEDL

**Figure S2D.**

**Vectors for expressing (modified) trout cytokines and trout IL-15Rα in insect cells using a baculovirus system.**

#### pFBD-P10Uhis-ieGFP-*trout-IL-2-FLAG*

This expression vector encodes rainbow trout FLAG-tagged IL-2 (named IL-2).
Cloned from cDNA. The *IL-2* part of the sequence is identical with GenBank accession FJ571512.

CCCGGGACCATGGACCGTCGTTACAGGATTTCCTTTTTGACGCTTTTTCTCACCGGTTG
TCTACAAGGAAACCCAATTCCCAGACTCCTAGCTGGAATCGATTATCTAGAAGAAAATA
TTACTTGTCCAGATTCAGTCTTCTATACACCAACTGATGTAGAGGATAGTTGCATTGTT
GCAGCATTGGCCTGTTCCATTAAGGAACTGGACACTGTGAAAGTAGAATGCCTCGATAA
CGCGGTCCATCTGGAAAGTATGCAACACCACATCAGCATGACTGCCACGGACCTACAAA
AGACGATTGATAAGGAGAACAGCACAACGGACACTTCAGAATGCATCTGTGAAGACAAG
CGGTTGGAAAAGTCTTTCAAGGACTTCATTCAGAACATAAGACATTTAACTCAAGCTCA
TGCTGCAAAGCGTCTAAGTTCAGATTACAAGGATGACGACGATAAGTAACCCGGG

MDRRYRISFLTLFLTGCLQGNPIPRLLAGIDYLEENITCPDSVFYTPTDVEDSCIVAAL
ACSIKELDTVKVECLDNAVHLESMQHHISMTATDLQKTIDKENSTTDTSECICEDKRLE
KSFKDFIQNIRHLTQAHAAKRLSSDYKDDDDK

#### pFBD-P10Uhis-ieGFP-*trout-IL-15-FLAG*

This expression vector encodes rainbow trout FLAG-tagged IL-15 (named IL-15).
The sequence was commercially ordered (Invitrogen). The signal peptide is replaced with that of rainbow trout IL-2. The *IL-2* part of the sequence is identical with GenBank accession FJ571512. The *IL-15* part of the sequence is identical with GenBank accession NM_001124395. The encoded IL-2 leader sequence is shaded blue.

CCCGGGACCATGGACCGTCGTTACAGGATTTCCTTTTTGACGCTTTTTCTCACCGGTTG
TCTACAAGGAGCTGAAACACACGGGATGACAATAGATGACGTTAAAGAGCTTCAGTCGG
AGTTGAAAAACTTGAAAAGTACCATAGAAAAATCAGATGCCTGTTTGTATGCTCCTACC
AACGATGACATCTACAATGACCACTGCATCTTTAAGTTCATGCACTGTTATTTATTGGA
GTTGGAGGTTGTCCTGTTTGAGGACATGTCGGTCACAGATAACTACCATGACGAAATAA
AAACATCCATCTACCATCGGAAAAAACATTTGGAAGAACATGAACGCCAATATAACAGT
TCTAGATGCTCACCGTGTGAGGCACAGAGAGTTGCCAACTCCACAATATTCCTCTACAA
CCTGGAGCGCCTTTTGGAGAGAATAGGGCAAACTGTCAGTGATTACAAGGATGACGACG
ATAAGTGACCCGGG

MDRRYRISFLTLFLTGCLQGAETHGMTIDDVKELQSELKNLKSTIEKSDACLYAPTNDD
IYNDHCIFKFMHCYLLELEVVLFEDMSVTDNYHDEIKTSIYHRKKHLEEHERQYNSSRC
SPCEAQRVANSTIFLYNLERLLERIGQTVSDYKDDDDK

#### pFBD-P10Uhis-ieGFP-*trout-IL-15La-FLAG*

This expression vector encodes rainbow trout FLAG-tagged IL-15La (named IL-15La). Cloned from cDNA. The *IL-15La* part of the sequence is identical with GenBank accession MK619679.

CCCGGGACCATGCTGAGGAGACAGAGGACTGACACTCTTCTAGCCCTTTTGCTGTGGTT
TCTCTTCTTCATCGCCATGACAATGAAACAGGCATATGGAAAATCCATGTGCAGTAAAG
AACTTCCCGGAATTGTGCGAAAATGCATTGAGGAAGTTCACAAGATGGAATCATTTGAT
TGCAGACTGTACACCCCAACTTTGGCTGATTATCAGAAGTGCCCCACGTCCACACTCAT
ATGCTTTGAAAAAGAAGTGAATGTCCTGGTGTTAGAATCTGGGAATAAGTCCTCACCCA
TATACAAGCCAAAACTATCCATCCGGCTGAAGTCCTTGATCAAACAGAAAGAAGGTGCC
AACTGTCCAGACTGTGAGGCCCACAGAGAAAGGGCAGCAAAGGATTTCTTAACAACATT
GCAAACAATTCTGGAGTGGATGAACGATCAGGGGTGTCGGAAGCCATCCAGCCATGATT
ACAAGGATGACGACGATAAGTAACCCGGG

MLRRQRTDTLLALLLWFLFFIAMTMKQAYGKSMCSKELPGIVRKCIEEVHKMESFDCRL
YTPTLADYQKCPTSTLICFEKEVNVLVLESGNKSSPIYKPKLSIRLKSLIKQKEGANCP
DCEAHRERAAKDFLTTLQTILEWMNDQGCRKPSSHDYKDDDDK

#### pFBD-P10Uhis-ieGFP-*trout-solIL-15Rα-Myc*

This expression vector encodes rainbow trout soluble Myc-tagged IL-15Rα (named soluble IL-15Rα or sIL-15Rα).
Cloned from cDNA. The *IL-15Rα* part of the sequence is, except for 1 silent nucleotide exchange, identical with GenBank accession GFIN01030410.

CCCGGGACCATGCGGCTGCGTCTCCTTTCCCTCCTTCTTATCGTCAAAGTGTGTCGACT
CAATACTGTCAGTCCAGACGTGTGCCCACCCCTGTCTCATTGGAAGCTCACCAAGCCTT
TGAACACGAGGGAAATTTATCAAGAAAACGAATCCCTGCGCTACCAGTGTGTGGATGGC
TATGTGAGGAAGGCGGGAACCTCCAATCTCATCAAGTGCAAACGGATTGGTCAGGTTCT
TCAGTGGACACTCCCAACACTGATATGCATACCTGACCCATCCATTCCAACTACGATAG
AACCTTCAACAACCCACAAGACAACTCAAGAACAAAAACTCATCTCAGAAGAGGATCTG
TGACCCGGG

MRLRLLSLLLIVKVCRLNTVSPDVCPPLSHWKLTKPLNTREIYQENESLRYQCVDGYVR
KAGTSNLIKCKRIGQVLQWTLPTLICIPDPSIPTTIEPSTTHKTTQEQKLISEEDL

#### pFBD-P10Uhis-ieGFP-*trout-IL-15-RLI*

This expression vector is based on the RLI construct described by Mortier *et al.* 2006 (29), and encodes a genetic fusion of a human IL-2 leader, a FLAG-tag, a large part of the trout IL-15Rα ectodomain, a glycine-serine linker (shaded blue), and the rainbow trout IL-15 sequence (named IL-15-RLI).
The sequence was commercially ordered (Invitrogen). The *IL-15Rα* part of the sequence is, except for 1 silent nucleotide exchange, identical with GenBank accession GFIN01030410. The *IL-15* part of the sequence is identical with GenBank accession NM_001124395.

CCCGGGACCATGTACAGGATGCAACTCCTGTCTTGCATTGCACTAAGTCTTGCACTTGT
CACAAACAGTGACTACAAGGATGACGATGACAAGAATACTGTCAGTCCAGACGTGTGCC
CACCCCTGTCTCATTGGAAGCTCACCAAGCCTTTGAACACGAGGGAAATTTATCAAGAA
AACGAATCCCTGCGCTACCAGTGTGTGGATGGCTATGTGAGGAAGGCGGGAACCTCCAA
TCTCATCAAGTGCAAACGGATTGGTCAGGTTCTTCAGTGGACACTCCCAACACTGATAT
GCATACCTGACCCATCCATTCCAACTACGATAGAACCTTCAACATCTGGAGGCTCCGGT
GGAGGTGGGAGTGGCGGTGGATCCGGTGGCGGAGGCAGCGAAACACACGGGATGACAAT
AGATGACGTTAAAGAGCTTCAGTCGGAGTTGAAAAACTTGAAAAGTACCATAGAAAAAT
CAGATGCCTGTTTGTATGCTCCTACCAACGATGACATCTACAATGACCACTGCATCTTT
AAGTTCATGCACTGTTATTTATTGGAGTTGGAGGTTGTCCTGTTTGAGGACATGTCGGT
CACAGATAACTACCATGACGAAATAAAAACATCCATCTACCATCGGAAAAAACATTTGG
AAGAACATGAACGCCAATATAACAGTTCTAGATGCTCACCGTGTGAGGCACAGAGAGTT
GCCAACTCCACAATATTCCTCTACAACCTGGAGCGCCTTTTGGAGAGAATAGGGCAAAC
TGTCAGTTGACCCGGG

MYRMQLLSCIALSLALVTNSDYKDDDDKNTVSPDVCPPLSHWKLTKPLNTREIYQENES
LRYQCVDGYVRKAGTSNLIKCKRIGQVLQWTLPTLICIPDPSIPTTIEPSTSGGSGGGG
SGGGSGGGGSETHGMTIDDVKELQSELKNLKSTIEKSDACLYAPTNDDIYNDHCIFKFM
HCYLLELEVVLFEDMSVTDNYHDEIKTSIYHRKKHLEEHERQYNSSRCSPCEAQRVANS
TIFLYNLERLLERIGQTVS

#### pFBD-P10Uhis-ieGFP-*trout-IL-15La-RLI*

This expression vector is based on the RLI construct described by Mortier *et al.* 2006 (29), and encodes a genetic fusion of a human IL-2 leader, a FLAG-tag, a large part of the rainbow trout IL-15Rα ectodomain, a glycine-serine linker (shaded blue), and the rainbow trout IL-15La sequence (named IL-15La-RLI).
The sequence was commercially ordered (Invitrogen). The *IL-15Rα* part of the sequence is, except for 1 silent nucleotide exchange, identical with GenBank accession GFIN01030410. The *IL-15La* codons were optimized for expression in insect cells.

CCCGGGACCATGTACAGGATGCAACTCCTGTCTTGCATTGCACTAAGTCTTGCACTTGT
CACAAACAGTGACTACAAGGATGACGATGACAAGAATACTGTCAGTCCAGACGTGTGCC
CACCCCTGTCTCATTGGAAGCTCACCAAGCCTTTGAACACGAGGGAAATTTATCAAGAA
AACGAATCCCTGCGCTACCAGTGTGTGGATGGCTATGTGAGGAAGGCGGGAACCTCCAA

TCTCATCAAGTGCAAACGGATTGGTCAGGTTCTTCAGTGGACACTCCCAACACTGATAT
GCATACCTGACCCATCCATTCCAACTACGATAGAACCTTCAACATCTGGAGGCTCCGGT
GGAGGTGGGAGTGGCGGTGGATCCGGTGGCGGAGGCAGCAAGCAAGCTTACGGCAAGTC
CATGTGCTCCAAGGAACTGCCCGGTATCGTGCGCAAGTGCATCGAAGAGGTGCACAAGA
TGGAATCCTTCGACTGCCGTCTGTACACCCCCACCCTGGCTGACTACCAGAAGTGCCCT
ACCTCCACCCTGATCTGCTTCGAGAAGGAAGTGAACGTCCTGGTCCTCGAGTCCGGCAA
CAAGTCCTCCCCAATCTACAAGCCCAAGCTGTCCATCCGTCTGAAGTCCCTGATCAAGC
AGAAGGAAGGCGCTAACTGCCCCGACTGCGAGGCTCACAGAGAGCGTGCTGCTAAGGAC
TTCCTGACCACCCTGCAGACCATCCTCGAGTGGATGAACGACCAGGGTTGCCGCAAGCC
CTCCTCCCACTAACCCGGG

MYRMQLLSCIALSLALVTNSDYKDDDDKNTVSPDVCPPLSHWKLTKPLNTREIYQENES
LRYQCVDGYVRKAGTSNLIKCKRIGQVLQWTLPTLICIPDPSIPTTIEPSTSGGSGGGG
SGGGSGGGGSKQAYGKSMCSKELPGIVRKCIEEVHKMESFDCRLYTPTLADYQKCPTST
LICFEKEVNVLVLESGNKSSPIYKPKLSIRLKSLIKQKEGANCPDCEAHRERAAKDFLT
TLQTILEWMNDQGCRKPSSH

### Supplementary figure 3 (Fig. S3)

Isolation of rainbow trout lymphocyte subpopulations by flow cytometry

**Table of Contents** **Page**

Introduction and general explanation of the figure 49

Fig. S3A: Establishment of a new monoclonal antibody against rainbow trout CD8α 51

Fig. S3B: Representative sortings of CD8α$^+$ versus CD8α$^-$ rainbow trout lymphocytes 53

Fig. S3C: Representative sorting of CD4$^-$CD8$^-$ (double negative, DN), CD4$^+$CD8$^+$ (double positive, DP), CD4 single positive (SP) and CD8SP rainbow trout lymphocytes from thymus 55

Fig. S3D: Representative sorting of CD4SP, CD8SP, IgMSP and CD4$^-$CD8$^-$IgM$^-$ triple negative (TN) rainbow trout splenocytes 57

Fig. S3E: Percentages of CD4SP, CD8SP, IgMSP and TN cells among rainbow trout spleen lymphocytes 58

#### Introduction and general explanation of the figure.

Teleost fish lack lymph nodes [e.g. (33)], but they possess a bona fide spleen and thymus [e.g. (33-38)], while the head kidney (HK; pronephros) simultaneously is a hematopoietic organ and a secondary lymphoid organ [e.g. (36, 39, 40)]. It has also been shown that gills as the respiratory organ of teleosts [e.g. (36, 38)] and the intestine [e.g. (36, 41)] are mucosal organs that contain numerous lymphocytes. Therefore, in this study, rainbow trout leukocytes were isolated from the lymphoid tissues spleen, thymus and HK, and from the mucosal tissues gill and intestine.

There are several lines of evidence that indicate that teleost fish cytotoxic TCRαβ T cells express CD8α as known in mammals [e.g. (20, 42-44)]. Whereas mammals have only one CD4 molecule, teleost fish have two quite diverged CD4-1 and CD4-2 molecules which in lymphocytes are mostly co-expressed [e.g. (22, 45-47)]; both CD4-1 and CD4-2 show expression patterns and molecular features consistent with a CD4 marker function of helper and regulatory T cells as known in mammals [e.g. (22, 45-51)]. Fish B cells do not undergo an immunoglobulin class switch thus lacking IgG, IgA and IgE, and the main B cell population expresses IgM (e.g. [52]).

In the present study, monoclonal antibodies (mAbs) against trout CD8α, CD4-1, CD4-2 and IgM were used to label leukocytes that had been isolated by Percoll density gradient centrifugation, after which cells with lymphocyte features ($FSC^{low}$ and $SSC^{low}$ gated cells; described below as "lymphocytes") were flow sorted into subpopulations depending on their labeling. To allow distinguishing usage of anti-CD8α together with other mAbs, a new mAb was established against trout CD8α possessing a different IgG isotype than the already available mAb 13.2D (20); this is explained in Fig. S3A. For CD4 labeling, a mixture containing both mAb 4.1.2 which recognizes trout CD4-1 and mAb 4.2.12 which recognizes trout CD4-2 (22) was used. For IgM labeling the mAb 1.14 was used (23). Doublets and dead cells were excluded by FSC-A/FSC-H gating and by propidium iodide (PI) or 4', 6-diamidino-2-phenylin-dole (DAPI) staining, respectively.

Details on the antibodies used for flow cytometry are:

*Primary antibodies:*

mAb 13.2D: anti-trout CD8α, rat IgG2a isotype (20)
mAb 7α8c: anti-trout CD8α, rat IgG1 isotype, established in this study (Fig. S3A)
mAb 4.1.2: anti-trout CD4-1, rat IgG2a isotype (22)
mAb 4.2.12: anti-trout CD4-2, rat IgG2b isotype (22)
mAb 1.14: anti-trout IgM, mouse IgG1 isotype (23)

*Fluorochrome-conjugated secondary antibodies:*

anti-rat IgG Alexa Fluor 488 (Thermo Fischer Scientific)
anti-rat IgG1-FITC (BD Bioscience)
anti-rat IgG2a-PE (eBioscience)
anti-rat IgG2a-eFluor 660 (eBioscience)
anti-rat IgG2b-PE (eBioscience)
anti-rat IgG2b-eFluor 660 (eBioscience)
anti-mouse IgG1-Brilliant Violet 421™ (Biolegend)

In the various experiments, different sets of the above listed antibodies were used for the flow sorting of lymphocyte subpopulations. The figures S3B-to-S3E are representative results for the flow sorting used in the present study, concerning lymphocytes from different tissues and different antibody combinations.

Fig. S3B shows representative sorting results for lymphocytes of various tissues when using primary antibody anti-trout CD8α (mAb 13.2D) plus secondary antibody anti-rat IgG Alexa Fluor 488. Two subpopulations were obtained, being $CD8^+$ and $CD8^-$ lymphocytes. This type of sorting was used for experiments of which results are shown in Fig. S5C.

Fig. S3C shows representative sorting results for lymphocytes of various tissues when using primary antibodies anti-trout CD8α (mAb 7α8c), anti-trout CD4-1 (mAb 4.1.2), and anti-trout CD4-2 (mAb 4.2.12), plus secondary antibodies anti-rat IgG1-FITC, anti-rat IgG2a-PE and anti-rat IgG2b-PE. Four subpopulations were obtained from thymus, being CD4SP, CD8SP, $CD4^-CD8^-$ (DN) and $CD4^+CD8^+$ (DP) lymphocytes while three subpopulations were obtained from spleen and intestine, being CD4SP, CD8SP and DN. This type of sorting was used for experiments of which results are shown in main text Fig. 9, and in Figs. S5C-6, S5C-11, S5D and S5E.

Fig. S3D shows representative sorting results for lymphocytes of various tissues when using primary antibodies anti-trout CD8α (mAb 7α8c), anti-trout CD4-1 (mAb 4.1.2), anti-trout CD4-2 (mAb 4.2.12), and anti-trout IgM (mAb 1.14), plus secondary antibodies anti-rat IgG1-FITC, anti-rat IgG2a-eFluor 660, anti-rat IgG2b-eFluor 660 and anti-mouse IgG1-Brilliant Violet 421™. Four subpopulations were obtained, being CD4SP (aka $CD4^+$), CD8SP (aka $CD8^+$), IgMSP (aka $IgM^+$), and $CD4^-CD8^-IgM^-$ (TN) lymphocytes. This type of sorting was used for experiments of which results are shown in main text Fig. 11 and in Figs. S6A and S6B.

Fig. S3E summarizes the relative numbers of the CD8SP (aka $CD8^+$), CD4SP (aka $CD4^+$) IgMSP (aka $IgM^+$), and $CD4^-CD8^-IgM^-$ (TN) cells among the spleen lymphocytes used for the experiments resulting in main text Fig. 11 (and Fig. S6B). This summary intends to help with the interpretation at the whole tissue level of the gene expression patterns observed within cell subpopulations.

*At a note:* Like in mammals, teleost lymphocytes can be distinguished from monocytes/macrophages and granulocytes by flow cytometry according to their scatter characteristics ($FSC^{low}$ and $SSC^{low}$) as small non-granulated cells [e.g. (20, 22, 40)]. Cells within this population are referred to as “lymphocyte gate cells” or “morphological lymphocytes”, and often are simply called “lymphocytes” in the present paper. However, it should be mentioned that teleost thrombocytes are nucleated cells with similar scatter characteristics as lymphocytes [e.g. (53)] and thus be contained in the corresponding negative populations.

#### **Fig. S3A** Establishment of a new mAb against rainbow trout CD8α.

A new anti-trout CD8α mAb 7α8c with another Ig isotype than the previously published anti-trout CD8α mAb 13.2D (20) was established because its Ig isotype was more suitable for multicolor staining in combination with mAbs against other molecules.

(a) Flow cytometry analysis of HEK293T cells transfected for expression of HA-tagged trout CD8α (HEK293T-CD8α-HA) and parental HEK293T cells stained with anti-HA (upper plots) and anti-trout CD8α 7α8c mAb (lower plots). (b) Rainbow trout leukocytes from intestine were stained with anti-CD8α 7α8c mAb and FITC-conjugated goat anti-rat IgG, and sorted into 7α8c-negative (-) and 7α8c-positive (+) lymphocytes. RNA was extracted from both populations and analyzed by RT-PCR using specific primers for *CD8α* (upper panel) and *β-actin* (lower panel). Distilled water served as a negative control. Numbers of cycles are provided on the right side of the figure.

(a) *Confirmation of 7α8c antibody specificity by flow cytometry analysis of HEK293T cells transfected for rainbow trout CD8α*

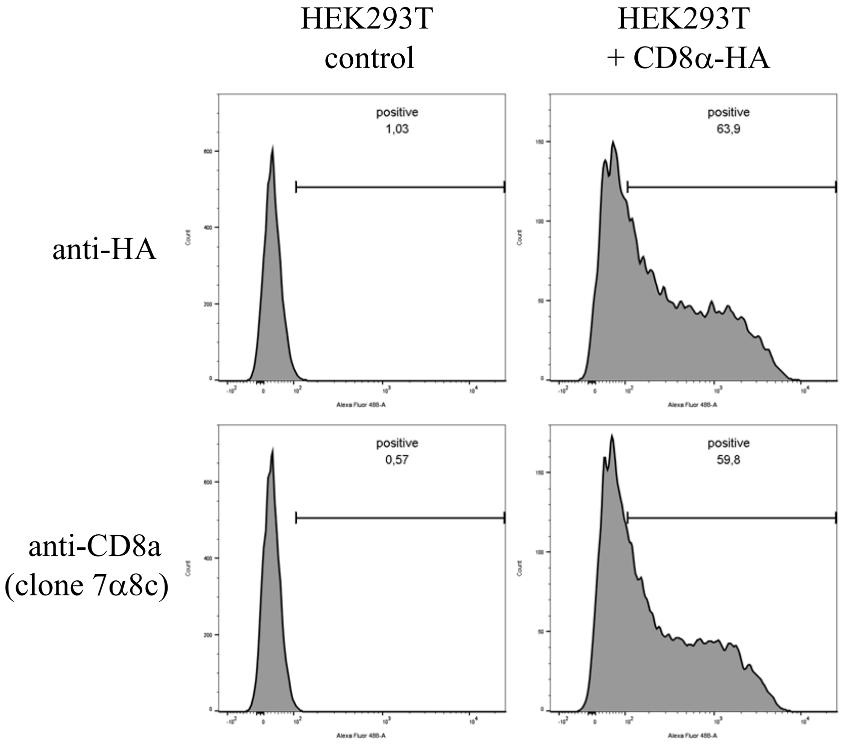

*(Fig. S3A continued)*

(b) *Confirmation of 7α8c antibody specificity by semi-quantitative RT-PCR analysis of flow-sorted rainbow trout intestinal lymphocytes*

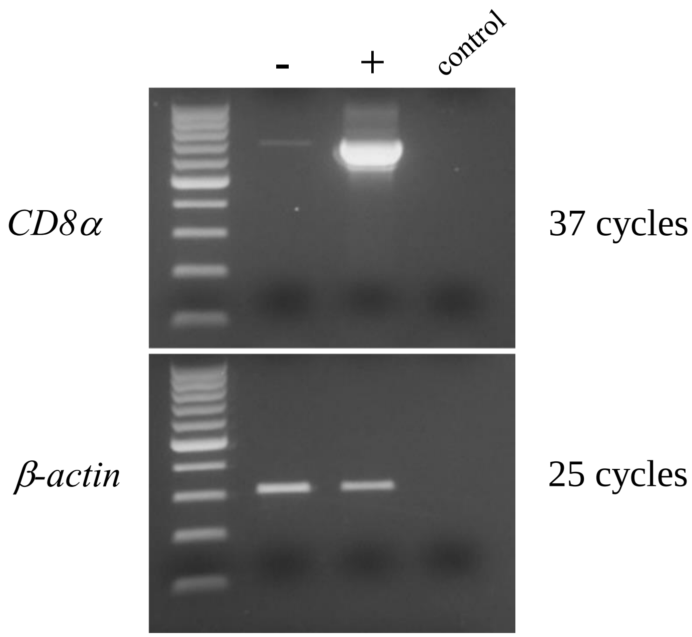

**Fig. S3B** Representative sortings of CD8α$^+$ versus CD8α$^-$ rainbow trout lymphocytes.

Leukocytes from thymus (Fig. S3B-1), intestine (Fig. S3B-2), head kidney (Fig. S3B-3), spleen (Fig. S3B-4) and gill (Fig. S3B-5) were stained with anti-CD8α mAb. The X-axis depicts relative cell size (FSC; forward scatter) while the Y-axis represents fluorescent intensity with the anti-CD8α mAb. The Y-axis is set to a biexponential scale. FSC$^{low}$ and SSC$^{low}$ lymphocytes were gated and fluorescence intensities from unsorted leukocytes (Pre-sort; left panel), sorted CD8α$^-$ lymphocytes (Negative; middle panel) and sorted CD8α$^+$ lymphocytes (Positive; right panel) were plotted. CD8α$^+$ lymphocytes are depicted in the upper right quadrants while the CD8α$^-$ cells appear in the lower right quadrants, respectively. Data are from a single representative of at least two independent experiments. In each experiment, leukocytes from 4 - 6 clonal individual trout were pooled.

*Fig. S3B-1. Rainbow trout thymocytes*

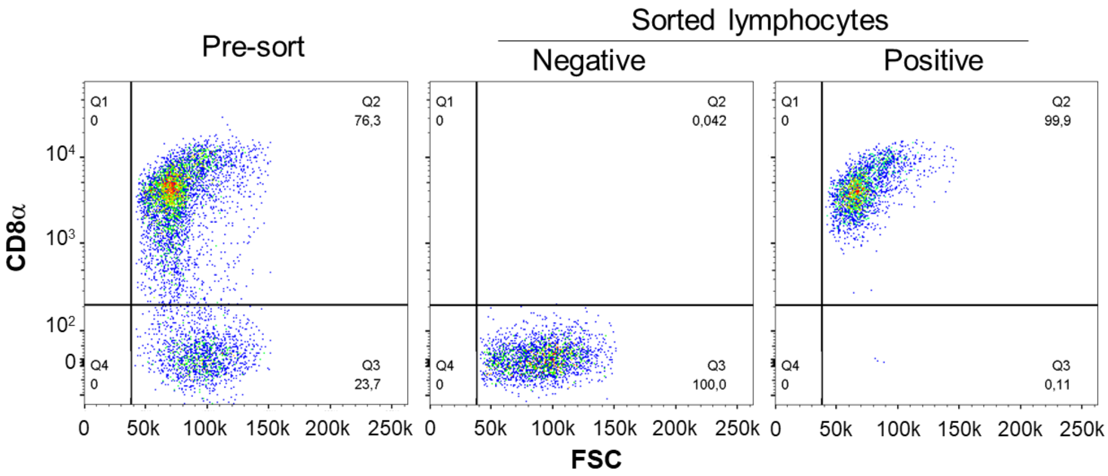


*Fig. S3B-2. Rainbow trout intestine lymphocytes*

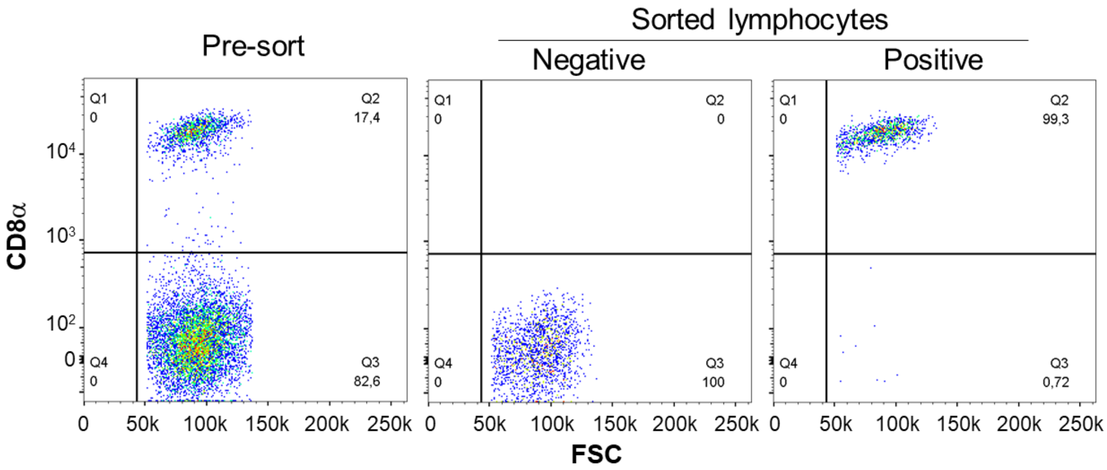

*Fig. S3B-3. Rainbow trout head kidney lymphocytes*

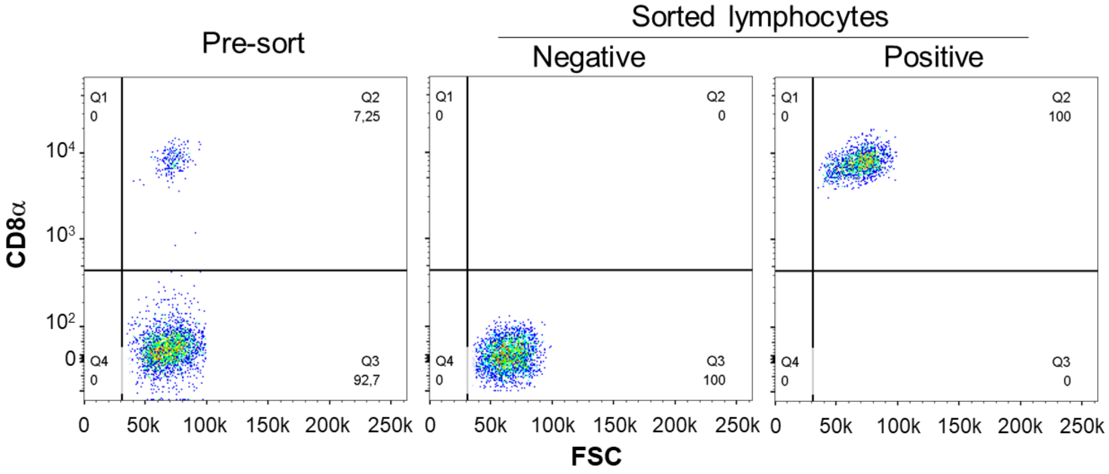


*Fig. S3B-4. Rainbow trout spleen lymphocytes*

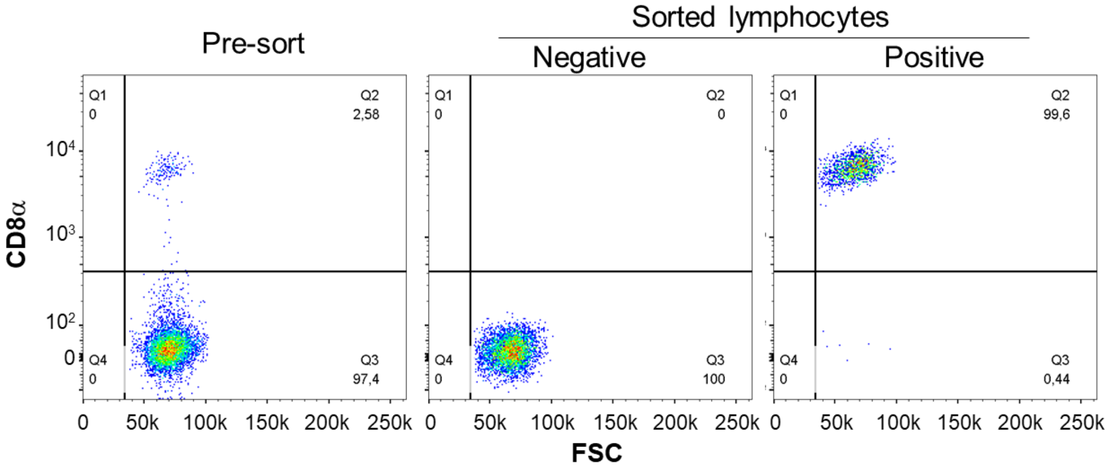


*Fig. S3B-5. Rainbow trout gill lymphocytes*

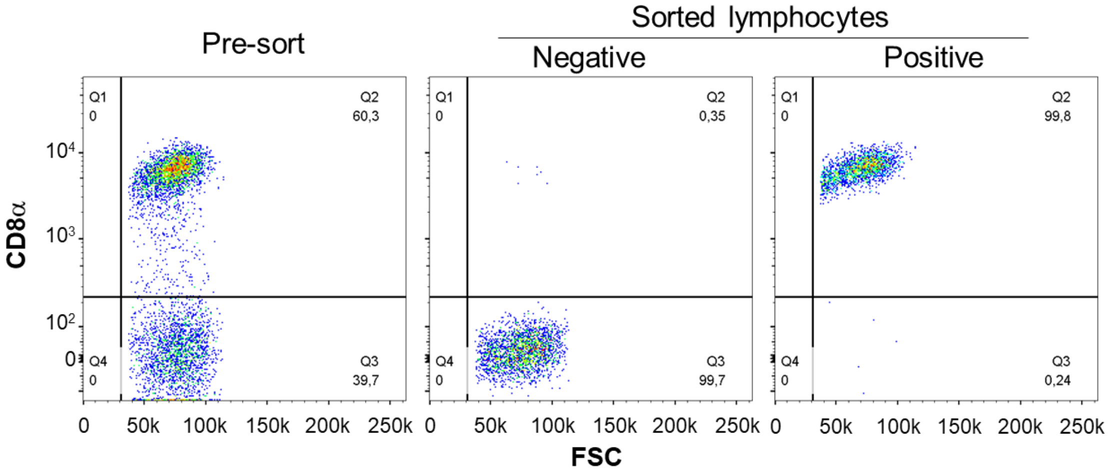

**Fig. S3C** Representative sortings of DN, DP, CD4SP and CD8SP rainbow trout lymphocytes from thymus, intestine and spleen.

Leukocytes from thymus (Fig. S5C-1), intestine (Fig. S5C-2) and spleen (Fig. S5C-3) were stained with anti-CD8α, anti-CD4-1 and anti-CD4-2 mAbs and sorted into the respective subpopulations. The X-axis depicts fluorescent intensity of staining with anti-CD8α mAb while the Y-axis represents fluorescent intensity with the anti-CD4 mAbs. Both axes are set to a biexponential scale. $FSC^{low}$ and $SSC^{low}$ lymphocytes were gated and their fluorescence intensities were plotted as follows: (S5C-1) stainings of thymocytes prior to sorting (Pre-sort; left panel), sorted DN thymocytes (upper middle panel), sorted DP thymocytes (upper right panel), sorted CD8SP thymocytes (lower middle panel) and sorted CD4SP thymocytes (lower right panel). (S5C-2 and 3) stainings of lymphocytes prior to sorting (Pre-sort; far left panel), sorted DN lymphocytes (left panel), sorted CD8SP lymphocytes (CD8SP; right panel) and sorted CD4SP lymphocytes (far right panel). Data are from a single representative of 5 independent experiments. In each experiment, leukocytes from 6 - 8 clonal individual trout were pooled.

*Fig. S3C-1. Rainbow trout thymocytes*

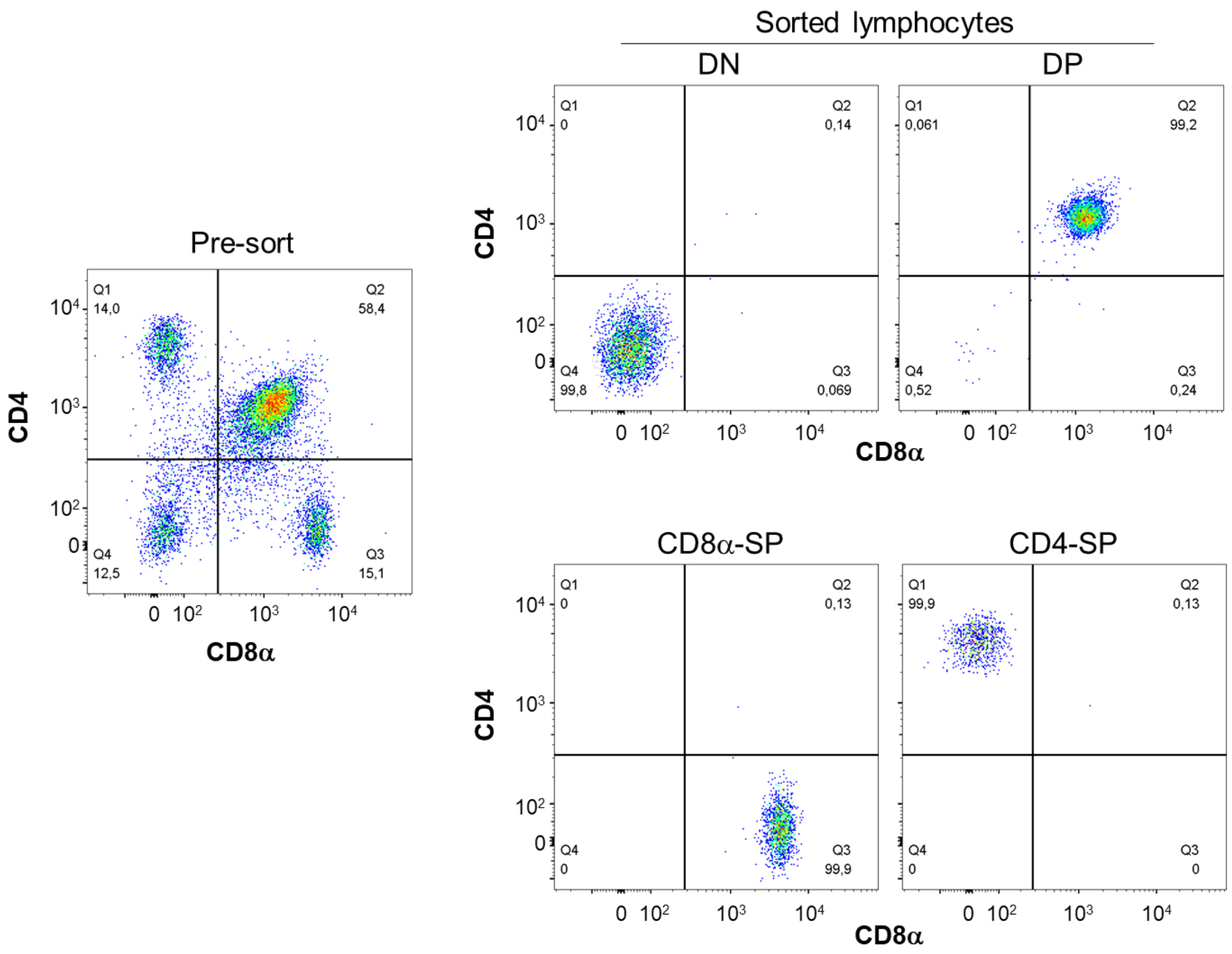

*Fig. S3C-2. Rainbow trout intestine lymphocytes*

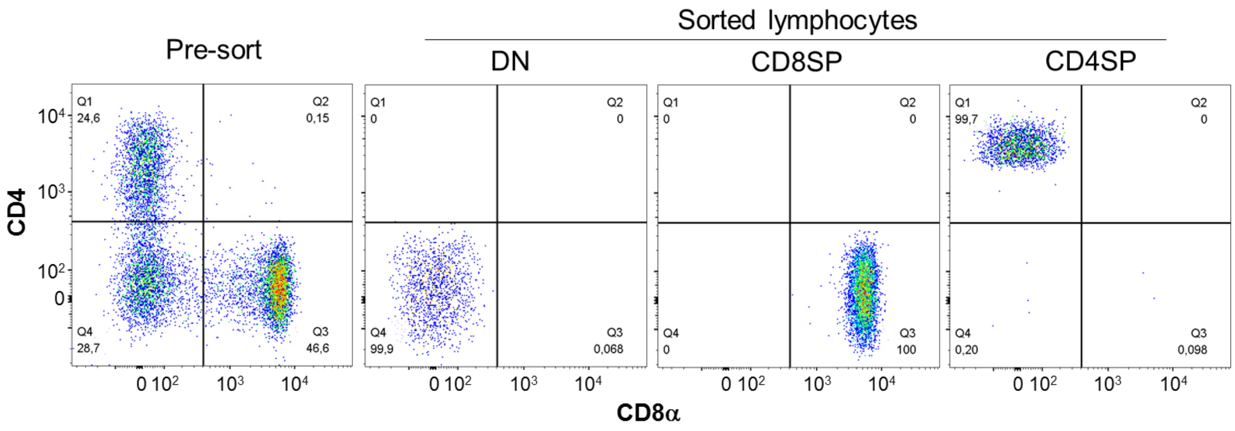


*Fig. S3C-3. Rainbow trout spleen lymphocytes*

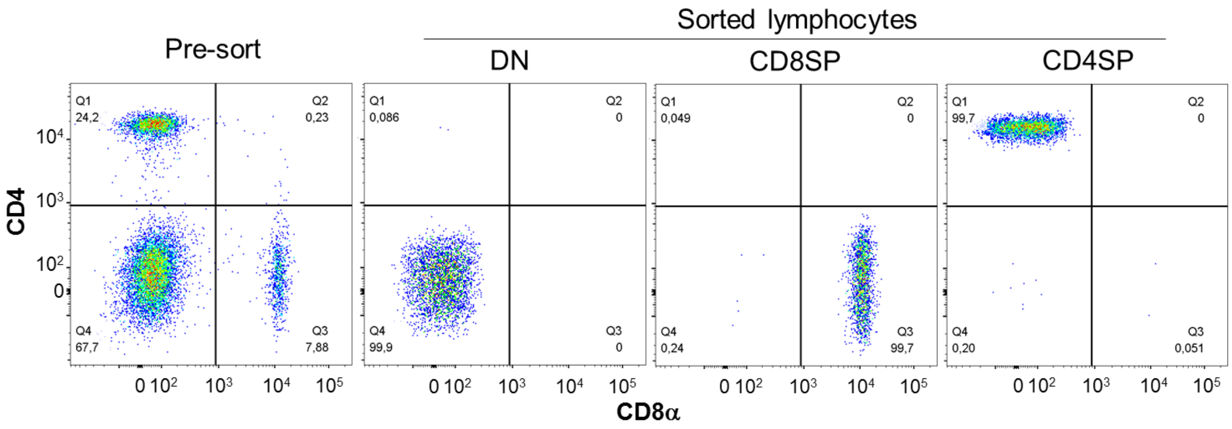

**Fig. S3D** Representative sorting of CD4SP, CD8SP, IgMSP and TN (triple negative; CD4⁻CD8⁻IgM⁻) rainbow trout splenocytes.

Splenocytes were stained with anti-CD8α, anti-CD4-1, anti-CD4-2 and anti-IgM mAbs and flow sorted. $FSC^{low}$ and $SSC^{low}$ lymphocytes were gated, and considered for further sorting steps. Representative dot plots show CD4 versus CD8α (Fig. S3D-1), IgM versus CD8α (Fig. S3D-2), and IgM versus CD4 (Fig. S3D-3). The fluorescence intensities of stainings prior to sorting (left Pre-sort panels) was plotted. $CD4^-CD8^-$ double negative (DN) populations were further gated and used for sorting into $IgM^+$ and $IgM^-$ splenocytes. Fluorescence intensities of sorted cells are depicted in the right panels: TN splenocytes (far left panels), sorted CD4SP (left panels), sorted CD8SP splenocytes (right panels) and sorted IgMSP splenocytes (far right panels). Both axes are set to a biexponential scale. Data are from a single representative of 5 independent experiments. In each experiment, leukocytes from 8 individuals were pooled.

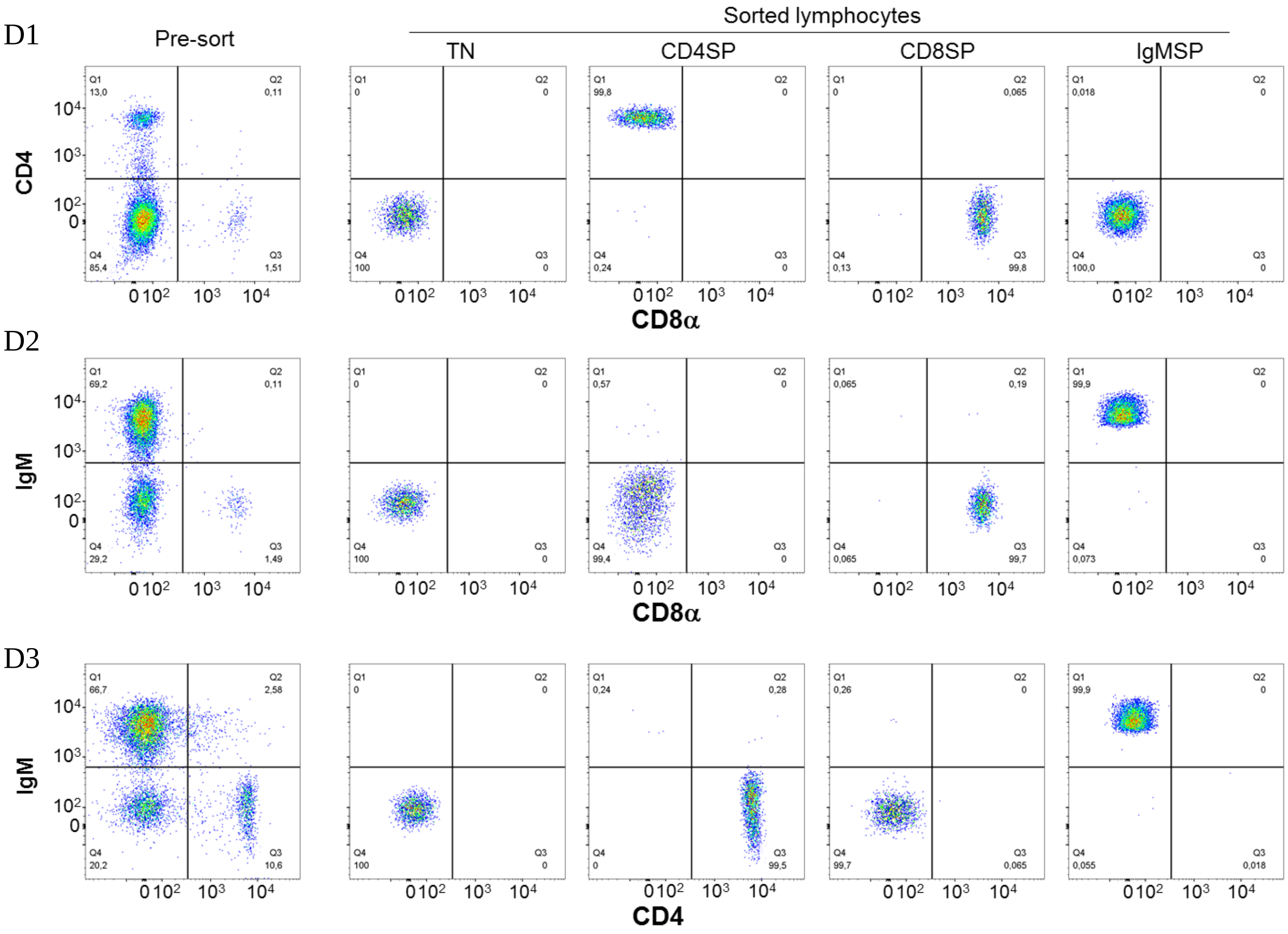

**Fig. S3E** Percentages of CD4SP, CD8SP, IgMSP and TN cells among rainbow trout spleen lymphocytes.

For the experiments of which the results are shown in main text Fig. 11, spleens of 8 individuals were pooled in four independent experiments. For one of those experiments, the sorting results into CD4SP, CD8SP, IgMSP and TN cell subpopulations are shown in Fig. S3D. The tables below summarize the percentages of subpopulation cells relative to spleen lymphocytes for each of the four experiments. Table (a) shows the raw data provided by the cell sorter, and Table (b) shows calculated percentages of $IgM^+$ and TN splenocytes, based on dividing the percentage of DN splenocytes according to the ratio of $IgM^+$ and $IgM^-$ splenocytes.

(a)

| % in lymphocyte gate | CD8SP | CD4SP | DN | | within DN | |
|---|---|---|---|---|---|---|
| | | | | | IgM+ | IgM- (TN) |
| #1 | 7.6 | 29.3 | 62.9 | | 53.3 | 46.7 |
| #2 | 6.1 | 23.4 | 70.3 | | 52.8 | 47.2 |
| #3 | 6.5 | 22.0 | 71.4 | | 59.8 | 40.2 |
| #4 | 5.7 | 20.2 | 74.0 | | 56.0 | 44.0 |

(b)

| % in lymphocyte gate | CD8SP | CD4SP | IgMSP | TN |
|---|---|---|---|---|
| #1 | 7.6 | 29.3 | 33.5 | 29.4 |
| #2 | 6.1 | 23.4 | 37.1 | 33.2 |
| #3 | 6.5 | 22.0 | 42.7 | 28.7 |
| #4 | 5.7 | 20.2 | 41.4 | 32.6 |
| Av. | **6.5** | **23.7** | **38.7** | **31.0** |

### Supplementary figure 4 (Fig. S4)

Preparations of recombinant trout cytokines, soluble IL-15Rα (sIL-15Rα) and RLI fusion proteins from insect cells

**Table of Contents** **Page**


Fig. S4A: Co-purification by anti-FLAG agarose reveals binding of trout IL-15 to trout sIL-15Rα 60

Fig. S4B: Co-purification by anti-FLAG agarose reveals binding of trout IL-15La to trout sIL-15Rα 61

Fig. S4C: Co-purification by anti-FLAG agarose reveals binding of trout IL-2 to trout sIL-15Rα 62

Fig. S4D: Analysis of purified recombinant trout IL-15-RLI 63

Fig. S4E: Analysis of purified recombinant trout IL-15La-RLI 66

Fig. S4F: Analysis of purified recombinant trout IL-2 69

**Fig. S4A** Co-purification by anti-FLAG agarose reveals binding of trout IL-15 to trout sIL-15Rα.

Approximately 100 ml of supernatant each of insect cells infected with recombinant baculovirus(es) for inducing expression of FLAG-tagged trout IL-15 and/or Myc-tagged trout sIL-15Rα (IL-15, sIL-15Rα, and IL-15+sIL-15Rα) were purified using agarose-bound anti-FLAG and chromatography columns. The flow-through sample from the column was collected ("Flow-through"). After washing, elution and a final buffer exchange for 300 μl PBS, the purified product was obtained ("Eluted"). Of both the Eluted and Flow-through samples 15 μl per lane was analyzed by SDS-PAGE followed by anti-FLAG (a) and anti-Myc (b) Western blot (WB) analyses. M, size marker. The results for (a) and (b) were obtained using separate gels.

The results show that, in case alone, sIL-15Rα could not be retained and subsequently eluted using the anti-FLAG agarose [see lane sIL-15Rα-Eluted in (b)], whereas IL-15 was readily detectable in the Eluted fraction [see lane IL-15-Eluted in (a)]. In contrast, from the supernatants in which IL-15 and sIL-15Rα were present together (IL-15+sIL-15Rα), also sIL-15Rα was readily detectable in the Eluted fraction [see lane IL-15+sIL-15Rα-Eluted in (b)]. The results provide evidence for binding between trout IL-15 and IL-15Rα.

(a) *anti-FLAG WB for detection of IL-15*

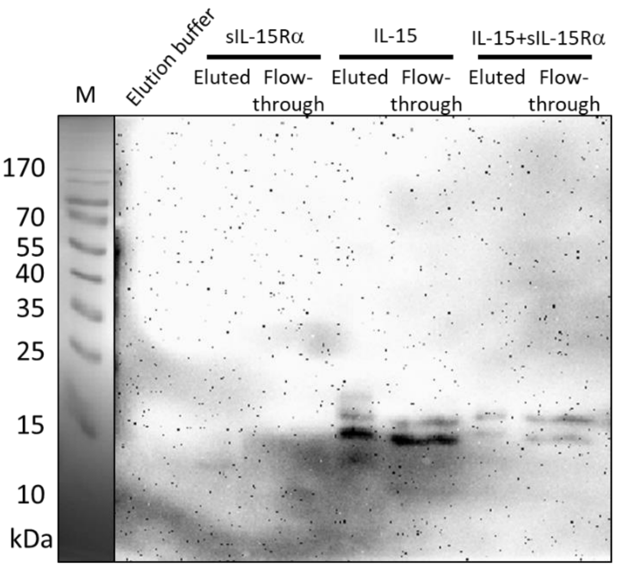


(b) *anti-Myc WB for detection of sIL-15Rα*

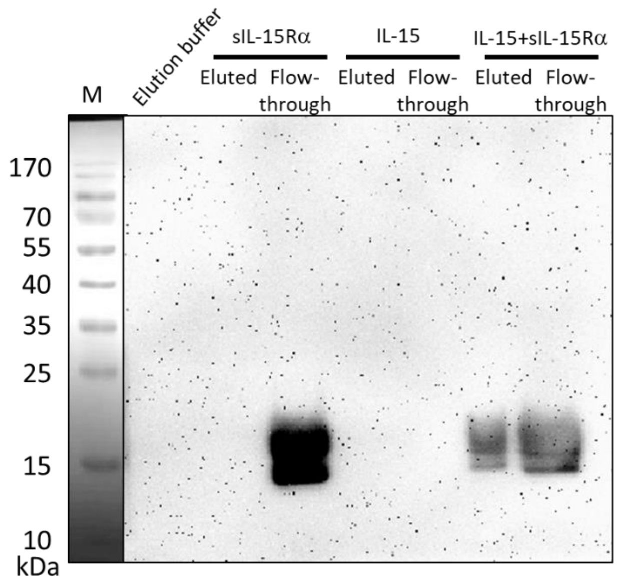

**Fig. S4B** Co-purification by anti-FLAG agarose reveals binding of trout IL-15La to trout sIL-15Rα.

Approximately 100 ml of supernatant of insect cells infected with recombinant baculoviruses for inducing expression of FLAG-tagged trout IL-15La plus Myc-tagged trout sIL-15Rα was purified using agarose-bound anti-FLAG and chromatography columns. The flow-through sample from the column was collected ("Flow-through"). After washing, elution and a final buffer exchange for 300 μl PBS, the purified product was obtained ("Eluted"). Of both the Eluted and Flow-through samples 15 μl per lane was analyzed by SDS-PAGE followed by non-specific protein staining using Coomassie blue (a), as well as by anti-FLAG (b) and anti-Myc (c) Western blot (WB) analyses. M, size marker. The results for (a), (b) and (c) were obtained using separate gels.

The results indicate that both IL-15La and sIL-15Rα were efficiently eluted. Because, under similar conditions, from supernatants in which sIL-15Rα was present alone the sIL-15Rα protein was not detectably eluted [see Fig. S4A(b)], the results provide evidence for binding between trout IL-15La and IL-15Rα. The higher weight band (or bands) detected with anti-FLAG WB might be a homodimer form (or forms) of IL-15La.

(a) *Coomassie blue staining*

(b) *anti-FLAG WB for detection of IL-15La*

(c) *anti-Myc WB for detection of sIL-15Rα*

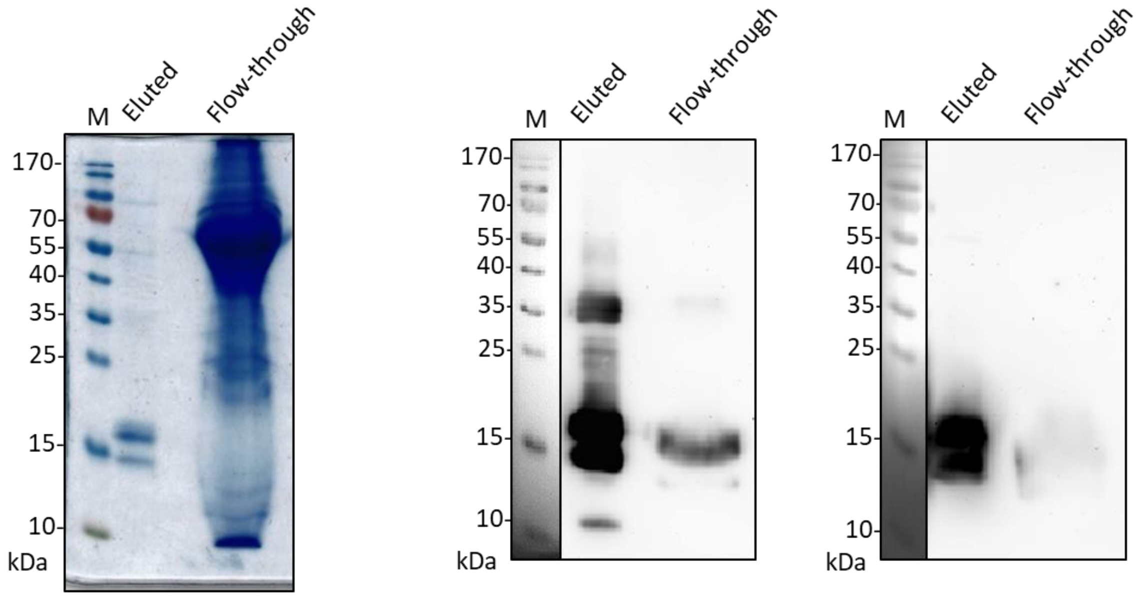

**Fig. S4C** Co-purification by anti-FLAG agarose reveals binding of trout IL-2 to trout sIL-15Rα.

Approximately 100 ml of supernatant of insect cells infected with recombinant baculoviruses for inducing expression of FLAG-tagged trout IL-2 plus Myc-tagged trout sIL-15Rα was purified using agarose-bound anti-FLAG and chromatography columns. The flow-through sample from the column was collected ("Flow-through"). After washing, elution and a final buffer exchange for 300 µl PBS, the purified product was obtained ("Eluted"). Of both the Eluted and Flow-through samples 15 µl per lane was analyzed by SDS-PAGE followed by non-specific protein staining using Coomassie blue (a), and anti-FLAG (b) and anti-Myc (c) Western blot (WB) analyses. M, size marker. The results for (a), (b) and (c) were obtained using separate gels.

The results indicate that both IL-2 and sIL-15Rα were efficiently eluted. Because, under similar conditions, from supernatants in which sIL-15Rα was present alone the sIL-15Rα protein was not detectably eluted [see Fig. S4A(b)], the results provide evidence for binding between trout IL-2 and IL-15Rα. The higher weight band (or bands) detected with anti-FLAG WB might be a homodimer form (or forms) of IL-2, and/or maybe a complex of IL-2 and sIL-15Rα.

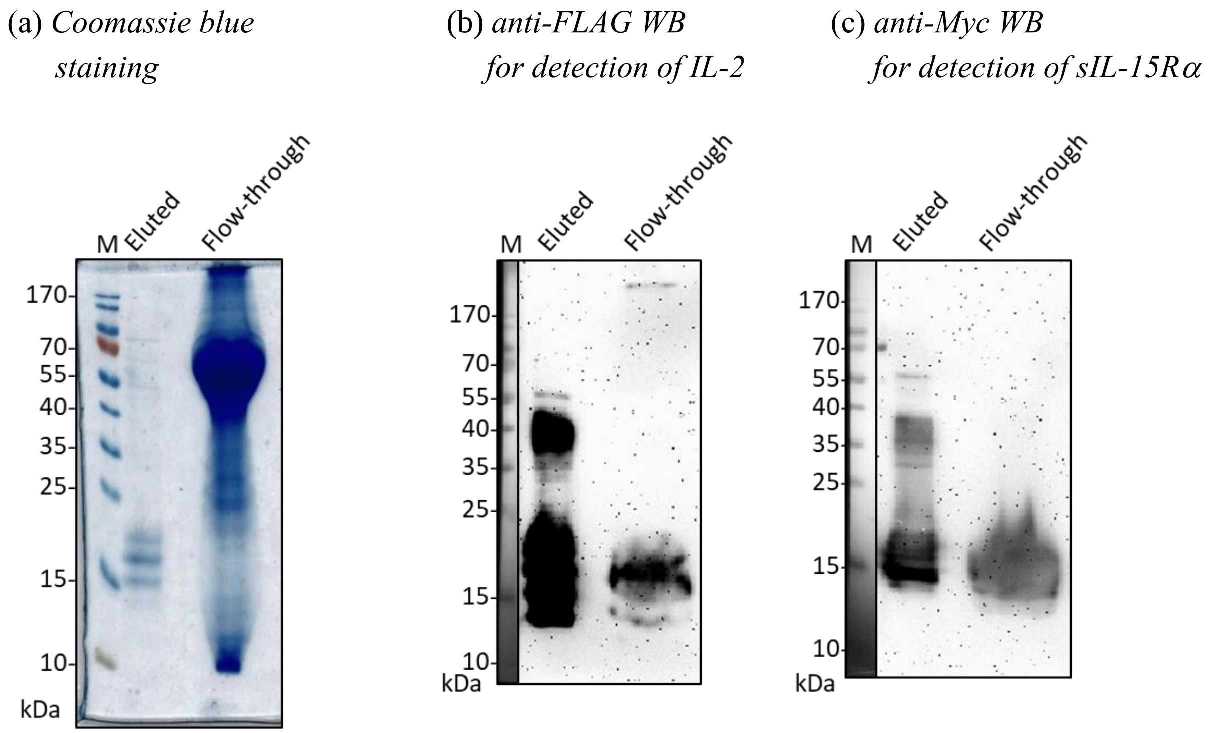

**Fig. S4D** Analysis of purified recombinant trout IL-15-RLI.

Approximately 300 ml of supernatant of insect cells infected with recombinant baculovirus for inducing expression of FLAG-tagged trout IL-15-RLI (a fusion protein of trout IL-15 and trout sIL-15Rα) was purified using agarose-bound anti-FLAG and chromatography columns. After washing, elution and a final buffer exchange for 300 μl PBS, the purified product was obtained ("Purified IL-15-RLI"). Part of this product was mixed with glycerol and BSA and stored at -20 °C for later use in functional assays, while another part was investigated for preparation quality by size exclusion chromatography (aka "gel filtration") [(a), with (b) for estimating the protein size corresponding with the major peak F2 in (a)], and by SDS-PAGE followed by non-specific protein staining using Coomassie blue (c) and anti-FLAG Western blot analysis (d). Not only the "Purified IL-15-RLI" sample was analyzed by Coomassie blue staining and Western blot analysis, but also the fractions F1, F2 and F3 which were eluted using gel filtration (a) and concentrated before loading onto gels for SDS-PAGE [(c) and (d)]. (b) Molecular weights of the purified recombinant proteins were estimated by gel filtration. The red dot represents the F2 peak shown in (a) and blue dots represent the following standard proteins: bovine serum albumin, 66 kDa; carbonic anhydrase, 29 kDa; cytochrome c, 12.4 kDa; aprotinin, 6.5 kDa. (e) N-glycosylation of proteins in the purified IL-15-RLI preparation was shown by anti-FLAG Western blot analysis of samples of this preparation that were not treated [Purified IL-15-RLI], subjected to treatment with PNGaseF [Purified IL-15-RLI (+)], or subjected to the corresponding mock treatment [Purified IL-15-RLI (-)] (e). For the SDS-PAGE analyses in (c) and (d), PBS was loaded as negative control [(c) and (d)]. M, size marker. The results for (c) and (d) were obtained using separate gels and loading 15 μl sample per lane, and the results for (e) were obtained by loading amounts corresponding to 1.2 μl (~160 ng) of the purified IL-15-RLI sample per lane.

The results showed that the purified IL-15-RLI preparation was rather pure [see the "Purified IL-15-RLI" lane in (c)]. The bands with an apparent molecular weight of ~35 kDa observed upon SDS-PAGE analysis [(c), (d) and (e)] can be concluded to include N-glycosylated (e) monomeric proteins, considering that the predicted molecular weight of the protein backbone (without leader sequence) is 26.4 kDa. Upon gel filtration the bulk of the protein behaved as having an approximate molecular weight of 42.6 kDa [(a) and (b)] which agrees best with the assumption that the bulk of the protein is present as a soluble monomer. Upon sensitive Western blot analysis, a small band of ~13 kDa was detected which may represent a breakdown product.

*Fig. S4D*

(a) *Gel filtration analysis of purified IL-15-RLI*

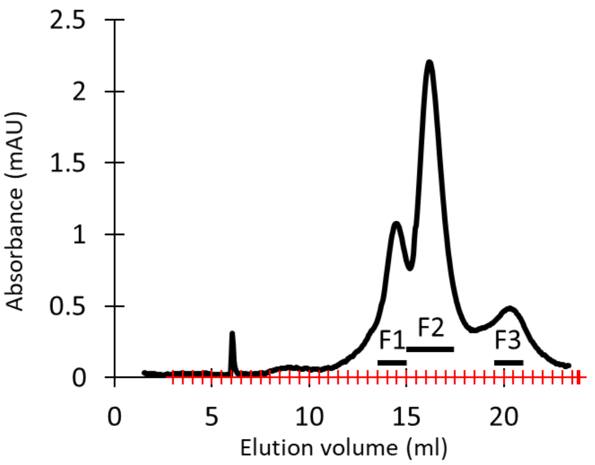


(b) *Gel filtration analysis for estimating the size of proteins in the F2 peak in (a)*

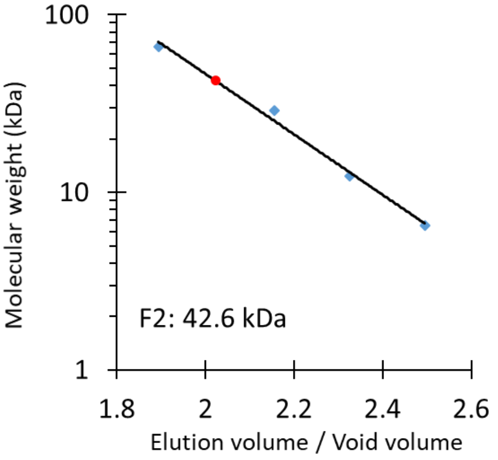


(c) *Coomassie blue staining*

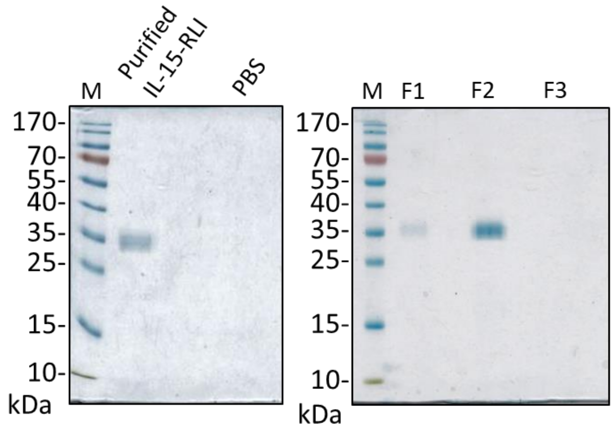


(d) *Anti-FLAG Western blot analysis*

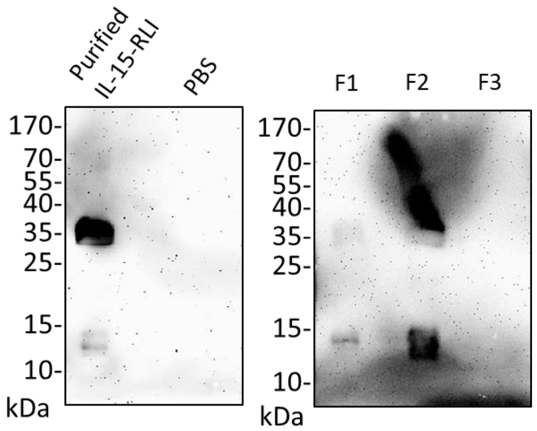

*(Fig. S4D continued)*

(e) *Analysis of N-glycosylation by PNGaseF treatment*

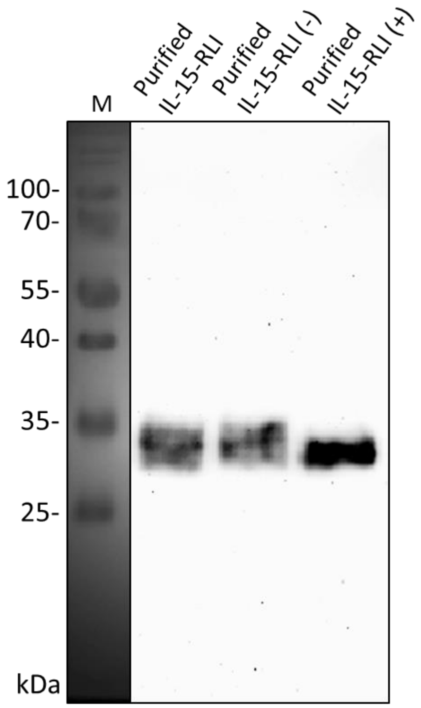

#### **Fig. S4E** Analysis of purified recombinant trout IL-15La-RLI.

Isolation from insect cells and subsequent analysis of recombinant FLAG-tagged IL-15La-RLI (a fusion product of trout IL-15La and trout sIL-15Ra) was performed as done for IL-15-RLI (see the legend of Fig. S4D). The results are shown in figures (a)-to-(e).

The results showed that the purified IL-15La-RLI preparation was rather pure [see the "Purified IL-15La-RLI" lane in (c)]. The bands with an apparent molecular weight of ~35 kDa observed upon SDS-PAGE analysis [(c), (d) and (e)] can be concluded to include N-glycosylated (e) monomeric proteins, considering that the predicted molecular weight of the protein backbone (without leader sequence) is 26.0 kDa. Upon gel filtration the bulk of the protein behaved as having an approximate molecular weight of 31.4 kDa [(a) and (b)] which agrees best with the assumption that the bulk of the protein is present as a soluble monomer. Upon sensitive Western blot analysis, a band of ~70 kDa was detected in the purified IL-15La-RLI preparation which may represent a homodimer (d). Western blot analysis of the F1 fraction isolated by gel filtration detected a small band of ~17 kDa which may represent a breakdown product resulting from treatment (d).

(a) *Gel filtration analysis of purified IL-15La-RLI*

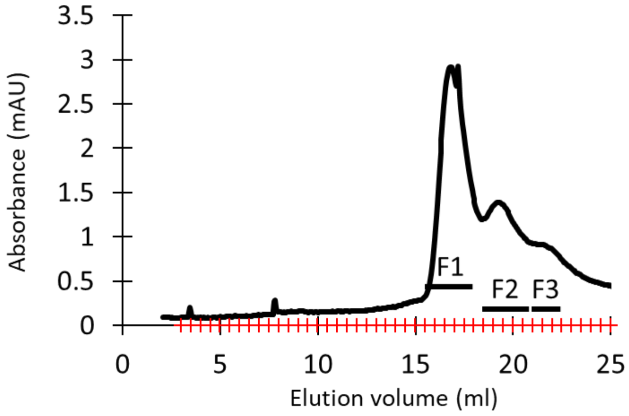


(b) *Gel filtration analysis for estimating the size of proteins in the F1 peak in (a)*

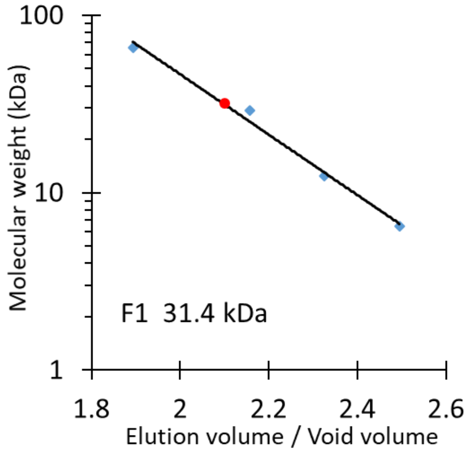

*(Fig. S4E continued)*

(c) *Coomassie blue staining*

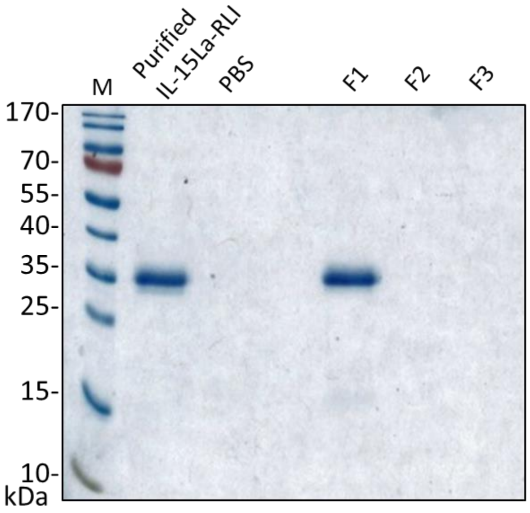


(d) *Anti-FLAG Western blot analysis*

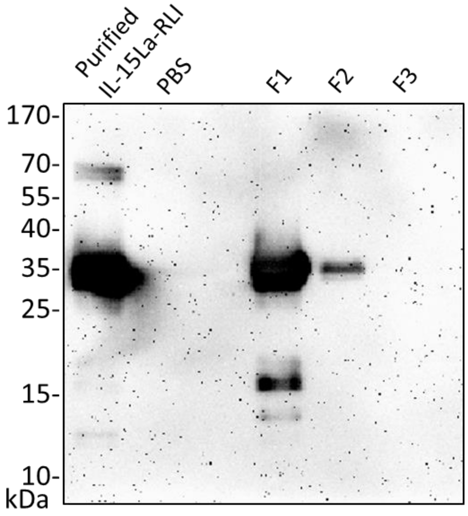


(e) *Analysis of N-glycosylation by PNGaseF treatment*

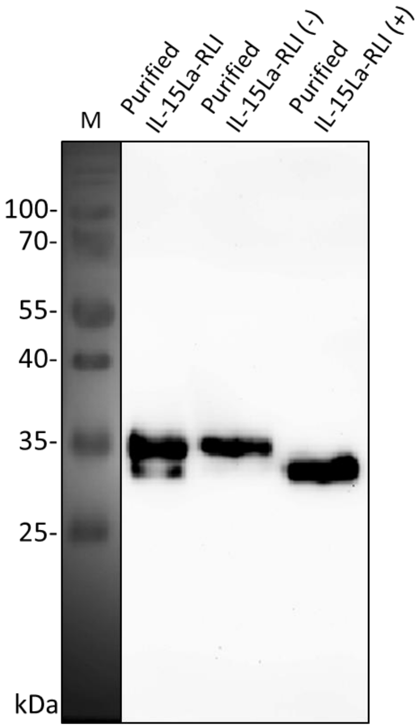

#### **Fig. S4F** Analysis of purified recombinant trout IL-2.

Isolation from insect cells and subsequent analysis of recombinant FLAG-tagged trout IL-2 was performed as done for IL-15-RLI (see the legend of Fig. S4D). The results of those analyses are shown in figures (a)-to-(e).

The results showed that the purified IL-2 preparation was rather pure [see the "Purified IL-2" lane in (c)]. The four major bands in the purified IL-2 preparation with an apparent molecular weight of approximately 13-19 kDa that were observed upon SDS-PAGE analysis [(c) and (d)] may include different levels of glycosylation of monomeric proteins, since treatment with PNGaseF affected the running behavior of at least the two biggest protein among these bands (e) and the predicted molecular weight of the protein backbone (without leader sequence) is 13.2 kDa. We do not understand all four bands in the 13-19 kDa range, and they may also represent other modifications besides N-glycosylation and/or breakdown products. Upon gel filtration the bulk of the protein behaved as having an approximate molecular weight of 31.4 kDa [(a) and (b); (c) shows that peak F4 contained most protein] which agrees best with the assumption that the bulk of the protein is present in soluble homodimer form. Upon sensitive Western blot analysis, a band of ~40 kDa was detected in the purified IL-2 preparation which may represent a homodimer form (d).

(a) *Gel filtration analysis of purified IL-2*

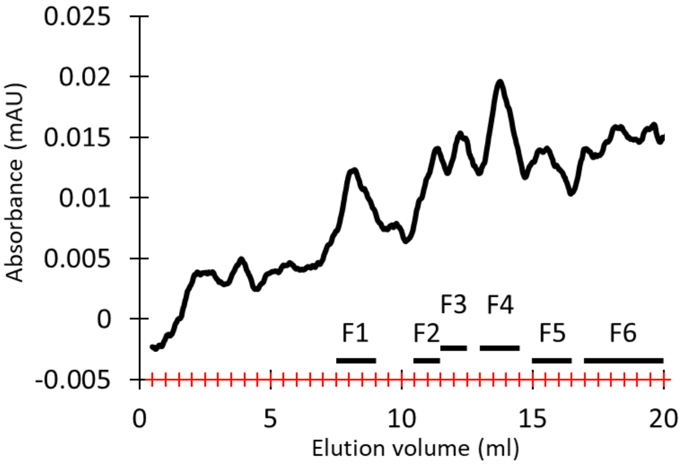


(b) *Gel filtration analysis for estimating the size of proteins in the F3 and F4 peaks (a)*

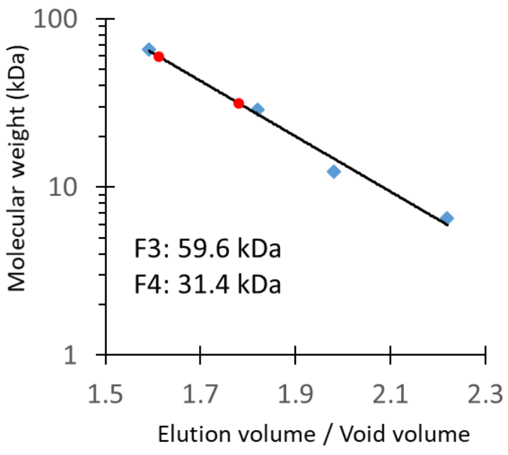

*(Fig. S4F continued)*

(c) *Coomassie blue staining*

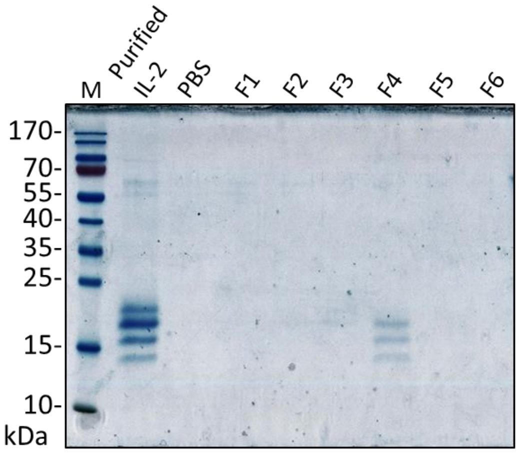


(d) *Anti-FLAG Western blot analysis*

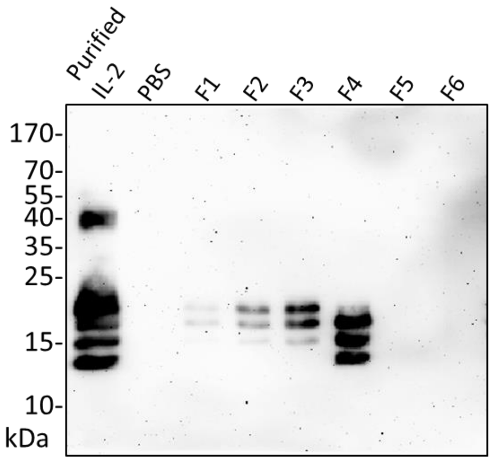


(e) *Analysis of N-glycosylation by PNGaseF treatment*

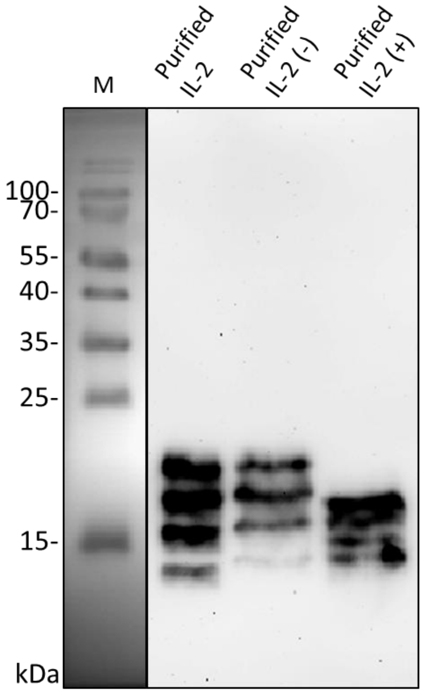

### Supplementary figure 5 (Fig. S5)

Additional Western blot results

**Table of Contents** **Page**

| | |
|---|---|
| Fig. S5A: Cytokine deglycosylation analyses *(extended version of main text Fig. 4)* | 72 |
| Fig. S5B: Secretion of recombinant cytokines and alpha receptor chains from transfected HEK293T cells (*information supporting the main text Fig. 6 results*) | 74 |
| Fig. S5C: Phosphorylation of STAT5 in various trout lymphocyte populations induced after incubation with recombinant trout cytokine containing HEK293T supernatants (*information supporting the main text Fig. 7 results*) | 80 |
| Fig. S5D: Phosphorylation of STAT5 in DN, DP, CD4SP and CD8SP fractions of trout thymocytes induced after incubation with recombinant cytokine containing HEK293T supernatants (*information supporting the main text Fig. 8 results*) | 92 |

Fig. S5E: Phosphorylation of STAT5 in lymphocyte fractions of trout thymus, intestine and spleen, induced after incubation with purified recombinant trout cytokines (produced in insect cells) IL-2, IL-15-RLI and IL-15La-RLI at 5, 25 and 125 nM (*information supporting the main text Fig. 9 results*) 95

Fig. S5F: Titration of the recombinant trout cytokine (produced in insect cells) concentration necessary for the induction of pSTAT5 98

**Fig. S5A.** Cytokine deglycosylation analyses *(extended version of main text Fig. 4)*.

Bovine IL-2 and IL-15, and trout IL-2, IL-15 and IL-15La, can be N-glycosylated.

Lysates of HEK293T cells that were transfected with DNA plasmids encoding FLAG-tagged cytokines were incubated overnight in digestion buffer (+) or not (-), and the incubated samples were digested with PNGase-F (+) or not (-). After that, the samples were analyzed by anti-FLAG Western blotting. Figures S5A-1 and S5A-2 show uncropped versions of the blot results of which parts of the lanes for the PNGase-F treated and mock treated samples are shown in main text Fig. 4; in addition to the lanes shown in Fig. 4, results for a PNGase-F treated untransfected HEK293T lysate control (the left lanes) and a transfected but untreated control (the second lanes from the left) are included. (Fig. S5A-1) Bovine IL-2 and IL-15, but not bovine IL-15L, show a shift in apparent molecular weight consistent with removal of N-linked sugar chains. (Fig. S5A-2) Trout IL-2, IL-15, and IL-15La, but not trout IL-15Lb, show a shift in apparent molecular weight consistent with removal of N-linked sugar chains. Some of the bands were also sensitive to mock treatment alone (e.g. the largest bovine IL-2 band, the smallest trout IL-15 band, and the trout IL-15Lb band), possibly because of aggregation. For the untreated sample of trout IL-15La expressing HEK293T cells, a band of ~37 kDa is observed which might represent an IL-15La homodimer.

**Fig. S5A-1.** Bovine IL-2 and IL-15 can be N-glycosylated.

*anti-FLAG Western blots*

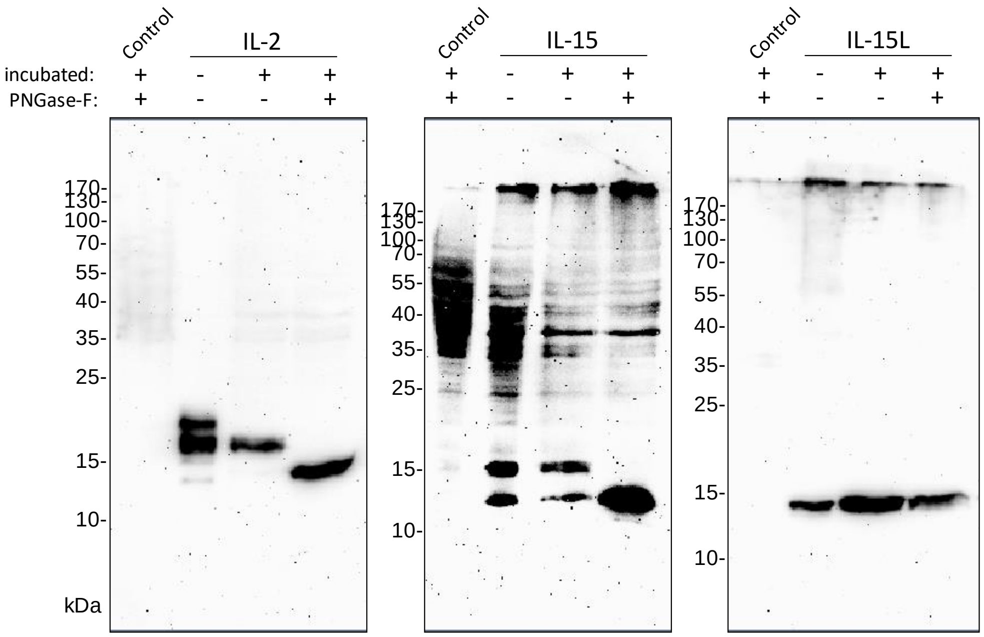


**Fig. S5A-2.** Trout IL-2, IL-15 and IL-15La can be N-glycosylated.

*anti-FLAG Western blots*

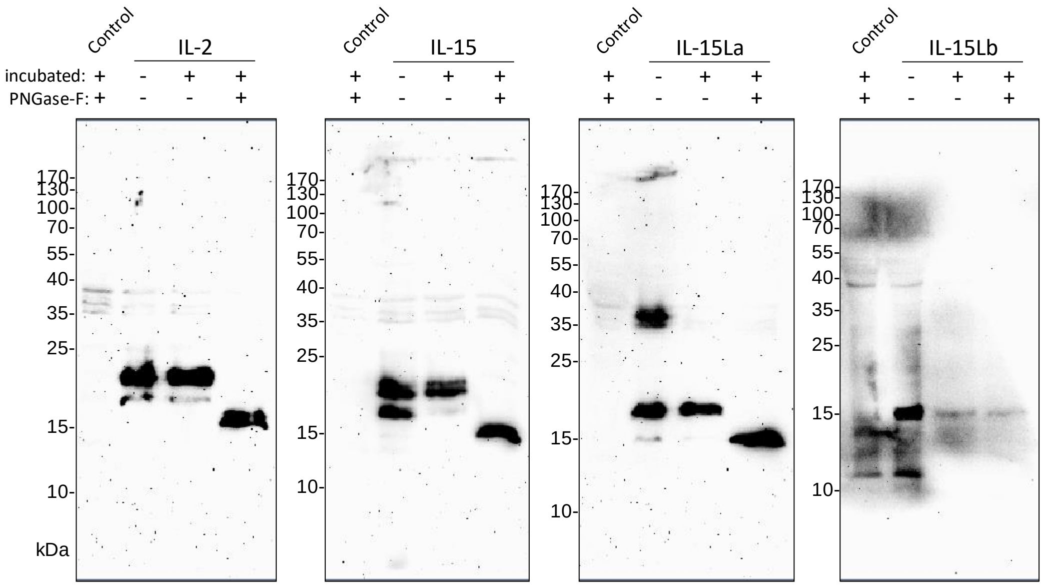

**Fig. S5B.** Secretion of recombinant cytokines and alpha receptor chains from transfected HEK293T cells *(information supporting the main text Fig. 6 results).*

Co-expression with soluble IL-15Rα causes enhanced presence of bovine and trout IL-15 and IL-15L in the supernatant of transfected cells.

HEK293T cells were transfected with DNA plasmids for expression of FLAG-tagged cytokines and/or species-specific (bovine in the bovine panel, trout in the trout panel) Myc-tagged soluble IL-2Rα or IL-15Rα. Matching supernatants (upper blots; the supernatants were concentrated before loading) and cell lysates (lower blots) were compared by Western blot analyses using antibodies against FLAG (left blots) or Myc (right blots). The data shown are from two independent experiments for expression of the bovine molecules (Cattle experiments 1 and 2), and three independent experiments for expression of the trout molecules (Trout experiments 1, 2 and 3). Cropped versions of the anti-FLAG results of Cattle experiment 1 and Trout experiment 1 are shown in main text Fig. 6; a reason for the cropping was because the uncropped blot results include data for RLI protein versions in which modified bovine IL-15L or trout IL-15La were linked with human sIL-15Rα (bov.IL-15Lhyb-h-RLI and IL-15La-h-RLI). In our experiments we first used a combination of human sIL-15Rα and trout IL-15La in order to more precisely copy the previously published all-human RLI version (39), and only after we had found stability and function this fusion version (e.g. Figs. S5B-3, S5B-4 and main text Fig. 8) we progressed to making a fusion between trout IL-15La and trout sIL-15Rα which was produced in insect cells (IL-15La-RLI used for the experiments resulting in main text Figs. 9-11) We deleted the RLI-form results from the blot pictures shown in main text Fig. 6 because the explanation would have needed too much text space and might be confusing to readers. For more explanation of the bovine IL-15L modification in the bov.IL-15Lhyb-h-RLI sequence see Fig. S2. In the cell lysates with trout IL-15La a band of ~37 kDa is observed which might represent an IL-15La homodimer (Figs. S5B3-to-5).

**Fig. S5B-1.** Cattle, experiment 1.

*Western blots*

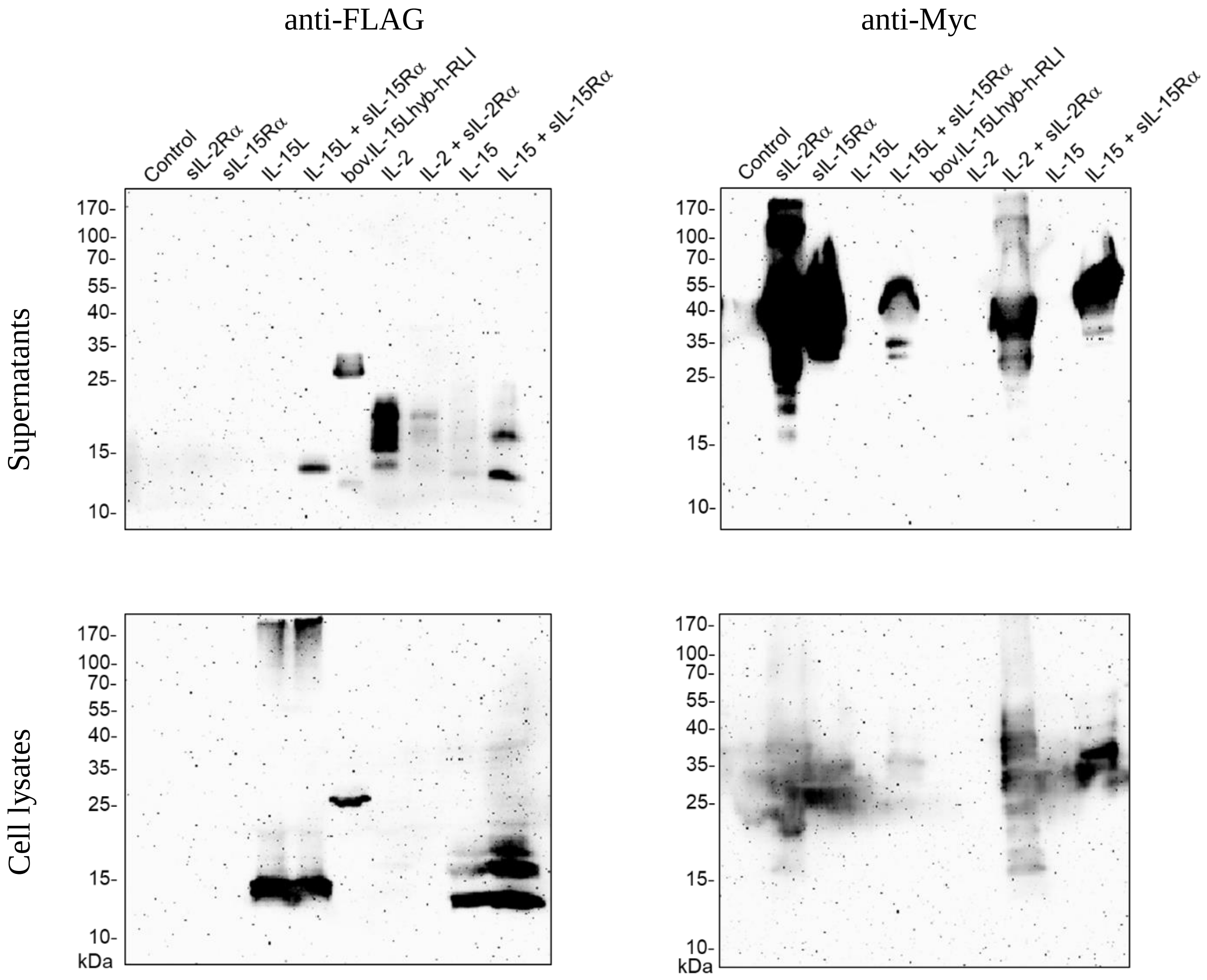

**Fig. S5B-2.** Cattle, experiment 2.

*Western blots*

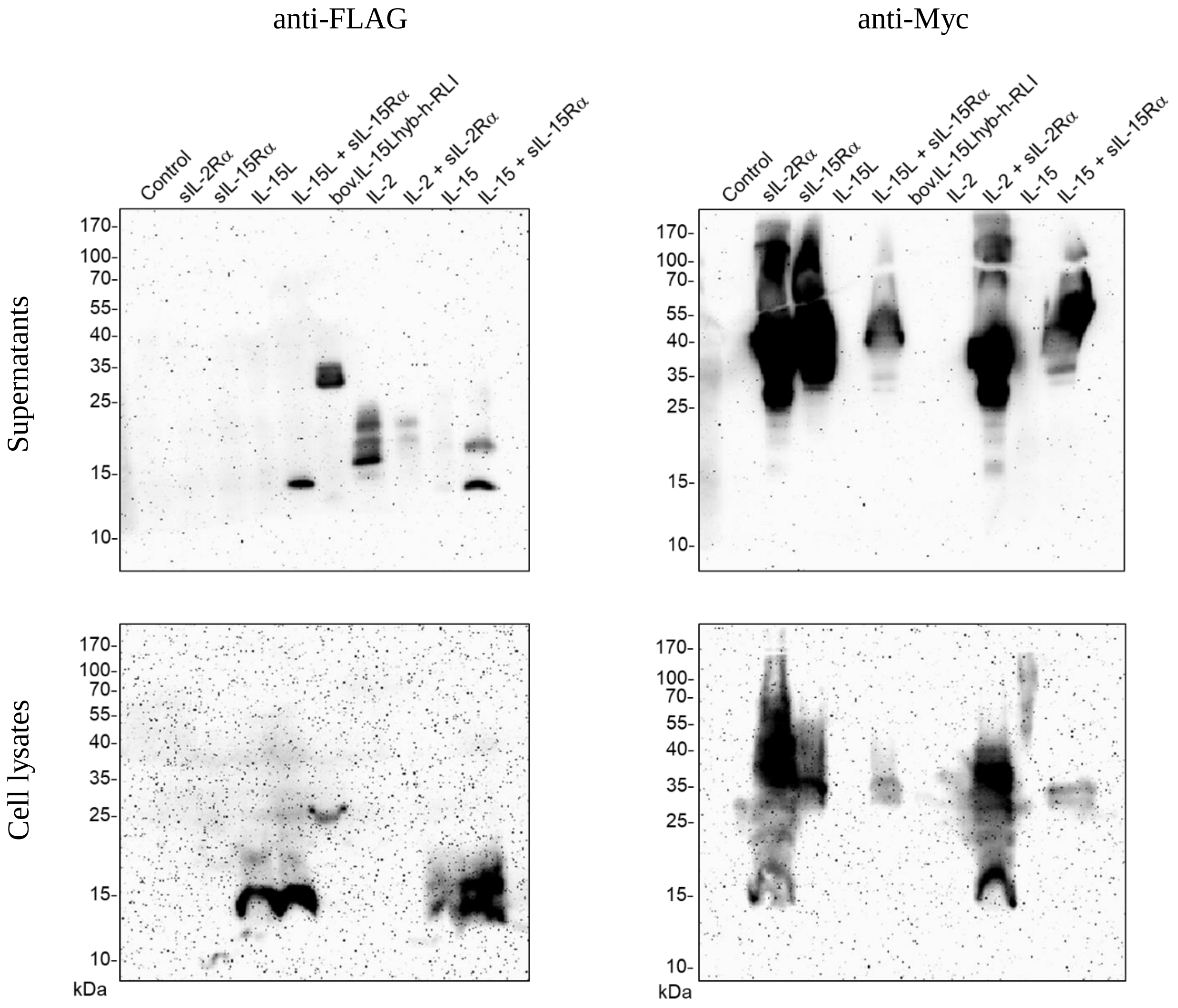

**Fig. S5B-3.** Trout, experiment 1.

*Western blots*

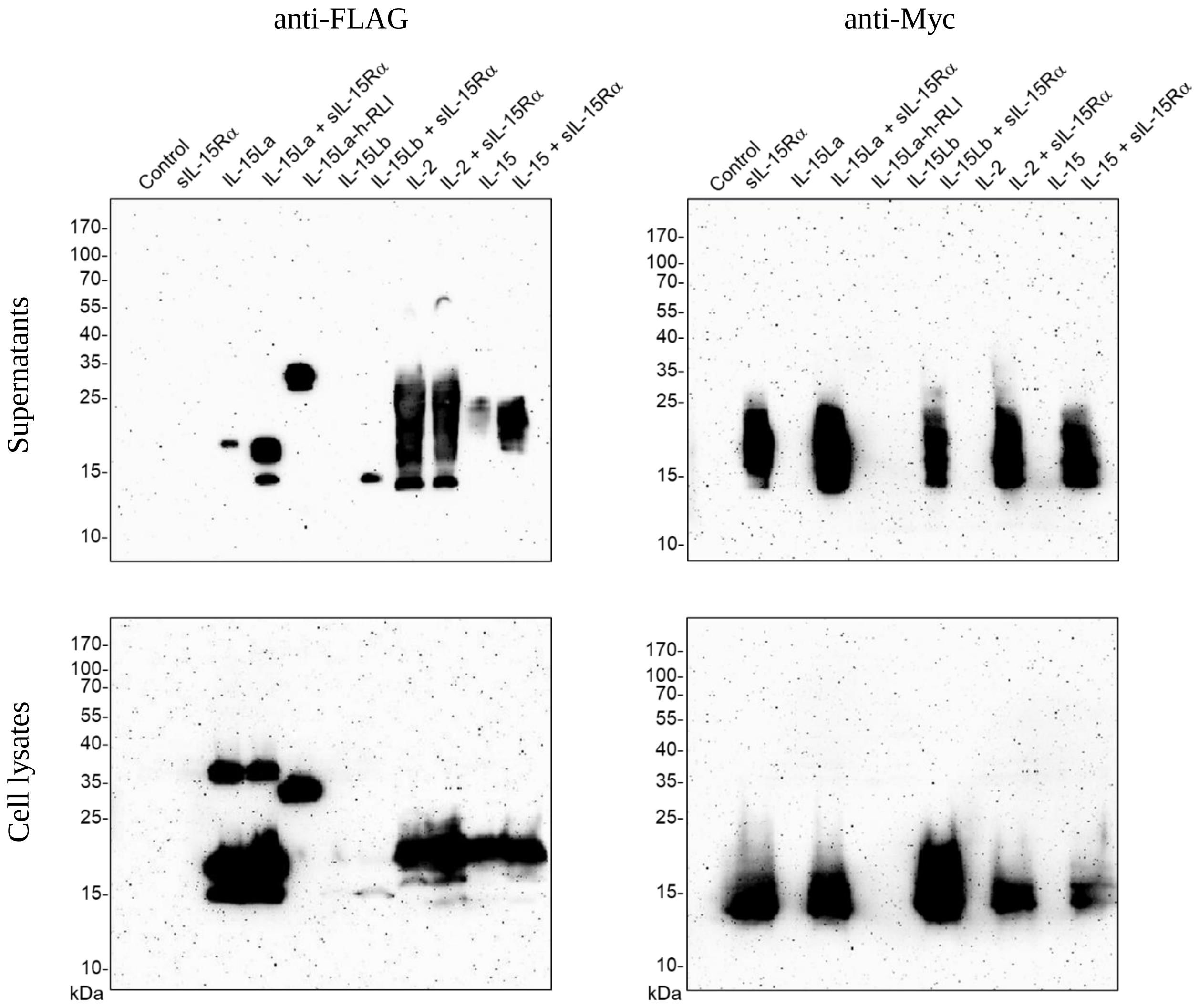

**Fig. S5B-4.** Trout, experiment 2.

*Western blots*

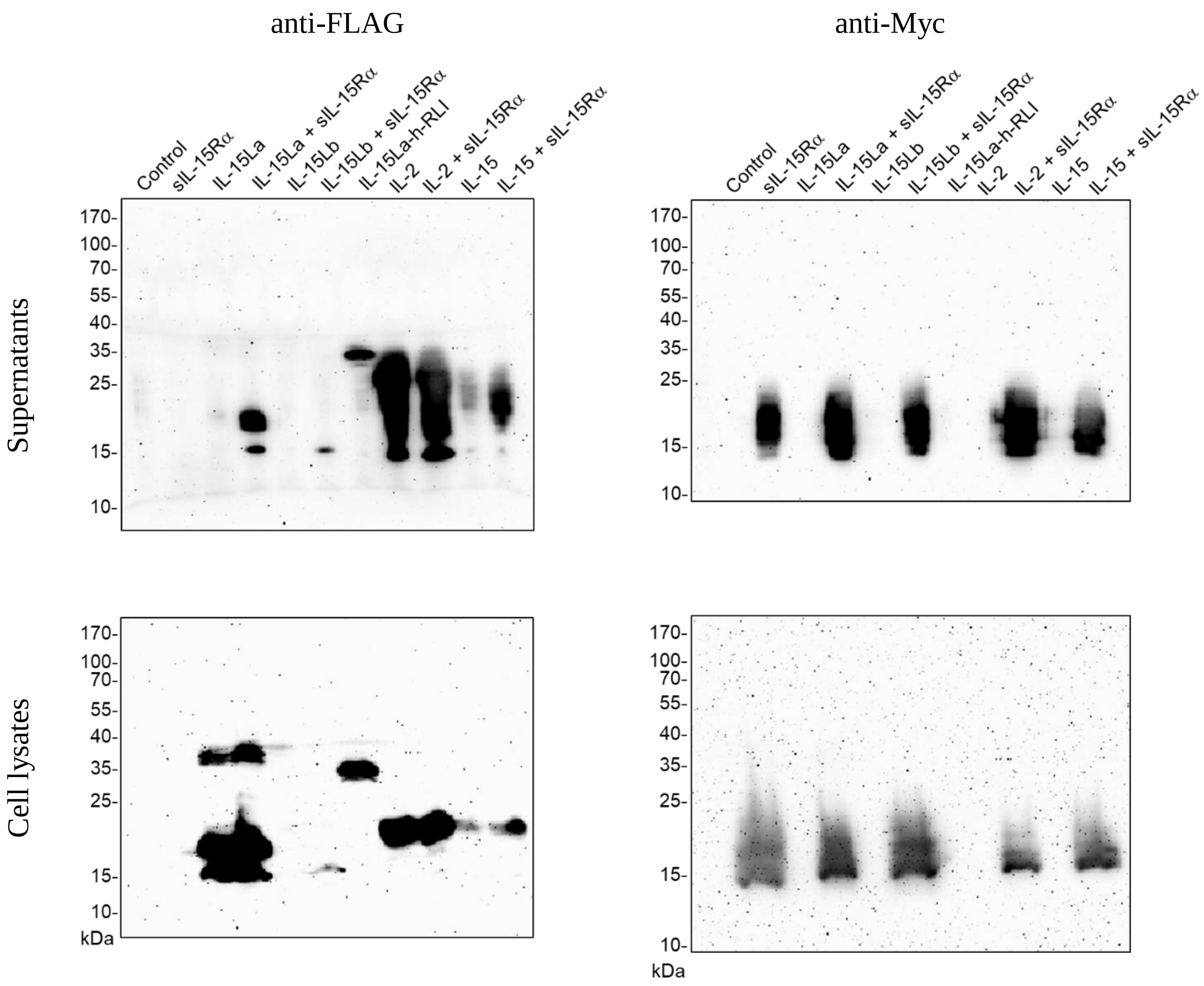

**Fig. S5B-5.** Trout, experiment 3.

*Western blots*

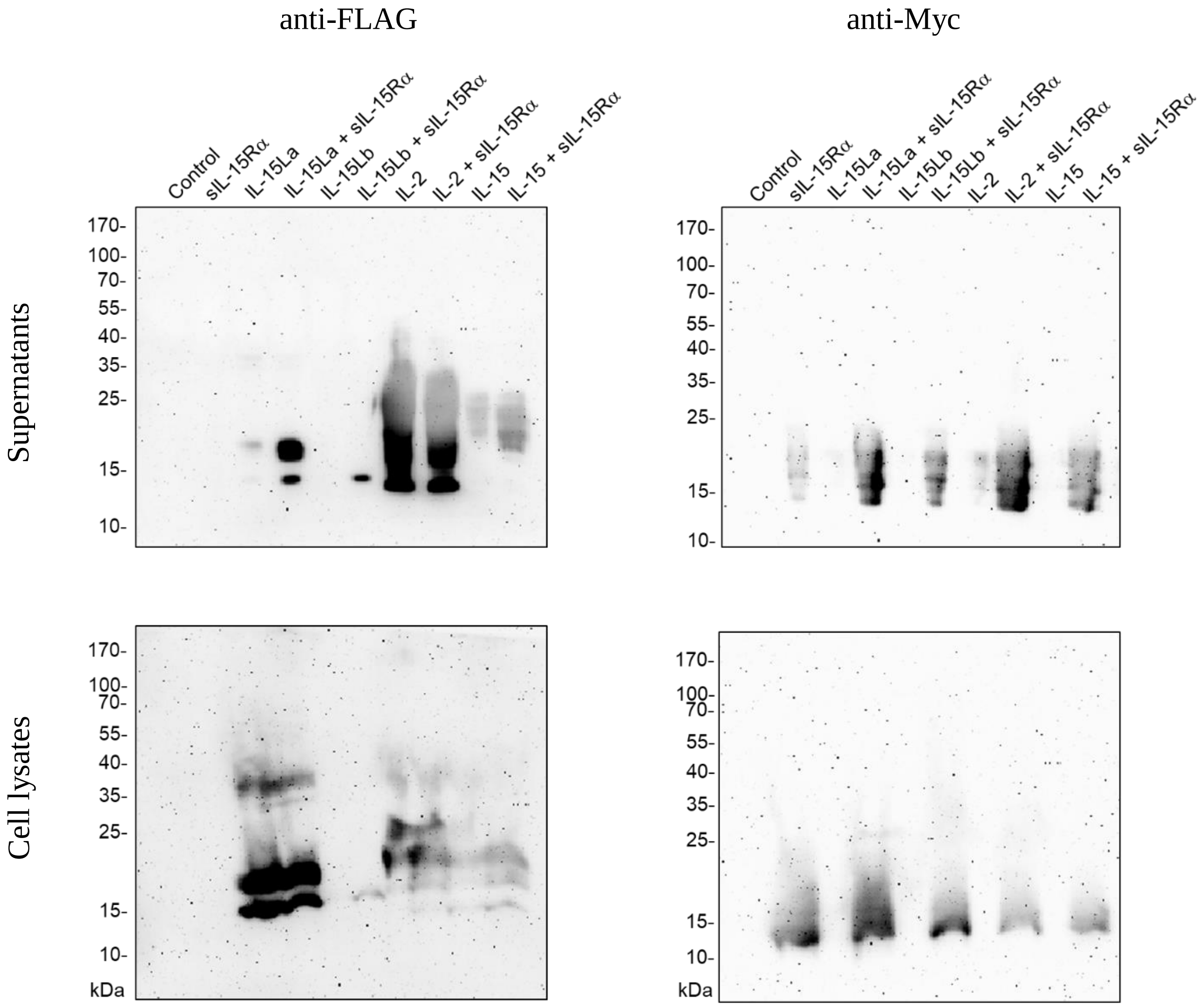

**Fig. S5C.** Phosphorylation of STAT5 in various trout lymphocyte populations induced after incubation with recombinant trout cytokine containing HEK293T supernatants *(information supporting the main text Fig. 7 results)*.

Western blot analyses of phosphorylated STAT5 (pSTAT5) in CD8$\alpha^+$ and CD8$\alpha^-$ fractions of trout morphological lymphocytes that had been isolated from several tissues using flow sorting (for examples see Fig. S3) and had been incubated with recombinant trout cytokine containing supernatants from transfected HEK293T cells.

The results shown in this figure show uncropped versions of the Western blot results shown in main text Fig. 7, as well as confirmation of those results by independent experiments. In addition, in many of the experiments the HEK293T supernatants used for stimulation were investigated for the presence of recombinant cytokines by anti-FLAG Western blot analysis.

Lymphocytes were incubated for 15 min at 15 °C with the supernatants of HEK293T cells transfected for trout IL-2, IL-15 or IL-15La, with (+) or without (-) trout sIL-15Rα. In most cases FLAG-tagged cytokines were used, but in some instances also cytokines without an added tag were used [IL-2(N), IL-15 (N), IL-15La(N) and IL-15Lb(N)]; furthermore, in some experiments an RLI fusion product was used which consisted of a fusion between human sIL-15Rα and trout IL-15La (IL-15La-h-RLI; for explanation see the Fig. S5B legend). Treatment controls consisted of supernatants of HEK293T cells transfected with empty vector (Control), or with vector for trout sIL-15Rα alone (sIL-15Rα). Independent experiments for lymphocytes of the same tissue pooled from other trout individuals are numbered #1 etc. Results are shown for: Intestine#1 (Fig. S5C-1), Intestine#2 (Fig. S5C-2), Gill#1 (Fig. S5C-3), Gill#2 (Fig. S5C-4), Spleen#1 (Fig. S5C-5), Spleen#2 (Fig. S5C-6), Head kidney#1 (Fig. S5C-7), Head kidney#2 (Fig. S5C-8), Thymus#1 (Fig. S5C-9), Thymus#2 (Fig. S5C-10) and Thymus#3 (Fig. S5C-11). (a) After stimulation, lysates were subjected to Western blot analysis using antibodies against phosphorylated STAT5. After stripping and washing, the same blots were analyzed by Western blot analysis using anti-actin as a loading control. (b) In many but not all instances the (unconcentrated) HEK293T supernatants used for stimulation were subjected to anti-FLAG Western blot analysis; because the proteins were not concentrated first, the cytokine concentrations were often too low for allowing detection. However, a relevant finding was that in the investigated supernatants the IL-2 concentrations were higher than those of IL-15, concluding that in cases where IL-15 showed higher activity than IL-2 that could not be due to lower concentrations of IL-2. In some instances, more than one pooled tissue was analyzed for the same group of trout individuals using the same HEK293T supernatants (Spleen#1 and Gill#2; Intestine#1 and Head kidney#1; Gill#1 and Thymus#2); in such cases, for convenience of the reader, the Western blots of the HEK293T supernatants are repeatedly shown in the relevant figures. The experiments of which cropped blot results are shown in main text Fig. 7 are: Intestine#1, Gill#1, Spleen#1, Head kidney#1, and Thymus#1.

As exceptions, for the Spleen#2 and Thymus#3 experiments (Figs. S5C-6 and -11) cells were not only separated by using anti-CD8 but also by anti-CD4. For the Thymus#3 experiment only the stimulation of CD4$^-$CD8$^-$ (DN) thymocytes was investigated.

**Fig. S5C-1** Intestine#1

*Western blots*

(a) *Results of the lymphocyte stimulation experiments*

CD8⁺ lymphocytes CD8⁻ lymphocytes

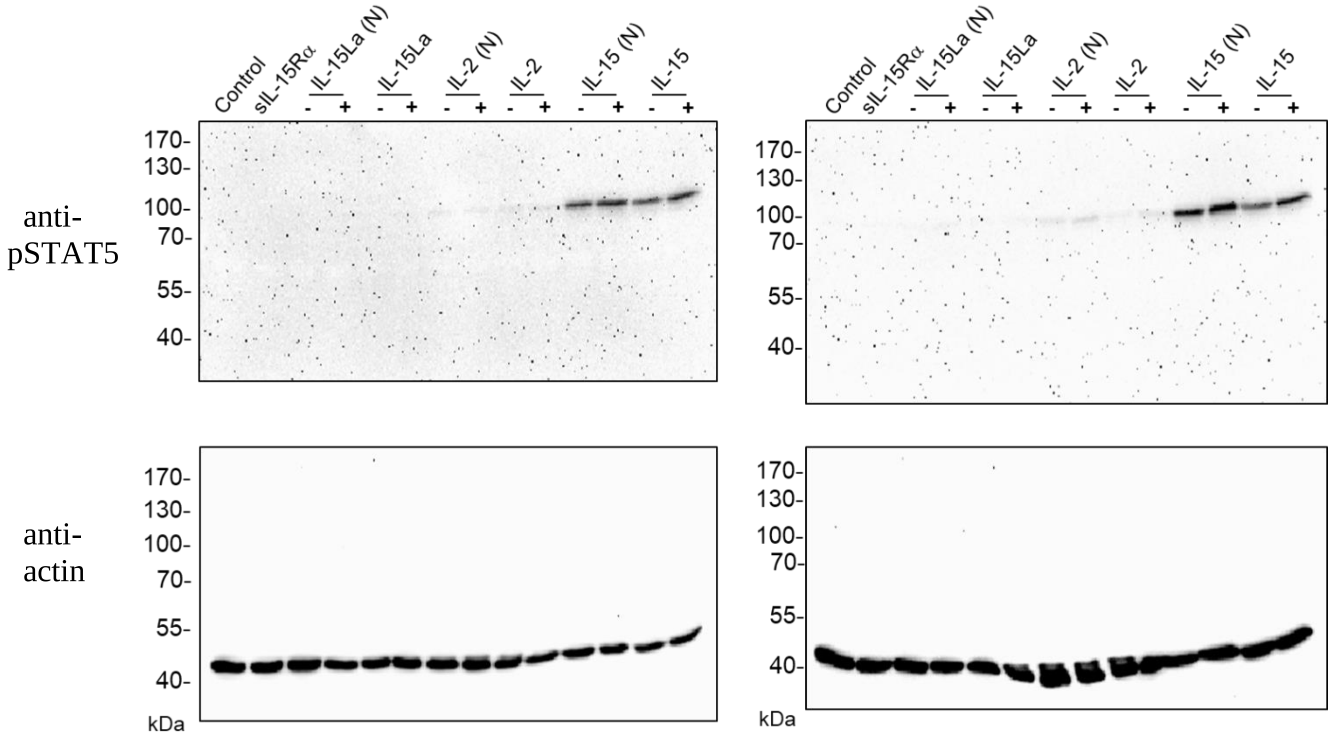


(b) *anti-FLAG Western blot of the HEK293T supernatants used for the stimulation*

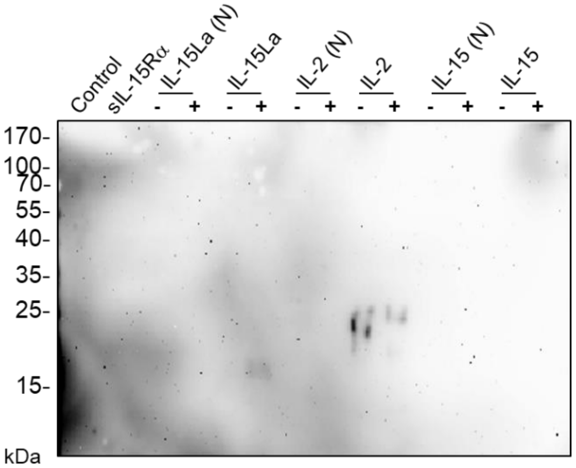

**Fig. S5C-2** Intestine#2

*Western blots*

(a) *Results of the lymphocyte stimulation experiments*

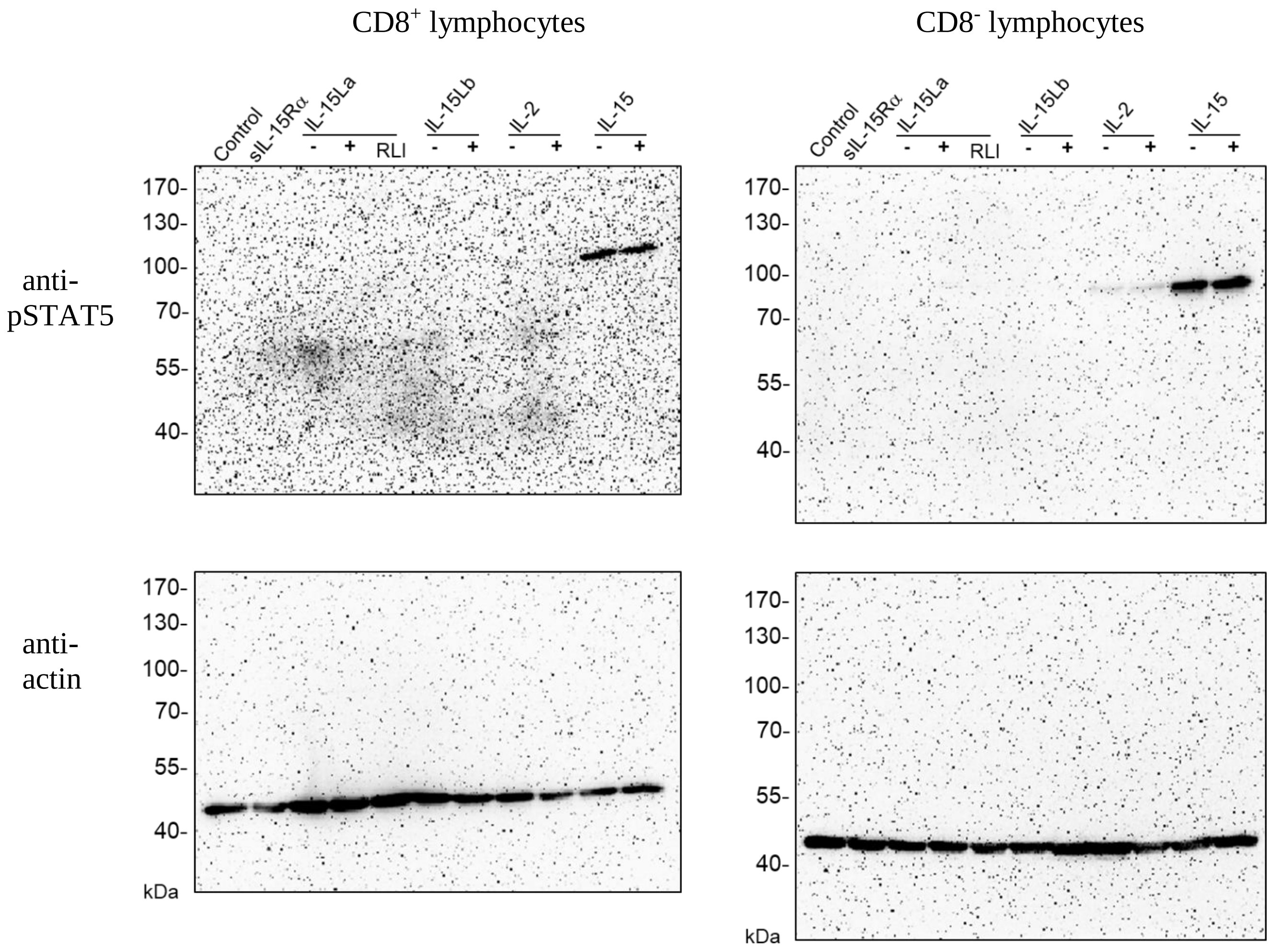

**Fig. S5C-3** Gill#1

*Western blots*

(a) *Results of the lymphocyte stimulation experiments*

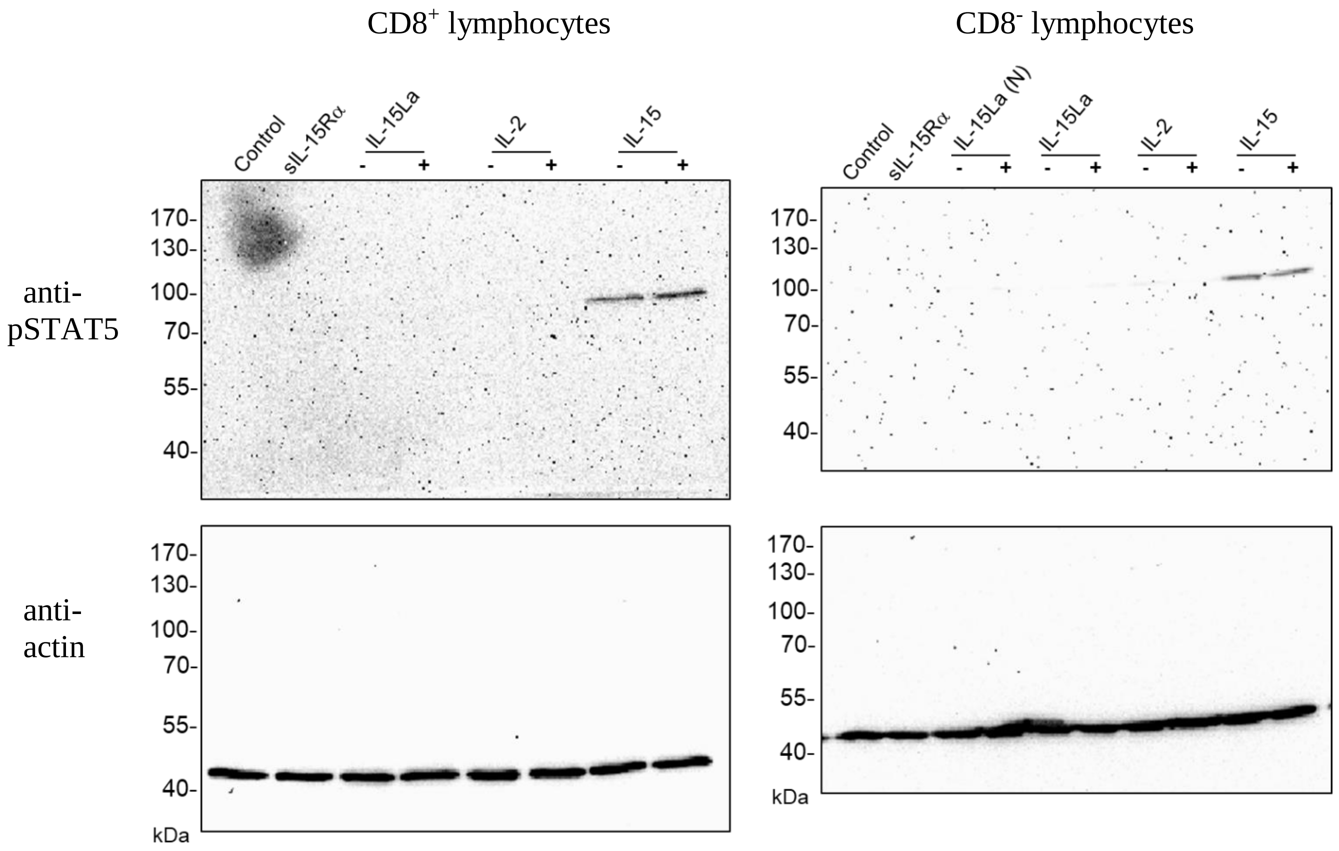


(b) *anti-FLAG Western blot of the HEK293T supernatants used for the stimulation*

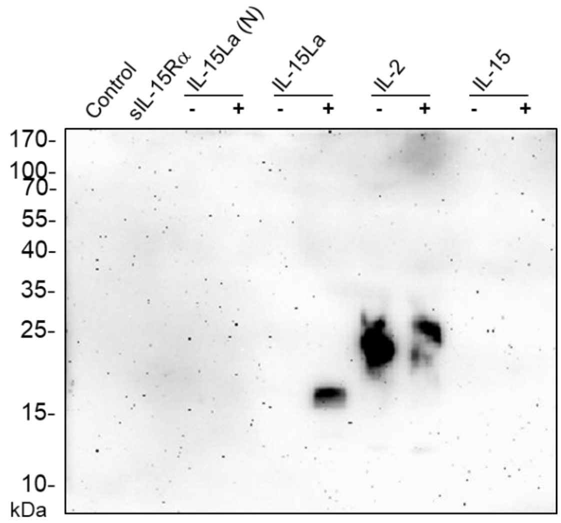

**Fig. S5C-4** Gill#2

*Western blots*

(a) *Results of the lymphocyte stimulation experiments*

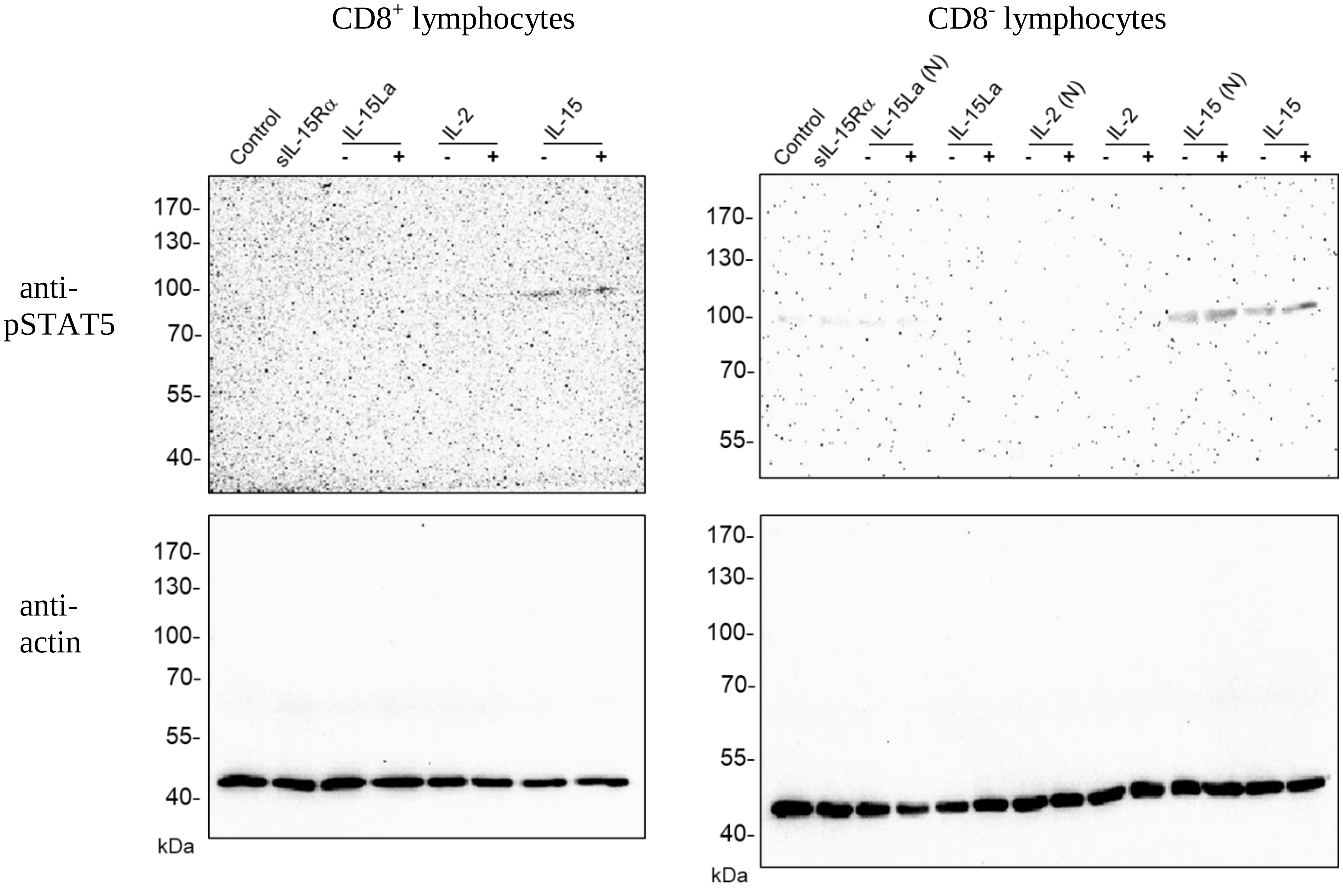


(b) *anti-FLAG Western blot of the HEK293T supernatants used for the stimulation*

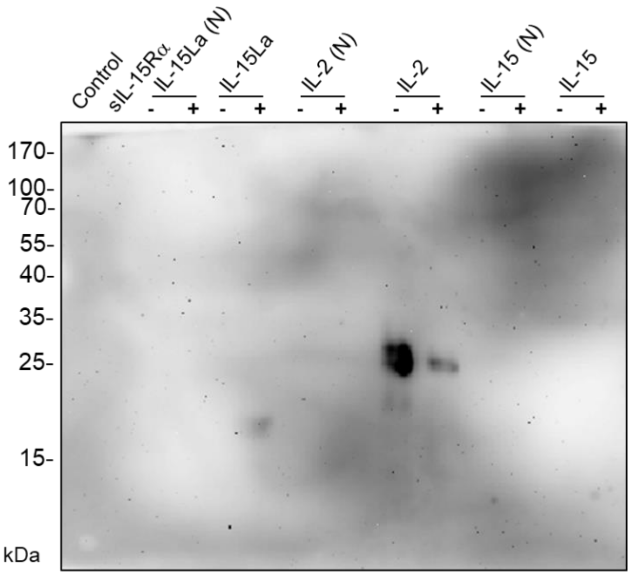

#### **Fig. S5C-5** Spleen#1

*Western blots*

(a) *Results of the lymphocyte stimulation experiments*

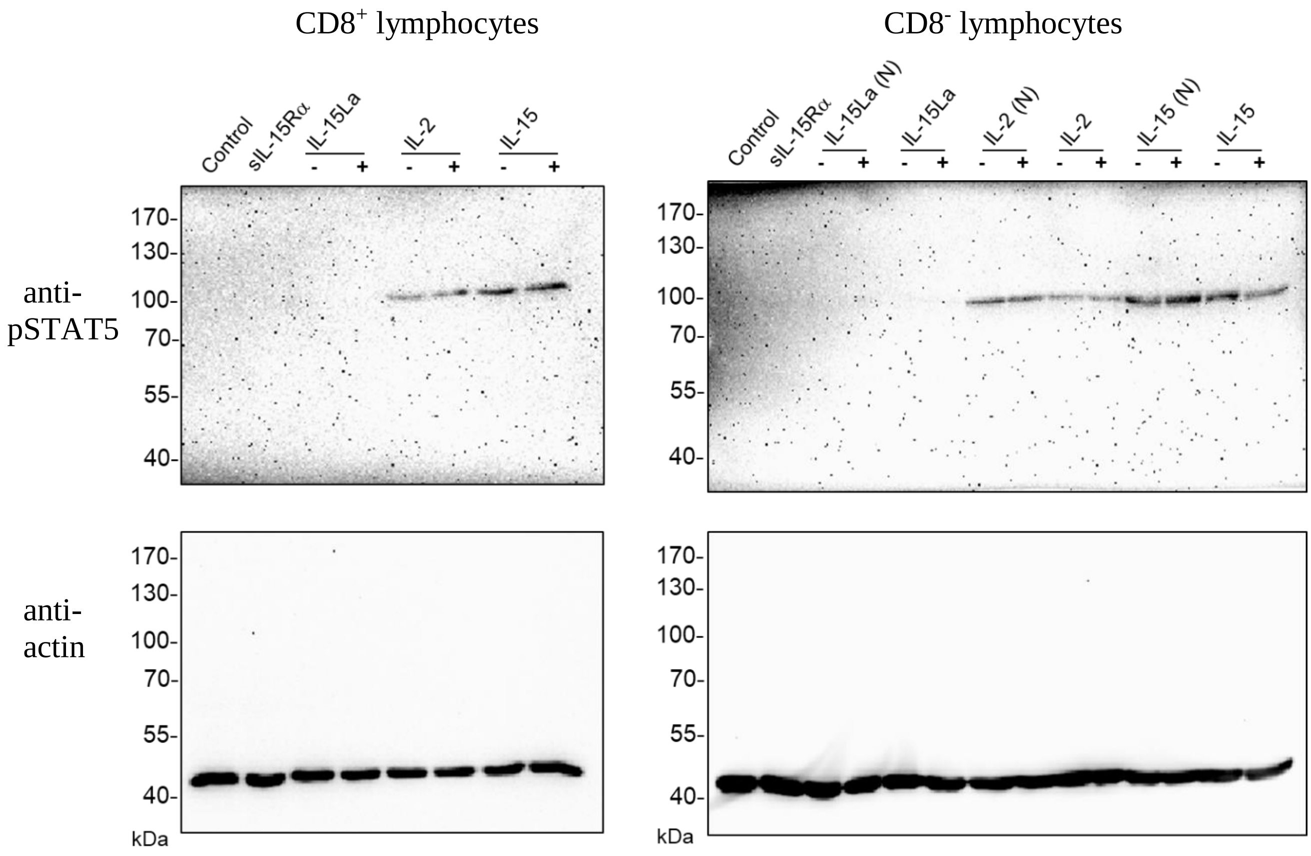


(b) *anti-FLAG Western blot of the HEK293T supernatants used for the stimulation*

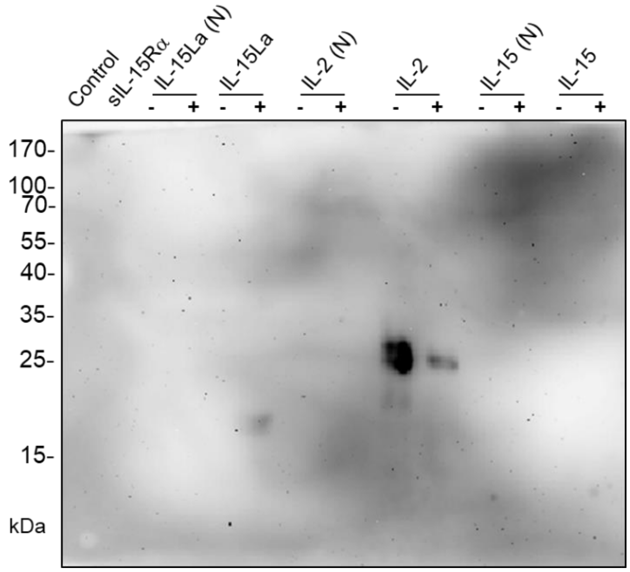

**Fig. S5C-6** Spleen#2

*Western blots*

(a) *Results of the lymphocyte stimulation experiments*

CD8$^{+}$ lymphocytes

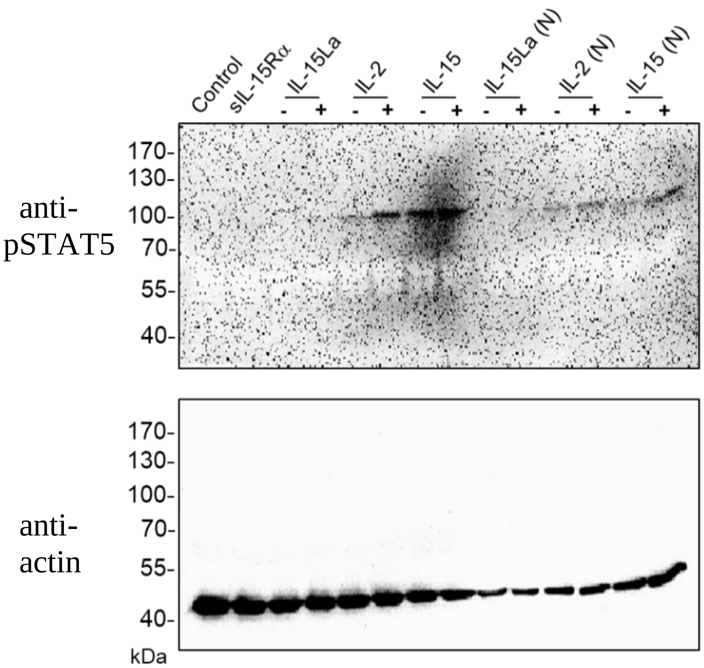


CD8$^{-}$ lymphocytes

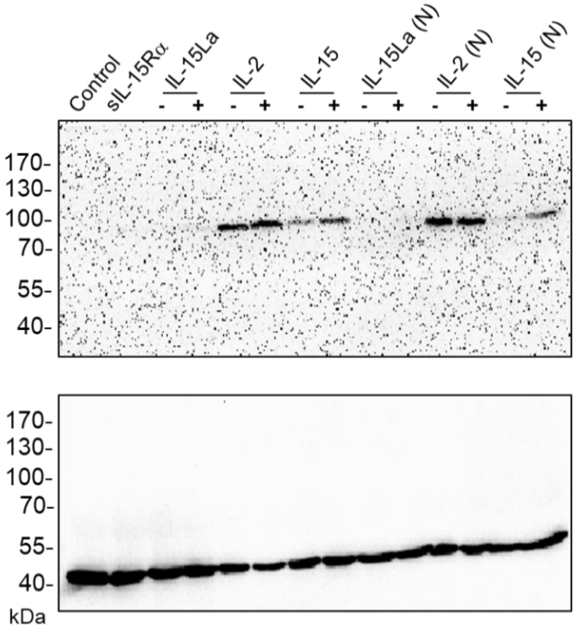


CD4$^{-}$CD8$^{-}$(DN) lymphocytes

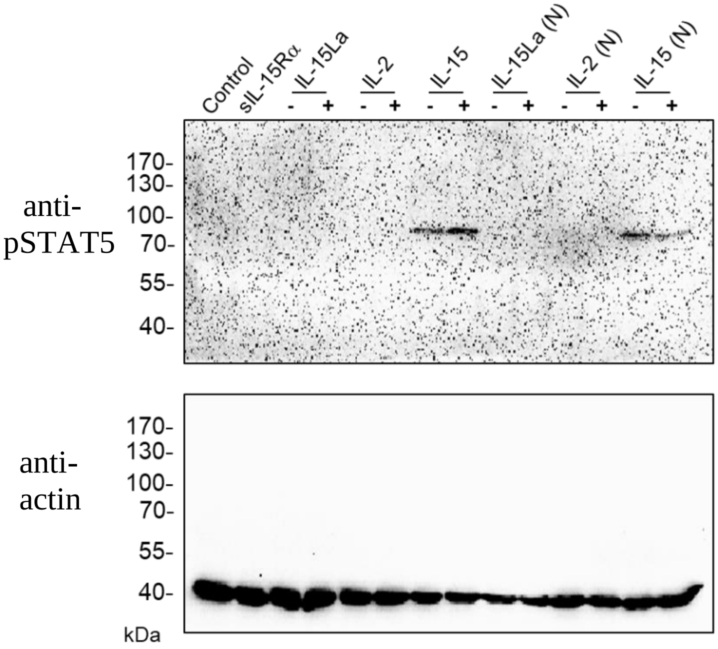


(b) *anti-FLAG Western blot of the HEK293T supernatants used for the stimulation*

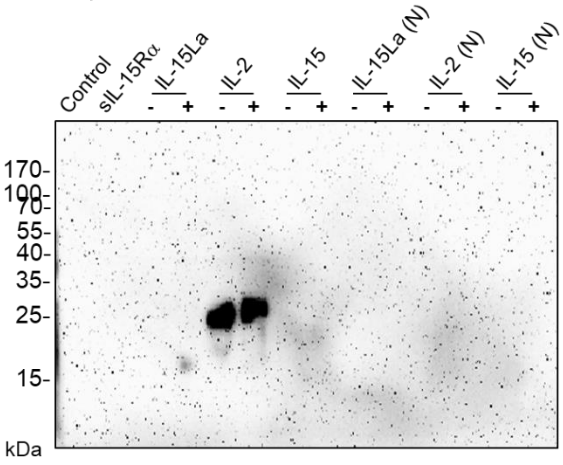

**Fig. S5C-7** Head kidney#1

*Western blots*

(a) *Results of the lymphocyte stimulation experiments*

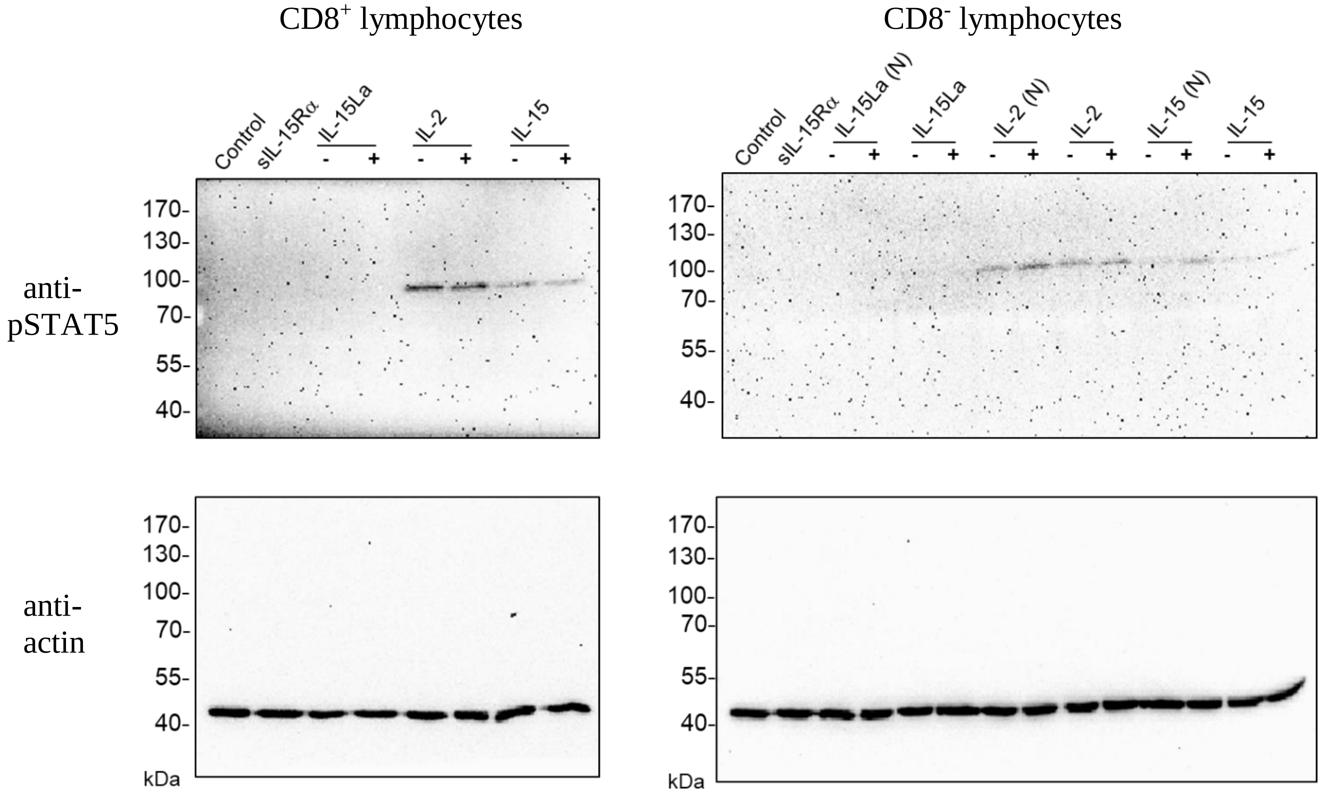


(b) *anti-FLAG Western blot of the HEK293T supernatants used for the stimulation*

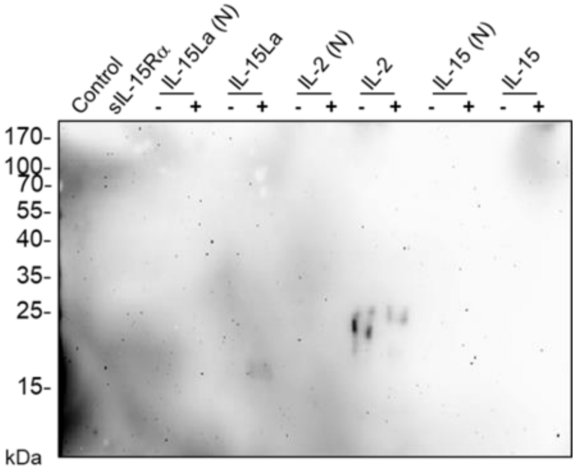

#### **Fig. S5C-8** Head kidney#2

*Western blots*

(a) *Results of the lymphocyte stimulation experiments*

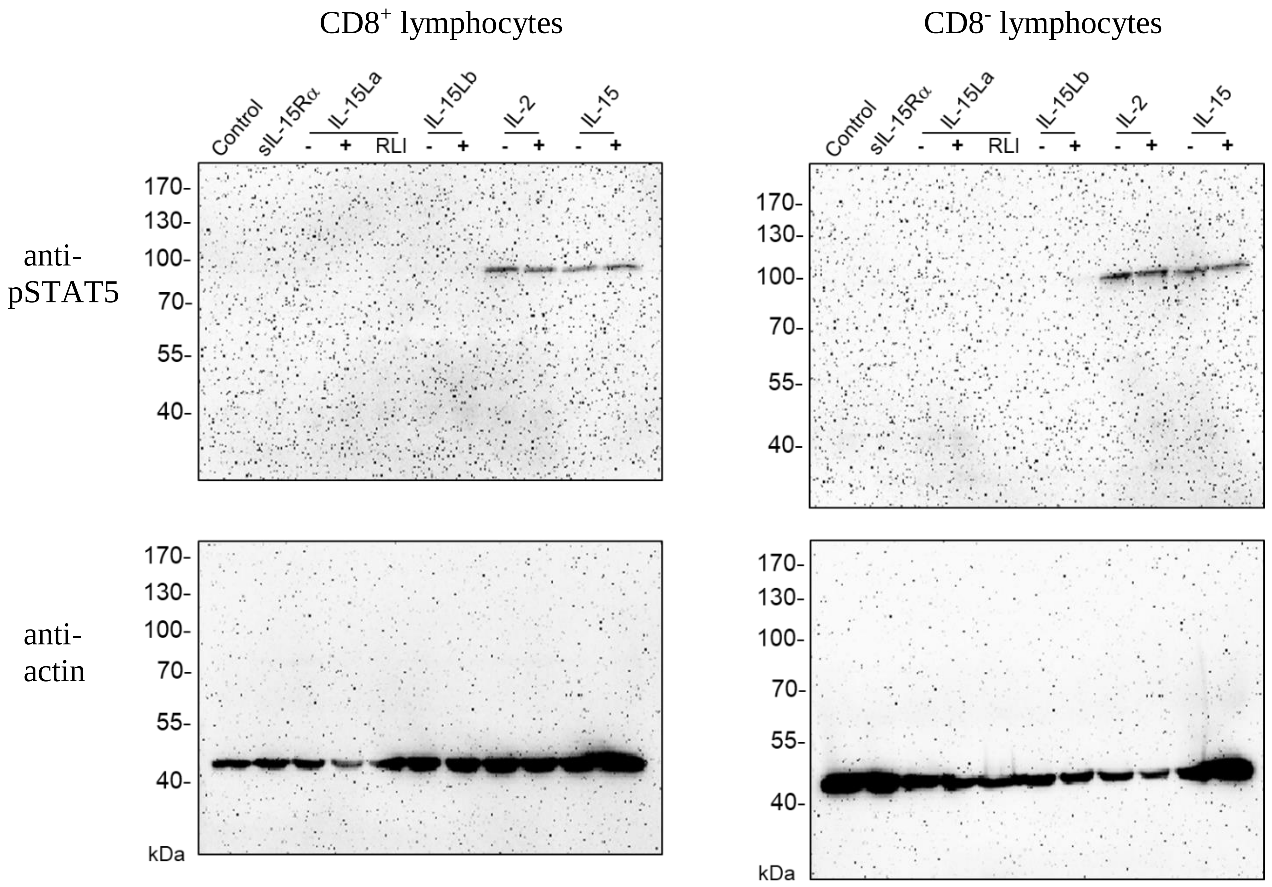


(b) *anti-FLAG Western blot of the HEK293T supernatants used for the stimulation*

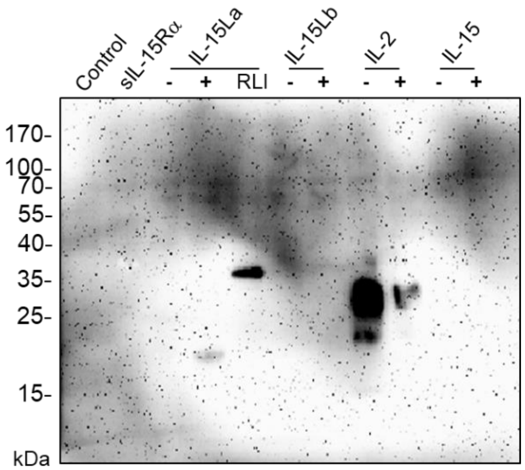

**Fig. S5C-9** Thymus#1

*Western blots*

(a) *Results of the lymphocyte stimulation experiments*

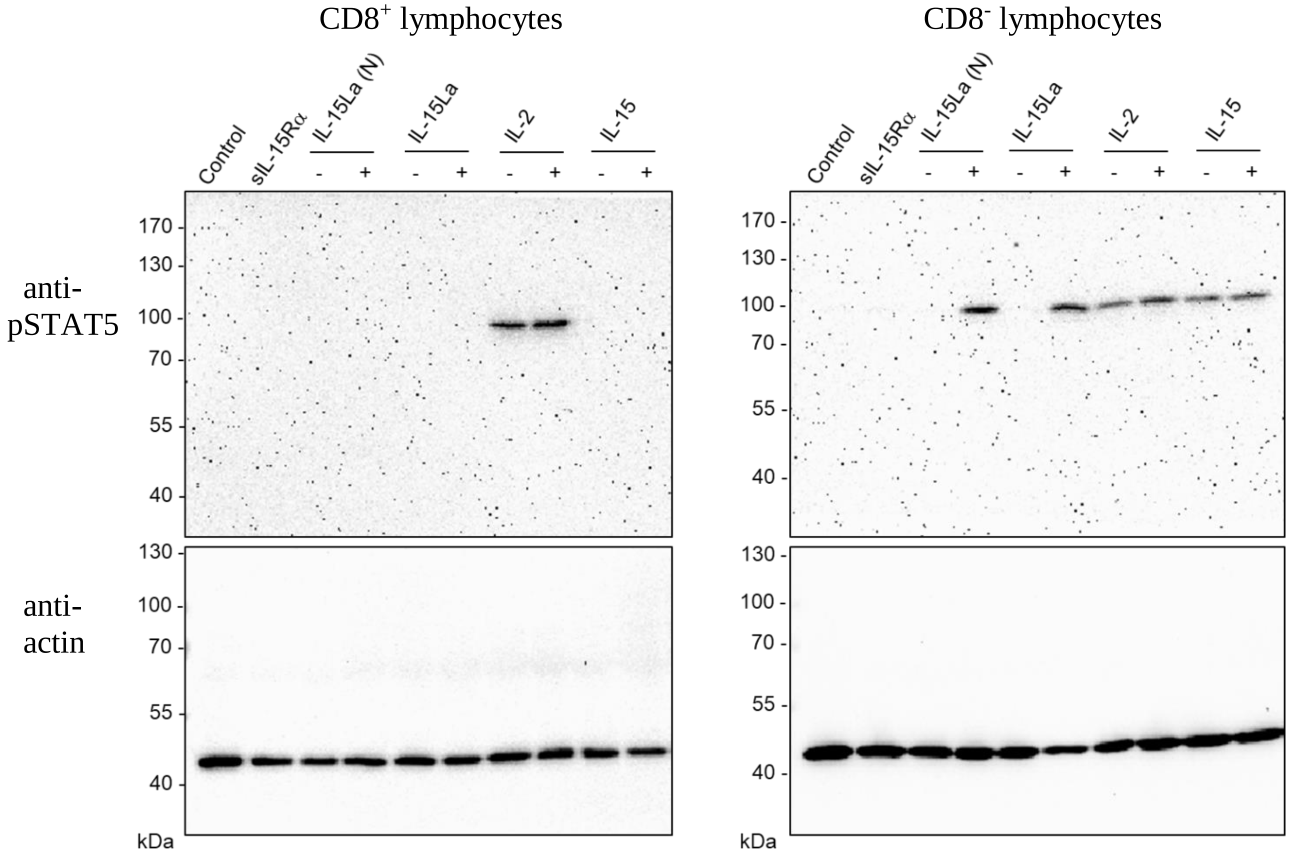


(b) *anti-FLAG Western blot of the HEK293T supernatants used for the stimulation*

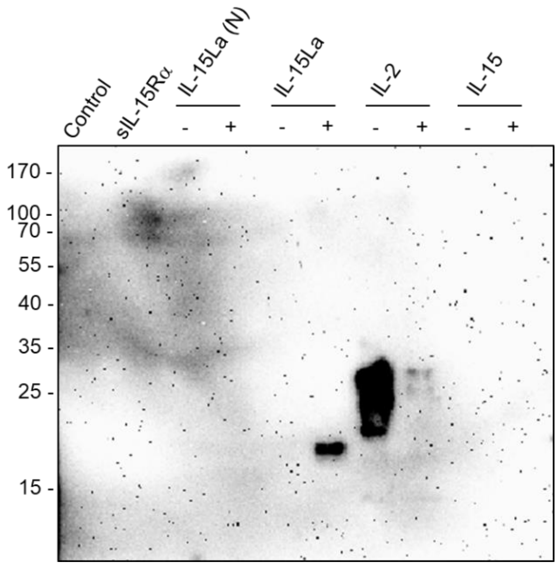

**Fig. S5C-10** Thymus#2

*Western blots*

(a) *Results of the lymphocyte stimulation experiments*

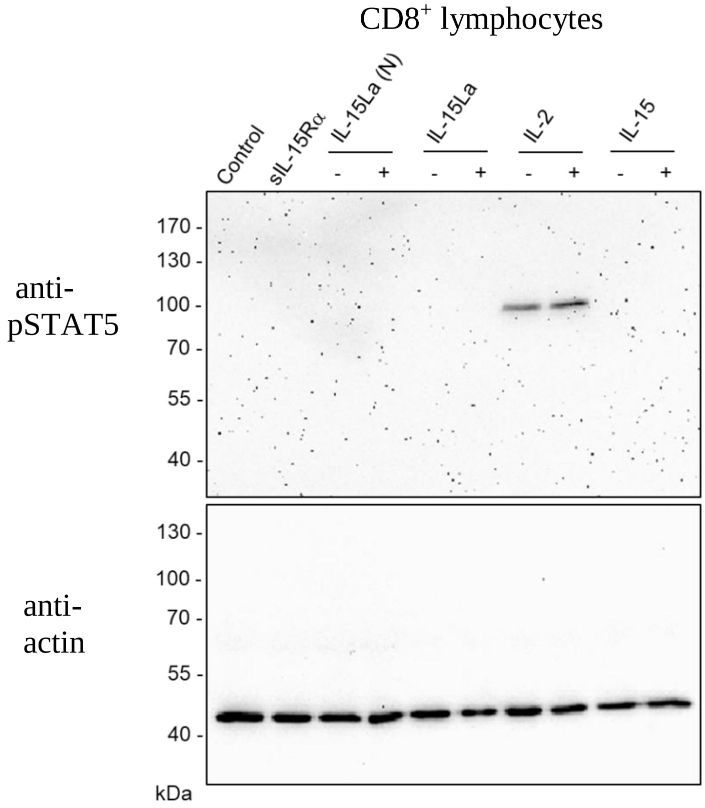


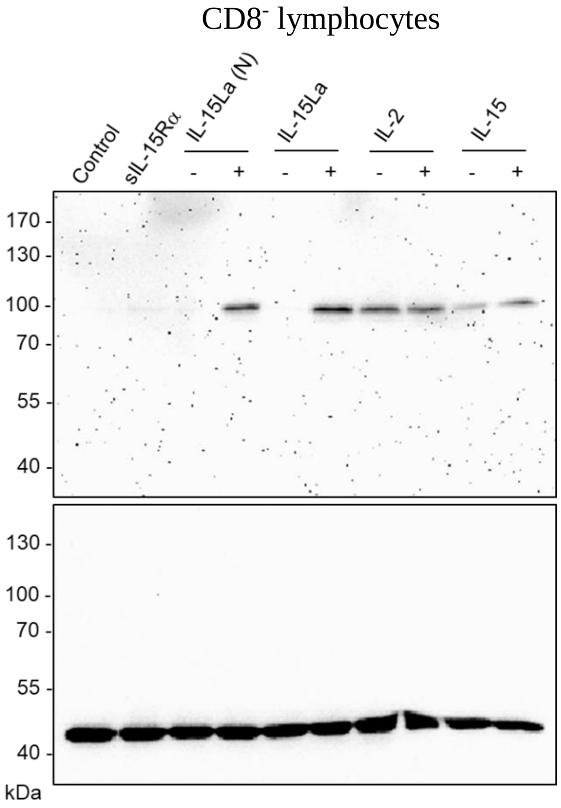


(b) *anti-FLAG Western blot of the HEK293T supernatants used for the stimulation*

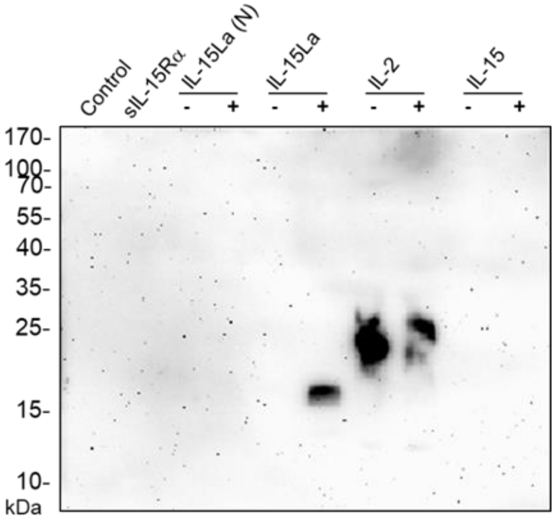

**Fig. S5C-11** Thymus#3

*Western blots*

(a) *Results of the lymphocyte stimulation experiments*

CD4⁻CD8⁻ (DN) lymphocytes

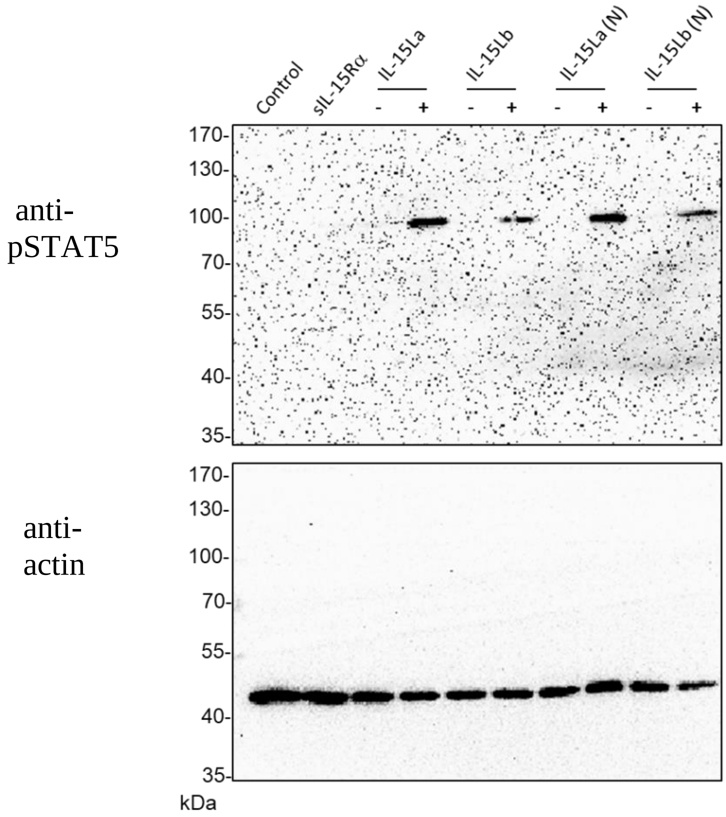

**Fig. S5D.** Phosphorylation of STAT5 in DN, DP, CD4SP and CD8SP fractions of trout thymocytes induced after incubation with recombinant cytokine containing HEK293T supernatants (*information supporting the main text Fig. 8 results*).

Western blot analyses of phosphorylated STAT5 (pSTAT5) in $CD8^-CD4^-$ (DN), $CD8^+CD4^+$ (DP), $CD8^+CD4^-$ (CD8SP), and $CD8^-CD4^+$ (CD4SP) fractions of morphological lymphocytes that had been isolated from trout thymus using flow sorting (for an example see Fig. S3) and had been incubated with recombinant trout cytokine containing supernatants from transfected HEK293T cells.

This figure supports the main text Fig. 8. Fig. S5D-1 is identical to main text Fig. 8, except that now also the loading control results are shown. Fig. S5D-2 is an independent confirmation of the main text Fig. 8 result.

Trout thymocytes were separated using appropriate antibodies and flow sorting into $CD8^-CD4^-$ (DN), $CD8^+CD4^+$ (DP), $CD8^+CD4^-$ (CD8SP), and $CD8^-CD4^+$ (CD4SP) populations. Isolated populations were incubated for 15 min at 15 °C with the supernatants of HEK293T cells transfected for trout IL-2, IL-15 or IL-15La, with or without trout sIL-15Rα. In a few cases, supernatants of HEK293T cells transfected for RLI fusion products including human sIL-15Rα were used (see explanation in Fig. S5B legend). Experiments 1 and 2 were performed independently from each other, using thymocytes pooled from different trout individuals and HEK293T supernatants from different transfections.

Fig. S5D-1 (Experiment 1) is identical to main text Fig. 8, except that now also the anti-actin loading control blot results are shown. "-" and "+" refer to whether sIL-15Rα was co-expressed with the indicated cytokines. The RLI form used in this experiment was a fusion of human sIL-15Rα and trout IL-15La (IL-15La-h-RLI).

Fig. S5D-2 (Experiment 2) (a) In addition to the cytokines used in Experiment 1, also a fusion between human sIL-15Rα and modified bovine IL-15L was tested (bov.IL-15Lhyb-h-RLI). This incubation of the modified bovine molecule with trout lymphocytes was just one of our many different trials to find a function for bovine IL-15L, none of which produced positive results so far (this negative lane result in Fig. S5D-2 is not relevant for the present study, but was left to to show the intact blot result). (b) Different from Experiment 1, in Experiment 2 the unconcentrated HEK293T supernatants were subjected to anti-FLAG Western blot analysis. As discussed in the Fig. S5C legend, not for all cytokines the concentrations were sufficiently high for being detectable in this manner, but the blot result helps to see for example that the IL-2 concentrations used were higher than the IL-15 concentrations.

**Fig. S5D-1.** Experiment 1.

Main text Fig. 8 but now also showing the anti-actin loading control results.

*Western blots*

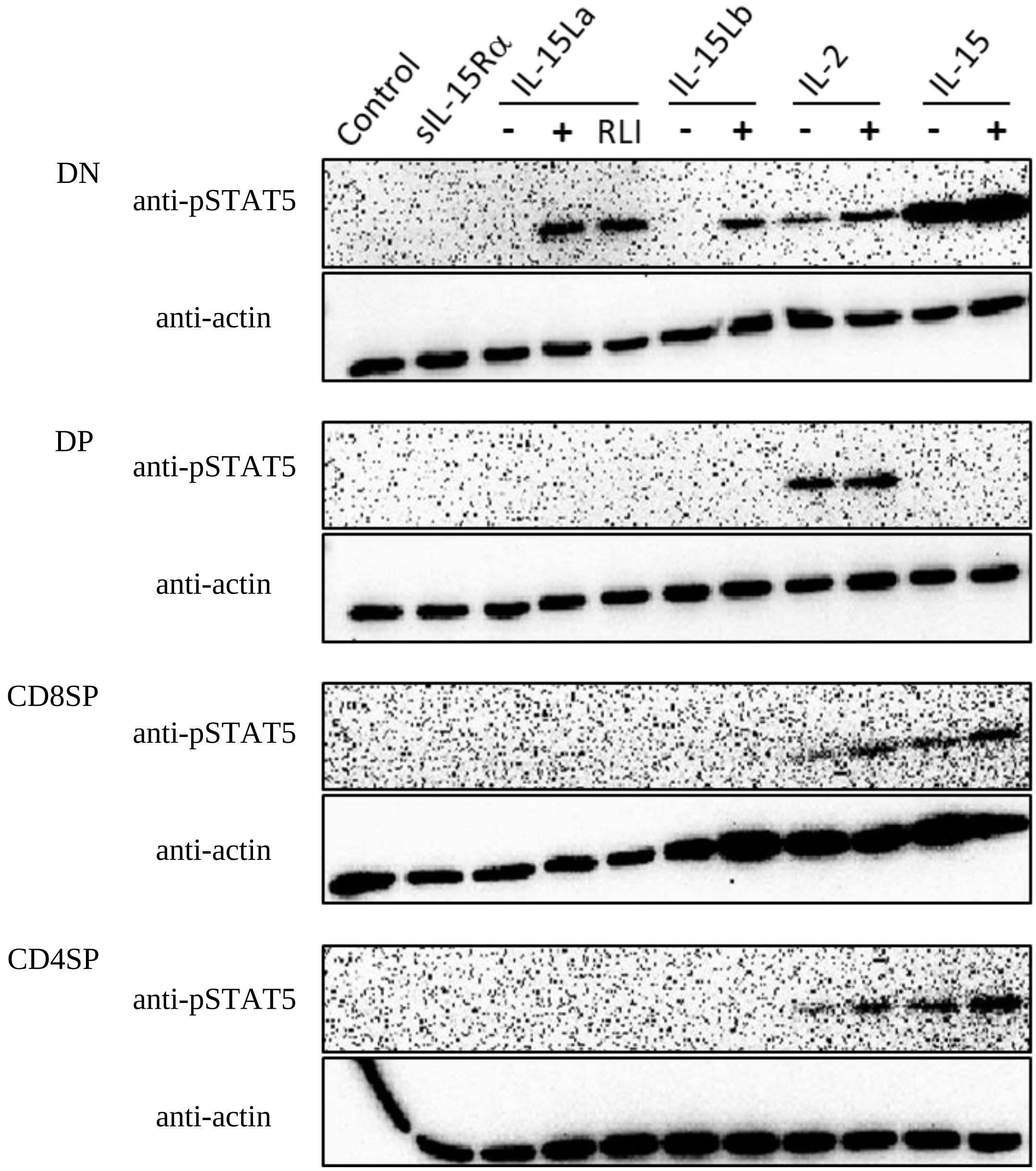

**Fig. S5D-2.** Experiment 2.

Independent confirmation of the main text Fig. 8 result. *Western blots*

(a) *Results of the lymphocyte stimulation experiments. Upper blots show anti-pSTAT5 results, lower blots show the anti-actin loading control results*

M, marker; 1, Vector control; 2, trout sIL-15Ra; 3, IL-15La-h-RLI (trout IL-15La linked with human sIL-15Rα); 4, bov.IL-15Lhyb-h-RLI (modified bovine IL-15L linked with human sIL-15Rα); 5, trout IL-15La; 6, trout IL-15La+sIL-15Rα; 7, trout IL-15Lb; 8, trout IL-15Lb+sIL-15Rα; 9, trout IL-2; 10, trout IL-2+sIL-15Rα; 11, trout IL-15; 12, trout IL-15+sIL-15Rα.

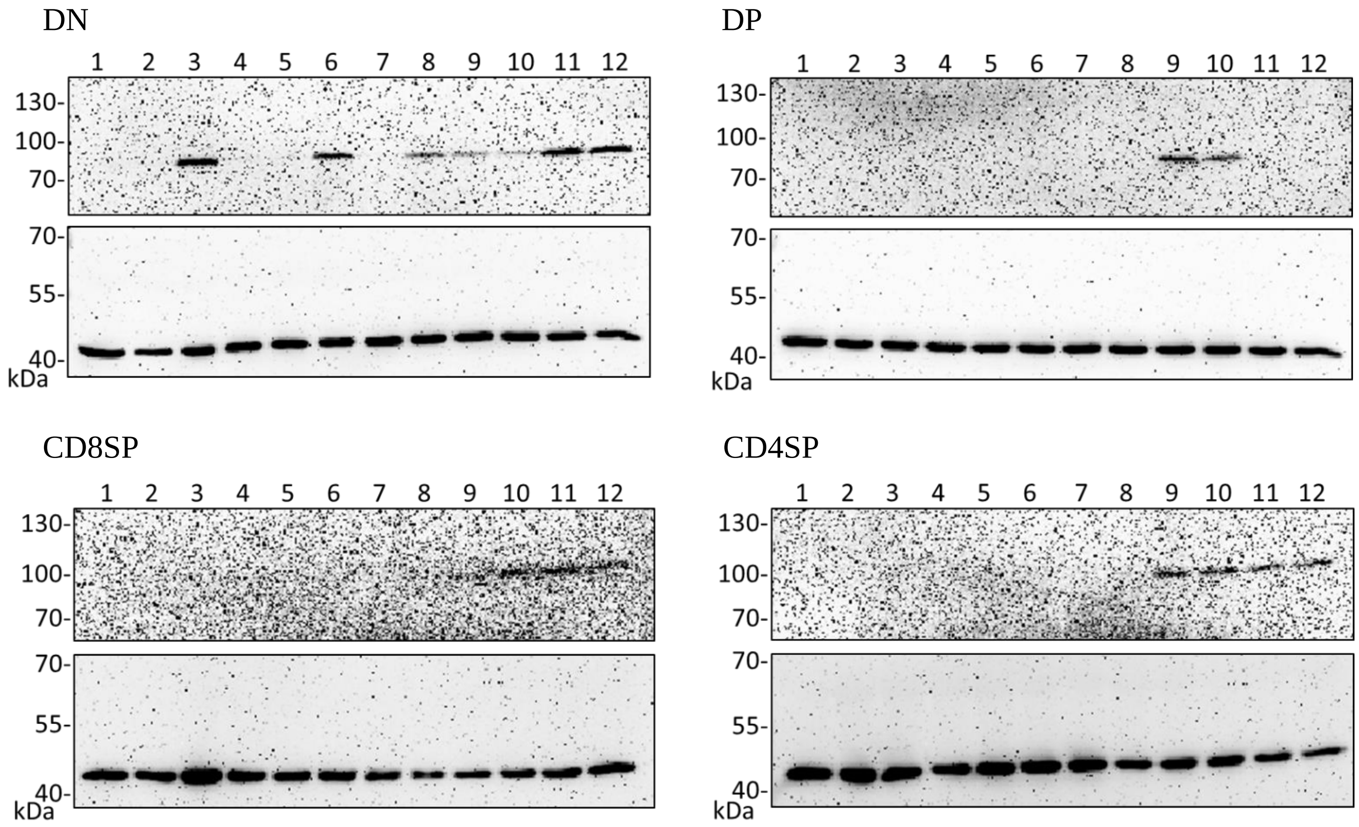


(b) *anti-FLAG Western blot of the HEK293T supernatants used for the stimulation*

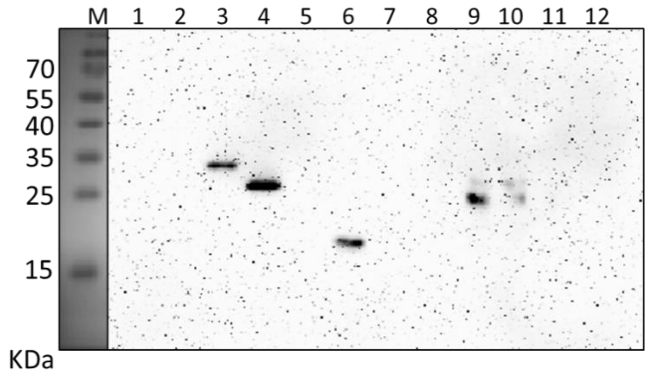

**Fig. S5E.** Phosphorylation of STAT5 in lymphocyte fractions of trout thymus, intestine and spleen, induced after incubation with purified recombinant trout cytokines (produced in insect cells) IL-2, IL-15-RLI and IL-15La-RLI at 5, 25 and 125 nM (*information supporting the main text Fig. 9 results*).

Western blot analyses of phosphorylated STAT5 (pSTAT5) in $CD8^{-}CD4^{-}$ (DN), $CD8^{+}CD4^{+}$ (DP; only isolated from thymocytes), $CD8^{+}CD4^{-}$ (CD8SP), and $CD8^{-}CD4^{+}$ (CD4SP) fractions of morphological lymphocytes that had been isolated from trout thymus, intestine and spleen using flow sorting (for examples see Fig. S3) and had been incubated for 15 min at 15 °C with recombinant trout cytokines purified from insect cells (for protein purification see Fig. S4). These cytokines were trout IL-2, IL-15-RLI (a fusion of trout IL-15 and trout sIL-15Rα) and IL-15La-RLI (a fusion of trout IL-15La and trout sIL-15Rα), and they were used in 5, 25 and 125 nM concentrations. As negative controls, cells were mock treated (Control). After stripping and washing, the same blots were analyzed by Western blot analysis using anti-actin as a loading control.

This figure supports the main text Fig. 9. Fig. S5E-1 is identical to main text Fig. 9, except that now also the anti-actin loading control results are shown. Fig. S5E-2 is based on an independent but similar experiment as done for Fig. S5E-1, although only done for thymocytes. The Fig. S5E-2 result confirms many of the observations in Fig. S5E-1 (main text Fig. 9) for thymocytes, for example that DN thymocytes are sensitive to IL-15La-RLI and that DP thymocytes are only sensitive to IL-2. However, comparison of the Fig. S5E-1 and Fig. S5E-2 results for thymocytes also shows that preparations pooled from different trout individuals can have different relative sensitivities to the different cytokines, as for example the DN thymocyte preparation used for the Fig. S5E-2 experiment was more sensitive to IL-15La-RLI than to IL-2 or IL-15-RLI. In the future, when hopefully the respective cytokine receptors will have been determined and antibodies will have been established against those receptors, the different sensitivities for cytokines between cells of different trout individuals may be addressed in more detail.

**Fig. S5E-1.** Main text Fig. 9 with anti-actin loading control.

*Western blots*

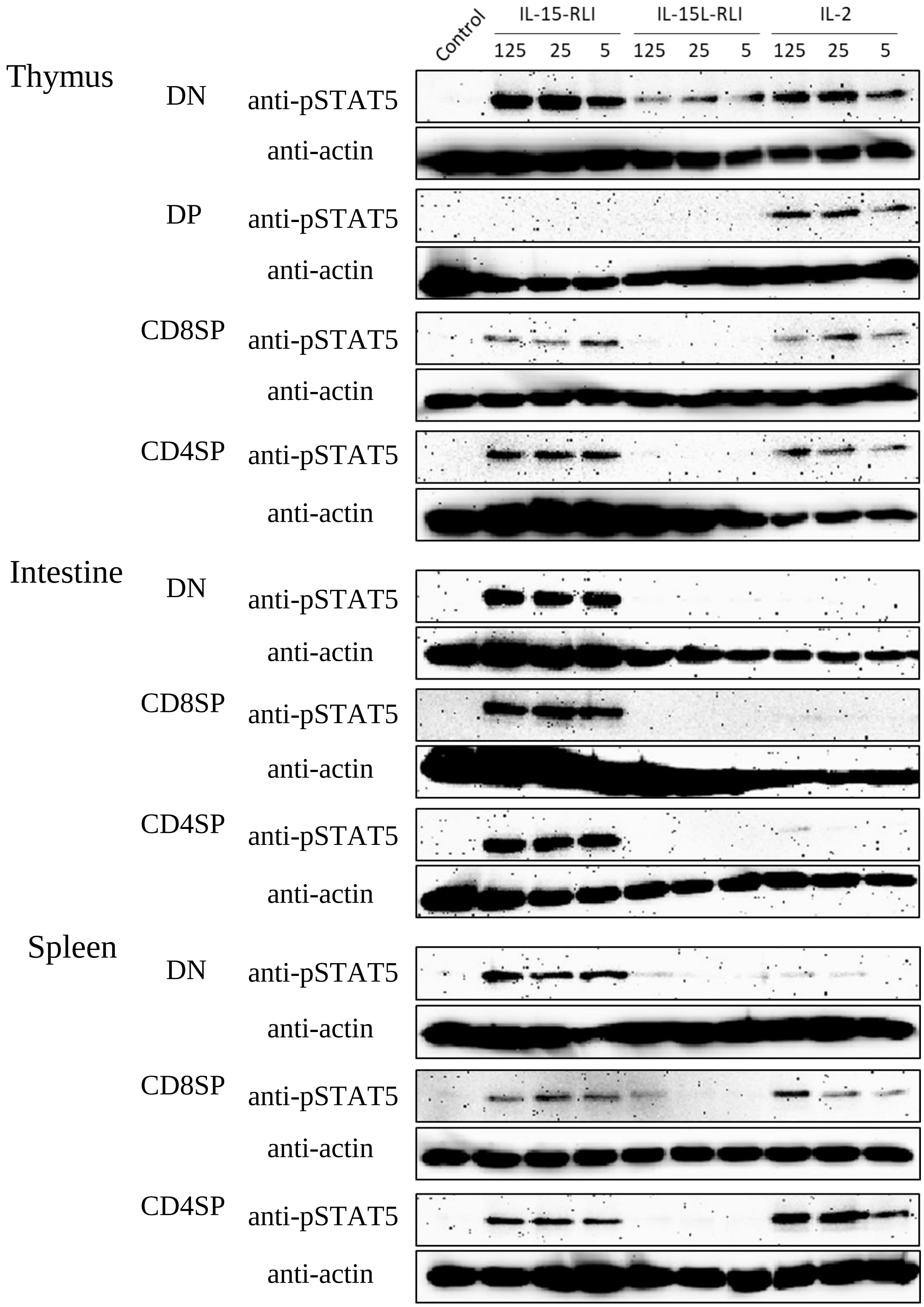

**Fig. S5E-2.** Stimulation of thymocytes pooled from different trout individuals than used for the Fig. S5E-1 experiment.

*Western blots*

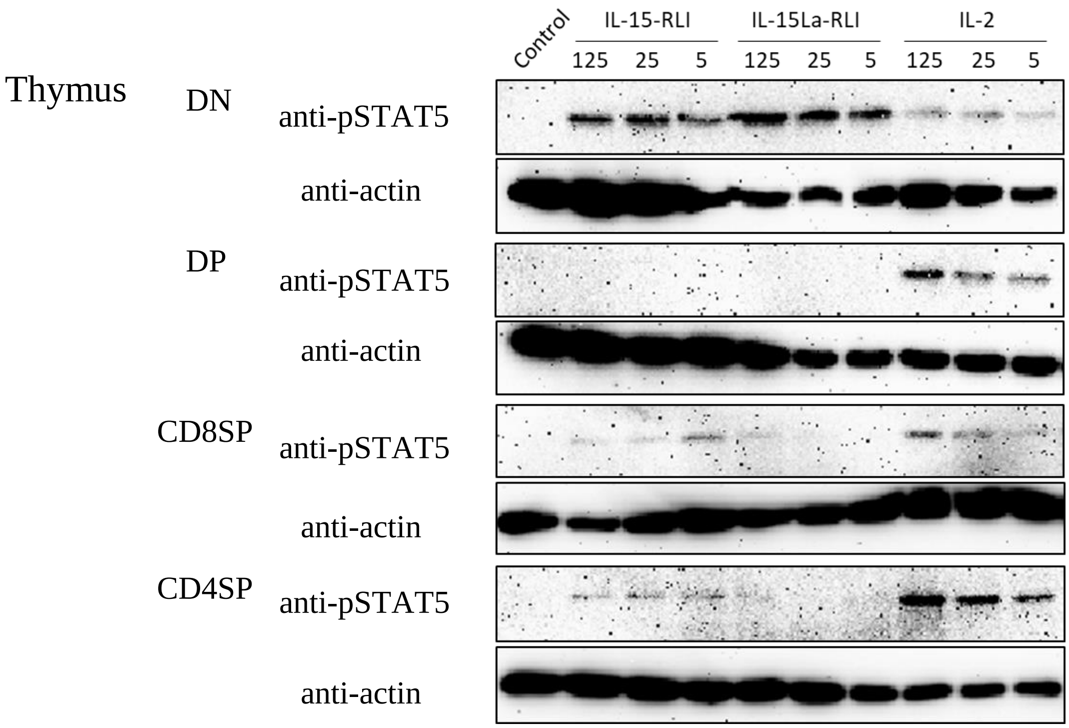

**Fig. S5F.** Titration of the recombinant trout cytokine (produced in insect cells) concentration necessary for the induction of pSTAT5.

Western blot analyses of phosphorylated STAT5 (pSTAT5) in sensitive trout lymphocyte populations that had been isolated using flow sorting (for examples see Fig. S3) and had been incubated for 15 min at 15 °C with recombinant trout cytokines purified from insect cells (for protein purification see Fig. S4). These cytokines were trout IL-2, IL-15-RLI (a fusion of trout IL-15 and trout sIL-15Rα), IL-15La-RLI (a fusion of trout IL-15La and trout sIL-15Rα), IL-15/sIL-15Rα (noncovalent complex of trout IL-15 and trout sIL-15Rα) and IL-15/sIL-15Rα (noncovalent complex of trout IL-15 and trout sIL-15Rα), and they were used in 0.008-to-25 nM concentrations. As negative controls, cells were mock treated (Control). After washing, the same blots were analyzed by Western blot analysis using anti-actin as a loading control.

If using sensitive cells, a weak induction of pSTAT5 could be observed even at 0.008 nM for IL-2, IL-15-RLI and IL-15La-RLI. When comparing IL-15-RLI with IL-15/sIL-15Rα, or IL-15La-RLI with IL-15La/sIL-15Rα, similar sensitivities were observed, which suggests that the fusion proteins were functionally similar to the corresponding noncovalently associated complexes.

(Fig. S5F-1) Comparison of the pSTAT5 inducing effects of IL-15-RLI and IL-15/sIL-15Rα on the $CD8^-$ fraction of intestinal lymphocytes that had been separated by into $CD8^+$ and $CD8^-$ populations by flow sorting. Both cytokine preparations were able to induce detectable pSTAT5 levels from a concentration of 0.2 nM, which suggests their functional similarity.

(Fig. S5F-2) Comparison of the pSTAT5 inducing effects of IL-15La-RLI and IL-15La/sIL-15Rα on the $CD8^-$ fraction of thymocytes that had been separated into $CD8^+$ and $CD8^-$ populations by flow sorting. Both cytokine preparations were able to induce detectable pSTAT5 levels already at a concentration of 0.008 nM which suggests that IL-15La-RLI is at least similarly potent as IL-15La/sIL-15Rα.

(Fig. S5F-3) Analysis of the pSTAT5 inducing effects of IL-15-RLI on the $CD8^-$ fraction of head kidney lymphocytes that had been separated into $CD8^+$ and $CD8^-$ populations by flow sorting. It was found that IL-15-RLI can induce detectable pSTAT5 levels already at a concentration of 0.008 nM.

(Fig. S5F-4) Analysis of the pSTAT5 inducing effects of IL-2 on the $CD8^-$ fraction of thymocytes that had been separated into $CD8^+$ and $CD8^-$ populations by flow sorting. It was found that IL-2 can induce detectable pSTAT5 levels already at a concentration of 0.008 nM.

**Fig. S5F-1.** Similar potencies of trout IL-15-RLI fusion protein and non-covalent association of trout IL-15 with trout sIL-15Rα.

*Western blots*

CD8$^{+}$ Intestinal lymphocytes

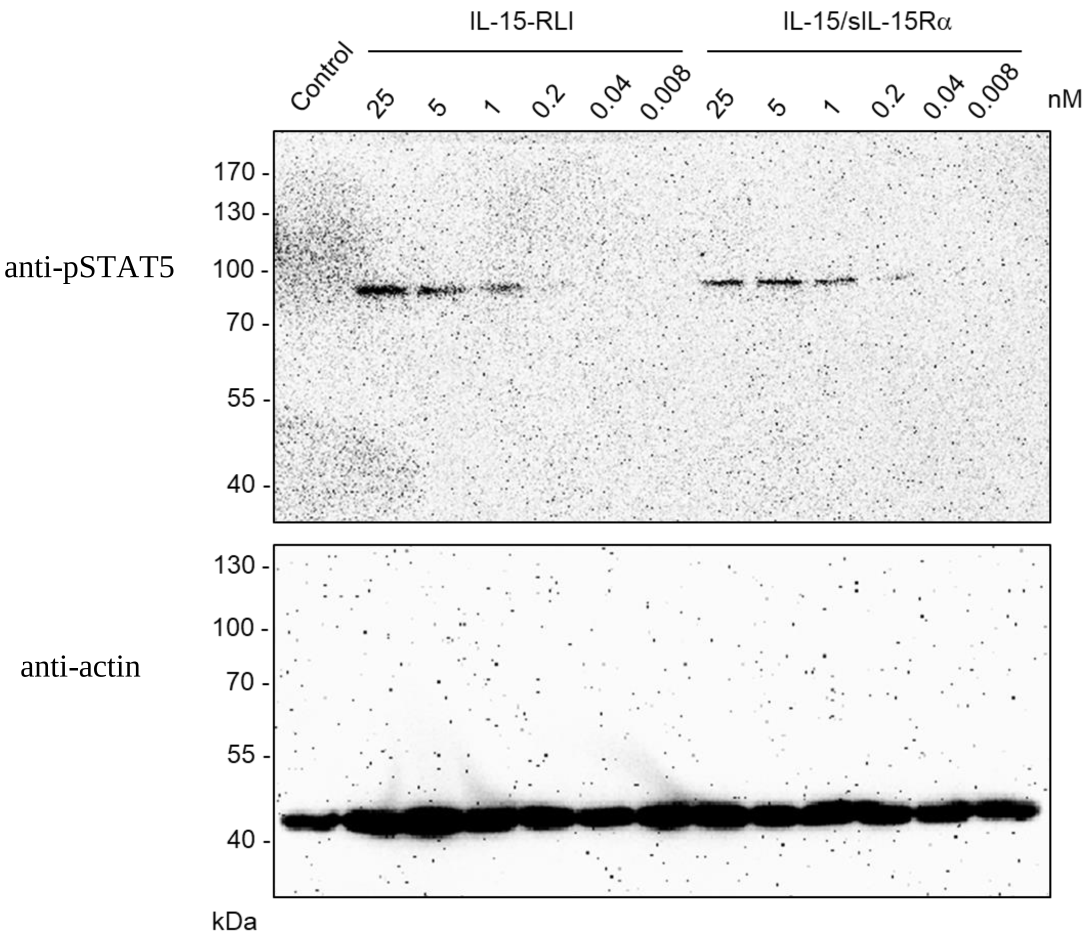

**Fig. S5F-2.** Similar potencies of trout IL-15La-RLI fusion protein and non-covalent associations of trout IL-15La with trout sIL-15Rα.

*Western blots*

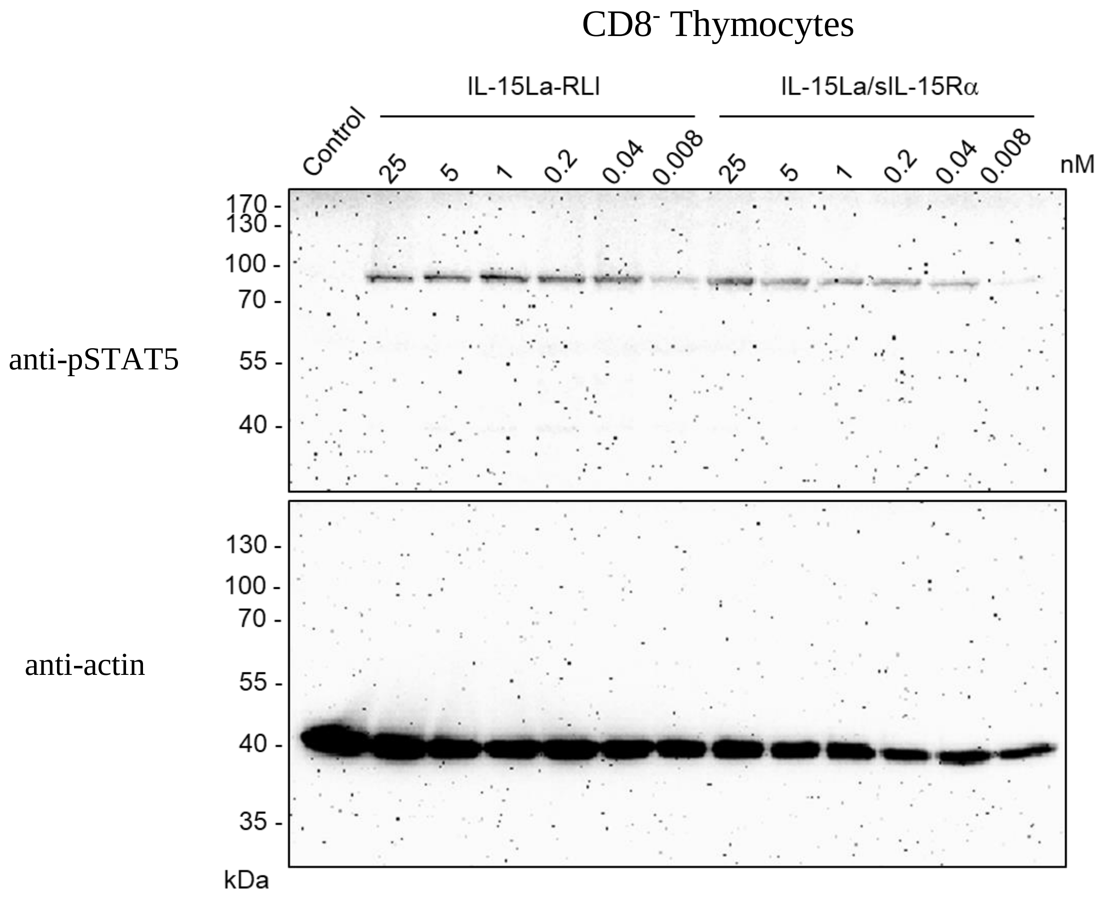

**Fig. S5F-3.** High sensitivity of trout CD8$^-$ head kidney thymocytes to trout IL-15-RLI.

*Western blots*

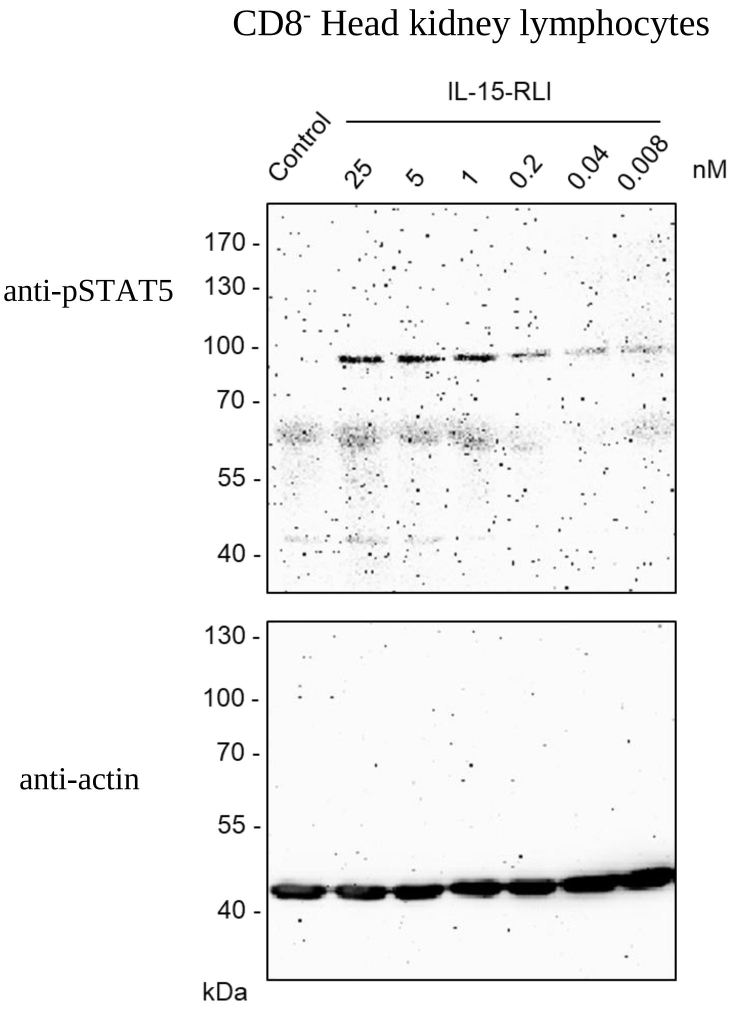

**Fig. S5F-4.** High sensitivity of trout CD8$^-$ thymocytes to trout IL-2.

*Western blots*

CD8$^-$ Thymocytes

anti-pSTAT5

anti-actin

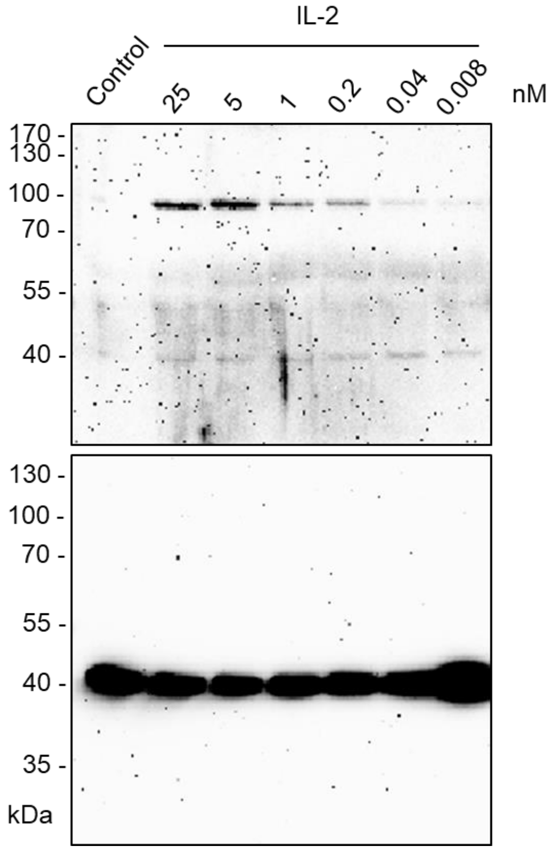

### Supplementary figure 6 (Fig. S6)

Additional information on RT-qPCR experiments

**Table of Contents** **Page**

Fig. S6A: RT-qPCR analysis of immune marker gene expression in trout total splenocytes after incubation with recombinant trout cytokine containing supernatants from transfected HEK293T cells 104

Fig. S6B: RT-qPCR analysis of immune marker gene expression in trout splenocyte subpopulations after stimulation with purified recombinant trout IL-2, IL-15-RLI and IL-15La-RLI produced in insect cells (*information supporting main text Fig. 11*) 107

**Fig. S6A.** RT-qPCR analysis of immune marker gene expression in trout total splenocytes after stimulation with recombinant trout cytokine containing supernatants from transfected HEK293T cells.

The immune marker gene expression profiles exhibited by trout total splenocytes after incubation with trout cytokine containing supernatants from transfected HEK293T cells (Fig. S6A) agree with the results obtained when using purified (fusion-type) trout cytokines produced in insect cells (main text Fig. 10), providing evidence that the observations cannot be explained by cytokine preparation artefacts.

Two independent experiments, using two trout individuals (Trout #1 and Trout #2) and supernatants from independently transfected HEK293T cells were performed. Trout total splenocytes were incubated for 4 h and 12 h at 15 °C with supernatants from HEK293T cells that had been transfected with DNA plasmids encoding trout IL-2, IL-15, IL-15La or IL-15Lb, with (+) or without (-) trout sIL-15Rα. As negative controls, supernatants from HEK293T cells transfected for sIL-15Rα alone (sIL-15Rα) or with an empty control vector (control) were used. After incubation, total RNA was isolated from the splenocytes and reverse transcribed into cDNA, followed by quantitative PCR (qPCR) for expression analysis of *interferon γ (IFNγ), perforin, IL-4/13A, IL-4/13B1, IL-4/13B2* and *EF1A*. Expression levels were equilibrated against *EF1A* expression and the values for the mock-treated controls were set to 1 in each experimental panel.

Results of both experiments, Fig. S6A-1 for Trout #1 and Fig. S6A-2 for Trout #2, show that: (1) IL-15 enhances the expression of type 1 immune genes *IFNγ* and *perforin*, and doesn't need the co-presence of sIL-15Rα to do so; (2) IL-15La and IL-15Lb enhance the expression of type 2 immune genes *IL-4/13A*, *IL-4/13B1* and *IL-4/13B2*, but only do so in the co-presence of sIL-15Rα; (3) IL-2 clearly enhances the expression of *IFNγ* and may slightly enhance the expression of the other investigated genes, and IL-2 doesn't need the co-presence of sIL-15Rα to do so.

*Fig. S6A-1.*

***Relative expression levels of IFNγ, perforin, IL-4/13A, IL-4/13B1 and IL-4/13B2 in total splenocytes of Trout #1 after incubation for 4 h and 12 h with recombinant trout cytokine containing supernatants from transfected HEK293T cells***

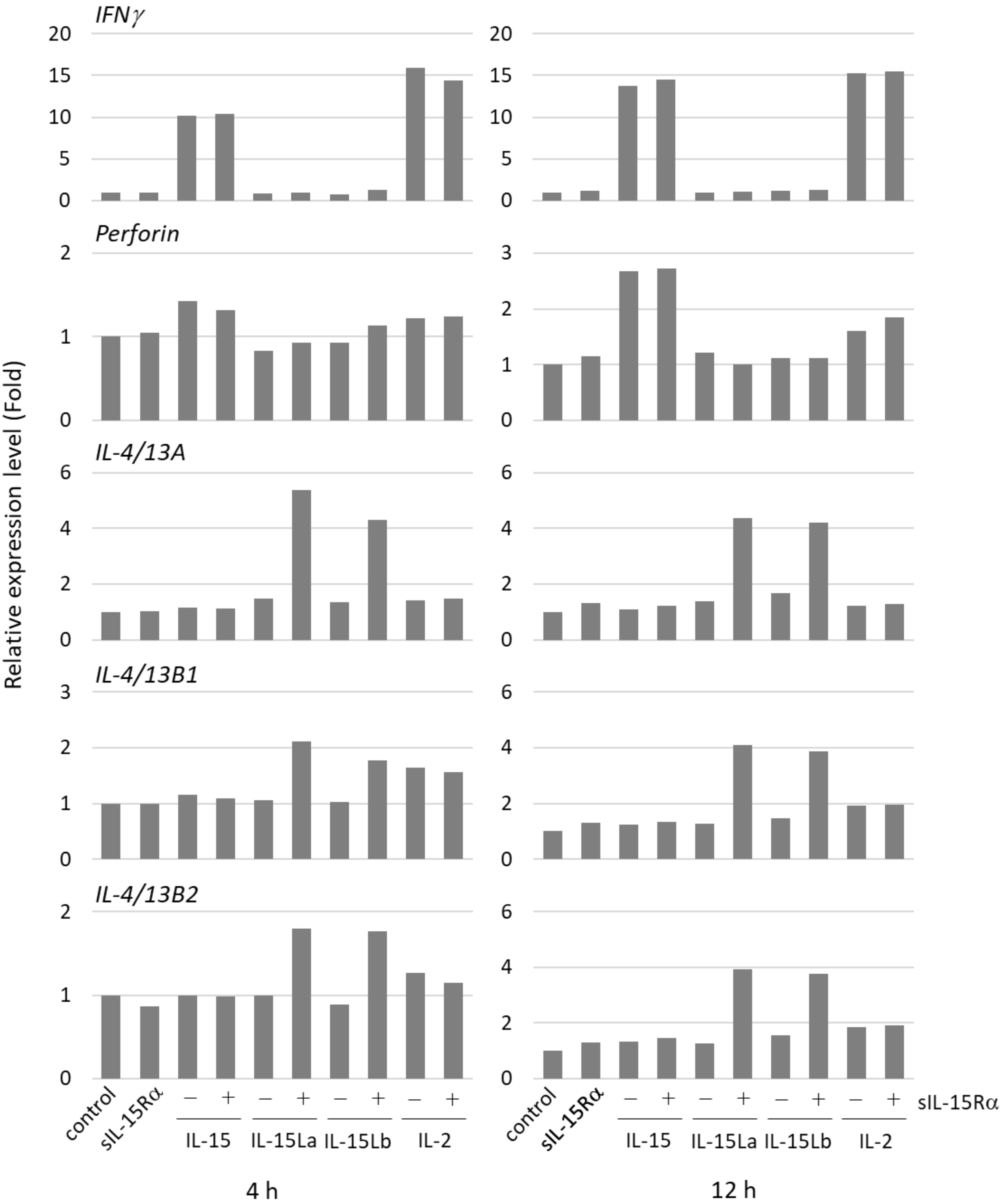

*Fig. S6A-2.*

*Relative expression levels of IFNγ, perforin, IL-4/13A, IL-4/13B1 and IL-4/13B2 in total splenocytes of Trout #2 after incubation for 4 h and 12 h with recombinant trout cytokine containing supernatants from transfected HEK293T cells*

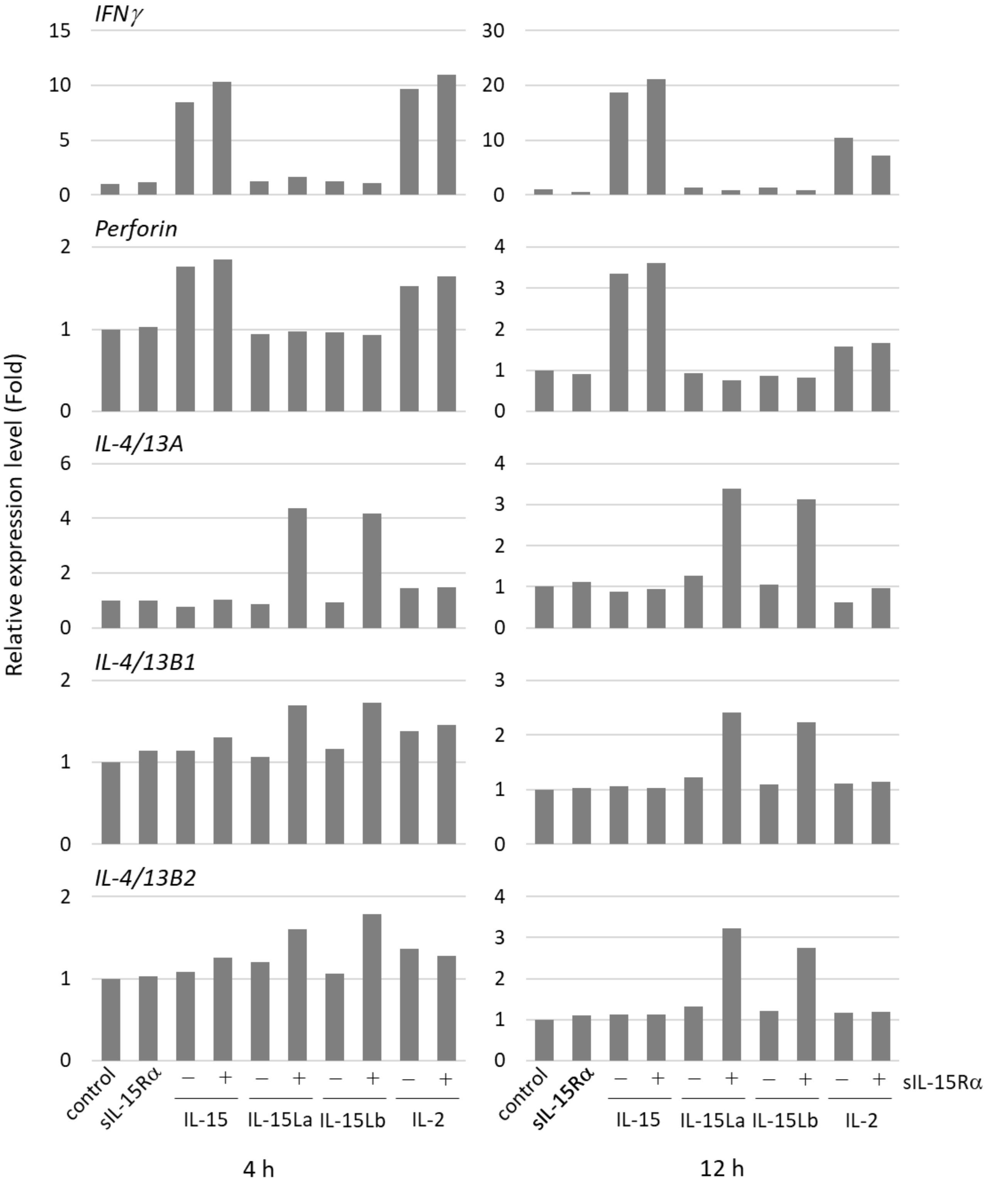

**Fig. S6B.** RT-qPCR analysis of immune marker gene expression in trout splenocyte subpopulations after stimulation with purified recombinant trout IL-2, IL-15-RLI and IL-15La-RLI produced in insect cells (*information supporting main text Fig. 11*).

In comparison to main text Fig. 11, the graphs in Figs. S6B (b) and (c) provide additional information on the gene expressions in $CD4^+$, $CD8^+$ and $IgM^+$ cells. The graphs in Fig. S6B(b) are identical to those in main text Fig. 11, except for being shown at a larger scale, while the graphs in Fig. S6B(c) are based on the same dataset but using a different approach of observations with high Ct values.

$CD4^+$, $CD8^+$, $IgM^+$ and $CD4^-CD8^-IgM^-$ (triple negative; TN) lymphocyte populations were isolated from trout spleen using flow sorting (for an example see Fig. S3) and stimulated for 12 h at 15 °C with purified recombinant trout cytokines IL-2, IL-15-RLI and IL-15La-RLI (produced in insect cells; see Fig. S4) at 0.2 and 5 nM. After incubation, total RNA was isolated and reverse transcribed into cDNA, followed by quantitative PCR (qPCR) for expression analysis of *IFNγ*, *perforin*, *IL-4/13A*, *IL-4/13B1*, *IL-4/13B2* and *EF1A*. The determined Ct values are shown in a table [Fig. S6B(a)], with Ct values $\geq$35 highlighted by pink color because the reliability of those values may be considered questionable.

Using the data shown in the table (a), expression levels were equilibrated against *EF1A* expression and the values for the mock-treated TN control ("0") were set to 1 in each experimental panel. In the graphs [main text Fig. 11 and Figs. S6B (b) and (c)], the average values of four experiments are shown together with error bars representing SD. The differences between the graphs shown in (b) and (c) are based on different approaches of the Ct $\geq$35 values. For the calculations resulting in the main text Fig. 11 and Fig. S6B(b) graphs, the Ct values were used as indicated in table Fig S6B(a), with "No Ct" interpreted as zero expression. For the calculations resulting in the Fig. S6B(c) graphs also the Ct values indicated in table Fig. S6B(a) were used, but now all Ct values $\geq$35 (and "No Ct" observations) were equalized as 35.

We are showing Figs. S6B (b) and (c) in order to provide complete information. However, overall, the reliability and functional relevance of the quantitative comparisons between cases of very low expression levels can be questioned, and many of the data shown in Figs. S6B (b) and (c) should not be overinterpreted (which is also why we refrained from including statistical analysis in these two figures). The important finding of our RT-qPCR analyses of trout total and subpopulation splenocytes (Figs. S6A and S6B, and main text Figs. 10 and 11) is that in splenocytes from healthy trout most *IFNγ*, *IL-4/13A*, *IL-4/13B1* and *IL-4/13B2* is expressed by $CD4^-CD8^-IgM^-$ lymphocytes, and that expression levels of these cytokine gene transcripts are differentially affected by IL-15(+sIL-15Rα) versus IL-15L+sIL-15Rα.

*Fig. S6B.*

(a) *Table with the Ct values used for main text Fig. 11*

| | | CD8$^+$ | | | | | | | CD4$^+$ | | | | | | |
|---|---|---|---|---|---|---|---|---|---|---|---|---|---|---|---|
| | | | IL-15-RLI | | IL-15La-RLI | | IL-2 | | | IL-15-RLI | | IL-15La-RLI | | IL-2 | |
| | | Empty | 0.2 nM | 5 nM | 0.2 nM | 5 nM | 0.2 nM | 5 nM | Empty | 0.2 nM | 5 nM | 0.2 nM | 5 nM | 0.2 nM | 5 nM |
| #1 | EF1A | 23.84 | 22.88 | 23.43 | 23.7 | 23.33 | 23.39 | 23.33 | 20.37 | 20.94 | 20.51 | 21.05 | 20.76 | 20.92 | 20.83 |
| | IFNg1/2 | 37.88 | 34.49 | 32.95 | 37.64 | 34.51 | 33.34 | 33.76 | 35.46 | 33.64 | 34.51 | 36.9 | 34.53 | 34.41 | 32.71 |
| | Perforin | 28.66 | 27.55 | 28.02 | 28.74 | 28.39 | 28.04 | 27.93 | 28.18 | 28.73 | 28.31 | 28.79 | 28.6 | 28.85 | 28.74 |
| | IL-4/13A | 34.6 | 34.71 | 34.71 | 34.83 | 35.79 | 34.56 | 34.64 | 32.13 | 32.92 | 32.22 | 32.71 | 32.36 | 32.92 | 32.74 |
| | IL-4/13B1 | 37.35 | 38.62 | 39.12 | 35.09 | No Ct | 37.42 | 30.24 | 28.32 | 29.61 | 28.47 | 28.63 | 30.6 | 28.69 | 27.93 |
| | IL-4/13B2 | No Ct | No Ct | No Ct | No Ct | No Ct | No Ct | 33.92 | 33.75 | 34.36 | No Ct | 34.35 | 32.78 | 32.54 | 31.94 |
| #2 | EF1A | 22.57 | 22.72 | 22.21 | 23.17 | 22.98 | 22.92 | 22.89 | 20.32 | 20.21 | 20.47 | 20.38 | 20.47 | 20.85 | 20.46 |
| | IFNg1/2 | 37 | 34.12 | 34.28 | 35.43 | 34.49 | 33.42 | 33.7 | 34.6 | 34.21 | 32.98 | 34.31 | 34.95 | 36.2 | 33.75 |
| | Perforin | 28.62 | 28.41 | 28.01 | 29.03 | 28.94 | 28.72 | 28.6 | 29.41 | 28.86 | 29.48 | 29.47 | 29.39 | 29.54 | 29.46 |
| | IL-4/13A | 32.88 | 32.76 | 31.77 | 32.93 | 32.7 | 33.25 | 32.93 | 29.55 | 29.71 | 30.29 | 29.72 | 29.81 | 30.01 | 30.02 |
| | IL-4/13B1 | 36.21 | 31.6 | No Ct | 39.31 | 37.33 | 33.65 | No Ct | 29.86 | 28.73 | 30.92 | 29.31 | 28.52 | 30.53 | 31.16 |
| | IL-4/13B2 | No Ct | No Ct | No Ct | No Ct | No Ct | No Ct | No Ct | 34.22 | 33.65 | 33.74 | 32.85 | 31.86 | 34.44 | 32.48 |
| #3 | EF1A | 22.82 | 23.97 | 23.74 | 23.39 | 23.83 | 23.72 | 22.96 | 21.99 | 21.53 | 21.24 | 21.51 | 21.76 | 21.48 | 21.54 |
| | IFNg1/2 | 34.49 | 35.35 | 34.64 | 34.61 | No Ct | 35.67 | 33.08 | 35.01 | 34.67 | 39.47 | 34.32 | 34.34 | 33.69 | 34.12 |
| | Perforin | 30.09 | 30.43 | 30.16 | 30.43 | 30.71 | 30.37 | 29.6 | 31.58 | 31.09 | 30.79 | 30.92 | 31.06 | 31.44 | 31.18 |
| | IL-4/13A | 35.03 | 35.88 | 34.89 | 34.68 | 36.35 | 39.48 | 35.09 | 33.92 | 33.18 | 33.02 | 33.56 | 33.63 | 34.37 | 33.92 |
| | IL-4/13B1 | 38.07 | No Ct | 34.33 | 31.31 | 37.23 | 35.81 | No Ct | No Ct | 31.37 | 33.85 | 33.42 | 34.24 | 29.8 | 31.17 |
| | IL-4/13B2 | No Ct | No Ct | No Ct | 35.21 | No Ct | No Ct | No Ct | 35.08 | 35.44 | 33.83 | No Ct | 35.18 | 34.17 | 34.67 |
| #4 | EF1A | 23.92 | 23.83 | 24.42 | 23.93 | 24.19 | 24.49 | 23.98 | 21.71 | 22.16 | 22.01 | 22.62 | 21.92 | 22.62 | 21.66 |
| | IFNg1/2 | No Ct | 33.69 | 35.42 | 35.27 | 35.08 | 35.37 | 32.88 | 38.75 | 35.82 | 35.74 | No Ct | 33.49 | 34.31 | 35.4 |
| | PFN1 | 29.85 | 29.78 | 30.22 | 29.74 | 30.23 | 30.17 | 29.89 | 30.95 | 31.69 | 31.14 | 31.51 | 31.39 | 31.43 | 30.87 |
| | IL-4/13A | 33.88 | 33.25 | 33.93 | 33.3 | 33.98 | 35.41 | 34.2 | 31.97 | 31.94 | 31.95 | 32.11 | 31.81 | 31.73 | 31.37 |
| | IL-4/13B1 | 37.98 | 32.48 | 38.04 | 33.58 | 38.32 | 39.22 | 39.16 | 33 | 34.03 | 31 | 30.37 | 31.75 | 39.45 | 29.74 |
| | IL-4/13B2 | No Ct | No Ct | No Ct | 39.22 | No Ct | No Ct | No Ct | 36.01 | 36.93 | 34.13 | 35.89 | 37.95 | No Ct | 36.95 |

| | | IgM$^+$ | | | | | | | TN | | | | | | |
|---|---|---|---|---|---|---|---|---|---|---|---|---|---|---|---|
| | | | IL-15-RLI | | IL-15La-RLI | | IL-2 | | | IL-15-RLI | | IL-15La-RLI | | IL-2 | |
| | | Empty | 0.2 nM | 5 nM | 0.2 nM | 5 nM | 0.2 nM | 5 nM | Empty | 0.2 nM | 5 nM | 0.2 nM | 5 nM | 0.2 nM | 5 nM |
| #1 | EF1A | 21.38 | 21.82 | 21.46 | 21.47 | 21.78 | 22.27 | 21.71 | 21.47 | 22.06 | 22.01 | 21.78 | 21.89 | 23.01 | 21.86 |
| | IFNg1/2 | 36.7 | No Ct | No Ct | 38.03 | 38.35 | 39.69 | No Ct | 30.61 | 29.21 | 28.69 | 30.96 | 30.81 | 31.6 | 29.77 |
| | Perforin | 34.17 | 33.9 | 33.19 | 38.19 | No Ct | 35.96 | 34.33 | 27.45 | 26.7 | 26.77 | 27.74 | 28.17 | 28.72 | 27.16 |
| | IL-4/13A | 35.01 | 35.5 | 34.67 | 34.59 | 34.77 | 36.66 | 35.01 | 28.08 | 28.19 | 27.96 | 25.6 | 26.26 | 29.33 | 28.04 |
| | IL-4/13B1 | 32.43 | No Ct | No Ct | No Ct | 30.63 | 37.65 | 39.6 | 20.2 | 20.35 | 20.33 | 18.96 | 19.44 | 21.26 | 19.85 |
| | IL-4/13B2 | No Ct | No Ct | No Ct | No Ct | 34.66 | No Ct | No Ct | 24.63 | 25.12 | 24.95 | 23.59 | 23.45 | 25.27 | 24.64 |
| #2 | EF1A | 21.85 | 20.5 | 21.61 | 21.6 | 21.52 | 21.91 | 20.99 | 21.36 | 20.91 | 21.08 | 21.85 | 21.38 | 22.26 | 21.72 |
| | IFNg1/2 | No Ct | No Ct | No Ct | No Ct | 37.78 | No Ct | No Ct | 31.1 | 28.86 | 29.35 | 31.12 | 31.44 | 30.69 | 29.76 |
| | Perforin | 37.94 | 34 | 33.65 | 34.51 | 34.92 | 35.56 | 34.21 | 28.83 | 27.29 | 27.42 | 28.58 | 28.69 | 28.78 | 27.94 |
| | IL-4/13A | 32.99 | 31.97 | 32.1 | 33.53 | 32.05 | 32.09 | 32.67 | 27.26 | 26.14 | 26.59 | 24.17 | 24.1 | 27.11 | 26.4 |
| | IL-4/13B1 | 30.01 | No Ct | 36.61 | No Ct | No Ct | No Ct | No Ct | 22.02 | 21.46 | 21.75 | 20.87 | 20.96 | 21.95 | 21.25 |
| | IL-4/13B2 | No Ct | 38.93 | No Ct | No Ct | No Ct | No Ct | No Ct | 25.45 | 24.74 | 25.08 | 24.7 | 24.51 | 25.91 | 24.97 |
| #3 | EF1A | 19.35 | 20.49 | 19.63 | 19.91 | 19.92 | 21.42 | 21.05 | 21.08 | 20.81 | 20.89 | 21.04 | 20.92 | 21.4 | 21.75 |
| | IFNg1/2 | 38.55 | 36.8 | 38.39 | 36.37 | 35.85 | 37.47 | 38.33 | 30.36 | 28.41 | 28.67 | 31.04 | 31.4 | 31.12 | 30.62 |
| | Perforin | 33.76 | 33.09 | 33.91 | 33.76 | 33.43 | 33.88 | 33.4 | 29.72 | 28.24 | 27.83 | 29.98 | 29.81 | 29.76 | 29.51 |
| | IL-4/13A | 34.94 | 34.81 | 34.63 | 36.11 | 35.23 | 35.36 | 34.18 | 31.46 | 30.32 | 30.27 | 27.36 | 27.1 | 31.95 | 32.03 |
| | IL-4/13B1 | 31.6 | 39.87 | No Ct | 34.3 | No Ct | 33.32 | No Ct | 23.32 | 22.6 | 22.5 | 21.6 | 21.41 | 23.14 | 22.54 |
| | IL-4/13B2 | No Ct | No Ct | No Ct | No Ct | No Ct | No Ct | No Ct | 27.03 | 26.4 | 26.39 | 25.65 | 25.56 | 27.35 | 27.29 |
| #4 | EF1A | 21.62 | 20.73 | 22.06 | 21.92 | 21.53 | 21.52 | 21.47 | 22.74 | 21.86 | 21.84 | 22.43 | 22.17 | 22.32 | 22.14 |
| | IFNg1/2 | 37.5 | No Ct | No Ct | No Ct | 38.93 | No Ct | No Ct | 32.92 | 31.31 | 31.05 | 32.27 | 32.7 | 31.63 | 31.16 |
| | PFN1 | 33.75 | 32.34 | 32.95 | 36.88 | 33.79 | 36.11 | 34.83 | 31.53 | 28.85 | 28.51 | 30.7 | 30.87 | 30.31 | 30.01 |
| | IL-4/13A | 32.62 | 32.91 | 32.83 | 33.46 | 33.51 | 32.48 | 33.16 | 33.27 | 30.82 | 30.79 | 27.74 | 27.44 | 32.04 | 31.83 |
| | IL-4/13B1 | 33.55 | 39.17 | No Ct | 37.95 | 35.92 | 29.13 | 39.86 | 23.92 | 23.02 | 22.81 | 22.2 | 22 | 23.46 | 22.88 |
| | IL-4/13B2 | No Ct | No Ct | No Ct | No Ct | No Ct | No Ct | No Ct | 29.56 | 28.89 | 28.88 | 28.1 | 28.29 | 29.57 | 28.85 |

*(Fig. S6B continued)*

(b) *Relative expression levels of IFNγ, perforin, IL-4/13A, IL-4/13B1 and IL-4/13B2 in CD4⁺, CD8⁺ and IgM⁺ trout spleen morphological lymphocytes after incubation for 12 h with purified recombinant trout cytokines (produced in insect cells) IL-2, IL-15-RLI and IL-15La-RLI at 0.2 and 5 nM. Analysis as done for main text Fig. 11 but shown at a larger scale.*

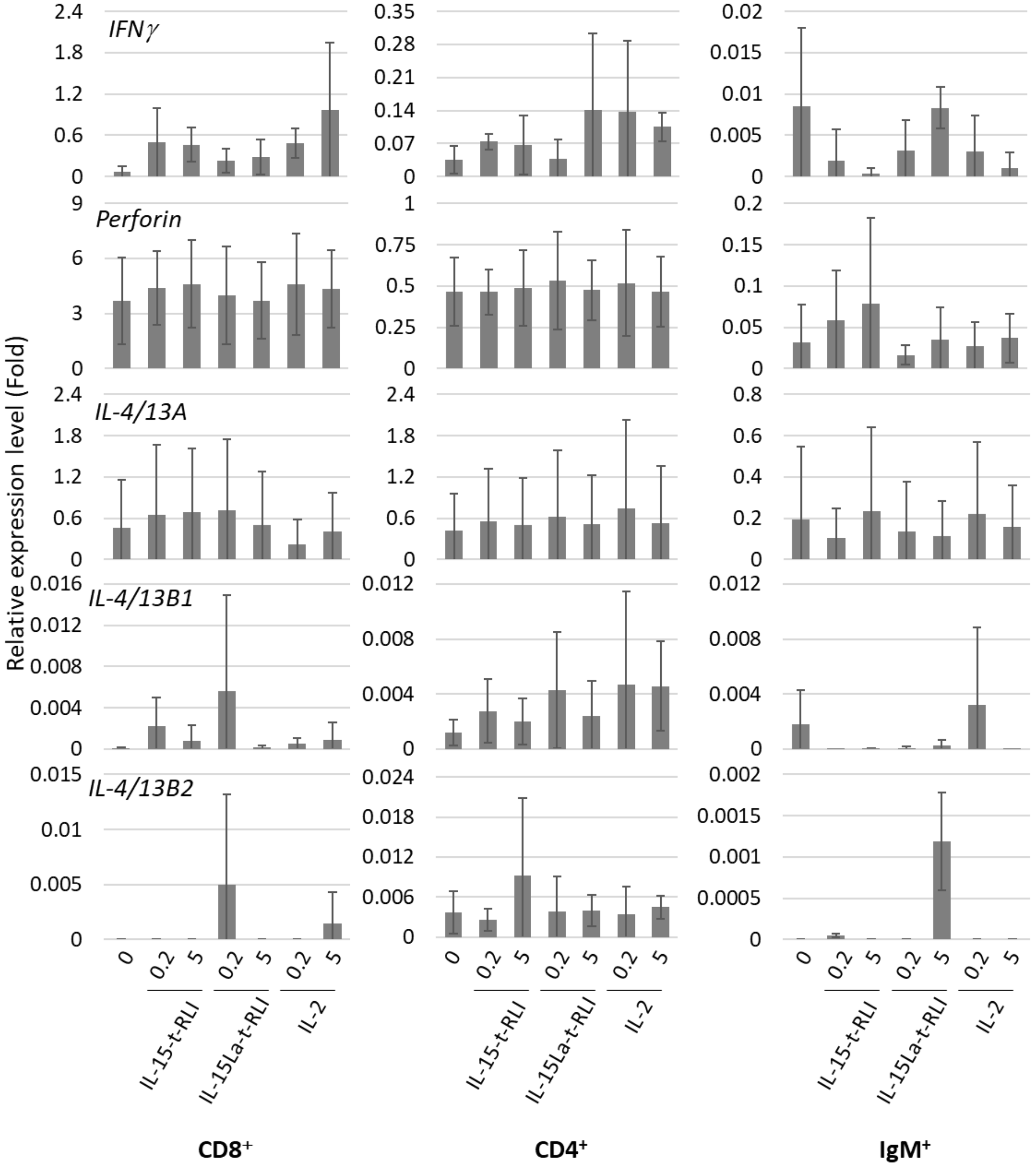

*(Fig. S6B continued)*

(c) *Relative expression levels of IFNγ, perforin, IL-4/13A, IL-4/13B1 and IL-4/13B2 in $CD4^+$, $CD8^+$ and $IgM^+$ trout spleen morphological lymphocytes after incubation for 12 h with purified recombinant trout cytokines (produced in insect cells) IL-2, IL-15-RLI and IL-15La-RLI at 0.2 and 5 nM. Analysis as done for main text Fig. 11 except for considering all Ct* $\geq$*35 values as 35.*

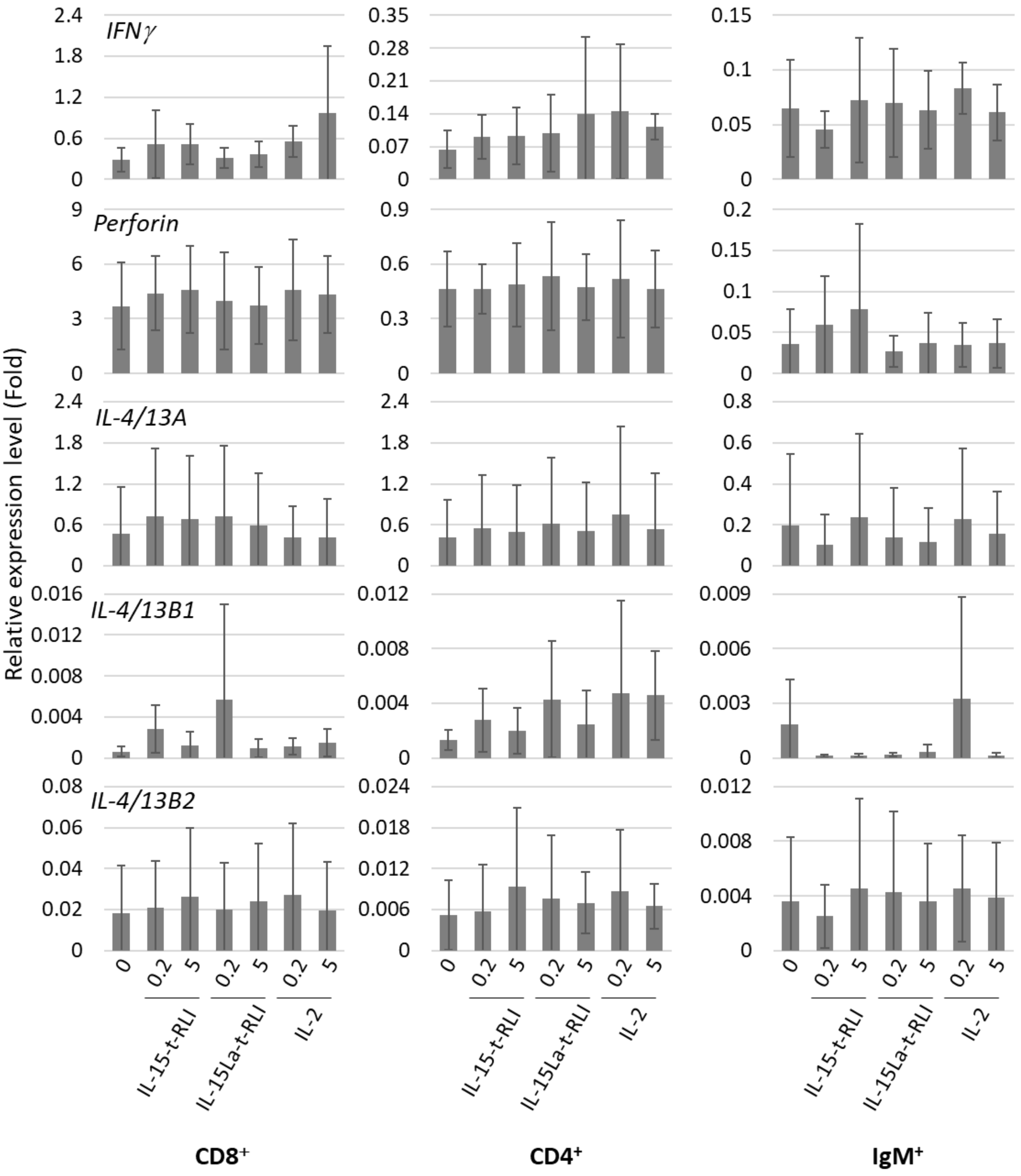

#### Supplementary Table 1 (Table S1).

Percentages of $FLAG^{+}$ cells among live HEK293T cells after their transfection for FLAG-tagged trout or bovine IL-2/15/15L-family cytokines with or without co-transfection for trout or bovine IL-15Rα or bovine IL-2Rα. The values were determined by FACS as shown in main Text Fig. 5, and include the data shown in Fig. 5 plus those of independent experiment repeats. Light orange color is used for highlighting values that were deemed indicative for cytokine to alpha chain receptor binding.

**Trout cytokines**

| IL-2 only | IL-2 + Trout IL-15Rα |
|---|---|
| 11.04 | 73.97 |
| 0.47 | 31.28 |
| 0.45 | 17.90 |
| 6.70 | 64.92 |

| IL-2 only | IL-2 + Bovine IL-2Rα | IL-2 only | IL-2 + Bovine IL-15Rα |
|---|---|---|---|
| 11.04 | 3.46 | 11.04 | 87.36 |
| 0.58 | 0.95 | 0.55 | 20.37 |
| 0.45 | 0.92 | 0.19 | 22.79 |

| IL-15 only | IL-15 + Trout IL-15Rα |
|---|---|
| 1.44 | 76.24 |
| 0.74 | 35.07 |
| 0.80 | 20.63 |
| 1.85 | 72.38 |

| IL-15 only | IL-15 + Bovine IL-2Rα | IL-15 only | IL-15 + Bovine IL-15Rα |
|---|---|---|---|
| 1.44 | 1.61 | 1.44 | 95.14 |
| 0.73 | 1.29 | 1.06 | 31.90 |
| 0.51 | 0.82 | 0.60 | 37.35 |

| IL-15La only | IL-15La + Trout IL-15Rα |
|---|---|
| 12.29 | 20.38 |
| 3.69 | 28.68 |
| 2.83 | 17.46 |
| 11.06 | 61.56 |

| IL-15La only | IL-15La + Bovine IL-2Rα | IL-15La only | IL-15La + Bovine IL-15Rα |
|---|---|---|---|
| 12.29 | 8.08 | 12.29 | 86.46 |
| 3.95 | 4.79 | 3.07 | 32.43 |
| 2.29 | 3.08 | 4.30 | 37.00 |

| IL-15Lb only | IL-15Lb + Trout IL-15Rα | IL-15Lb only | IL-15Lb + Bovine IL-2Rα | IL-15Lb only | IL-15Lb + Bovine IL-15Rα |
|---|---|---|---|---|---|
| 0.03 | 10.73 | 0.03 | 0.24 | 0.03 | 29.28 |
| 0.52 | 3.18 | 0.52 | 0.21 | 0.52 | 10.45 |
| 0.72 | 4.27 | 0.72 | 0.76 | 0.72 | 15.61 |

**Bovine cytokines**

| IL-2 only | IL-2 + Trout IL-15Rα | IL-2 only | IL-2 + Bovine IL-2Rα | IL-2 only | IL-2 + Bovine IL-15Rα |
|---|---|---|---|---|---|
| 0.35 | 0.18 | 0.35 | 27.70 | 0.35 | 0.43 |
| 0.51 | 0.45 | 0.39 | 42.48 | 0.39 | 0.35 |
| 0.64 | 1.67 | 0.48 | 23.80 | 0.26 | 0.49 |

| IL-15 only | IL-15 + Trout IL-15Rα | IL-15 only | IL-15 + Bovine IL-2Rα | IL-15 only | IL-15 + Bovine IL-15Rα |
|---|---|---|---|---|---|
| 54.46 | 68.94 | 54.46 | 51.34 | 54.46 | 91.19 |
| 15.41 | 29.22 | 15.41 | 8.24 | 15.41 | 44.42 |
| 17.88 | 34.62 | 17.88 | 10.37 | 17.88 | 51.70 |

| IL-15L only | IL-15L + Trout IL-15Rα | IL-15L only | IL-15L + Bovine IL-2Rα | IL-15L only | IL-15L + Bovine IL-15Rα |
|---|---|---|---|---|---|
| 1.89 | 22.71 | 1.89 | 2.29 | 1.89 | 68.90 |
| 1.72 | 6.17 | 2.94 | 3.88 | 2.94 | 45.82 |
| 2.21 | 5.86 | 2.17 | 2.72 | 5.73 | 43.82 |
| 0.16 | 12.77 | 0.16 | 0.44 | 0.16 | 35.40 |
